# Comprehensive identification, isolation, and culture of human breast cell types

**DOI:** 10.1101/2022.09.20.508726

**Authors:** Kate Thi, Katelyn Del Toro, Yamhilette Licon-Munoz, Rosalyn W. Sayaman, William C. Hines

**Affiliations:** Department of Biochemistry and Molecular Biology; University of New Mexico School of Medicine, 1 University of New Mexico MSC08 4670, Albuquerque NM 87131, USA; Life Sciences Division, Lawrence Berkeley National Laboratory; Mailstop 977R225A, 1 Cyclotron Road; Berkeley, CA 94720, USA.; Department of Laboratory Medicine, Helen Diller Family Comprehensive Cancer Center, University of California, San Francisco, San Francisco, CA 94143, USA.

**Keywords:** Breast, flow cytometry, FACS, primary culture models, RNA-sequencing (RNA-seq), human mammary epithelial cells (HMEC), luminal epithelial cells, myoepithelial, vascular smooth muscle, pericytes, endothelial, lymphatic, fibroblasts, adipocyte-derived mesenchymal cells, adipocytes, leukocytes, erythrocytes, comparative analysis

## Abstract

Tissues are formed and shaped by cells of many different types and are orchestrated through countless interactions among the cells—and the myriad of molecules they synthesize. Deciphering a tissue’s biological complexity thus requires studying it at cell-level resolution, where molecular and biochemical features of different cell types can be explored and thoroughly dissected. Unfortunately, the lack of comprehensive methods to identify, isolate, and culture each cell type from many tissues has impeded progress. Here, we present a method for the breadth of cell types composing the human breast. Our goal has long been to understand the essence of each of these different breast cell types, that is, to reveal the underlying biology explaining their intrinsic features, the consequences of interactions, and their contributions to the tissue as a whole. This biological exploration has required cell purification, deep-RNA sequencing—and a thorough dissection of the genes and pathways defining each cell type, which we present in an adjoining article. Here, we present an exhaustive cellular dissection of the human breast, where we explore its cellular composition and histological organization. Moreover, we introduce a novel fluorescence-activated cell sorting (FACS) antibody panel and rigorous gating strategy capable of isolating each of the twelve major breast cell types to purity. Finally, we describe the creation of primary cell models from nearly every one of these breast cell types—some being the first of their kind— and submit these as critical tools for studying the dynamic cellular interactions within breast tissues and tumors. Together, this body of work and derived resources deliver a unique perspective of the breast, revealing insights into its cellular, molecular, and biochemical composition.

## Introduction

Modern advances in single-cell technologies and cytometry permit high dimensional analyses of numerous simultaneous markers and are capable and well suited for interrogating complex cell populations^1, 2^. These kinds of investigations have led to a more considerable appreciation of the types and properties of cells in complex tissue systems (e.g., the hematopoietic system), have given us insight into normal cell development and physiology, and have been crucial to developing a broader understanding of the malignant state^3, 4^. While emerging technologies like single-cell RNA-sequencing and digital spatial profiling are helping unravel the cellular heterogeneity in tissues, complementary deep RNA-sequencing of purified populations provides a broader and more thorough assessment of expressed genes and proteins. Unfortunately, the full range of capabilities offered by coupling complex multicolor FACS strategies and deep RNA-sequencing has yet to be applied *fully* to many solid tumor and tissue types, including the breast. Obstacles include the relatively limited availability and amounts of freshly isolated specimens, technicalities associated with producing single-cell suspensions from solid tissues^5^, the sparsity of specific cell types causing them to be easily overlooked, as well as the immense amount of sort-time and labor required to collect enough rare cells for deep analysis. Progress in the field is further restricted by the expertise and technical knowledge needed to design and optimize intricate antibody panels^6, 7^, properly operate a sophisticated flow sorter, and interpret the resulting complex multi-parameter datasets—information that can be dangerously misleading if not handled carefully^6–9^. These are indeed time-consuming and challenging endeavors^10^.

Nonetheless, the most notable impediment to analyzing breast tissues by flow cytometry is the absence of an accepted set of markers and strategy for purifying the entire collection of cell types constituting this tissue. This is true for adult stem cells and most other cells in the breast, and many technical caveats and perplexing and contradictory claims persist in the literature^5, 11–13^. These challenges and enduring questions have long contributed to controversies regarding the nature, relationship, and function of different cell types in breast tissues^5, 14, 15^, collectively placing the field—and our understanding of these different cell types—into a state of uncertainty^16^.

Resolving the types and properties of all breast cells is vital to understanding tissue biology and the emergence of diseases such as cancer. Achieving such a comprehensive understanding will initially rely on analyzing each cell type’s molecular profile and functionally characterizing these cells. Thus, inclusive and reliable methods are needed to identify, purify, and culture each breast cell lineage. To establish such protocols and cell models, we have revisited our understanding of the breast’s cellular composition and organization through exhaustive tissue immunostaining and have matched results with cytometric analyses performed in parallel. We have used this knowledge to develop a comprehensive FACS antibody panel and gating strategy that permits objective identification and isolation of every major cell type in the breast. We isolated twelve cell populations that include two distinct luminal epithelial fractions, myoepithelial cells, adipocytes, leukocytes (counted here as a single population), pericytes, vascular smooth muscle cells, erythrocytes, adipose-derived mesenchymal stem cells (fibroblasts), lymphatic and vascular endothelial cells, and a newly identified epithelial cell type. This method considerably expands the number and types of cells isolated and cultured from the breast from the two to three commonly reported. Development and validation of this FACS method occurred over many years; Here, we present critical discoveries that guided its evolution—including how newly isolated cell fractions were identified and validated— and how critical pitfalls were encountered and overcome. Furthermore, we demonstrate how nearly every cell type can be seeded into culture and expanded for future analyses.

Most of the breast literature to date has dealt with either the epithelium or stroma, with a focus primarily confined to a few cell types within each of these compartments, such as luminal and myoepithelial cells—or fibroblasts and adipocytes (Figure 1a,b). Widespread adoption of primary cell models has also been limited—essentially to myoepithelial cells, fibroblasts, or mixed unsorted human mammary epithelial cultures (HMECs), which are commercially available and ultimately self-select for faster-growing myoepithelial cells (Figure 1 –figure supplement 1). Of course, it is understood that the tissue contains additional cell types, for example, cells that compose blood vessels, immune cells, and possibly others (Figure 2; Figure 3). Knowing where these different cell types reside in the breast is not always abundantly clear, and markers and methods to objectively classify and prospectively isolate each cell type from the breast have proved difficult to establish. This has left the field without precise methods to purify most cell types from breast tissues—and even less from tumors. Epithelial cells have received the bulk of past attention, with the most commonly reported marker-combination for discriminating these being ‘epithelial cell adhesion molecule’ (EpCAM) and alpha-6 integrin (CD49f)^17–25^. In our hands, this marker pair resolved four different cell populations from normal reduction mammoplasty breast tissues (Figure 4). However, reports using these markers have varied on the number, types, and functional capabilities of the cell populations identified (Ibid). Whether these disparities reflect technical differences or underlying biological diversity was unclear at the outset of this study, but the discord has generated uncertainty. To expand upon these current methods and make deductive inferences about each of the different cell types in the breast, we needed a comprehensive FACS strategy capable of objectively and unambiguously resolving each cell type from the tissue.

**Figure 1.**
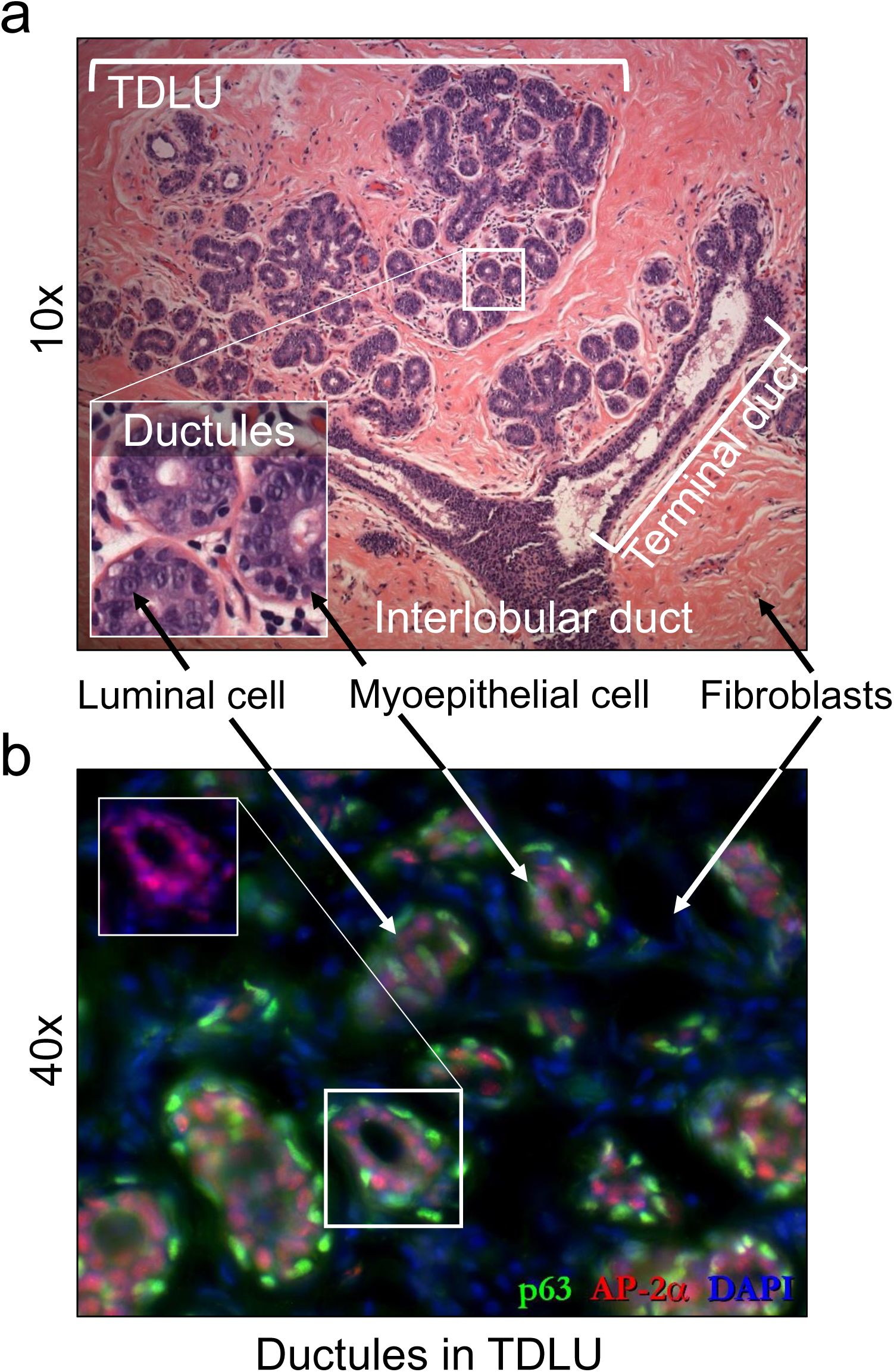
Normal Breast Histology. **(a)** H&E staining of a normal breast reveals the breast’s compound alveolar glandular architecture (10x objective; 34 y/o female). The terminal ductal lobular units (TDLUs)— also referred to as lobules— along with their associated terminal ducts and an interlobular duct are embedded in an eosin-staining collagenous stroma. The two major epithelial cell types (myoepithelial and luminal epithelial cells) are discernible within the magnified tube-shaped ductules that compose the TDLUs (magnified inset). **(b)** Immunostaining for the transcription factor AP-2α (red staining) reveals nuclei of all epithelial cell types, whereas the superimposed p63 staining (green) is limited to the basally located myoepithelial cells (40x objective, 35 y/o female). (b, inset) Removal of the p63 overlay reveals myoepithelial AP-2α expression.

**Figure 2.**
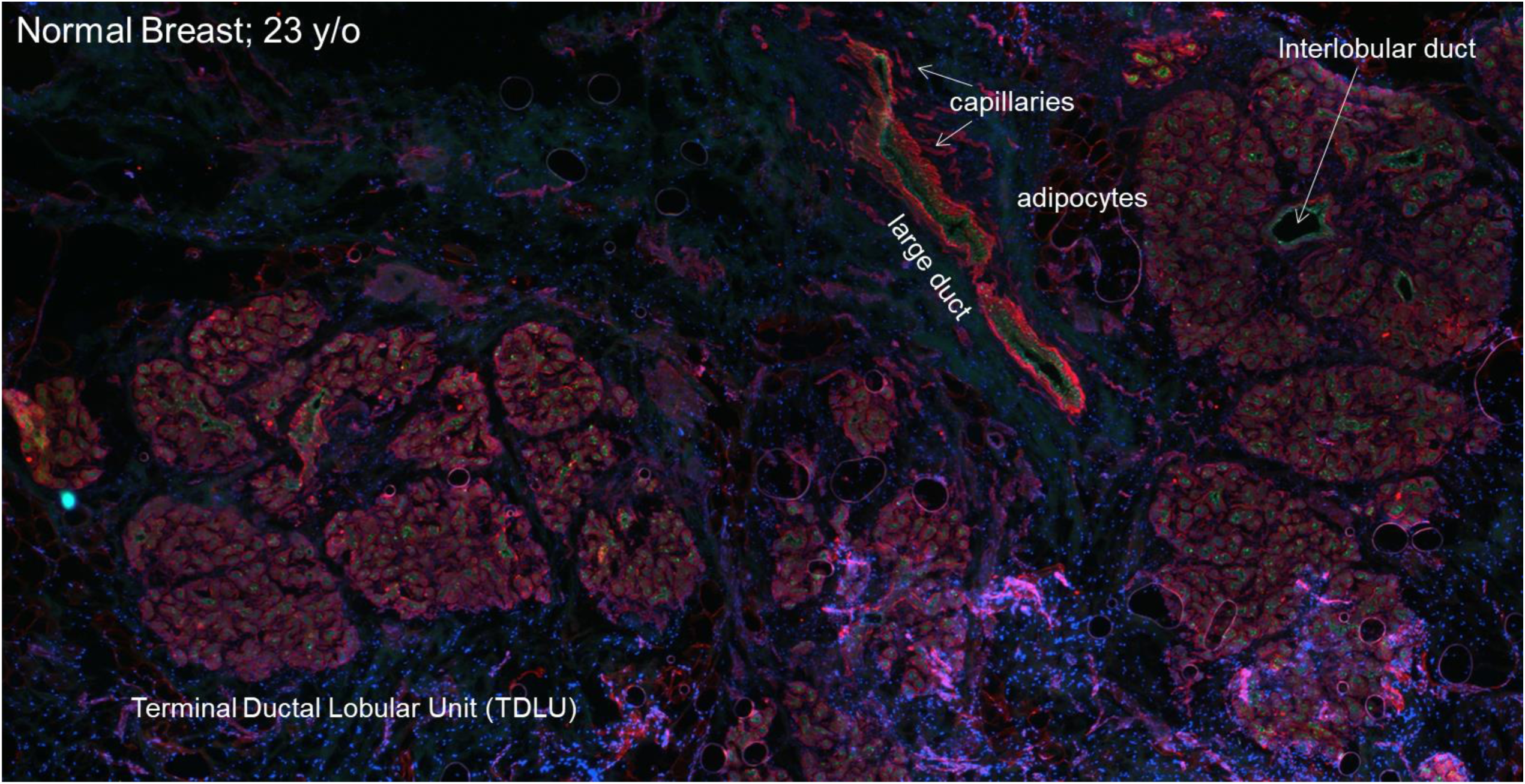
Breast Tissue Landscape. A low-power (4x) microscopic tiled scan of normal breast tissue (reduction mammoplasty, 23-year-old female) immunostained with keratin 18 (green) and pan-laminin (red) antibodies exposes the arrangement of cell types and tissue elements within the breast. Several large TDLUs, a large lactiferous duct, adipocytes, interlobular duct, and a dense epithelium-adjacent vasculature are present. Nerve fibers were not observed, consistent with previous anatomical reports describing the breast’s sensory innervation being limited to superficial fascia and an areolar subplexus^54,55^. EVOS FL Auto imaging scan; 4X Plan LWD Fluor 0.13NA objective.

**Figure 3.**
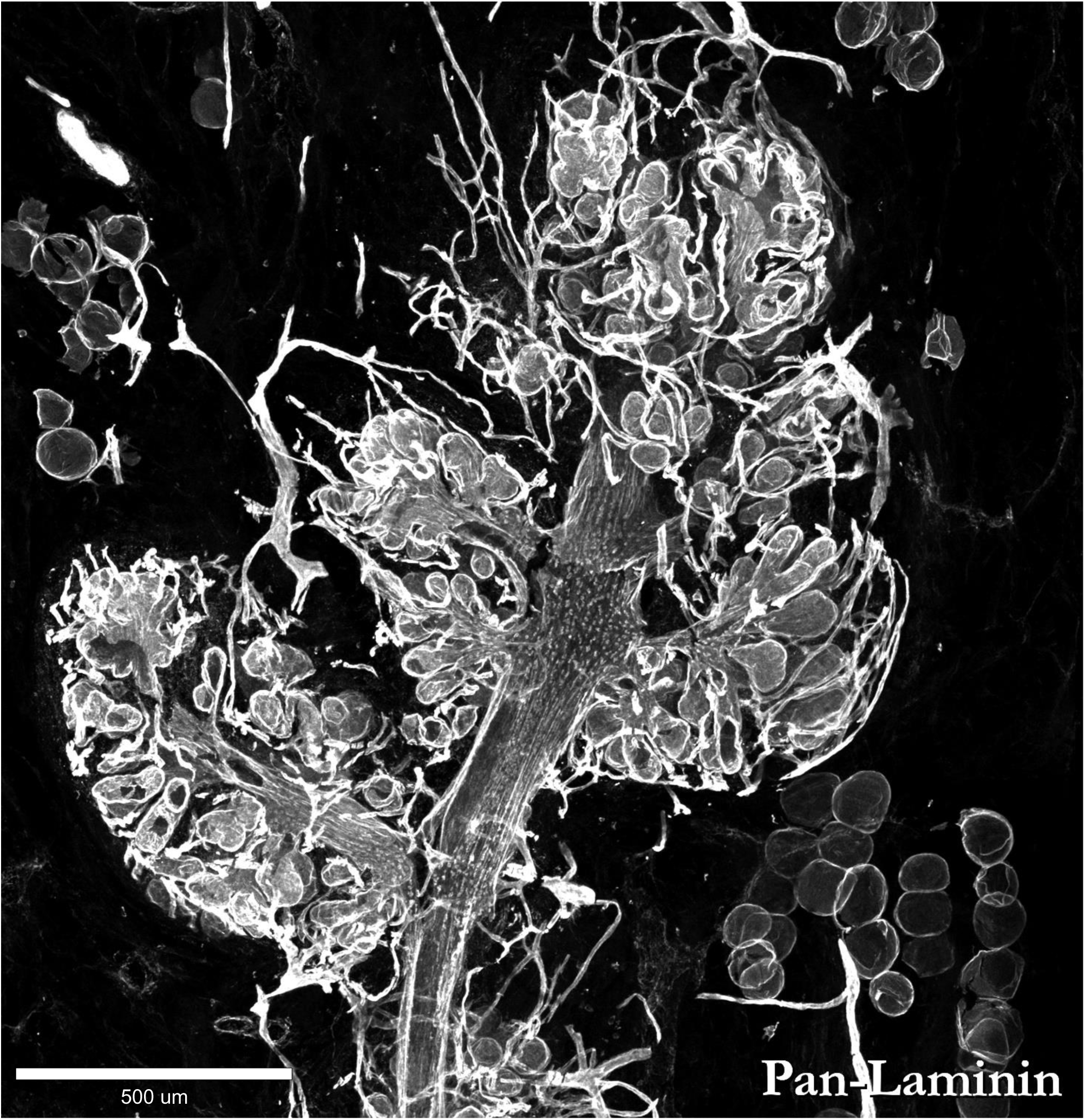
3-dimensional organization the breast’s stromal and epithelial components. A maximum intensity projection of a confocal Z-stack of a 72.25μm thick breast tissue section immunostained for pan-laminin. Staining reveals the basement membranes of the epithelium, blood vasculature, and adipocytes. This reduction mammoplasty tissue was from a 33-year-old female. Imaged on a Zeiss LSM 710 confocal microscope; Plan-Apochromat 10x/0.3 DIC M27 objective; 500μm scale.

**Figure 4.**
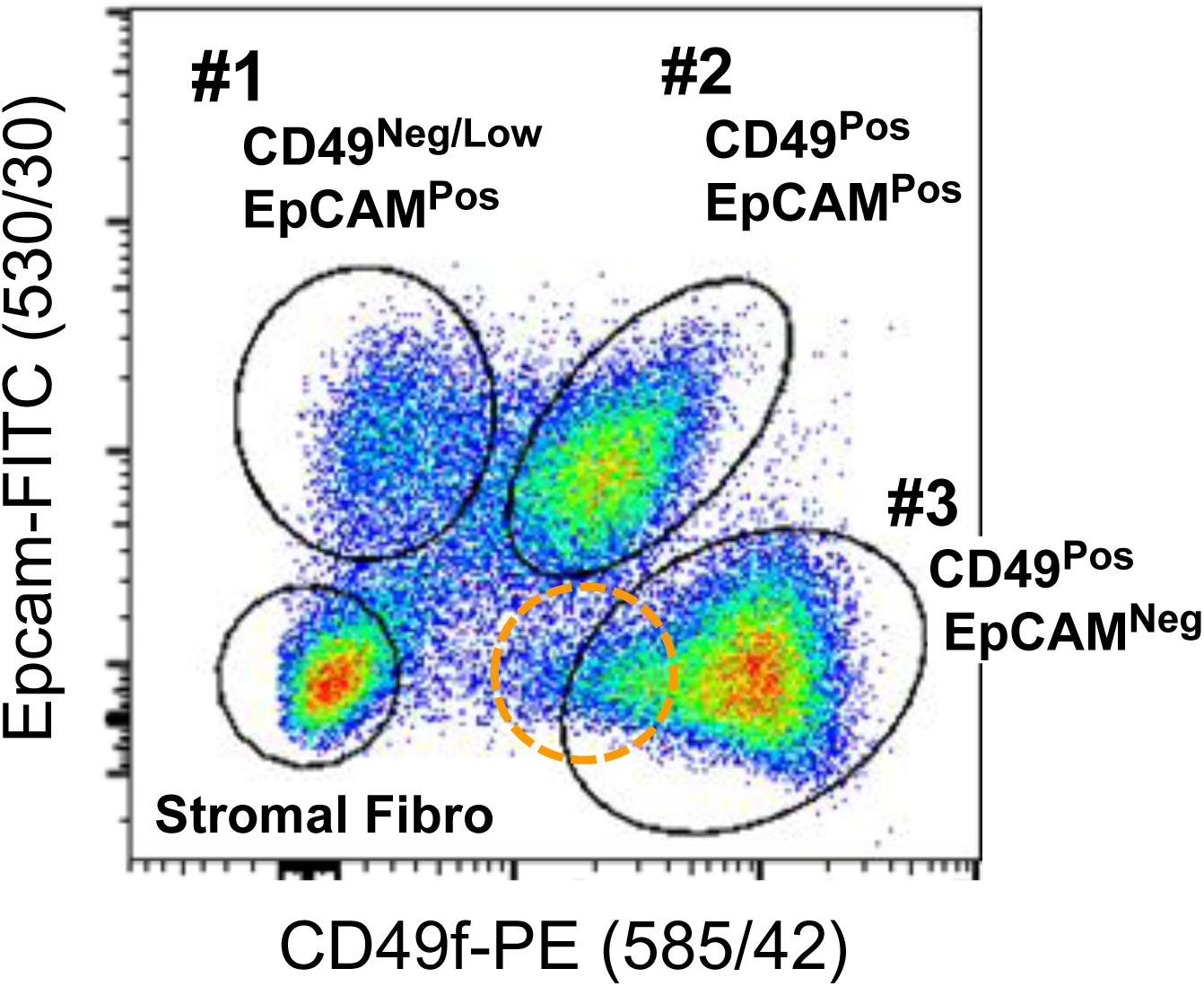
Flow cytometric analysis of breast cells. EpCAM and CD49f staining of normal breast cells resolves three epithelial cell populations by flow cytometry: #1 CD49f^Neg^ EpCAM^Pos^, #2 CD49^Pos^EpCAM^Pos^, and #3 CD49f^High^ EpCAM^Neg^. The dual negative CD49^Neg^EpCAM^Neg^ population is often described as ‘stroma’ or ‘stromal fibroblasts.’ During the development of our FACS panel, we discovered that if other markers and gates are used to categorize the cells, several other cell populations are revealed, which occupy the general area indicated by the orange dotted line. These cell types include two endothelial populations, myoepithelial cells (MEPs), luminal epithelial cells (LEPs), and a rare epithelial cell population. We have found also that the dual negative ‘stromal’ area contains fibroblasts, vascular smooth muscle cells, pericytes, luminal epithelial cells, red blood cells, and leukocytes—the relative abundance of which depends on the lineage-depletion and detection strategy employed. This reduction mammoplasty tissue was from a 22-year-old female (sample #N141).

## Results and Discussion

### FACS method development: Key examples and guiding principles

To explore the breast’s architecture and decipher its cellular composition—and also test different marker combinations we could use to FACS-isolate cells— we prospectively collected and analyzed normal tissues (reduction mammoplasty specimens) from 65 women, whose ages ranged from 16 to 61 years 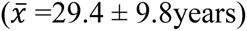. We froze, sectioned, and immunostained these tissues with over 60 different primary antibodies (Figure 5a-f, Figure 5–figure supplement 1a-f; supplement 2a-c; Figure 5–table supplement 1). We further analyzed antibodies displaying cell-type (and cell-surface) specificity by flow cytometry to determine if they could resolve viable cell populations. Our goal was to find markers that maximized resolution between cell types that would, in turn, increase the objectivity of the isolation strategy. Unfortunately, many antibodies failed in this regard despite their marked ability to expose different cell types when used on tissue. The luminal-cell-specific antibodies we tested are a prime example: That is, out of the eight we screened by tissue immunostaining, only three adequately resolved luminal cells when measured by FACS, with both CD24 and Muc1 staining more intensely and providing better resolution than the frequently used EpCAM antibody (Figure 6a-h; Figure 6–figure supplements 1-8).

**Figure 5.**
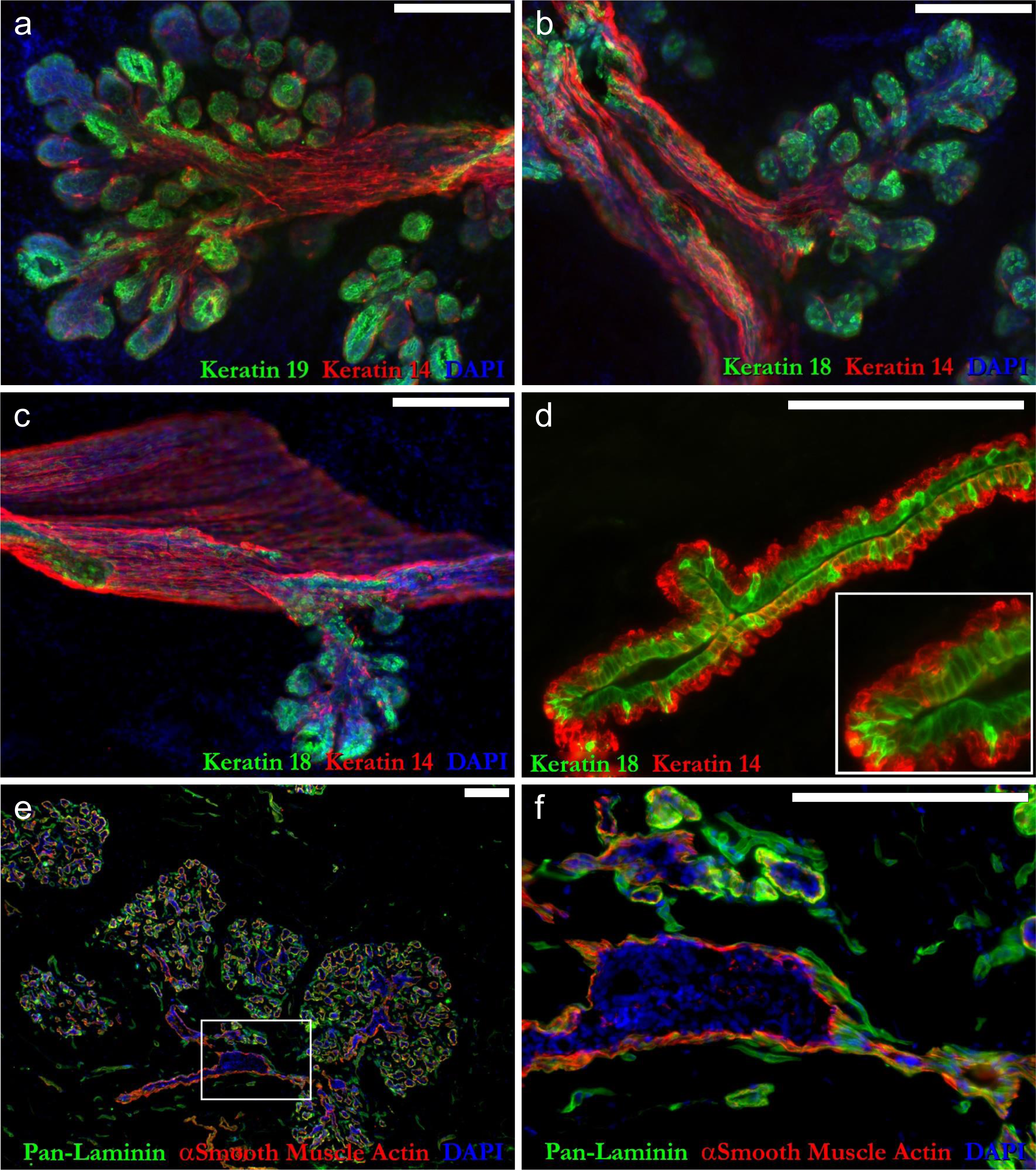
Cellular composition and architecture of breast tissues. **(a-c)** Representative immunostains of 80μm thick breast tissue sections illustrating the complex three-dimensional organization of cell types forming the lactiferous ducts and terminal ductal lobular unit structures in the breast. Myoepithelial cells, identified by intense keratin 14 staining, are arranged longitudinally along large, pleated ducts (a-c) and often appear as compressed tubes (c). Luminal epithelial cells are distinguished by their expression of keratin 18 and 19 (a-c). **(d)**10μm breast tissue section stained for keratins 18 and 14. Some luminal cell projections are seen extending basally to the basement membrane (d, magnified inset). **(e)** A low power image of 10μm breast section illustrates the branching glandular architecture and tortuous system of tubules within the TDLUs. **(f)** At higher magnification **(f, boxed area in e)**, the entangled pan-laminin staining capillary network running parallel to the glandular epithelium becomes evident. 1mm scale bar (all images).

**Figure 6.**
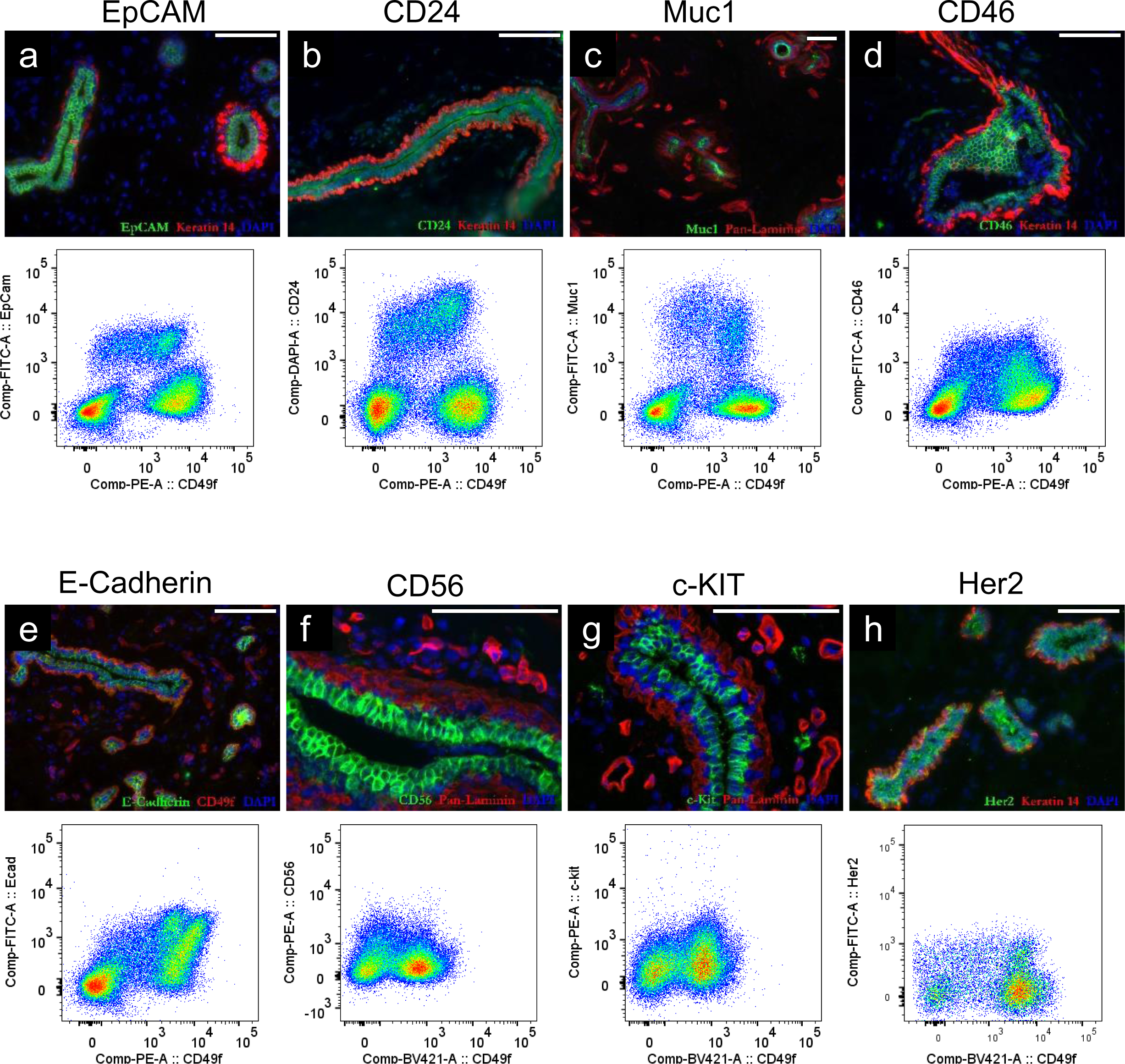
FACS antibody screen identifies CD24 and Muc1. (a-h, tissue immunostains); Normal breast tissue, immunostained with a panel of antibodies predicted to label luminal epithelial cells; Scale =0.1mm. **(a-h, FACS plots)** Flow cytometry scatter plots of breast cell suspensions (from reduction mammoplasty tissues) stained with each indicated antibody.

Why so few luminal-cell-specific antibodies could not adequately resolve luminal cells by FACS is puzzling, given that most performed exceptionally well when used on tissue sections. We attributed this discrepancy to the distinct detection strategies employed by these two methods, i.e., indirect staining of tissues vs. the direct staining used in FACS. However, the differential sensitivity of protein markers to trypsin—currently an unavoidable step in preparing cell suspensions for FACS analysis—may also contribute. Nevertheless, the antibody screen was successful, and through the exercise, we found that both CD24 and Muc1 antibodies enhanced the resolution of the luminal cell populations. We now needed to choose which of these two antibodies to use in our final staining panel—and testing them together led to a pivotal discovery.

When we co-stained breast cell suspensions with CD24 and Muc1 antibodies (each labeled with distinct fluorophores) and analyzed them by flow cytometry, most luminal cells bound both. However, we discovered that there was always a fraction of luminal cells that repeatedly and reproducibly escaped detection if we used either antibody in isolation. Some cells stained positively for one marker but not the other—and vice-versa (Figure 7a,b). This observation was critical. If we had used a single marker as is commonly practiced, most of these ‘missed’ luminal epithelial cells would have been incorrectly sorted as myoepithelial cells and gone unnoticed (Figure 7–figure supplement 1). This incorrect assignment would have confounded results and possibly produced erroneous conclusions. To avoid this problem and ensure proper luminal-cell discrimination, we discovered we needed to use both markers and take advantage of the redundancy. Therefore, we included both CD24 and Muc1 antibodies in all subsequent experiments. This unexpected finding was an object lesson and set a precedent as we moved forward with panel design. To ensure proper classification of the other cell types, we also needed to build redundancies into their sorting strategies. However, finding markers that would adequately separate each cell type by FACS was a persistent challenge.

**Figure 7.**
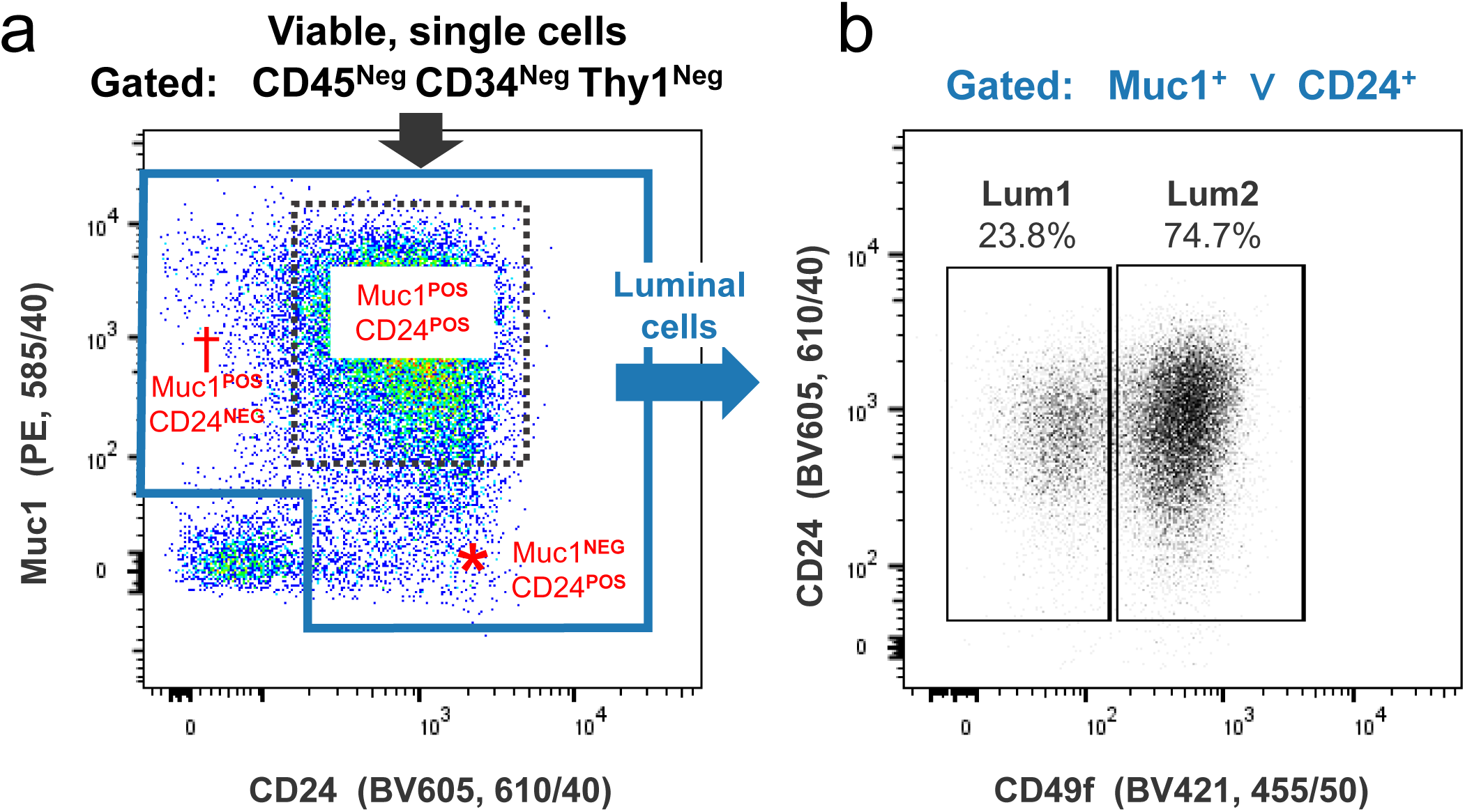
Luminal cells escape detection if single FACS marker is used. **(a)** Representative FACS analysis of normal breast cells co-stained with luminal epithelial cell markers Muc1 and CD24 (using independent markers: PE and BV605). Cells were backgated for the viability marker To-Pro-3^Neg^ (viable cells), SSCWidth^Low^ (single cells), CD45^Neg^, CD34^Neg^, and Thy1^Neg^ (which removed dead cells, cell clusters, leukocytes, fibroblasts, endothelial cells, pericytes, and myoepithelial cells from the analysis). Most remaining cells indeed stained for both Muc1 and CD24, identifying them as luminal cells (dotted box). However, a fraction of luminal cells stained only for one luminal marker, but not the other; i.e., they stained for Muc1^POS^CD24^NEG^ (†) or Muc1^NEG^CD24^POS^(*). The ‘luminal cell’ gate (blue outline) indicates cells that expressed Muc1, CD24, or both. **(b)** FACS plot (CD24 vs. CD49f) of the gated luminal cells from the previous plot. Cells were classified as being either ‘Luminal1’ (Lum1) or ‘Luminal2’ (Lum2) cells, based on their respective staining for CD49f (α6-integrin). The machine’s bandpass filter used for each fluorophore is indicated on its corresponding axis (e.g., a 610/40 filter was used for detecting the CD24-BV605 antibody).

A noteworthy example of mistaken identity and low resolution involved CD31—a lineage marker frequently used to identify endothelial cells. Like many other antibodies we tested, CD31 performed exceptionally well on frozen breast sections. The antibody clearly marked capillaries and other vessels in tissues (Figure 8a; Figure 8–figure supplement 1a). However, even after titrating and optimizing antibody staining, we found the resolution in our FACS experiments was often subpar. Identifying this problem was somewhat tortuous. It became apparent only after we had sorted epithelial fractions by FACS, isolated their RNA, and measured more than 100 transcripts using a custom quantitative RT-PCR array. Among the genes tested was the hematopoietic stem cell marker CD34, which we curiously found highly expressed in the sorted myoepithelial fraction (Figure 8–figure supplement 2a, Figure 8–Table supplement 1). This myoepithelial expression was at odds with our tissue immunostaining results, where we found no evidence of CD34 in myoepithelial cells (Figure 8b; Figure 8–figure supplement 1b; supplement 2b,c; and supplement 3a,b). Based on this peculiar result, we questioned the purity of our gated myoepithelial fraction and investigated further. We added a CD34-specific antibody to our FACS panel and repeated the analysis. Upon examining the FACS data, we immediately noticed that cells resolved by CD31 also stained for CD34 (Figure 8c,d). By comparing the inverse, i.e., exploring CD31 expression in the CD34^Pos^ gated populations, it became evident that the CD31 antibody was not resolving the entire endothelial cell population (Figure 8e). Because endothelial cells express CD49f at a level similar to myoepithelial cells, those that escaped detection by CD31 were being gated into—and were contaminating— the myoepithelial cell fraction (these were the unknown cells detected in Figure7, supplement 1). This endothelial contamination explained why we had initially detected CD34 mRNA in our FACS-isolated myoepithelial fraction. We rectified this problem by replacing the CD31 antibody with one specific to CD34.

**Figure 8.**
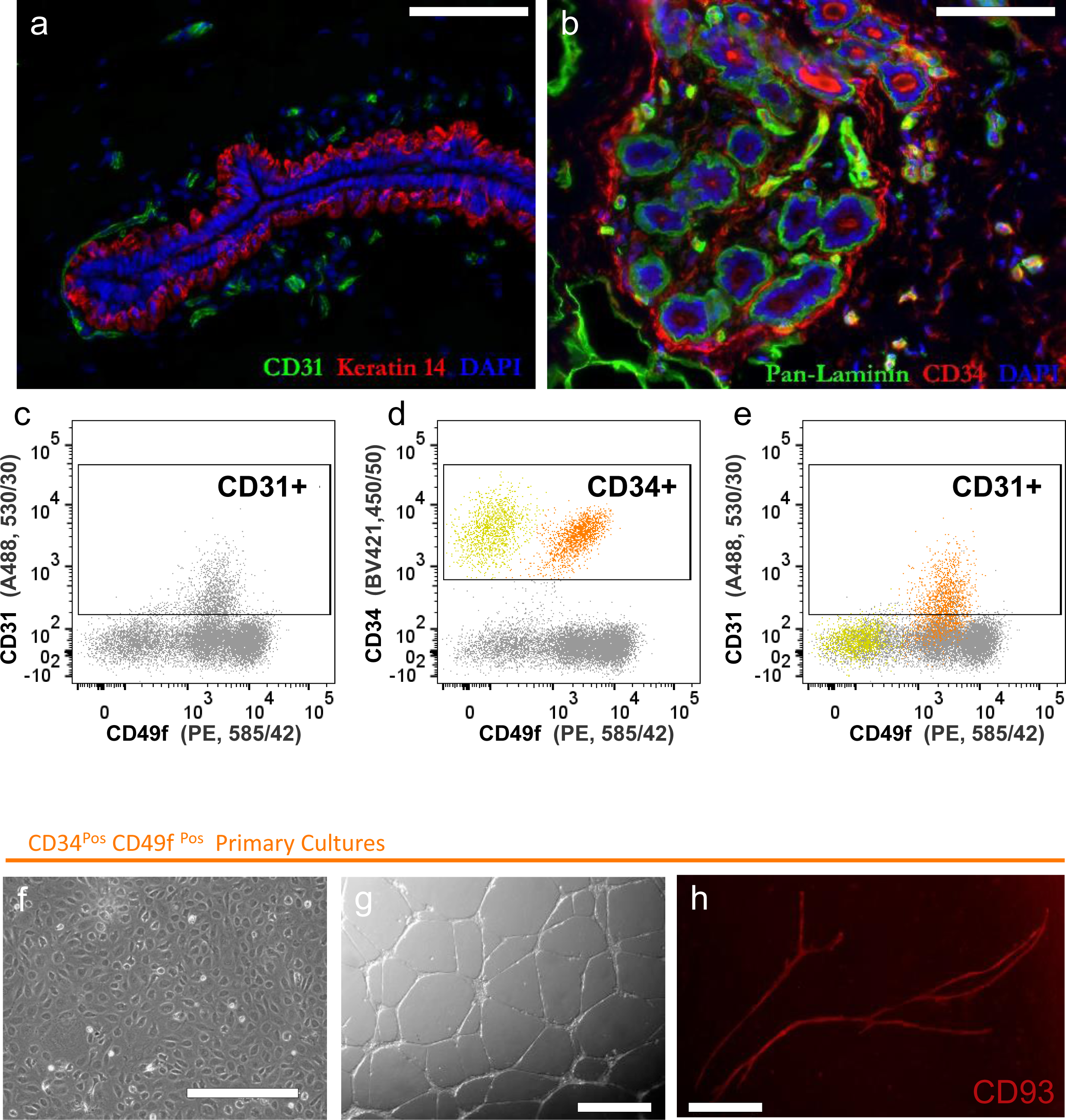
Isolation and validation of vascular endothelial cells. **(a)** Breast tissue immunostained for CD31 reveals capillaries adjacent to a mammary lactiferous duct. **(b)** CD34-staining labels endothelial and other stromal cell types. **(c-e)** FACS analysis of normal breast cells co-stained with CD31 (A488), CD34 (BV421), and CD49f (PE). (c) CD31 vs. CD49f scatterplot illustrates the incomplete resolution of endothelial cells by the CD31 antibody. (d) Same data as in c, now contrasting CD34 vs. CD49f. CD34 provides superior resolution and separates the entire endothelial cell fraction (orange) and another unknown cell population (yellow) that lacks CD49f. (e) Same data and plot as c, retaining the yellow/orange color scheme defined in d, which illustrates the incomplete separation of the orange endothelial cell fraction. **(f-h)** Primary CD34^Pos^CD49f^Pos^ endothelial cell cultures seeded (f) on 2D collagen I, (g) on top (O/T) of Matrigel™ with 5% overlay, or (h) with a fibroblast feeder layer, stained with CD93-PE to label endothelial cells and reveal vessel-like structures. Cells were cultured in endothelial growth medium (EGM-2, Lonza) supplemented with 10% FBS and incubated in a 5% CO2 humidified incubator at 37°C. Scale =100μm (a,b); 400μm (f); 200μm (g,h).

A silver lining to the above exercise is that we discovered that the resolution provided by the CD34 antibody was unmatched by any other marker we have tested to date. CD34 excelled at identifying endothelial cells when combined with CD49f. Indeed, sorting and culture of the CD34^Pos^ CD49f^Pos^ population from normal breast tissues produced homogeneous cell lines with prominent endothelial characteristics (Figure 8f). These cells formed endothelial cords and vessel-like structures when seeded on Matrigel or fibroblast feeder layers, respectively (Figure 8g,h). Because CD34 provided excellent resolution, we made it a central marker in our final FACS strategy that we outline in Figure 9.

**Figure 9.**
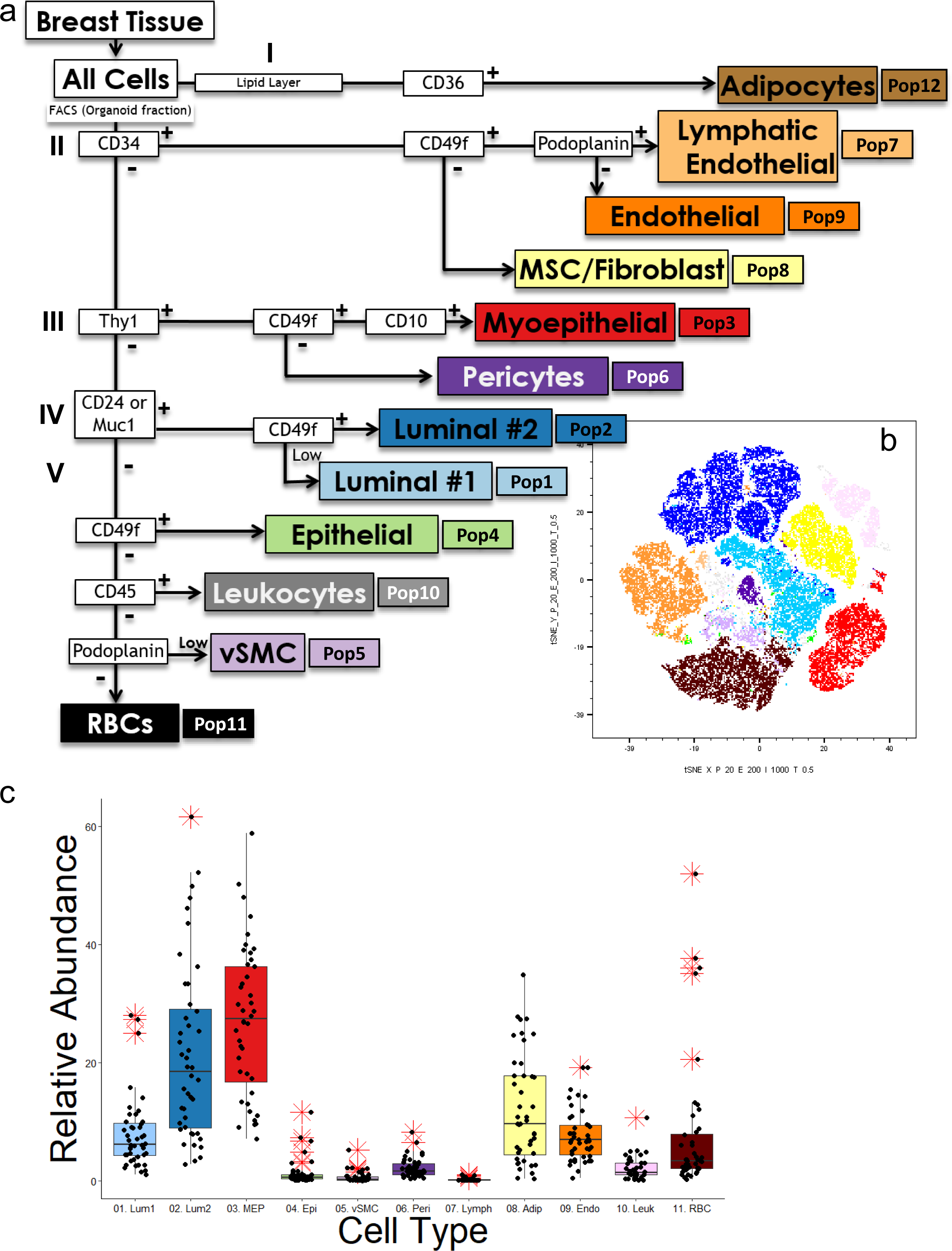
FACS strategy for comprehensive cell type isolation. **(a)** Purifying the breast cell types begins with enzymatic (collagenase) dissociation of normal breast tissues. This was followed by low-speed centrifugation, yielding an organoid pellet, supernatant, and an adipocyte-enriched lipid layer. **Node I)** We used the lipid layer for RNA isolation/RNA-sequencing and primary adipocyte culture. Cultured adipocytes were subsequently FACS purified using CD36, a marker identified by RNA-sequencing. All remaining breast cell types were FACS-purified from the organoid cell pellet, after first dissociating the organoids to a cell suspension and staining it with CD34, CD49f, Podoplanin, Thy1, CD10, CD24, Muc1, and CD45 antibodies, along with the viability marker To-Pro-3. As most FACS machines can sort into only four tubes simultaneously, purifying every population required multiple rounds of sorting. The first round of sorting isolated cells into groups (or nodes) based on their shared expression of key markers. These nodes were: **II)** The CD34^Pos^ fraction; **III)** The Thy1^Pos^ (CD34^Neg^) fraction; **IV)** The CD24^Pos^ and/or Muc^Pos^ (CD34^Neg^ Thy1^Neg^) fraction; and **V)** CD34^Neg^ Thy1^Neg^ CD24^Neg^ Muc^Neg^ fraction. After the first round of sorting, two additional rounds of sorting (required to ensure purity) were used to isolate each cell type from their respective nodes, using the indicated markers and Boolean logic scheme. For example, the CD34^Pos^ cells (node II) were FACS interrogated for CD49f and podoplanin. Lymphatic endothelial cells in this fraction were identified by their co-expression of both CD49f and podoplanin; Vascular endothelial cells were identified by their expression of CD49f and lack of podoplanin; and Fibroblasts, by their lack of CD49f. The plus (+) and minus (-) signs indicate visual differences and separation of stained cell fractions—absolute ‘negative’ expression was not explicitly evaluated. Each FACS-purified population was assigned a number, i.e., pop1-12. The identity of each of these populations was affirmed or later discovered by using the literature and prior knowledge or validation experiments, e.g., primary cell cultures, immunofluorescence staining, and RNA-sequencing. **(b)** An unsupervised tSNE projection of normal breast cells (stained and analyzed by FACS) segregated the cells into discrete populations. Each cell was overlaid with the color associated with its respective FACS gate, collectively providing a visual representation of each cell type’s relative proportion in the population. **(c)** Relative abundance of each cell type from 30 different individuals (44 independent FACS experiments); Outliers (*) are defined as points falling (1.5 x IQR) above/below the 75^th^ and 25^th^ quartile.

### Comprehensive identification and isolation of cell types

The central nodes of our sorting strategy, including markers used and cell types they each contain, are detailed in the following sections—along with evidence supporting their validation. For some cell types, we needed primary cultures to establish or confirm their identity. In other cases, molecular data from RNA-sequencing experiments provided the necessary clues. In the end, we used a panel of nine different antibodies and the viability dye To-Pro-3 to isolate twelve viable cell types. These antibodies are specific to CD36, CD34, CD49f, Podoplanin, Thy1, CD10, CD24, Muc1, and CD45. The development of this panel and the design of our final gating strategy were guided by tissue immunostaining using these and many other antibodies (Figure 5—Table Supplement 1). Select images of tissues stained with several of these markers—from the thousands collected— are provided in Figure 10, along with high-resolution images and RNA-seq derived transcript levels in each isolated cell fraction (Figure 10–figure supplements 1-12). The mRNA sequencing and thorough analysis of transcripts from each cell type is provided in an adjoining article (Del Toro, et.al., in preparation).

**Figure 10.**
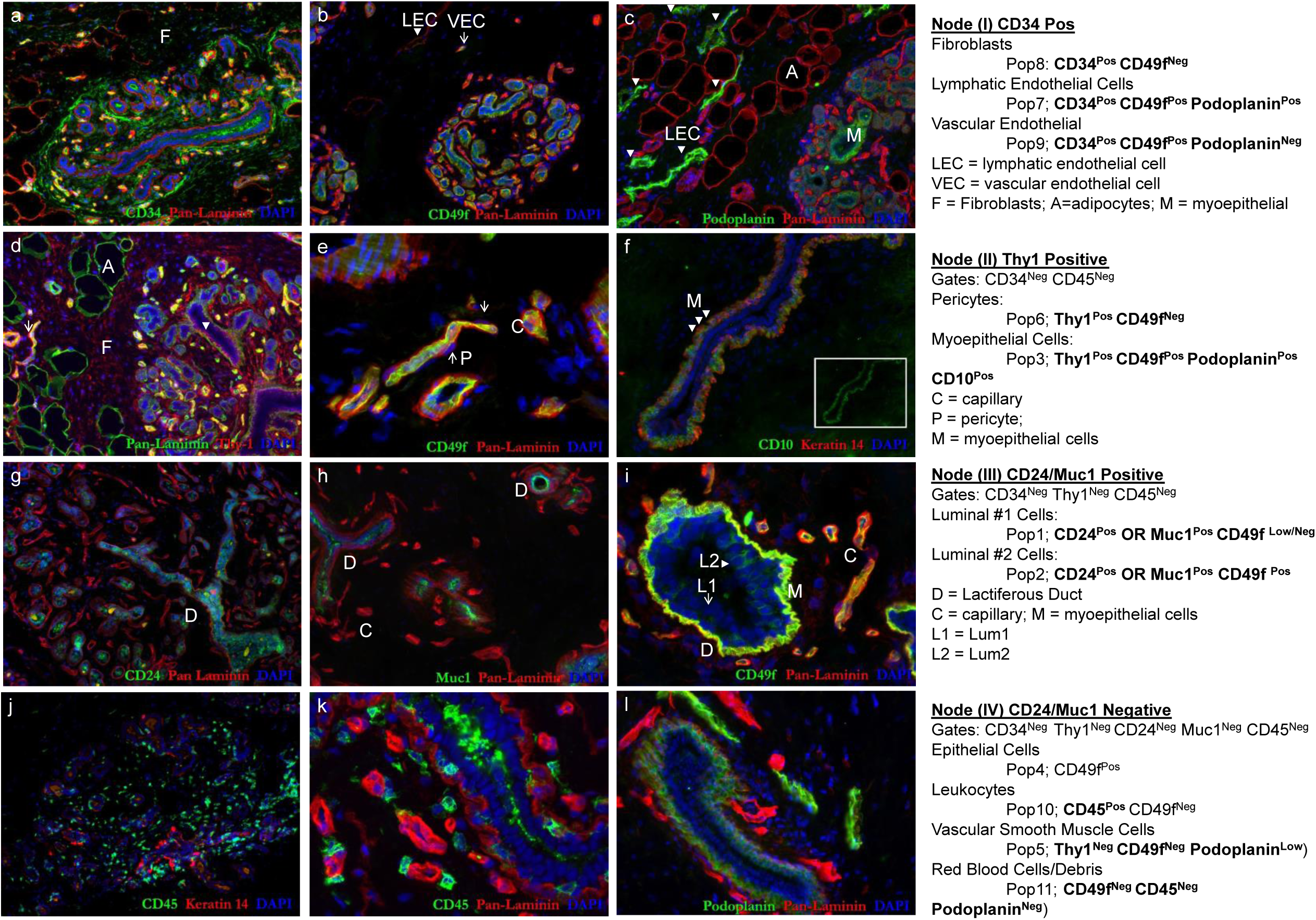
FACS marker expression in breast tissues. Representative immunofluorescence images of tissues stained for critical markers used in the FACS isolation strategy **(a)** CD34 is expressed by breast fibroblasts (Pop8) and both endothelial cell types (Pop7&9). **(b)** Among the CD34 expressing cells, CD49f is expressed by vascular (Pop9, 〈) and lymphatic (Pop7, ▾) endothelial cell types, which can be distinguished by their differentially high expression of podoplanin by the lymphatic endothelial cells (**c**, ▾). Podoplanin is also expressed by myoepithelial cells and fibroblasts—expression that does not interfere with cell sorting because other distinguishing markers identify these different cell types. **(d)** Thy1 is widely expressed. Among the CD34 negative cell populations, Thy1 is found in **(e)** pericytes (Pop6, ↑) and **(f)** myoepithelial cells (Pop3,▾); two cell types that are FACS separated by their differential expression of CD49f. **(g)** CD24 and **(h)** Muc1 are expressed exclusively by both luminal epithelial cell types. The two luminal cell types are themselves distinguished by their differential expression of **(i)** CD49f; as luminal cells are either CD49f^Neg/Low^ (Pop1, ↑) or CD49f^Pos^ (Pop2, ▾). **(j,k)** CD45 staining reveals leukocytes (Pop10). **(l)** Analysis of Podoplanin (and CD49f) is used in the final FACS node to separate the remaining cell types. All staining was performed on normal breast tissues (cryosectioned reduction mammoplasty tissues).

Our final FACS gating strategy, as presented in Figure 9, has five primary nodes: I) The adipocyte-containing lipid layer; II) The CD34-positive population containing fibroblasts and vascular and lymphatic endothelial cells; III) The Thy1-expressing fraction containing myoepithelial cells and pericytes; IV) The CD24 ∨ Muc1 positive (inclusive disjunction-‘or’) gate that defines the two luminal cell types; and finally, V) The combined CD24- and Muc1-null exclusion gate that contains CD45^Pos^ leukocytes, erythrocytes, vascular smooth muscle cells and a newly-identified epithelial cell population. We have successfully implemented this gating strategy over a hundred times on scores of samples. In addition, we have reproduced results on multiple FACS platforms using different fluorophore arrangements and strategies tailored to each machine’s capabilities (Figure 9–figure supplements 1, 2).

The isolation process begins by processing tissues to generate cell suspensions that can be stained and FACS analyzed. The isolation, identification, and validation of each cell type, including derivation of primary cell models, occurred as follows:

### Node1: Lipid Layer (Adipocytes)

To prepare cell suspensions for FACS analysis, we processed and collagenase-digested normal breast tissues from reduction mammoplasties, as described previously^5, 26^. Centrifugation of these digests separated an oily adipocyte-containing surface layer from the remaining organoid fraction. We collected these lipid layers, evaluated their cellular content by fluorescent microscopy (staining a small aliquot with Hoechst 33342), isolated total RNA from each oil fraction, and archived the RNA for later sequencing. Furthermore, we created primary adipocyte cultures by seeding small portions of the oil (100-300μl) into collagen-coated T-25 flasks, using a ceiling culture method^27^. Emerging adherent colonies contained intracellular lipid-containing vesicles that stained positively for BODIPY™ (Figure 11a,b). Furthermore, transcriptome analyses identified CD36 as a marker differentially expressed by adipocytes that we found could be applied to purify cultured adipocytes from potentially contaminating fibroblasts (Figure 11– figure supplement 1a-c; Del Toro, et.al., in preparation).Genes we found uniquely expressed in the lipid layer included leptin (LEP) and glycerol-3-phosphate-dehydrogenase 1(GPD1)— a key enzyme required for re-esterification of fatty acids to form triacylglycerol—that, along with other genes, demonstrated the oil layer was indeed highly enriched with adipocytes.

**Figure 11.**
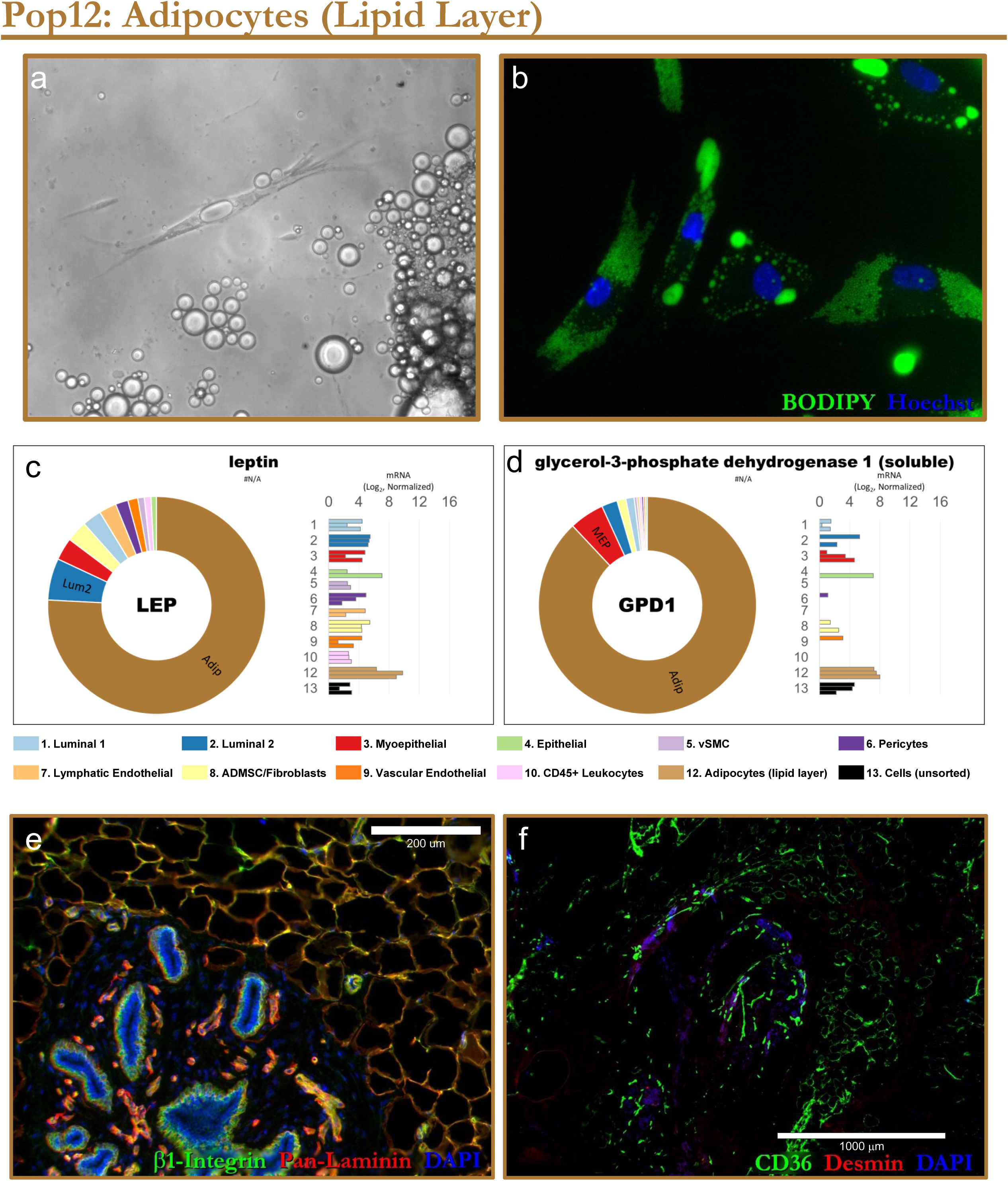
Isolation and validation of adipocytes. **(a)** Adipocytes (Pop12) were isolated from the oil layer derived from collagenase digested tissues. Roughly 250μl of oil was injected into tissue culture flasks (filled to the brim with medium) and cultured upside-down for 1-2 weeks— known as a ‘ceiling’ culture. Adipocytes floated to the top and attached to the collagen-I coated surface; After 1-2 weeks, most of the medium was removed, the dish flipped over, and the cells propagated as a typical adherent cell line. (a) phase-contrast image of lipid vesicles within of the primary adipocytes. **(b)** Nonpolar BODIPY^®^ (493/503) staining of lipid droplets within the primary adipocyte cultures. **(c,d)** RNA from uncultured adipocytes (oil layer from tissue digests) was extracted and subjected to RNA-sequencing. Among the genes uniquely or differentially expressed by these adipocyte fractions included: **(c)** the adipocyte hormone leptin (LEP) and **(d)** glycerol-3-phosphate dehydrogenase 1 (GPD1)— an enzyme critical to triglyceride synthesis. Normalized mRNA values (rlog, DESeq2) are provided on a log2 scale (horizontal bar graph showing each biological replicate) and linear scale (donut graph of median values), each color-coded by cell type. mRNA from unsorted cells served as a comparator control (Pop13, black bars). **(e,f)** Adipocytes within breast tissues are revealed by staining tissues for **(e)** β1-integrin (CD29), **(e,f)** Pan-laminin or **(f)** CD36.

To purify the remaining breast cell types by FACS, we dissociated the organoid fractions (from the above collagenase digestions) with trypsin. We stained cell suspensions with antibodies specific to cell surface markers and interrogated the cells using an advanced flow sorter. Unused organoids were viably archived in liquid nitrogen to permit multiple repeat FACS analyses. Before collagenase digestion, we thoroughly rinsed the tissues with phosphate-buffered saline (PBS) to visibly remove as much blood as possible. We did not use hypotonic RBC-lysis reagents on cell preparations because we were unsure if the osmotic stress would negatively impact other cell types or trigger significant changes in gene expression. Furthermore, we did not apply any method that would alter the proportion of cell types in these cell suspensions, e.g, by removing cell lineages with magnetic beads or using any other method, as our goal was to identify every cell type in the tissue. Finally, we kept the cells chilled throughout the entire process, except during collagenase digestion, and used the viability marker (To-Pro-3) during the sorting to exclude compromised cells. Essentially, every cell in these samples was analyzed and binned into 11 FACS-defined cell populations (the adipocytes from the lipid layer bring the total to twelve identified cell types). The first marker we used in our FACS gating scheme to isolate each cell fraction was CD34 (Figure 9).

### Node2: CD34 (Vascular and Lymphatic Endothelial cells and Fibroblasts)

CD34 was chosen as a central marker in our sorting strategy because of its pronounced ability to resolve endothelial cells, when combined with CD49f, as introduced in Figure 8. However, when we stained and analyzed breast cells by FACS, we noticed CD34 resolved yet another cell fraction on 2D scatterplots (Figure 12a, yellow box). This unknown cloud of cells lacked CD49f. Because its identity was a mystery, we named it ‘Population 8,’ or ‘Pop8’ for short.

**Figure 12.**
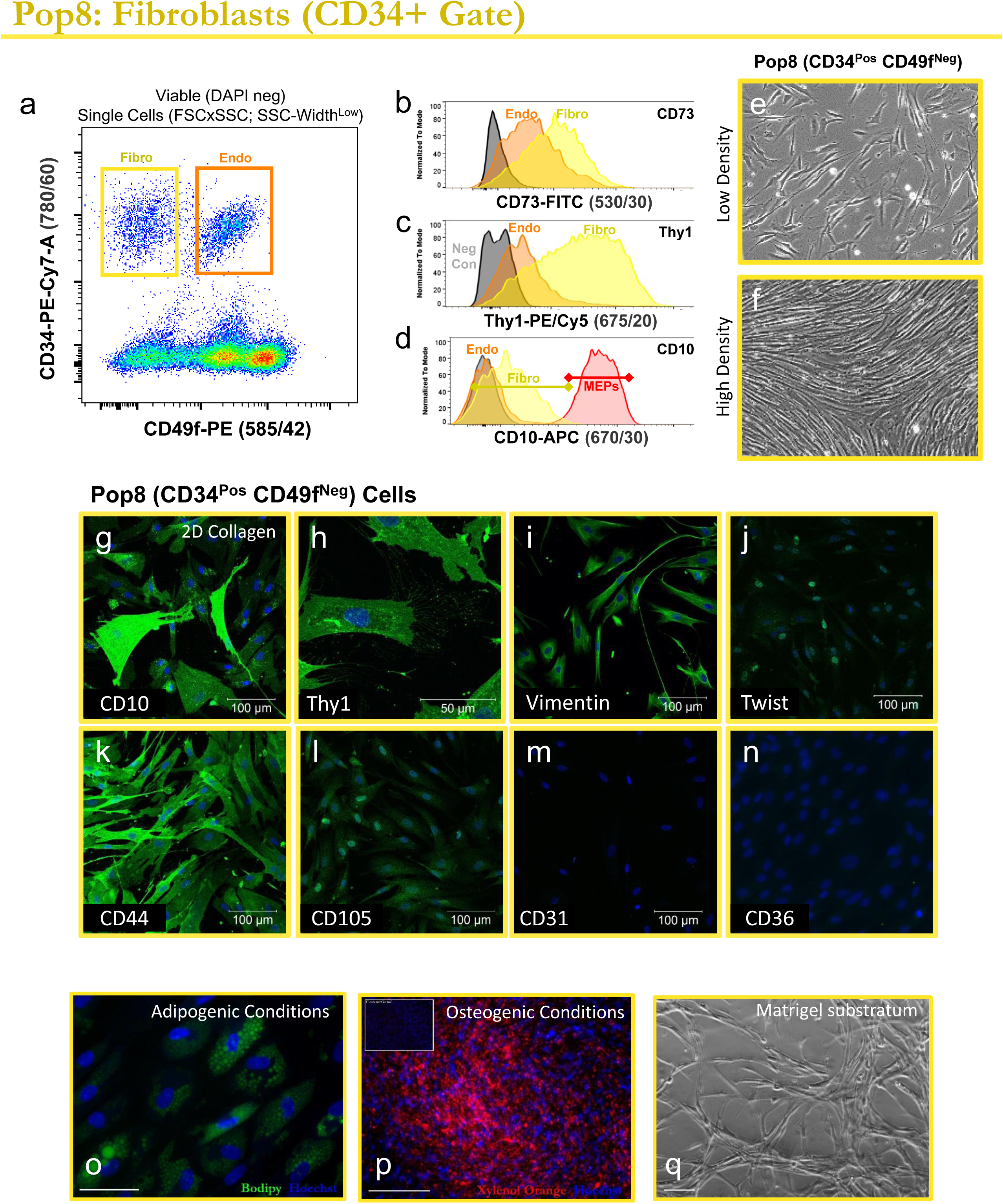
Identification and validation of fibroblasts. **(a)** Flow cytometry analysis of normal breast cells, backgated for viable single cells. A discreet cell population (Pop8, later discovered to be fibroblasts), was discovered in the CD34^Pos^ FACS fraction, due to its lack of CD49f staining (yellow box), which distinguished it from CD49f^Pos^ endothelial cells (Pop9, orange box). **(b-d)** Discovering the identity of Pop8 cells was aided by recognizing this population also stained positive for (b) CD73, (c) Thy1, and (d) CD10, markers frequently used to isolate mesenchymal stem cells. Each histogram overlay displays gated Pop8 fibroblasts (‘Fibro’, yellow), Pop9 endothelial cells (‘Endo’, orange), unstained cells (‘neg con’, gray), and in (d) Pop3 myoepithelial cells (‘MEPs’, red). **(e,f)** Primary Pop8 cultures, imaged at (e)low and (f)high densities, display a characteristic mesenchymal phenotype. **(g-n)** Primary Pop8 (CD34^Pos^CD49f ^Neg^) cultures immunostained for CD10, Thy1, vimentin, twist, CD44, CD105, CD31, and CD36. **(o-q)** Pop8 fibroblasts cultured under (o) adipogenic or (p) osteogenic conditions; or (q) on top of Matrigel™ substratum.

During the development of our FACS strategy, we assigned numbers to each FACS gate to make it easier to label and track samples at the benchside. As was the case for Pop8, these names were often assigned long before we had ascertained the gated population’s identity. The numbering system began with the epithelial types (Pops1-3) and evolved from there. We have retained these shorthand notations for brevity and clarity, as they refer, specifically, to the cell fractions defined by the FACS gating scheme outlined in Figure 9 (Figure 9–figure supplements 1&2).

To establish the identity of ‘Pop8’ (CD34^Pos^ CD49f^Neg^) cells, we co-stained them in subsequent flow cytometry experiments using a collection of antibodies we had assembled while developing our FACS panel. Through this screen, we found Pop8 cells uniformly expressed CD73, Thy1(CD90), and moderate levels of CD10 (Figure 12 b-d). Staining by CD10 and Thy1 was surprising because we anticipated the expression of these markers would be limited to myoepithelial cells, as they are frequently used for this purpose^28–30^. Yet, Pop8 cells were clearly not myoepithelial cells. When seeded into culture, we found these cells were elongated and had a mesenchymal spindle shape (Figure 12e,f). Immunostaining of these cultures demonstrated the cells indeed stained positively for CD10 and Thy1, along with Vimentin, Twist, CD44, CD105, and alpha-1 type I collagen. However, they did not express CD31 or CD36 (Figure 12g-n; Figure 11–figure supplements 1,2). These data collectively indicated that Pop8 cells were fibroblasts. When we examined marker expression in breast tissues, we found fibroblasts indeed expressed the pattern of markers defining the pop8 FACS gate (i.e., they were CD34^Pos^ and CD49f^Neg^; Figure 12–supplementary figures 4a,b; 5a). We also observed stromal fibroblasts in the tissues stained faintly for CD10. We were initially surprised by this finding, as we had overlooked this weak fibroblast staining in the past—likely because intense staining of myoepithelial cells overshadows it. Previous reports have indeed described CD10 expression in fibroblasts (within breast tumors)^31^, and we should have expected this, as CD10 is an established marker of bone-marrow-derived mesenchymal stem cells (MSC)^32^. Interestingly we discovered Pop8 shares more than just CD10 expression with MSCs.

While developing our FACS panel, and attempting to identify Pop8 cells, we noticed that this cell fraction expressed other prominent markers frequently used to define bone-marrow and adipose-derived MSCs^33–35^. Along with CD10, these included CD34, CD73, Thy1, CD44, and CD105 (Figure12 a-d,g-n). This unexpected connection prompted us to explore whether Pop8 fibroblasts exhibited lineage-differentiation potentials commonly ascribed to MSCs^36–38^. We found they did. Primary cultures of early-passage Pop8 cells, placed under classic adipogenic or osteogenic conditions, exhibited tell-tale signs of directed differentiation, i.e., deposition of calcium and lipid accumulation (Figure 12o-q). Because Pop8 cells exhibited these abilities— and to be consistent with current literature—we defined Pop8 as ‘adipose-derived mesenchymal stem cells’ (ADMSC); however, we most often refer to them as ‘fibroblasts.’

After identifying the CD34-expressing CD49f^Neg^ fraction as fibroblasts and the CD49f ^Pos^ fraction as endothelial cells (Figure 12a), it appeared that we had revealed all CD34-expressing cell types in the breast. We later discovered that this was not the case. We found that lymphatic endothelial cells also express CD34 and that we needed another marker to resolve them. Lymphatic endothelial cells have received little attention in the breast literature, especially in the context of normal breast tissues. As such, we could not locate literature on how to identify, isolate, or culture these cells from the breast. A search of the vascular literature led us to a few potential markers^39, 40^, and after testing several antibodies, we discovered anti-podoplanin worked beautifully. Other antibodies, such as those specific to lyve1 and vegfr3, indeed stained lymphatic endothelium when used to stain tissue sections, but the staining was either not specific or did not produce sufficient resolution in FACS experiments (Figure 13a-e; Figure 13–figure supplement 1,2). Podoplanin staining, in contrast, resolved the entire lymphatic endothelial cell fraction (Figure 13e). We labeled this CD34^Pos^ CD49f ^Pos^ podoplanin^Pos^ cell fraction ‘Pop7’ and called the remaining podoplanin ^Neg^ vascular endothelial population ‘Pop9’. When we triple-sorted and seeded Pop7 cells into culture, these rare cells produced uniform colonies with a classical endothelial morphology that retained their differential expression of podoplanin and cultures also differentially stained for NG2 (Figure 13f-h; Figure 13–figure supplement 3a-f).

**Figure 13.**
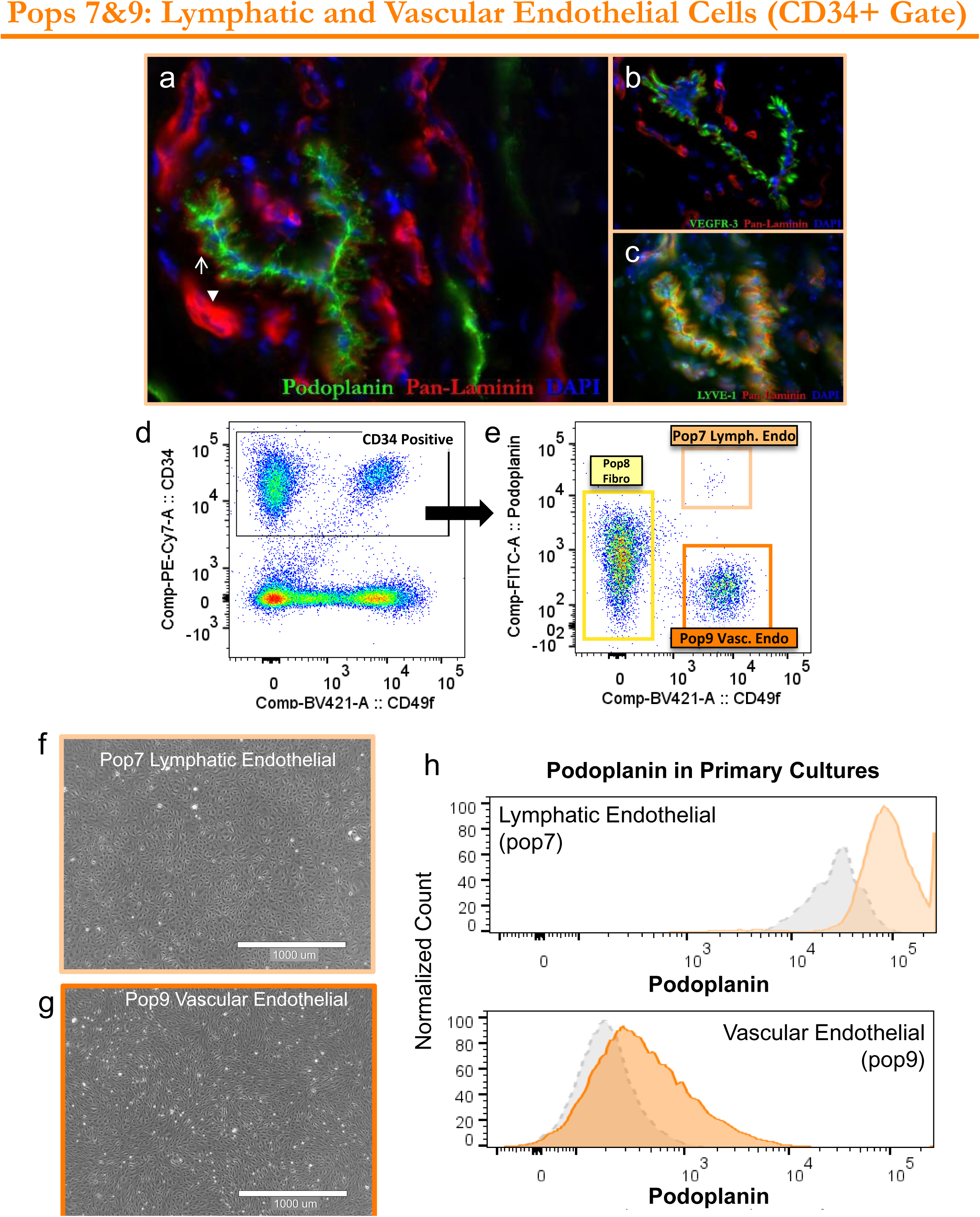
Identification and validation of lymphatic endothelial cells. **(a-c)** Normal breast tissue sections (adjacent, serial sections) immunostained for established lymphatic markers: **(a)**podoplanin, **(b)**vascular endothelial growth factor receptor 3 (VEGFR-3), and **(c)**Lymphatic Vessel Endothelial Hyaluronan Receptor 1 (LYVE-1). Lymphatic vessels expressed minimal levels of laminin, as indicated by faint pan-laminin staining (↑), which contrasted with intensely stained vascular capillaries (▾). **(d)** Flow cytometry analysis (scatterplot) of normal breast cells, backgated for viable single cells. CD34 resolved two groups of cells: pop8 (CD49f^Neg^) fibroblasts and CD49f^Pos^ endothelial cells. **(e)** High podoplanin expression in the CD34-gated fraction distinguishes Pop7 lymphatic endothelial cells (light orange box) from Pop9 vascular endothelial cells (dark orange box). Pop8 fibroblasts are resolved by their lack of CD49f expression, but express low to moderate levels of podoplanin which does not interfere with FACS purification (yellow box). **(f,g)** Primary cultures of (f) Pop7 lymphatic and (g) Pop9 vascular cells both display a classic endothelial phenotype and are themselves indistinguishable by phase microscopy (4x objective; scale =1000μm). **(h)** FACS histograms of passage 3 Pop7 (lymphatic, top) and Pop9 (vascular, bottom) endothelial cells, stained with podoplanin. Each trace is contrasted with the podoplanin levels (gray trace) measured when the cells were originally sorted two months prior (55 days). These data are of isogenic cells sorted from normal breast tissue of a 20-year-old female (sample #N218).

We confirmed the two FACS-isolated endothelial fractions (Pop7 and Pop9) by sequencing RNAs extracted from freshly sorted (uncultured) cells. We found that both populations expressed the gene encoding the endothelial marker CD31 (PECAM1) and that the two cell types differentially expressed podoplanin (PDPN), LYVE1, endothelial-associated Selectin-E (SELE), among others (Figure 13 – figure supplement 4). These data confirmed the different vascular origins of Pop7 (lymphatic) and Pop9 (vascular) and validated our FACS isolation strategy. Based on the above tissue and cell staining results and transcriptome data collected from sorted cells, we concluded that three CD34-expressing cell types exist in the breast: fibroblasts and vascular and lymphatic endothelial cells.

We have sorted breast cells from over thirty individuals using our sorting scheme–some many times over (using liquid nitrogen archived specimens for multiple repeat analyses). The CD34^Pos^ cell fraction accounts for roughly 7% of all cells in these breast samples, but this proportion varied widely between individuals, ranging between <1-19% (Figure 9c; Figure 9 - table supplement 1). The different amount of red blood cells (RBCs) that remain after rinsing the cells likely contributes to some of this variability, as RBCs are included in the entire FACS cell count. We attempted to remove as much external blood as possible by rinsing samples with PBS, but the RBC fraction still ranged between 0-52% of all cells (which also includes debris particles; 3±11%, median±sdev). Due to the indefinite nature of negative gates, it is generally more useful to consider cell proportions within defined gates—instead of calculating proportions from all ‘detected events’, a.k.a. ‘total cells.’ For example, fibroblasts compose most of the CD34^Pos^ gate (51±21%), followed closely by vascular endothelial cells (43±21%). Lymphatic endothelial cells were in the minority and composed less than 1% of all CD34^Pos^ cells (0.46±1.2%), or between 0.01-1.02% of all cells in the tissue, which establishes it as the least abundant cell type in the breast. To our knowledge, this is the first report describing how to identify, isolate, and culture lymphatic endothelial cells from breast tissues.

To resolve the remaining breast cell types, i.e., those that do not express CD34, we focused our attention on myoepithelial cells and capitalized on their differential expression of Thy1, which we established as the next marker in our gating scheme (Figure 9).

### Node3: Thy1 (Myoepithelial cells and Pericytes)

Myoepithelial cells form the epithelial compartment’s basal layer and reside between a laminin-rich basement membrane and a sheet of luminal epithelial cells (Figures 3 and 6). Intracellular markers frequently used to identify myoepithelial cells (MEPs) in breast tissues include alpha-smooth muscle actin, keratin 14, and the transcription factor p63 (Figure 14 a,b; Figure 14–figure supplement 1a,b; Figure 5–figure supplement 1,2). Surface markers include Thy1 and CD10^19, 26, 28, 30^, which we incorporated into our staining panel to provide redundancy and ensure proper myoepithelial cell identification (Figure 9; Figure 10 – figure supplements 4,6). However, when we FACS-analyzed Thy1 staining within the CD34-null fraction (Figure 14c,d), we surprisingly discovered that Thy1 resolved two populations, which were distinguished by their different levels of CD49f expression in FACS scatterplots (Figure 14 d-f). Whereas the CD49f-null population’s identity was a complete mystery, we were confident the CD49f^Pos^ cells, which we designated Pop3, were myoepithelial cells—as high CD49f expression is a characteristic feature (e.g., Figure 4, Figure 8—figure supplement 2a, Figure 10b,i). Furthermore, gated Pop3 cells also expressed significant levels of podoplanin and CD10, which corresponded to myoepithelial staining observed in tissue immunostains (Figure 14f,g; Figure 10f,l). Notably, the CD49f-null fraction, which we named Pop6, did not express either of these two markers, indicating Pop6 was a discreet cell population. (Figure 14f,g,i; Figure 14–figure supplement 3a; Figure 10 – figure supplement 3,6). Although Thy1 resolved both Pop3 and Pop6 in our FACS strategy, it is important to emphasize that Thy1 was also expressed by Pop8 fibroblasts and, to a lesser degree, by Pop9 endothelial cells (Figure 14h). However, these cell types are removed by the preceding CD34 gate, which is why we placed CD34 analysis before Thy1 in our gating strategy.

**Figure 14.**
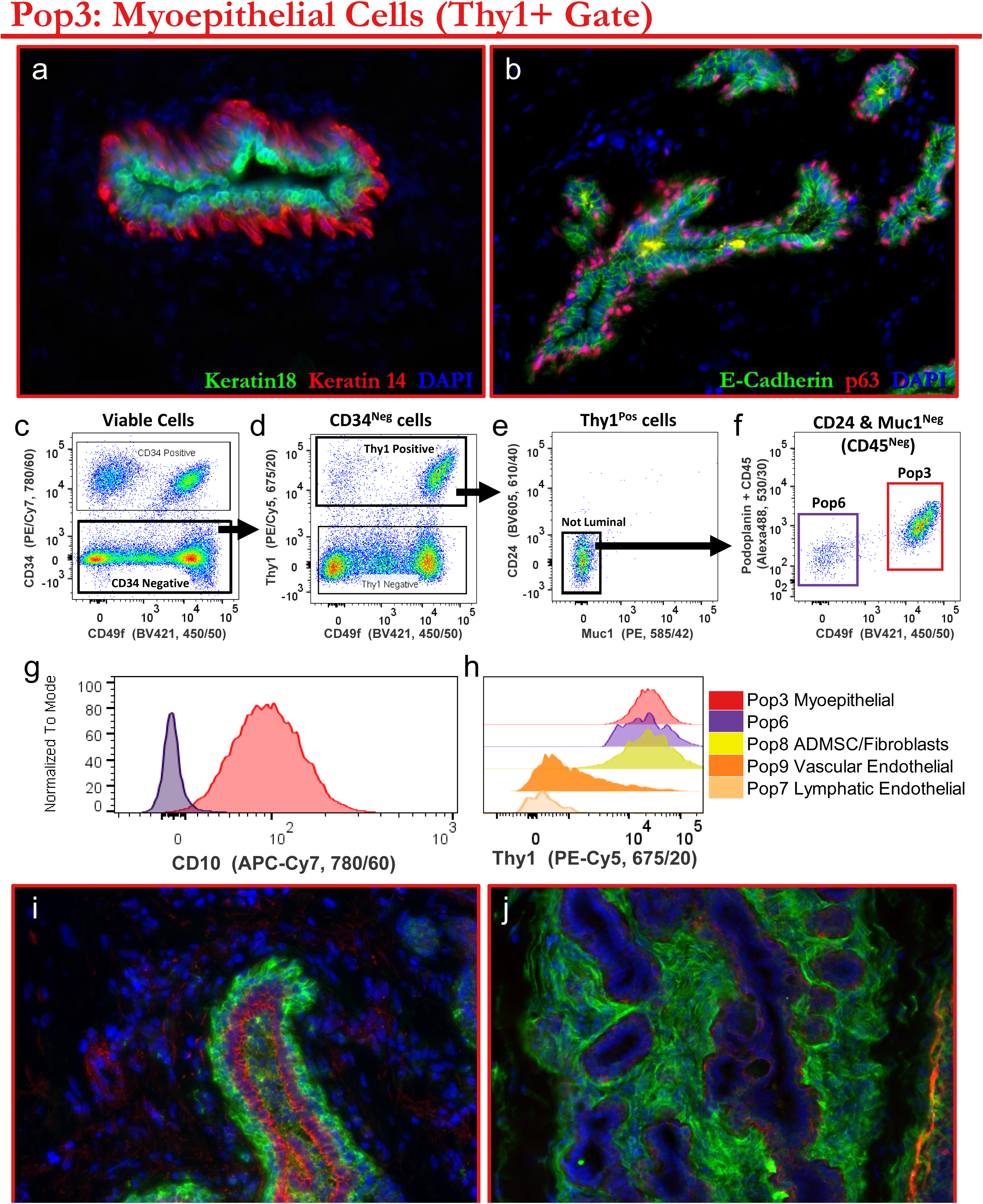
Identification and validation of myoepithelial cells. **(a,b)** Normal breast tissues immunostained for myoepithelial-associated proteins **(a)** keratin 14 and **(b)** p63, counterstained with keratin 18 and E-cadherin, respectively. **(c-f)** FACS gating strategy for isolating myoepithelial cells: (c) CD34 Negative cells are gated and analyzed for (d) Thy1 expression. (e) The CD34^Neg^Thy1^Pos^ cell fraction was refined by excluding any cell staining positively for CD24 or Muc1 luminal markers. (f) The resulting CD34^Neg^ Thy1^Pos^ CD24^Neg^ Muc1^Neg^ cell fraction was then interrogated for CD49f and podoplanin (along with CD45 when using the 8-channel FACS strategy depicted in Figure 9-supplement 1). Intense CD49f staining distinguishes Pop3 myoepithelial cells (red box, f) from a CD49f^Neg^ cell fraction (Pop6; purple box, f—later identified to be pericytes). **(g)** CD10, included as a redundant and confirmatory marker for myoepithelial cell isolation, was indeed differentially expressed between Pop3 myoepithelial and Pop6 cell fractions. **(h)** Although Thy1 expression is a defining feature of Pop6 and Pop3 myoepithelial cells, it is expressed also by Pop8 fibroblasts and, to a lesser extent, by Pop9 vascular endothelial cells (both of which are removed by the preceding CD34 gate). **(i,j)** Normal breast tissue immunostained with FACS panel markers (i) CD10 and (j) Thy1, counterstained with claudin-1 and keratin 14 staining.

To validate the myoepithelial identity and purity of Pop3 gated cells, we seeded these cells into culture and found, unsurprisingly, that they indeed exhibited the tell-tale cobblestone morphology of myoepithelial cells; Figure 14– figure supplement 3a,b). These cultures stained strongly and uniformly for myoepithelial markers p63 and keratin 14 (Figure 14 – figure supplement 3c,d; Figure 14 – figure supplement 4) consistent with RNA-seq measurements performed on freshly-sorted uncultured Pop3 cells (Figure 14 – figure supplement 3e,f). Myoepithelial cells are thus defined in our sorting strategy by eight markers, with their full designation being: CD34^Neg^ Thy1^Pos^ CD24^Neg^ Muc1^Neg^ CD45^Neg^ CD49f ^Pos^ Podoplanin^Pos^ CD10^Pos^. After solidifying the myoepithelial cells’ isolation strategy, we turned our attention to the mysterious CD49f-null (Pop6) fraction that we had previously uncovered (the CD49f^Neg^ population in the Thy1^Pos^ gate, Figure 14f).

The identity of Pop6 cells proved to be much more challenging to establish, more so than any other population we had so far encountered. We discovered this population somewhat serendipitously, so we were curious and determined to learn more about these cells. Representing less than 2% of all cells in the breast (1.6±2%, median±sdev), the scarcity of Pop6 cells contributed to our difficulties in identifying them. When we initially seeded Pop6 cells into culture, they did not attach or grow well, if at all. This was frustrating, yet itself revealing, as it contrasted with the ease and success we had in culturing fibroblasts, myoepithelial cells, and the other cell types to date. It did, however, demonstrate that Pop6 cells were different from the others. After testing different substrata and media, we eventually managed to get Pop6 cells to grow—something we now do routinely. As with every other cell type we cultured, we triple-sorted the cell populations to avoid potential contamination. Sorting multiple cycles was time-consuming yet essential to ensure we had pure populations, and it was especially crucial here as any contaminating cell type could easily overtake this slow-growing culture.

Once Pop6 cells had attached to the culture dishes and begun to grow, we discovered they exhibited a spindle-like mesenchymal phenotype that looked very much like fibroblasts. (Figure 15a). However, there were noticeable differences between pop6 cells and pop8 fibroblasts. First, Pop6 cells did not express podoplanin or CD10 like fibroblasts, and, in culture, they required more aggressive trypsinization and significant tapping of the culture dish to get them to detach. Pop6 cells also possessed a bi- and tri-polar phenotype and had a broader and more pronounced cell body than fibroblasts, especially at low cell densities (Figure 15b, Figure 15 –figure supplement 1a,b). Pop6 cells also occasionally exhibited a unique astrocyte-like stellate morphology (Figure 15c; Figure 15–figure supplement 1c-e) and would present prominent stress-actin fibers (Figure 15d-g), all of which were entirely distinct from Pop8 fibroblast cultures grown in parallel. We were perplexed.

**Figure 15.**
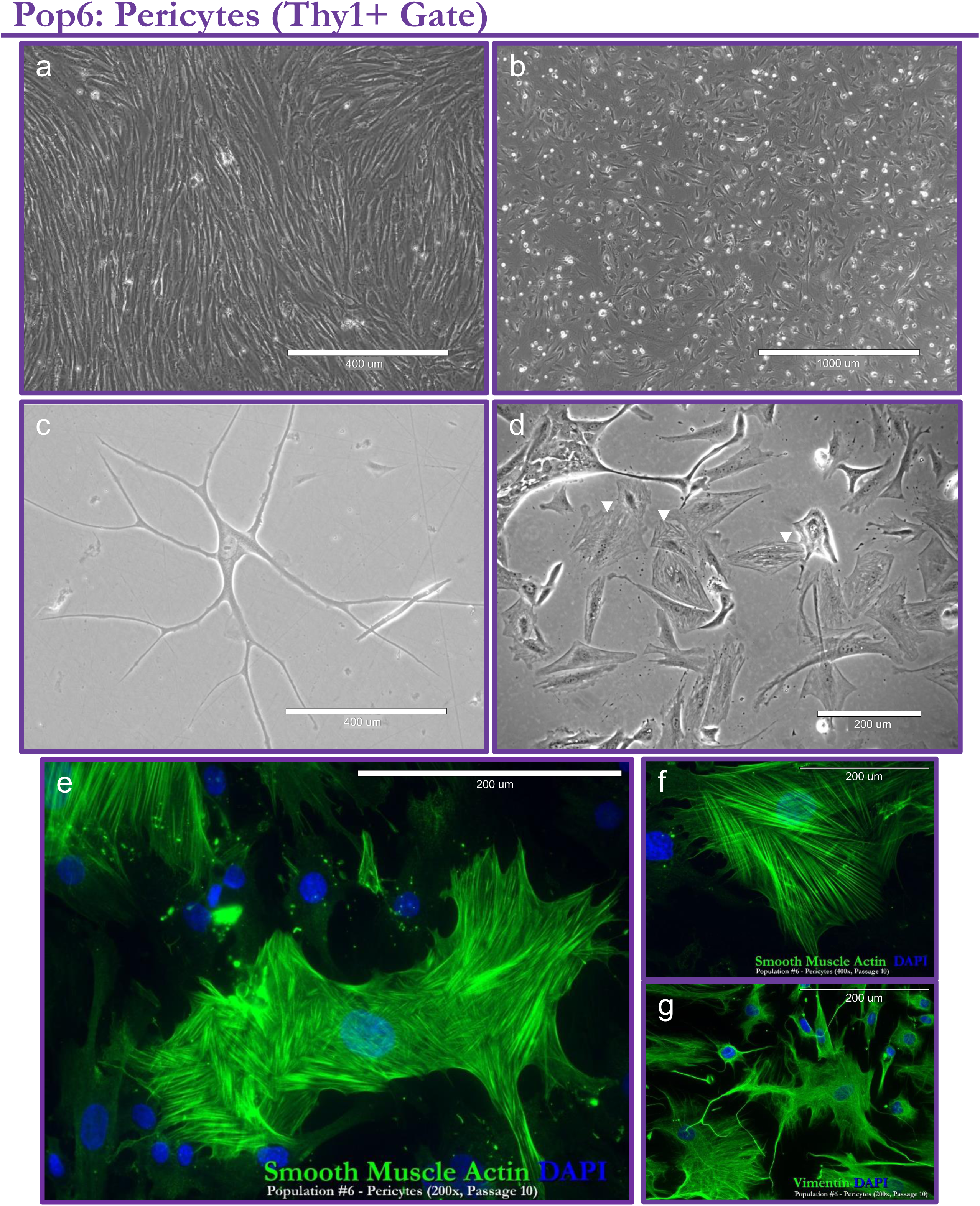
Identification and validation of pericytes I. **(a-d)** Phase contrast images of primary Pop6 cultures (later identified as pericytes). Prominent stress fibers were often observed, e.g., (d, ‘▾’). Pop6 cells were FACS-purified from normal breast tissues surgically excised from (a,b) a 23-year-old, sample #N239; (c) a 37-year-old, sample #N274; a (d) 22-year-old, sample# N141 female, and a (e-g) 22-year-old female, sample# N255 (all reduction mammoplasties). **(e-g)** primary pop6 pericyte cultures immunostained for (e,f) smooth muscle actin and (g) vimentin. Scale bars = 200μm (d-g), 400μm (a,c), and1mm (b).

Searching the literature for clues, we identified a 1985 article from Herman and D’Amore ^41^ that described bovine retinal pericytes, and these cells shared a striking resemblance to our own primary Pop6 cultures (Figure 15e,f). This connection guided us to the solution. We returned to our breast tissue staining to determine if periendothelial cells embedded within the basement membrane of blood microvessels expressed the same pattern of markers used to FACS-isolate Pop6 cells. That is, did pericytes in the tissue lack CD34 and CD49f expression? Moreover, did they express Thy1? Remarkably we found they did, indicating Pop6 cells were likely pericytes (Figure 16a-d; Figure 16 – figure supplements 1-2).

**Figure 16.**
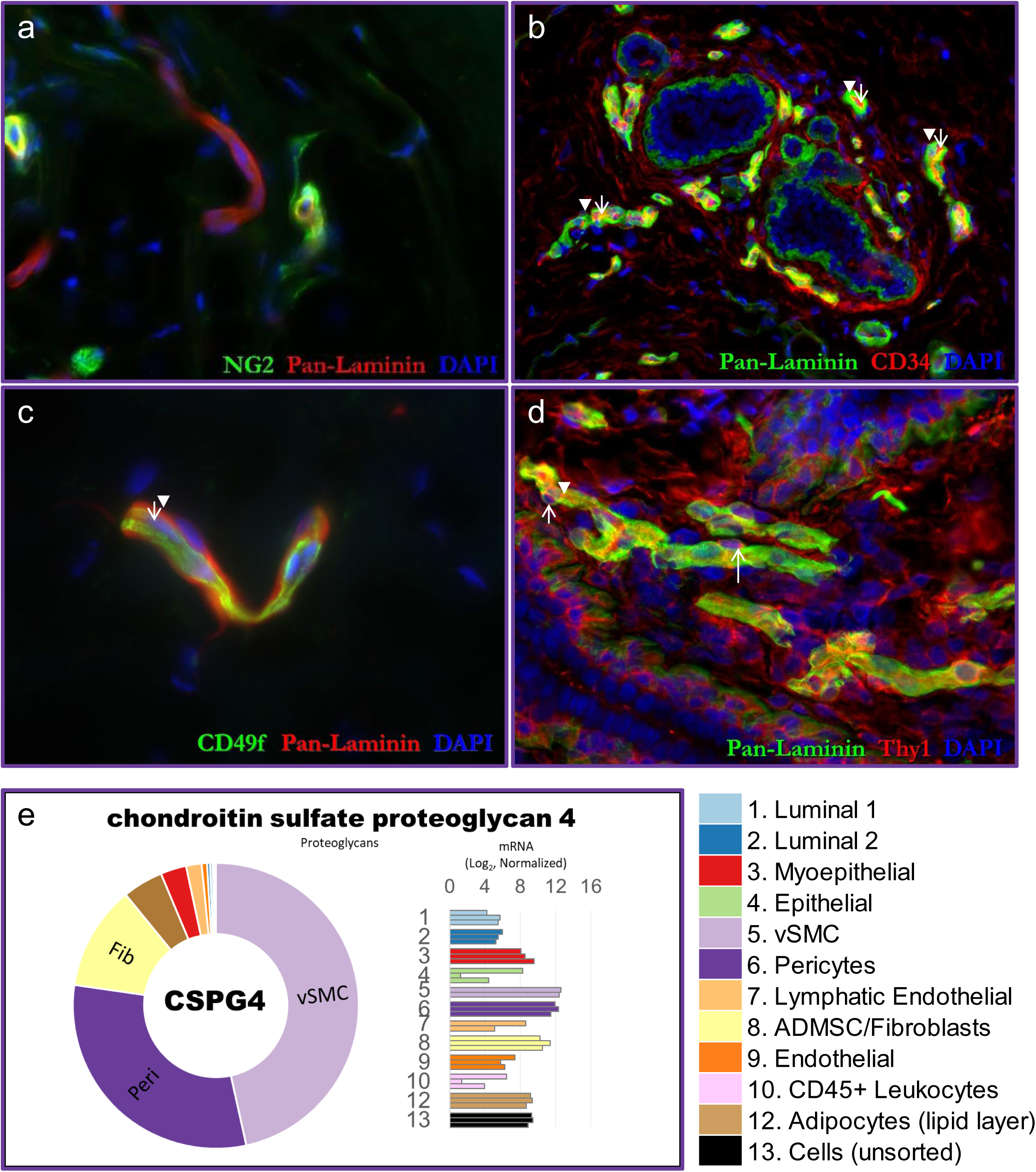
Identification and validation of pericytes II. **(a)** Normal breast tissue immunostained for pericyte marker NG2 proteoglycan, which labels a fraction of breast pericytes. **(b-d)** breast tissue immunostained for several key FACS markers defining the Pop6 gate (CD34^Neg^ Thy1^Pos^ CD24^Neg^ Muc1^Neg^ CD45^Neg^ CD49f^Neg^ Podoplanin^Neg^): (b) CD34, (c) CD49f, and (d) Thy1. Tissue staining indeed confirmed pericytes (embedded within the laminin-staining basement membrane of vessels) exhibited the pattern of staining observed by FACS. With respect to: (b) CD34, pericytes were CD34 negative (▾), whereas the endothelial cells were positive (↑); (c) CD49f, pericytes were CD49f negative (▾), whereas the endothelial cells stained positive (↑); (d) Thy1, pericytes stained positively for Thy1 (▾), along with the majority of endothelial cells (↑, as shown in Figure 12c). **(e)** Transcript levels of CSPG4 (the gene encoding NG2) within uncultured FACS-purified cell types, as determined by RNA-sequencing. Pop6 pericytes and Pop5 cells (later identified as vascular smooth muscle cells) indeed expressed the highest levels of this perivascular marker (CSPG4).

Absolute confirmation of Pop6’s pericyte-identity would not come for a couple of years until we finalized our overall gating strategy for the other cell types and sequenced and analyzed RNA isolated from each purified population. We discovered Pop6 cells indeed differentially expressed known pericyte markers that included: desmin (DES), CSPG4—the gene encoding neuron glial antigen 2 (NG2), basement membrane-associated collagen (COL4A4), RGS5, KCNJ8, and others (Figure 16 – figure supplement 3a-e). Pop6 cells also uniquely expressed serotonin receptor 1F (HTR1F, Figure 16 – figure supplement 3f). Serotonin is a vasoactive peptide and key modulator of vasoconstriction. The expression of HTR1F is notable because its unique expression by Pop6 pericytes helped us later identify Pop5, another mysterious population that uniquely expressed an HTR1F isoform.

The lack of defining markers for sorting pure pericytes has been an enduring problem that has raised doubts about the origins and identities of many primary cell models claimed to be pericytes^42^. Here, we reveal a technique that overcomes this persistent and nagging problem for breast pericytes, using combinations of markers and accounting for each cell type in the tissue. Like lymphatic endothelial cells presented above, this is to our knowledge, the first report describing how to identify, purify, and culture breast pericytes.

Although Pop6 pericytes were initially difficult to culture, we later found they would readily survive on collagen I substrate when seeded at higher densities (>5000 cells per cm^2^). This requirement stood in stark contrast to fibroblasts, which, under identical conditions, could achieve similar success rates at roughly 1/30^th^ the seeding density required by pericytes (Figure 15 – figure supplement 1f). Like adipocytes, primary pericytes in culture expressed CD36; They also retained pericyte-specific markers NG2 and did not express CD10, which distinguished them from cultures of isogenic fibroblasts grown in parallel (Figure 16 – figure supplement 4,5; Figure 15 –Supplementary figure 1e).

The Thy1 FACS node in our gating strategy thus contains two cell types: myoepithelial cells and pericytes. Cells in this node account for roughly 32% (32±13%) of all cells in the tissue, ranging between 10-61% (Figure 9-table supplement 1). Myoepithelial cells compose the bulk of this fraction (90±8%), and it is often, but not always, the most abundant cell type in the breast (27±13%). By contrast, pericytes are among the least abundant cell types in the breast (1.6±2%), composing only 8%±7% of cells in the Thy1 node on average.

Within the first three nodes in our isolation strategy (i.e., the lipid layer and the CD34 and Thy1 FACS nodes), we have identified and verified six cell types that can be isolated to purity and cultured successfully. These are adipocytes, fibroblasts, vascular and lymphatic endothelial cells, myoepithelial cells, and pericytes. To continue our quest in identifying and sorting each breast cell type, we set our sights on luminal epithelial cells— cells that possess a phenotype associated with the vast majority of breast cancers.

### Node4: CD24 ∨ Muc1(ER^Pos^ and ER^Neg^ Luminal cells)

Enveloping and forming the mammary ductal system’s interior luminal space are columnar and cuboidal epithelial cells that perform the gland’s main secretory role (Figure 17a,b; Figure 17– figure supplement 1a,b). To accurately identify luminal epithelial cells by FACS, we discovered at an early stage in our panel design that co-staining with both CD24 and Muc1 antibodies was essential to provide the necessary redundancy and assurance that luminal cells would be properly gated and not contaminate other cell populations (Figure 6, Figure 7). A common FACS strategy has been to segregate luminal cells into two populations based on their differential staining of CD49f^17–25^ (Figure 4). Although our upstream markers and gating strategy are distinct, we adhered to this approach (Figure 17c-f). We numbered the CD49f^Low/Neg^ and CD49f^Pos^ populations ‘Pop1’ and ‘Pop2’, respectively, with their full antibody designations being: CD34^Neg^Thy1^Neg^CD45^Neg^ (CD24 or Muc1)^Pos^ and CD49f^Low/Neg^ (for Pop1); or CD49f^Pos^ (for Pop2, Figure 17f; Figure 9). Others have designated similarly-isolated fractions as being respectively enriched for ‘mature luminal’ and ‘luminal progenitor’ cells— predominantly because the CD49f-null fraction has failed to produce actively growing primary cultures or outgrowths^19–21^. These designations have been supported by cell staining, which showed that the CD49f-null cell population has roughly twice as many estrogen receptor alpha expressing (ER^Pos^) cells than does the CD49f^Pos^ fraction (55 vs. 28%)^21^. Yet, we found that both would establish growing cultures when we triple-sorted and seeded Pop1 and Pop2 luminal cells into culture. Both primary cultures exhibited a squamous morphology distinct from the cobblestone appearance of Pop3 MEPs (Figure 17g,h). We attributed the successful culture of these two luminal cell types to the unique methods, stringent gates, medium, and conditions we used, which are indeed different from other reports^17, 19, 21, 23, 25, 43, 44^.

**Figure 17.**
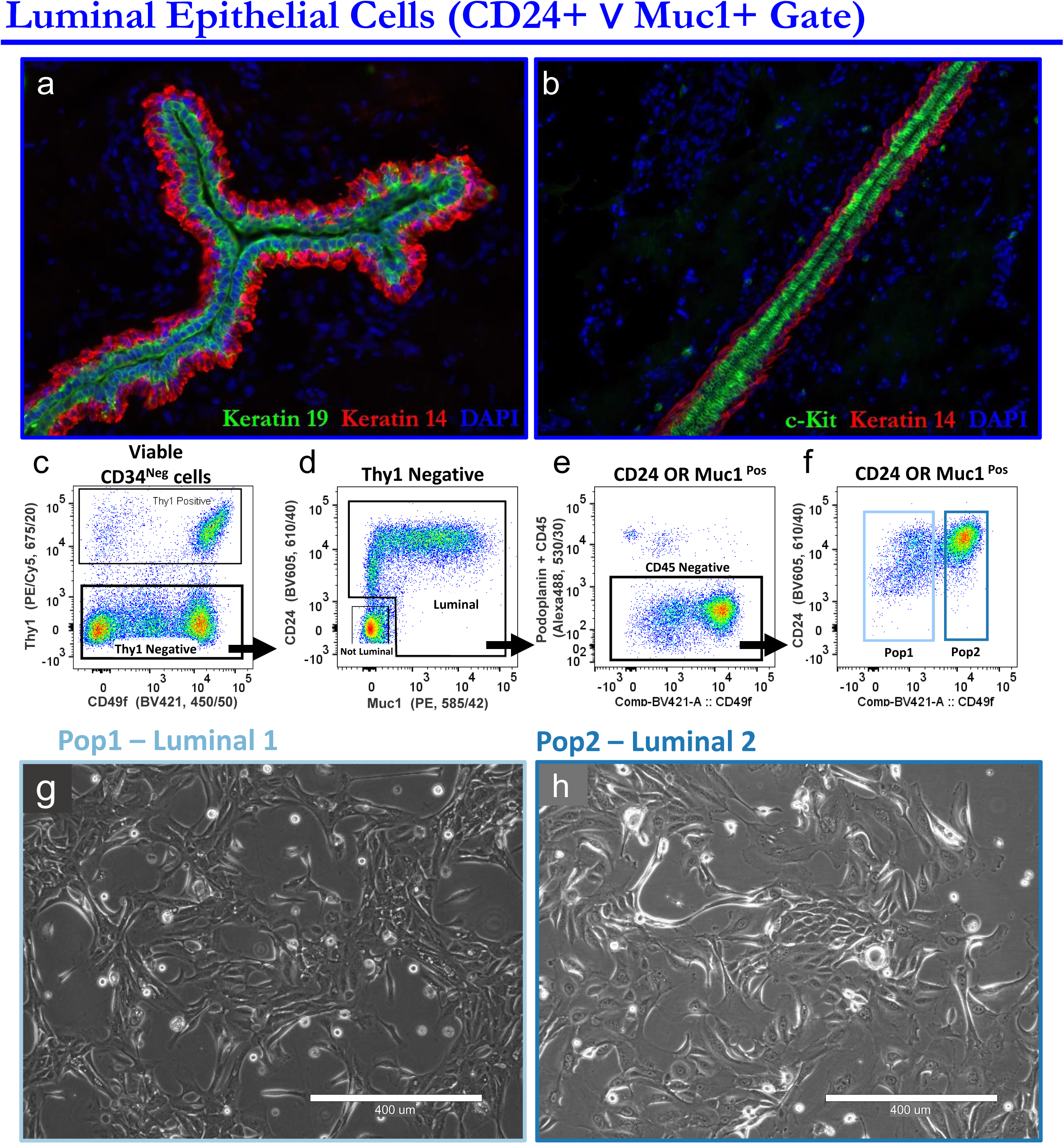
Identification and validation of the luminal cell types. **(a,b)** Normal breast tissues immunostained for luminal epithelial-associated proteins **(a)** keratin 19 and **(b)** c-kit, counterstained with keratin 14. **(c-f)** FACS gating strategy for isolating luminal epithelial cells: (c) CD34 negative cells (backgated for viability and single cells) were analyzed for Thy1 expression. (d) Luminal epithelial cells are present within the gated ‘Thy1 Negative’ fraction, identified by their expression of one, or both, luminal markers: CD24 and Muc1 (denoted by the inclusive disjunction operator, ‘∨’). (e) The resulting CD34^Neg^ Thy1^Neg^ CD24^Pos^ ∨ Muc1^Pos^ cell fraction (on the 8-color BD FACS machine, shown here) was further refined by gating out CD45^Pos^ leukocytes and focusing on the ‘CD45 Negative’ cells (Note: CD45Pos cells are gated earlier when using a FACS machine with additional channels). (f) Differential CD49f expression demarcates two distinguishable luminal cell populations within the CD45^Neg^ fraction: Pop1 (CD49f ^Neg/Low^, light blue box) and Pop2 (CD49f^Pos^, dark blue box). **(g)** Phase contrast images of Primary Pop1 (Luminal 1) and (h) Pop2 (Luminal 2) cultures (both Passage 2; 21 days post-seeding). Both cultures were initially seeded at 4700 cells/cm2 and sub-cultured at 32,500 cells/cm^2^ on bovine collagen I substrata, both of which (density and substrata) proved critical to successful culture of the luminal cells. To ensure purity, the cell populations were triple sorted (using conservative gating that avoided the transitional space between pop1 and pop2, gray dashed lines).

Although we found we could culture both luminal populations, there was a notable difference: Compared to their Pop2 counterparts, Pop1 luminal cells required a higher seeding density to establish a growing culture. When we tested colony-forming efficiencies by FACS sorting and seeding limiting numbers of luminal cells into collagen-coated 96 well dishes (Figure 17– figure supplement 2a), we found 83% of wells seeded with Pop2 cells (at the highest density) became confluent after 35 days (Figure 17– figure supplement 2a,b). In contrast, none of the wells seeded with Pop1 cells— at the same or lower densities (from 3 independent experiments)— ever produced large colonies. Although some individual Pop1 cells attached in these wells, a few produced small colonies of less than about ten cells each, but they did not exhibit the vigorous growth observed by Pop2 cells. These results seemingly confirmed prior claims that CD49f-null luminal cells lacked progenitor activity. However, we discovered that this was not the case. We had also seeded every other well in these experiments with irradiated fibroblasts—to serve as a non-confluent feeder layer. When we examined these neighboring wells, we found Pop1 cells grew dramatically, with half the wells reaching confluency (i.e., those seeded at the highest density; Figure 17 – figure supplement 2c). The support provided by the feeder layer extended to the lower seeding densities as well, for both Pop1 and Pop2 cells. Remarkably, 4% of wells seeded with just a single Pop2 cell produced clonal cultures. Still, the growth of Pop1 cells was striking and stood in stark contrast to the stagnant growth of those cultured without fibroblasts— and to prior claims that the CD49f-null ‘mature’ luminal cells do not have progenitor activity (Ibid). Our results demonstrate that they do, but it was not clear if stromal support was essential.

The progressive improvement in Pop2’s colony-forming efficiency, which occurred when we seeded them at higher densities (Figure 17—figure supplement 2b), made us wonder if Pop1 cells just needed an even higher seeding density than 1500 cells/well (4700 cells/cm^2^) to survive and thrive. We discovered this was indeed the case. When we seeded Pop1—and Pop2—cells at 15,000 cells/cm^2^ or greater (roughly 3x higher than the highest density in prior experiments), they all produced propagating cell lines (n=3 each). Along with the above experiments, these data provided evidence that Pop1 and Pop2 cells were functionally distinct and that both could establish *ex vivo* cell lines. Knowing where these different luminal cell types existed within the breast epithelium remained a vexing question, however.

The proportions of the two luminal types in our FACS-analyzed samples incidentally helped pinpoint their histological location. While we detected the two luminal populations in every breast sample (n=32 individuals), their relative abundance within the luminal cell compartment varied widely between individuals, ranging between 3% to 91% for Pop1 and 6% to 96% for Pop2. This unpredictable imbalance (i.e., the ratio of the two luminal cell populations) was notably the most variable feature among individuals—for any cell type (Figure 18a,b). A crucial clue to Pop1’s identity was uncovered when we stained and analyzed tissues for estrogen receptor. We noticed that one sample, N277, was teeming with ER^Pos^ luminal cells. Nearly every luminal cell in this sample stained positive for ERα (Figure 18c; Figure 18– figure supplement 1). This patient was notably taking TriNessa (norgestimate and ethinyl estradiol tablets) for contraceptive purposes; however, we are uncertain if this contributed to the overwhelming abundance of ER^Pos^ luminal cells in this patient’s breast tissue. The staining was nevertheless quite striking. When we examined the tissue by FACS, we noticed a similar luminal-cell imbalance: We found that it had, by far, the most CD49f^Low/Neg^ (Lum1) cells than any other tissue we have ever tested, and it was indeed the sole outlier (Figure 18b). The correlation between the abundance of ER^Pos^ and Pop1 cells in this tissue, measured respectively by tissue immunofluorescence and FACS, was glaring. In this sample, nearly every luminal-cell expressed ER (determined by tissue staining) and consisted, almost entirely, of Pop1 luminal epithelial cells (determined by FACS). This connection hinted that CD49f^Low/Neg^ luminal cell population (Pop1) were, in essence, ER^Pos^ luminal cells.

**Figure 18.**
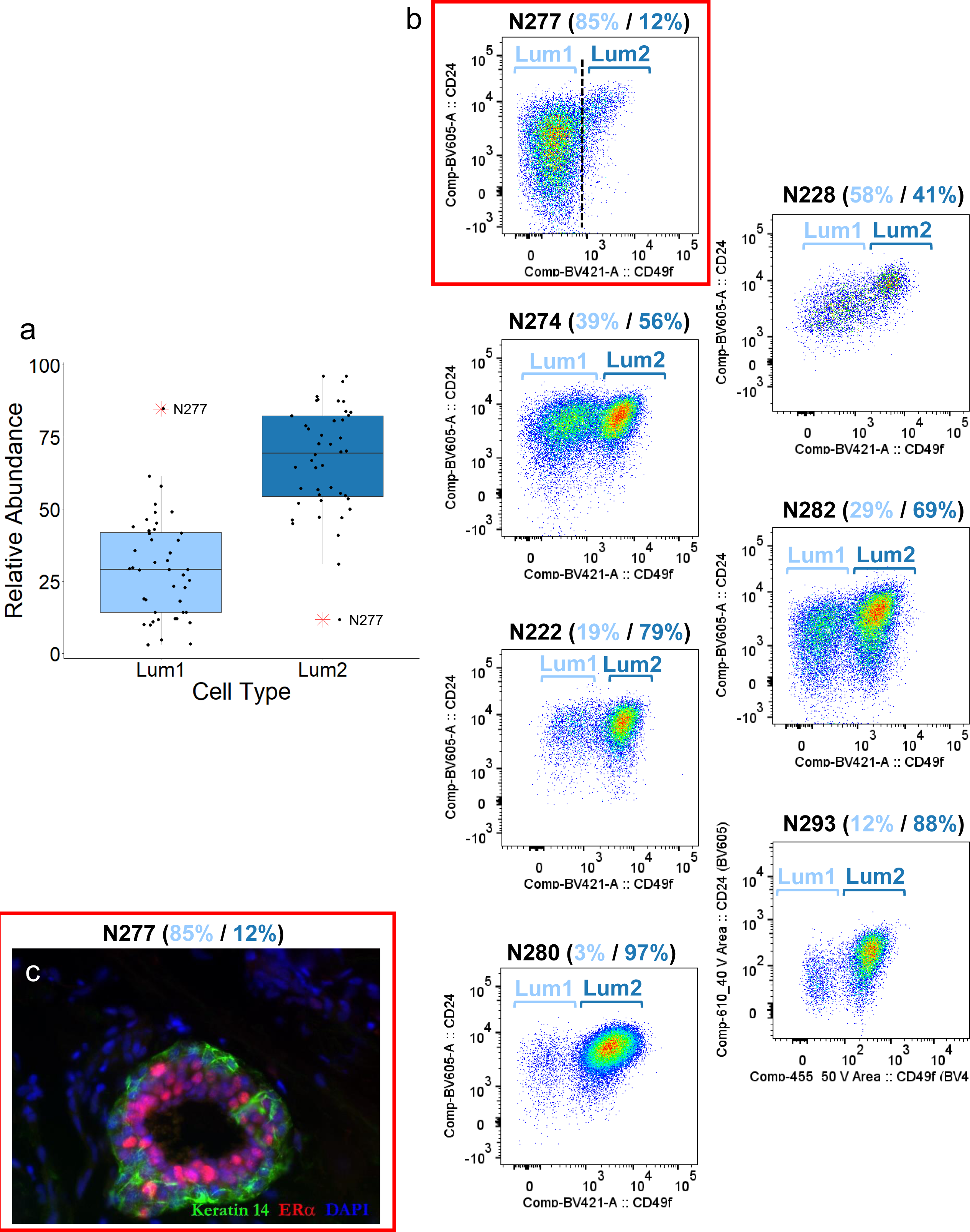
The relative abundance of Pop1 and Pop2 cells varies widely among individuals. **(a)** Relative abundance of CD49f ^Low/Neg^ Luminal 1 (Lum1) and CD49f^Pos^ Luminal 2 (Lum2) cells, measured by FACS, within the luminal cell compartment (all CD24/Muc1 staining breast cells). Displayed are 44 separate analyses of randomly selected normal breast tissues (reduction mammoplasty tissues) from 30 different individuals. The average age of the women was 27± 6 years (sdev), with ages ranging between 16 to 40 years**. (b)** FACS scatterplots of the luminal cell fractions, showing select samples with an uncommonly high proportion of Lum1 cells (N277, N228); nearly equivalent proportions of Lum1 and Lum2 cells (N274, N282, N222); and uncommonly high proportion of Lum2 cells (N293 and N280). (c) Immunofluorescent immunostaining of N277 breast tissue for estrogen receptor alpha indicated that the high proportion of Lum1 cells in this sample coincided with an uncommonly high proportion of ER-staining luminal cells (N277 outlined in red).

To explore the strong parallel relationship between the abundance of ER^Pos^ cells and FACS-defined CD49f^Neg^Pop1 cells, we co-stained breast tissues with both ER and CD49f antibodies. The results were clear: CD49f (alpha-6-integrin) was strongly expressed along the basal side of myoepithelial cells and nicely traced the outline of the basement membrane (Figure 19). The luminal cells were indeed split by those that stained for CD49f and those that did not. For those that did, expression was most intense along the cells’ basolateral membrane—and these cells did not express ER. Of the luminal cells that did express ER, we found they did not express CD49f – which we verified by confocal optical sectioning (Figure 19– video supplement 1). These results showed that, between the two luminal epithelial cell types, CD49f and ER expression was mutually exclusive. Therefore, we concluded that the FACS-defined CD49f^Low/Neg^ Pop1 cells are, in fact, ER^Pos^ luminal cells. Immunostaining tissues for markers differentially expressed between Pop1 and Pop2 cells (identified by RNA-seq analysis) solidified this conclusion. Staining results included the mutually coherent expression of estrogen receptor α and progesterone receptor in Pop1 cells (Figure 19 – figure supplement 1); and mutually exclusive expression between Pop1 and Pop2 of estrogen receptor α and either: keratin 15 or c-kit proto-oncogene (Figure 19 – figure supplement 2,3)—staining that was consistent with our RNA-sequencing results and with prior reports^45, 46^.

**Figure 19.**
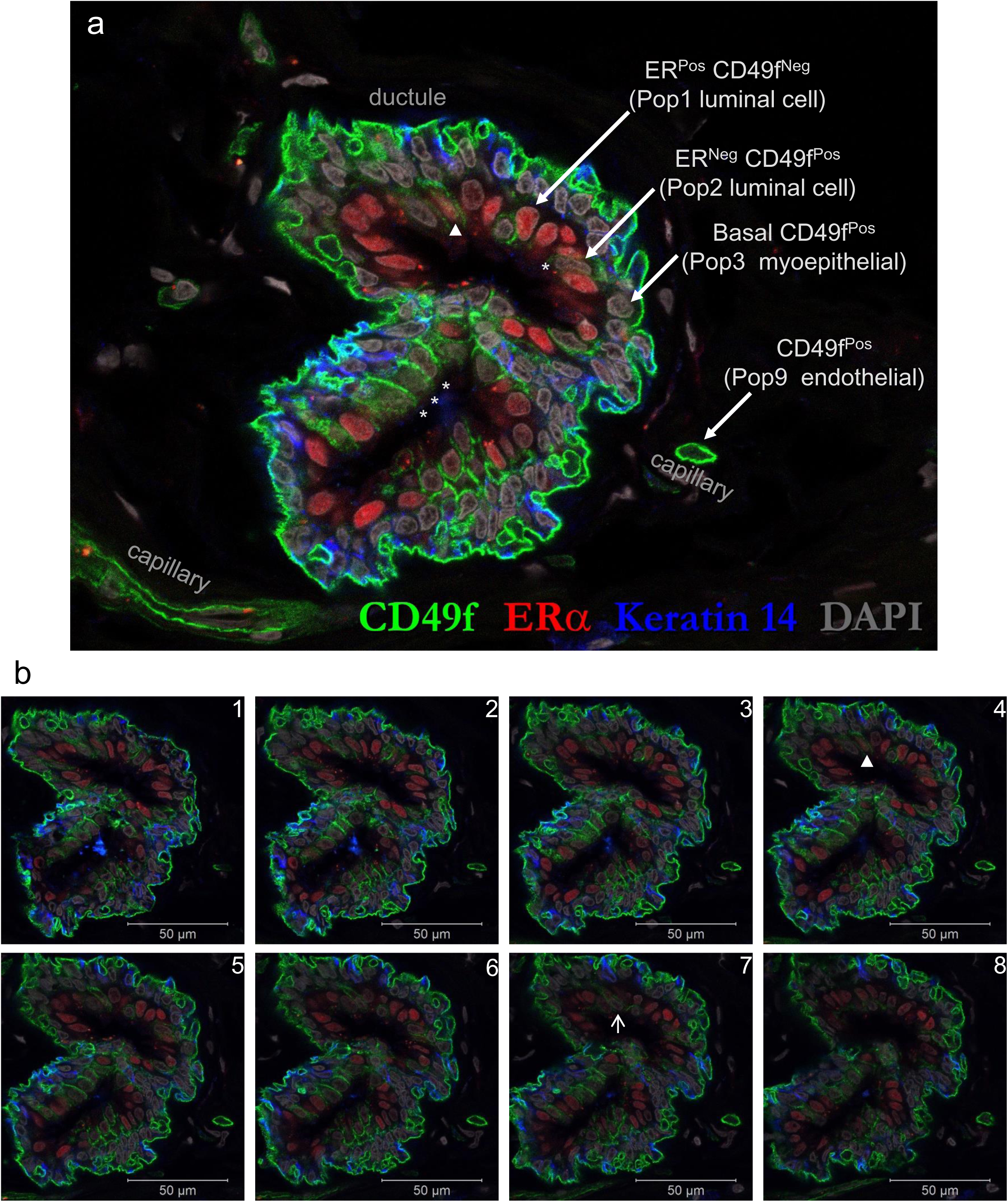
Pinpointing the histological location of Pop1 and Pop2 luminal cells. **(a)** Confocal image of normal breast tissue immunostained with ERα (red) and CD49f (green). CD49f (α6 integrin) was present within endothelial cells of small capillaries and along the basal side of myoepithelial cells, which is consistent with CD49f^Pos^ staining of these cell types in FACS experiments. Tissue staining also revealed CD49f expression along the basolateral membranes within a fraction of luminal epithelial cells, which is also consistent with FACS results, as this marker defines the Pop2 luminal cell gate (CD49f^Pos^). The tissue staining also showed that these same CD49f-expressing luminal cells did not express ERα. Four such cells are labeled with an asterisk, ‘*’. Estrogen receptor was instead confined to CD49f^Neg^ luminal cells that, by FACS, defines Pop1 luminal cells (CD49f^low/neg^). **(b)** Adjacent confocal planes (b1-8) illustrate how the cellular origin of ERα and CD49f staining is sometimes obscured by the proximity of cell types in tissues. For example, one ERα-stained luminal cell initially appeared to express CD49f (marked by ‘▾’ in a and b4), however, upon examination of adjacent Z-stacked planes, the CD49f staining was instead traced to an adjacent ER-negative luminal epithelial cell (marked by ‘↑’ in b6&7).

Both luminal cell types compose, on average, about 26% of all cells in the breast, ranging between 5 and 64% (26±16%), making them collectively the second most abundant cell type in the tissue–just behind myoepithelial cells (27%, Figure 9-table supplement 1). ER^Neg^ luminal cells (Pop2) were almost always the most prevalent of the two (38/44 cases), composing about 67% of all the luminal cells on average. However, as previously noted, the relative proportion of the luminal cell fractions varied widely between individuals. Why this is so is currently unknown. All FACS-analyzed samples were from assumedly premenopausal women (ages 16-40). Although we did not have the necessary data to stage each individual’s position in the estrous cycle, we predict that cycling estrogen and progesterone hormones influence the cell phenotype and the balance of the two luminal cell types.

We have demonstrated in this section that CD24 and Muc1, along with CD49f, resolve two luminal cell types within the Thy1^Neg^ FACS gate. The addition of these two cell types brings the running tally to eight cell types identified so far—all of which we have successfully cultured. The remaining Thy1^Neg^ cell types, which do not express CD24 or Muc1, compose the final node that we called the ‘Not-Luminal’ or ‘Null’ Gate.

### Node5: Null Gate (Leukocytes, Erythrocytes, Vascular Smooth Muscle Cells, and Pop4 Epithelial)

#### Leukocytes

Of the remaining cell populations to be isolated, those we had anticipated from the outset—and had still not sorted— were the leukocytes and erythrocytes (white and red blood cells). Although the leukocytes comprise a broad range of related cell types with distinct functions, resolving each subpopulation was not our priority. To isolate the bulk leukocyte population, we relied on the frequently used pan-leukocyte marker, CD45. However, we first needed to overcome a technical issue: The other markers in our panel were already occupying each of the fluorescent detectors on our 8-color FACS machine. These were: TO-PRO-3, CD34, CD49f, Podoplanin, Thy1, CD10, Muc1, and CD24. To add CD45, we needed to double-up and detect two markers in a single fluorescent channel. This strategy is possible if the two antibodies do not bind to the same cells and if the different cell types that they do bind can be resolved in another dimension by other antibodies in the panel. After a series of tests, we elected to use the same fluorescent tag (Alexa Fluor 488) for both CD45 and podoplanin. This strategy worked nicely, as adding CD45 to the panel cleanly separated the leukocyte population, which we numbered Pop10 (Figure 20a). This CD34^Neg^ Thy1^Neg^ CD24^Neg^Muc1^Neg^ CD45^Pos^ cell fraction exhibited a unique low side scatter FACS profile typical of leukocytes (Figure 20– figure supplement 1). After RNA-sequencing, we found Pop10 indeed differentially expressed the gene encoding the CD45 antigen, protein tyrosine phosphatase receptor (PTPRC, Figure20b), as well as many other leukocyte-specific markers (Figure 20– figure supplement 2). Collectively, these data validated our sorting strategy. Of course, if a cytometer has enough fluorescent detectors to accommodate each marker, doubling up antibodies would not be required, and one could instead add CD45 to an open channel. Using this alternative approach on a different FACS machine—one with 14 available channels (SONY SY3200)—we have successfully and repeatedly replicated the above results, demonstrating the rigor of the FACS strategy when tailored to the machine (Figure 9-figure supplement 1&2).

**Figure 20.**
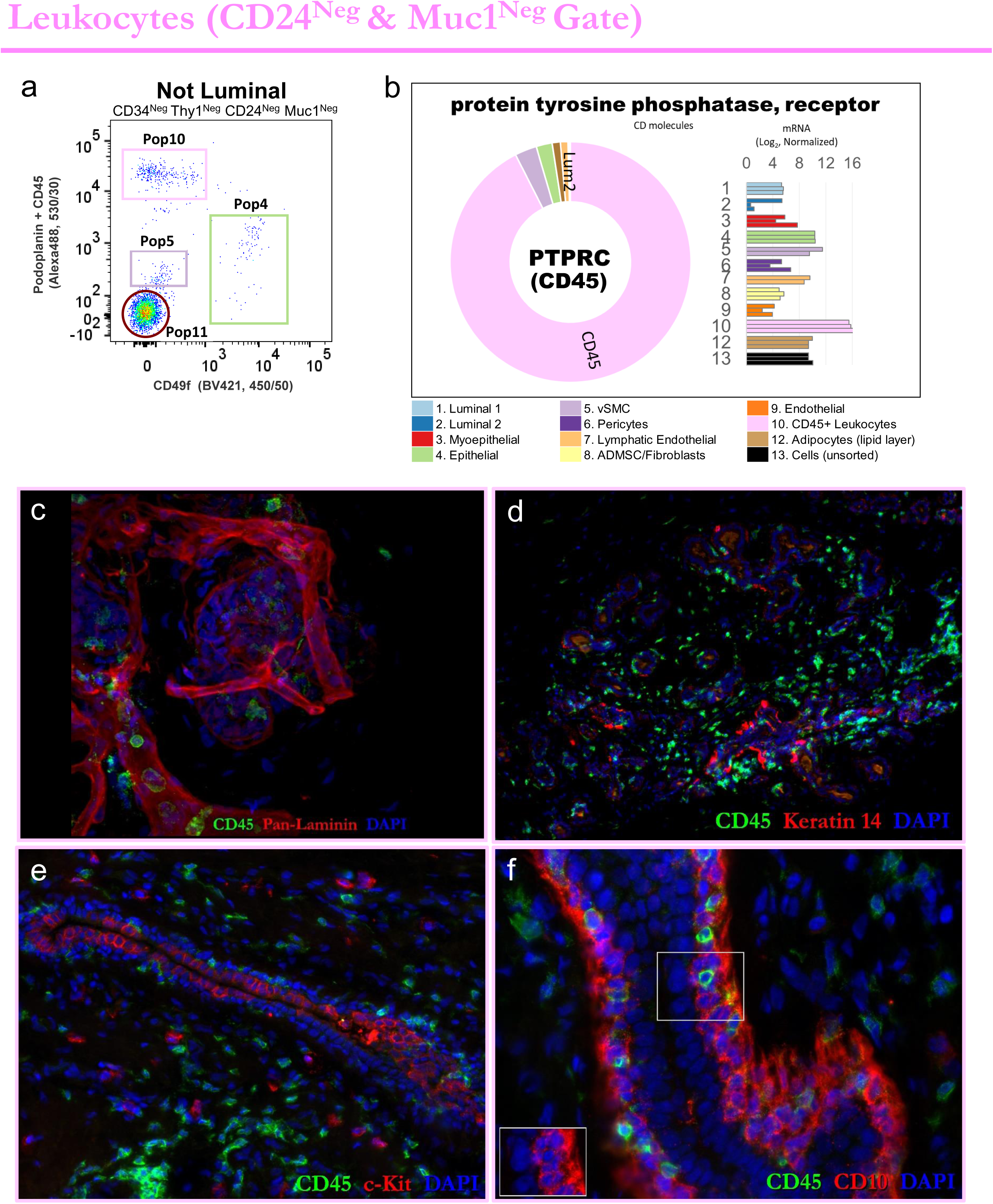
Identification and validation of Leukocytes. **(a)** FACS gating strategy for isolating leukocytes and remaining breast cell types. CD34, Thy1, CD24, and Muc1 negative cells (also backgated for viability and single cells) were analyzed for CD45/Podoplanin and CD49f expression. CD45 and podoplanin antibodies were labeled with the same fluorophore (Alexa488) because all 8 fluorescent channels on this FACS machine were in use. This analysis revealed four cell populations: Pop10 (leukocytes), Pop11 (red blood cells –and debris), and two additional, yet unknown, populations. We designated these two as Pop4 and Pop5 (which we later identified to be a rare epithelial cell type and vascular smooth muscle cells, respectively). **(b)** Transcript levels of PTPRC (the gene encoding CD45) within uncultured FACS-purified cell types, as determined by RNA-sequencing. As expected, this pan-leukocyte marker was clearly and differentially expressed by Pop10 leukocytes. **(c-f)** Leukocytes within breast tissues, revealed by staining for CD45 and (c) pan-laminin, to reveal white blood cells (WBCs) within- and outside the vasculature; (d) keratin 14, to reveal WBCs proximity to the epithelium; (e) c-kit, to reveal mast cells (and Pop2 luminal epithelial cells); and (f) CD10, which reveals WBCs that have transmigrated into the epithelial compartment and are found situated among CD10^Pos^ myoepithelial cells.

After applying the FACS strategy to our collection of breast tissues, we found that the proportion of leukocytes ranged between 0 and 11%, composing 2.1±2% of all cells in the tissue on average. Leukocytes were among the least abundant cell types in the breast (Figure 9-table supplement 1). Although we did not design our sorting strategy to identify each leukocyte type, we did observe Kit^Pos^ mast cells, CD56^Pos^ natural killer cells, and others when we immunostained tissues with epithelial markers that cross-reacted with these leukocyte types, i.e., CD117 (Kit), CD56, and CD10 (Figure 6– figure supplement 6,7; Figure 17b; Figure 14i). When we stained tissue sections, we found leukocytes throughout the tissue—inside and outside blood vessel lumina (Figure 20c; Figure 20– video supplement 1). Leukocytes were concentrated near epithelial structures and were especially prevalent within TDLUs (Figure 20d,e; Figure 20– figure supplement 3a). They were most frequently found in the stroma but were also conspicuously present on the epithelial side of the basement membrane, among the luminal and myoepithelial cells, and within the lumen of ducts and ductules (Figure 20f; Figure 20– figure supplement 3b). Co-staining tissues for CD45 and CD68, CD3e, CD20, or Kit demonstrated the vast majority of extravascular leukocytes were CD68^Pos^ macrophages, although a small fraction of Kit^Pos^ mast cells, CD3e^Pos^ T-cells, and CD20^Pos^ B-cells were also present (Figure 20– figure supplement 3c-i, figure supplements 4-10).

To ensure each of the other FACS-sorted cell types were free of leukocytes, we added a CD45^Neg^ gate to each of their respective isolation strategies, and these gates were included in the RNA-sequenced samples (Figure 9-figure supplement 1&2). We then returned to the final FACS plot in Figure 20a to investigate the remaining cell populations. As expected, red blood cells (erythrocytes) were one of these.

#### Erythrocytes

Erythrocytes do not express any of the markers within our staining panel. As such, they occupy the left lower quadrant of the final FACS scatter plot; i.e., they are CD34^Neg^ Thy1^Neg^ CD24^Neg^ Muc1^Neg^ Podoplanin^Neg^ CD45^Neg^ cells (Figure 20a). We named this null-staining cell fraction ‘Pop11,’ and found it was the only population that produced a red cell pellet after being centrifuged (Figure 21a). Microscopic examination of Pop11 confirmed the presence of biconcave red blood cells (Figure 21b). Small debris was often present as well, which is expected in a marker-negative gate such as this. Because debris particles, precipitates, and large salt crystals can sometimes trigger the FACS machine into falsely recognizing them as cells (FACS events), we filtered all buffers and collection media through 0.2um polyethersulfone (PES) filters prior to using them for cytometric analyses. We thoroughly rinsed the cells and strained them through 100 and 40um cell strainers—multiple times—prior to analysis, which reduced debris and false-positive events. When we seeded Pop11 into culture, nothing grew as expected -and RNA isolations did not yield any appreciable RNA. We, therefore, concluded Pop11 contained red blood cells and small debris particles. Additional purification using an erythrocyte-specific marker, e.g., CD235a, was not needed as this fraction was not further used or analyzed.

**Figure 21.**
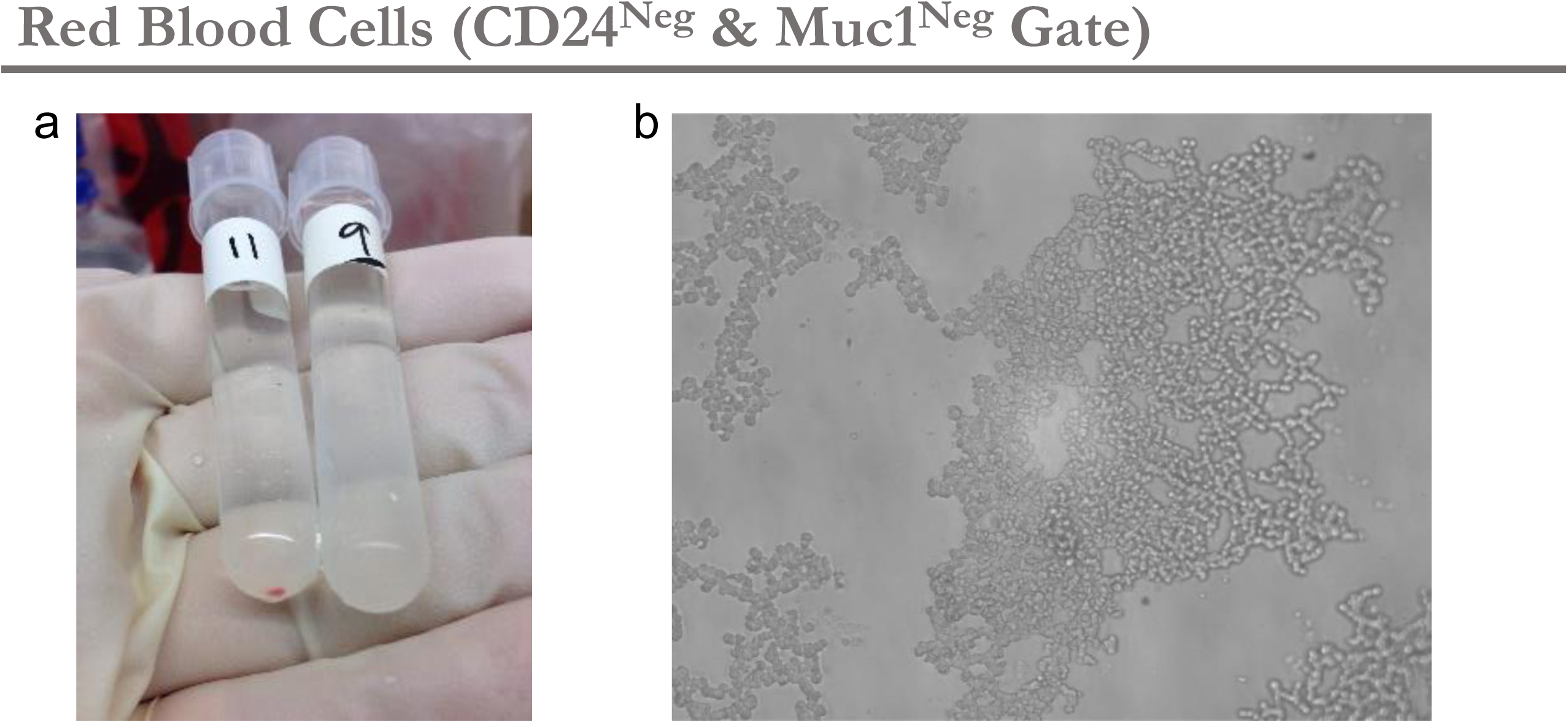
Identification and validation of Red Blood Cells. **(a)** As illustrated in the preceding figure (Fig.20), there was a single cell fraction that did not bind any of the eight antibodies used in our FACS panel. It was designated Pop11. This population was conspicuously the only cell fraction that produced a red cell pellet upon centrifugation, which indicated the presence of erythrocytes. **(b)** Phase-contrast microscopy confirmed Pop11 indeed consisted of biconcave erythrocytes and unidentifiable debris.

#### Population 5: Vascular Smooth Muscle Cells

When we viewed the final FACS plot of the ‘Not Luminal’ node (Figure 20a) after identifying leukocytes and erythrocytes, we were surprised that two cell populations remained. The first of these, we named ‘Pop5.’ This fraction was not well-defined but was located adjacent to the marker-negative RBCs on the final FACS plot (Figure 20a). Attempts to culture Pop5 cells were mainly unsuccessful, as single cells would sometimes attach to the dish and only a few cultures were established. Pop5’s lack of growth distinguished it from the other non-blood cell types, all of which we could successfully and routinely culture. Therefore, revealing Pop5’s identity required RNA-sequencing, and one of the earliest clues was its unique expression of 5-hydroxytryptamine (serotonin) receptor 1B (HTR1B). This gene was one of 13 uniquely expressed by Pop5 (at a 16x stringency level, Figure 22a). Pop5’s expression of HTR1B was notable because we recalled that Pop6 pericytes uniquely expressed two genes at this stringency, and one was HTR1F—an isoform of HTR1B (Figure 16 – figure supplement 3). Although circumstantial at the time, this association was our first clue that Pop5 and Pop6 cells were related, causing us to suspect Pop5 cells were vascular smooth muscle cells (vSMCs). Pericytes and vSMCs are both periendothelial cells and coexist on a related developmental spectrum^42^. A key feature of vSMCs is their strong expression of alpha-smooth muscle actin (αSMA, Figure 22b), which we discovered was expressed at high levels by Pop5 cells (Figure 22c; Figure 22– figure supplement 1) and was significantly higher than pericytes (adj. *p-*value= 1.03×10^-4^). We also discovered Pop5 differentially expressed additional vSMC-associated genes, including those encoding chymase and calponin (Figure 22– figure supplement 2). Other genes differentially expressed between Pop5 and Pop6 cells included melanoma cell adhesion molecule (MCAM, a.k.a. CD146) and CD36, the latter expressed more highly by Pop6 pericytes; and MCAM being highest in Pop5 (Figure 22d,e). When we immunostained tissues for these two markers, we found vSMCs and pericytes indeed displayed this expression pattern, further strengthening our conclusion that Pop5 cells were vSMCs (Figure 22f,g, Figure 22– figure supplement 3,4).

**Figure 22.**
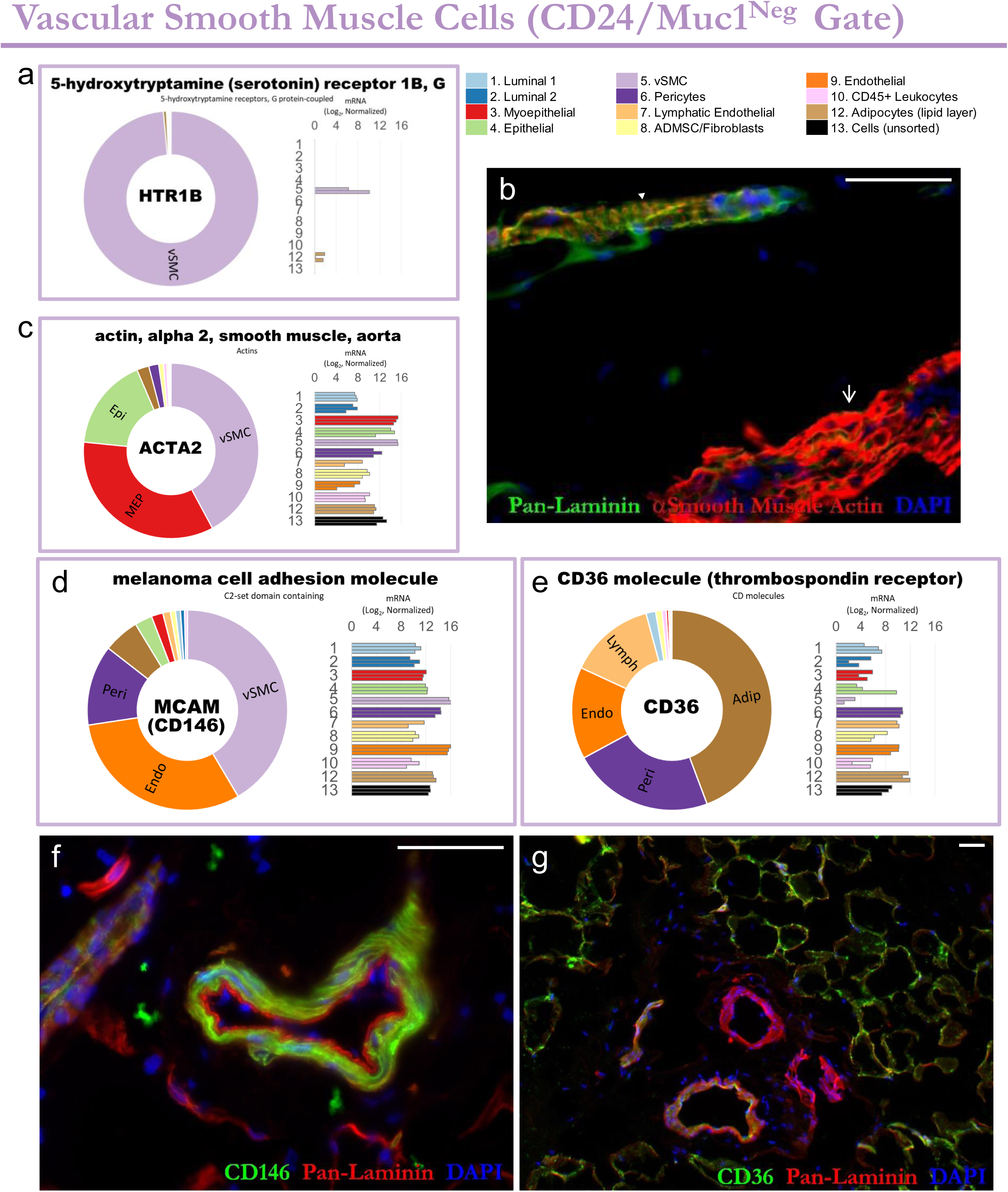
Identification and validation of Vascular Smooth Muscle Cells. **(a)** Transcript levels of HTR1B (the gene encoding 5-hydroxytryptamine receptor 1B) within uncultured FACS-purified cell types, as determined by RNA-sequencing. HTR1B is an isoform of serotonin receptor, similar to the HTR1F isoform, which was uniquely-expressed by breast pericytes. **(b)** Breast tissue immunostaining demonstrates alpha smooth muscle actin expression within arteriolar vascular smooth muscle cells (‘▾’, top left); and within ductal myoepithelial cells (‘↑’, bottom right). **(c)** ACTA2 (the gene encoding alpha smooth muscle actin) is differentially expressed between Pop5 vSMCs (light purple) and Pop6 Pericytes (dark purple). **(d,e)** Transcript levels of (d) MCAM (CD146) and (e) CD36 in uncultured FACS-purified cell types. **(f,g)** Normal breast tissue immunostained with (f) CD146 and (g) CD36.

Nevertheless, there was one more piece of supporting data: The primary marker separating Pop5 from Pop6 in the FACS strategy was thy1—which we used to identify and isolate Pop6 pericytes (and myoepithelial cells). Naturally, we were curious to know whether vSMCs in the tissue lacked thy1. We found they did—via thy1 immunostaining of breast tissues (Figure 22– figure supplement 5).

After analyzing each breast tissue sample by FACS, we found Pop5 vSMCs were relatively scarce in our preparations, constituting only 0.2%±1% of all cells in the tissue. Thus, vascular smooth muscle cells are the second least abundant cell type in the breast, slightly more prevalent than lymphatic endothelial cells. After our successful identification of Pop5, only one population in the final FACS plot remained — and it proved to be the most puzzling of them all.

#### Population 4: ‘Epithelial’ Cells

A single population stood apart from all other cell types in the ‘Not Luminal’ node, predominantly due to its expression of alpha-6 integrin (CD49f). We designated this fraction, Pop4 (Figure 20a). Its expression of CD49f was peculiar because we believed, based on our tissue immunostaining, that we had accounted for all CD49f expressing cell types in the tissue (Figure 10 – figure supplements 2&5). The previously characterized CD49f-expressing cell types included: a) ER^Neg^ luminal cells (Pop2—gated by CD24/Muc1), b) MEPs (Pop3—gated by Thy1), and c) both endothelial cell types (Pops 7&9—gated by CD34, Figure 9). Nevertheless, despite removing these cell types in the preceding gates, a small number of CD49f^Pos^ cells remained, which formed the Pop4 fraction. This population persisted even when we used extreme gating of the above markers to ensure that weakly-stained cells at each gate’s border were not slipping through, which would inadvertently divert them to the Pop4 gate.

After analyzing many samples, we found Pop4 cells to be quite rare, as they comprised only 0.6% of all cells in the breast (0.6±2%, median±sdev, Figure 9-table supplement 1). However, in contrast to Pop5 cells found in the same gate, Pop4 cells established growing cultures when seeded into culture. The phenotype of Pop4 cells was most similar to MEPs, but cultures often contained morphologically diverse phase-contrast phenotypes that were unlike homogeneous MEP cultures (Figure 23a,b; compare to Figure 14 –Figure supplement 3). Based on this phenotype, we determined Pop4 cells were epithelial. There were also several lines of evidence indicating Pop4 cells were distinct from luminal and myoepithelial cells. For example, in low-density 3D Matrigel cultures, Pop4 cells produced structures consisting of keratin 19 positive cells surrounded by a layer of cells co-expressing both keratin 14 and 19 (Figure 23c,d). In contrast, pure luminal and myoepithelial cells homogenously expressed keratin 19 or 14, respectively (Figure 23e,f). The mixed cell phenotypes persisted when we seeded Pop4 cells onto a 2D collagen substratum—at clonal density— that produced cultures that heterogeneously expressed Muc1, suggesting they possess bipotent progenitor activity (Figure 23– figure supplement 1a-c). Lastly, Pop4 cells demonstrated a unique beta-1 integrin (CD29) expression pattern distinct from all other CD49f expressing cell types (Figure 23g; Figure 23– figure supplement 1 d-g). How Pop4 cells related to the other epithelial cell types was a central question we aimed to clarify through RNA-sequencing.

**Figure 23.**
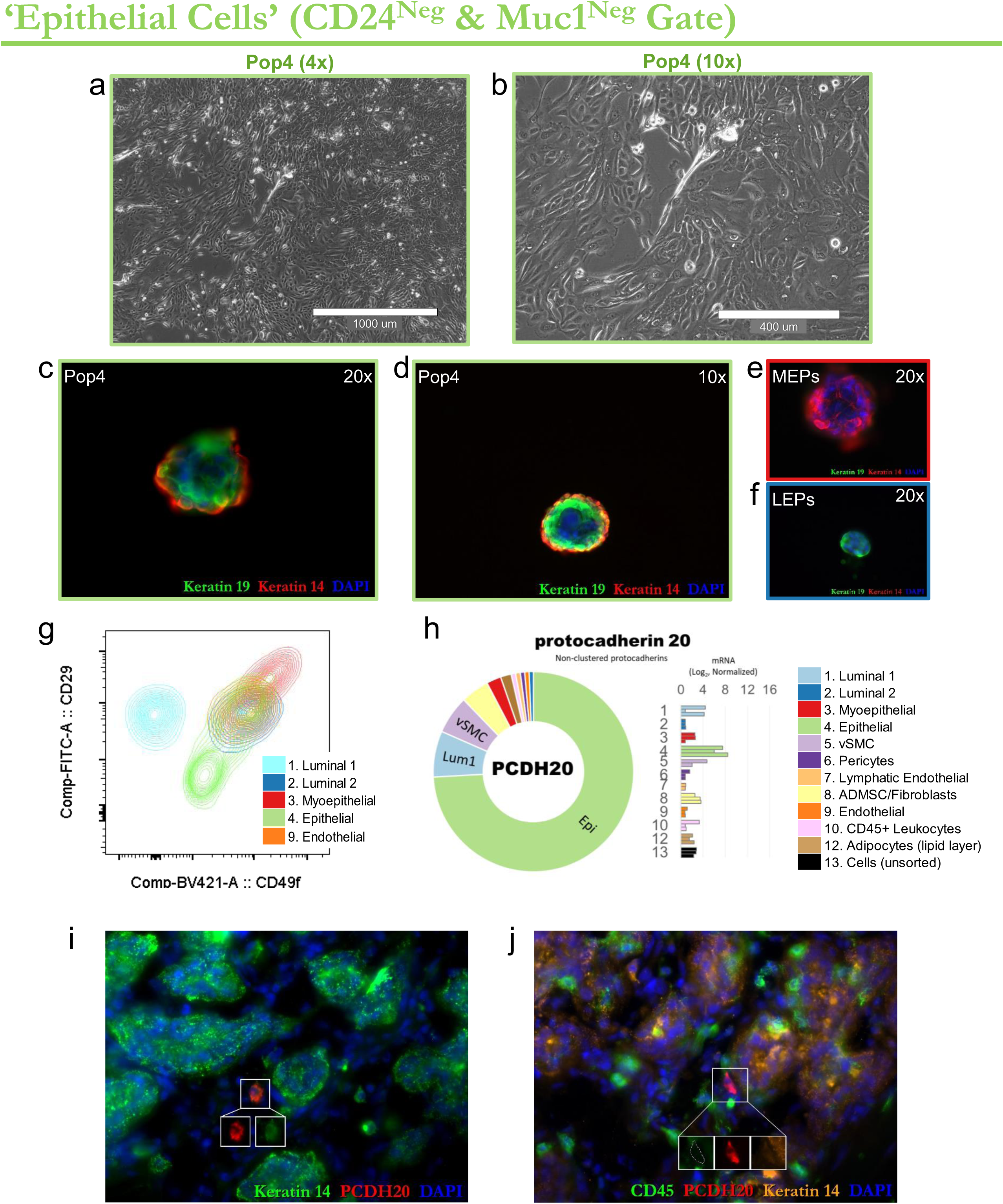
Identification and validation of ‘Pop4’ epithelial cells. **(a,b)** phase-contrast images of primary Pop4 epithelial cultures imaged with (a) 4x and (b) 10x objectives. **(c-f)** 3D matrigel cultures of (c,d) pop4 epithelial cells, (e) Pop3 MEPs, (f) Pop2 Luminal epithelial cells immunostained for keratin 14 and 19. **(g)** Flow cytometry scatter plot of β1-integrin (CD29) and α6-integrin (CD49f) in the five different CD49f-expressing breast cell types. **(h)** Transcript levels of protocadherin20 (PCDH20), a gene uniquely-expressed by Pop4 cells. Transcripts were measured in freshly sorted (uncultured) cell types by RNA-sequencing. Normalized mRNA values (rlog) are provided on log2 scale (bar graph of each biological replicate), as well as on a linear scale (donut graph of median value), each color-coded by cell type (Pop4 epithelial cells are green). **(i,j)** Pop4 epithelial cells within breast tissues, revealed by PCDH20 and keratin 14 co-staining, and absence of pan-leukocyte maker CD45 staining.

Unsupervised hierarchical clustering of transcripts from each cell population confirmed Pop4’s distinct epithelial nature, as we found Pop4 clustered most closely with MEPs. However, for a set of genes, Pop4 diverged and shared a more common expression pattern with luminal cells, highlighting the distinctiveness of Pop4 cells (Figure 23– figure supplement 1 d-h). Helping reveal the location of Pop4 epithelial cells in the breast was the transcriptional analysis that identified a transcript that Pop4, and no other cell type, expressed. This gene was protocadherin 20 (PCDH20, Figure 23h). When we stained breast tissues with a protocadherin 20 antibody, we found rare pcdh20-staining cells at a frequency of 1-3 cells per section. Surprisingly, these pcdh20^Pos^ cells resided within the stroma of TDLUs, near the basement membrane, but outside the epithelial compartment (Figure 23i). The lack of CD45 co-expression confirmed that these cells were not leukocytes (Figure 23j). Instead, we found they co-expressed keratin 14, establishing that these rare pcdh20^Pos^ stromal cells were indeed epithelial, consistent with the primary culture phenotype of Pop4 cells.

With the identification and characterization of Pop4 epithelial cells, we have now defined all cell populations in the ‘Null’ gate. Future isolations of Pop5 vSMCs and Pop4 epithelial cells would benefit from additional redundant markers to ensure their purity. Our characterization of the four cell types identified within this node (null gate) brings the running tally to twelve identified and purified breast cell types, leaving no other cell population for us to describe and thus concludes our isolation scheme.

## Summary

The number and different types of cells composing the breast has been clouded with uncertainty, as has been their origins and relationships to each other. We are not always sure how best to identify these different cell types, know where in the tissue they are located, how they communicate with each other and are maintained, or what all their respective functions may be. To these questions and more, we have some answers; but not all are sufficient or complete.

Since the emergence of methods permitting the outgrowth of cells from normal breast tissues in the late 1970s^47, 48^, the field’s most widely adopted primary cell models have essentially represented only two cell types. The breast, however, is an intricate structure whose function relies on a symphony of many different specialized cell types. Understanding how these cells interact—and knowing the consequences of their interactions—is essential to developing necessary insights into the tissue and the processes that go awry in malignancy. There is thus an urgent need for more context-specific models to study reciprocal interchanges between the cell and the ECM—and among other cell types^49^. Yet, despite this widespread acknowledgment, a method to comprehensively identify, isolate and culture the spectrum of cell types composing this elaborate tissue has not emerged.

To clarify the types of cells in the breast and build a solid foundation for future work, we have meticulously examined an extensive collection of normal breast tissues, microscopically and by flow cytometry. This article is the first in a pair dealing with the cellular composition of breast tissues. We present a thorough transcriptomic analysis in an adjoining article, where we identify and detail features that define each breast cell type identified herein. Although not required to interpret those transcriptomic analyses, it is advantageous to understand breast architecture, cellular composition, and considerations that ensured the purity and identity of each cell population. These topics, details, and an atlas of immunostaining results are presented here, along with primary cell models and preliminary RNA-sequencing data of markers that collectively helped identify and confirm cell identities.

To develop a comprehensive sorting strategy, we sought high-resolution markers and used redundancy rigorously to identify cell types (Figure 9 & supplements). We demonstrate the importance of this approach and show that many luminal epithelial cells would have otherwise been inappropriately sorted into the myoepithelial fraction had this not been discovered and alternative strategies employed (Figure 7 & Supplement 7-1). Analysis of FACS plots and immunostaining identified CD49f^Neg^ and CD49f^Pos^ luminal cells as ER^Pos^ and ER^Neg^ cells, respectively, and we show the proportion of these different luminal cell types varies considerably between individuals (Figures 17, 18, & 19, and supplements). Furthermore, we demonstrate that primary cultures of these different luminal cell fractions, though exhibiting different seeding requirements, can both be propagated in culture without the need for genetic manipulations or extraordinary effort (Figure 17g,h; Figure 17 supplement 2b,c). We have also identified a new and unusual epithelial cell with unknown functions (Pop4), which does not express CD24, Muc1, or Thy1—respective markers of mature luminal and myoepithelial cells (Figure 9,23 and supplements). Finally, we have identified and described methods for sorting and culturing perivascular pericytes and lymphatic and endothelial cells, that along with the other described models, significantly expand the repertoire of available breast cell models that may be employed. Now complete, our FACS isolation strategy resolves twelve distinct breast cell types— potentially more if refined to include the array of leukocyte populations residing in the breast (Figure 9 & 20 and supplements). We routinely sort and culture nine FACS-purified cell types from breast tissues using these procedures and methods. These are: ER^Pos^ and ER^Neg^ luminal epithelial cells, myoepithelial cells, a novel epithelial cell fraction, pericytes, fibroblasts, lymphatic and vascular endothelial cells, and adipocytes (Pops 1,2,3,4,6,7,8,9,&12; Figure 24). Several of these breast cell models are the first of their kind. The ability to purify and culture these breast cell types allows us to now explore the heterogeneity within each population (e.g., via single-cell RNA-sequencing) and investigate their functional properties—when cultured alone or in different combinations and contexts. We predict that the models, procedures, and concepts submitted herein will facilitate clarity and a deeper understanding of the types and properties of cells composing the breast. These tools should advance our ability to explore and define the rules governing normal cell and tissue function and may extend well beyond the breast to other tissues and organ systems to provide novel insights into disease processes.

**Figure 24.**
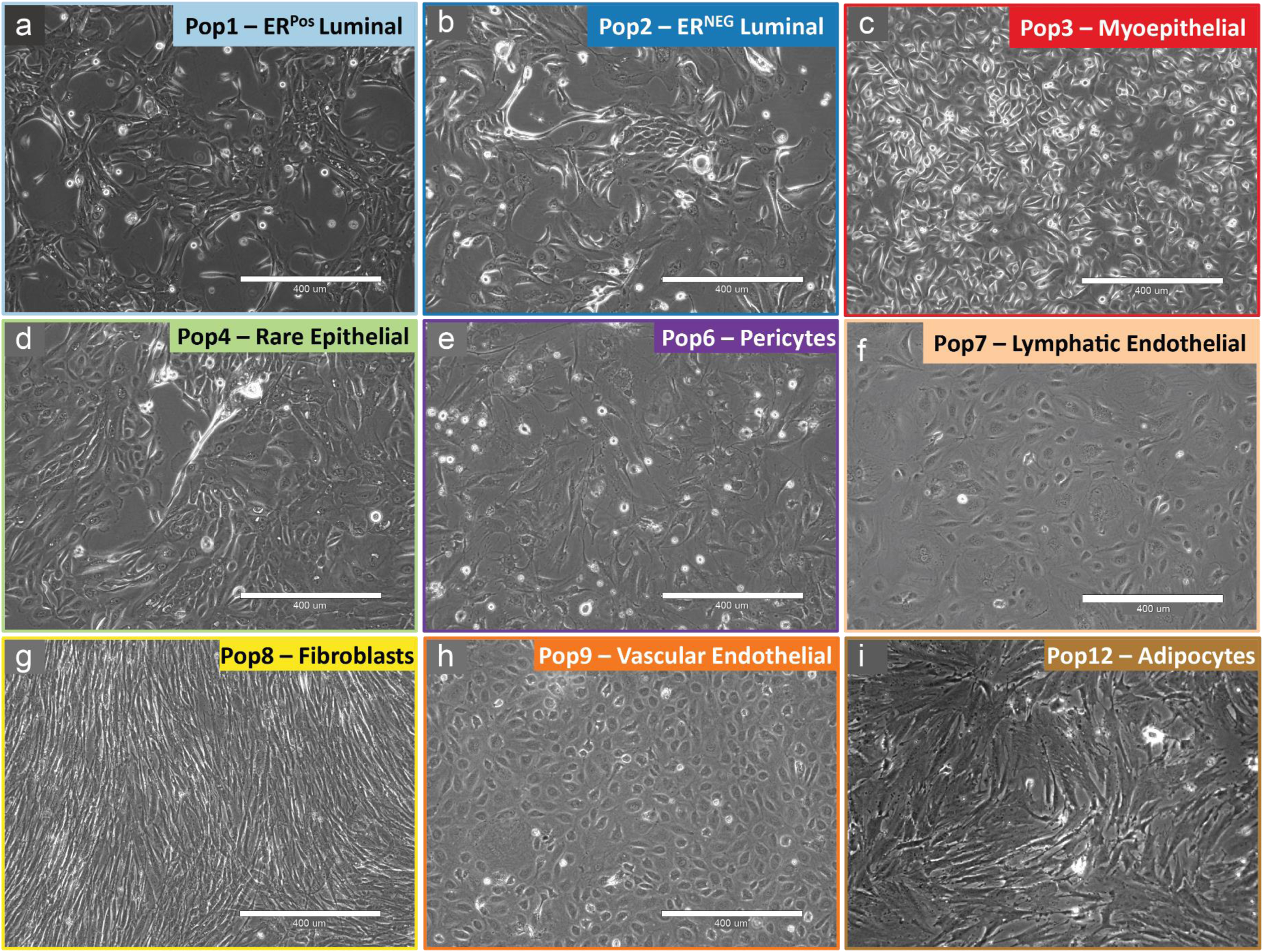
Primary cell models generated using the FACS strategy. Using the FACS isolation strategy detailed herein, cells were sorted and used to create primary cultures of a) Pop1 ER^Pos^ Luminal epithelial cells, b) Pop2 ER^Neg^ Luminal epithelial cells, c) Pop3 Myoepithelial cells, d) Pop4 Epithelial cells, e) Pop6 Pericytes, f) Pop7 Lymphatic Endothelial cells, g) Pop8 Adipocyte-derived Mesenchymal Stem Cells (Fibroblasts), h) Pop9 Vascular Endothelial cells, and i) Pop12 Adipocytes. Scale = 400μm.

## Methods

### Tissue acquisition and processing

Breast tissues from reduction mammoplasties were obtained from the Cooperative Human Tissue Network (CHTN), a program funded by the National Cancer Institute. All specimens were collected with patient consent and were reported negative for proliferative breast disease by board-certified pathologists. The University of California at Berkeley Institutional Review Board and University of New Mexico Human Research Protections Office granted use of anonymous samples through exemption status, according to the Code of Federal Regulations 45 CFR 46.101. Upon receipt, several fragments (roughly 2cm^2^) were embedded in optimal cutting temperature compound (OCT) in tissue cassettes, flash-frozen in nitrogen, and archived at −80°C for later cryosectioning and staining. The remaining tissue, typically 20-100g, was thoroughly rinsed with phosphate-buffered saline and processed to organoids as previously described (gentle agitation method)^5^. Briefly, this included manually mincing and scoring tissues with a scalpel and incubating them overnight (12-18 hrs, 37°C) with 0.1% collagenase I (Gibco/Invitrogen) in Dulbecco’s Modified Eagle Medium containing 100 U/ml penicillin, 100 μg/ml streptomycin, and 100 μg/ml Normocin™ (Invivogen, San Diego, CA). The resulting divested tissue fragments (organoids) were collected by centrifugation (100g x 2 min) and either archived in liquid nitrogen (90%FBS+10% DMSO) or immediately processed to cell suspensions for FACS analysis. Cells used for RNA-sequencing were derived from fresh unarchived tissues (not previously archived in nitrogen).

### Antibodies

A list of antibodies and reagents used in this study is provided in Figure 5—Table Supplement 1. The table includes antibody clone designations, conjugations, isotype, supplier product numbers, dilution factors used, and citations to figures.

### Cryosectioning and Immunostaining

Immunofluorescence was performed on 10-80 μm cryosectioned tissue sections (and cell cultures) by fixing samples with 4% paraformaldehyde for 5 minutes at 23°C, followed by 4% formaldehyde / 0.1% saponin for 5 minutes at 23°C. Tissues were subsequently incubated for 20 minutes in wash buffer (0.1% saponin/10% goat serum in PBS), and incubated with primary antibodies diluted in wash buffer at the indicated antibody dilution ratios (Figure 5-Table Supplement 1). Samples were incubated at empirically determined times, typically overnight at 4°C. Following the primary antibody incubations, samples were washed and incubated with anti-mouse, anti-rabbit, anti-chicken or anti-sheep secondary antibodies, respectively conjugated with Alexafluor 488, 568, 594, or 647 (Thermo), diluted 1:400 in wash buffer. After 1 hour incubation at 23°C, samples were rinsed in PBS and their nuclei counterstained with 300nM DAPI (4’,6-Diamidino-2-Phenylindole, Dihydrochloride, Thermo). Coverslips were mounted with Fluoromount G (Southern Biotech). Images were captured using a Zeiss LSM710 confocal microscope, Zeiss Axioscope, or EVOS FL Auto imaging station. Image contrast was applied to the entire image using Photoshop and were annotated with the antibodies used to stain the tissue. Thick tissue sections (30-80μm) were similarly treated, but primary antibody incubation times were extended to 3 days, and rinse times extended to 2-3 hours each.

### Cell Preparation for Flow Cytometry and FACS analysis

To prepare cell suspensions for FACS and flow cytometry analysis, organoids were rinsed twice in PBS and pelleted by centrifuging them at 100g x 2 min. After removing PBS by aspiration, the organoids were suspended in 1 ml of ‘Cell Dissociation Reagent’ (Sigma# C5914; or Thermo# 13150016), incubated for two minutes at room temperature, upon which 3ml trypsin (or TrypLE) was added (0.25%, Thermo 25200072, or TrypLE™, Thermo 12605010; Note that we have found TrypLE better preserves CD49f staining but worsens CD34 resolution). Samples were incubated by hand/body temperature, thus permitting visual inspection and brief (1-3sec) pulse vortexing every 30-60 seconds. After the mixture became cloudy (about 8-10 min), the cells were gently and repeatedly pipetted through a 16-gauge needle until the clumps of cells dissipated. Afterward, we filtered cell suspensions through a 100μm cell strainer and added 3ml 0.1% w/v soybean trypsin inhibitor to stop the digest (Sigma# T9128). The suspensions were then filtered through a 40μm cell strainer, rinsed in 10-20ml PBS, and pelleted by centrifugation at high centrifugal force (400g x 5 min.—or longer, until all cells had pelleted, which we confirmed by microscopic examination of the supernatant). The cells were then rinsed in 10 ml Hanks balanced salt solution/1% BSA(w/v), counted, and again pelleted by centrifugation. After centrifugation, nearly all the Hanks/BSA was aspirated, leaving the cell pellet with roughly 60μl of Hanks/BSA. The pellet was resuspended in this small volume, and we added FACS antibodies to the samples and incubated the samples on ice, covered, for 30 minutes (See Figure 5-Table Supplement 1 for antibodies). Following the incubation, the cells were rinsed in Hanks/1% BSA, centrifuged (400g x 5 min), and resuspended in Hanks/BSA with To-Pro-3 viability marker (diluted 1:4000, Thermo T3605). We did not use DNase enzyme or hypotonic RBC lysis solutions on the cell preparations because we found they were unnecessary, and we wanted to avoid any potential deleterious effect these treatments may have on the cells, such as reducing cell viability or altering gene expression levels. We also conscientiously chilled samples during cell preparation and the sorting procedure for similar concerns.

### Flow Cytometry

The FACS panel and gating strategy outlined herein was developed using a BD FACS Vantage 8-channel cytometer (FACSDIVA software) and has since been adapted and reproduced using a 14-channel SONY SY3200. PMT voltages were optimized on both instruments by analyzing individually stained compensation beads (AbC™ Anti-Mouse Bead Kit, Thermo # A10344) at increasing voltage intervals for each detector. Due to the higher autofluorescence of breast cells (compared to polystyrene beads), we found that defining the compensation spillover matrix was optimally set if individually stained breast cells were used as compensation controls. Breast cells were also used to titrate each antibody to identify the optimal staining dilutions, which are provided in Figure 9 Supplementary Figures 1&2. Because the FACS Vantage has only 8 available channels, it was necessary to use two markers (podoplanin and CD45) in a single channel (A488), which we experimentally validated. To-Pro-3 was used as the viability marker, so the corresponding 405nm channel (455/50) could be used for BV421. This channel was dedicated to CD49f to provide a maximum resolution of cell populations, which is essential to this sorting strategy. The considerable compensation typically required between the heavily overlapping dyes PE/Cy5 and To-Pro-3 was not needed, as the To-Pro-3 negative (viable) cells were used for downstream analyses, allowing us to use the PE/Cy5 channel for Thy1. Negative controls consisted of unlabeled beads and cells incubated with isotype control antibodies conjugated to each fluorophore. The additional channels available on the SONY SY3200 allowed us to move CD45 to an independent channel (PE/Cy5), which was available after we moved Thy1 to the PE-Dazzle-594 channel. This adjustment permitted sorting of the CD45^Pos^ leukocytes earlier in the gating scheme (Figure 9 supplement 2). Sort times typically ranged between 4-6 hours (∼10million cells stained), and 14-18 hours (50-120 million cells stained— needed to yield enough cells for RNA-sequencing). Cells were chilled at the sample intake and collection chambers, and all samples were chilled on ice during the entire experiment. All media and PBS were filtered through a 0.2μm filters to prevent the FACS machine from triggering events from crystals and debris. Staining profiles of cells collected at the beginning and end of the sorts (sometimes separated by as many as 18 hours) were identical and did not show any signal loss or appreciable decrease in viability. To purify cells for cell culture, we found it necessary to triple-sort the cells. Because our machines could sort cells into a maximum of 4 tubes, we naturally needed to sort the cells into four groups, containing 2-4 populations each. These mixtures were: Tube1 (CD34Pos) and contained Pops 7,8,9; Tube 2 (Thy1^Pos^) contained Pops 3,6; Tube 3 (Luminal) contained Pops 1&2; and Tube 4 (Null gate) contained Pops 4,5,10,11. During the first round of sorting, cells were typically sorted at approximately 8,000 cell/sec. We followed this initial enrichment with purity sorts for each cell type, using stringent sort masks, sorting each population into individual tubes at a rate of approximately 2,000 cells/sec. For cells earmarked for cell cultures, the cells were sorted a third time to ensure purity of the isolated fractions and were sorted directly into 96-well dishes containing the appropriate medium (at 1-1500 cells/well). In some experiments, irradiated fibroblast feeder layers (1500 cells/well, 30Gy X-ray) were used to support growth, and were applied into every other column of wells. FACS data were analyzed using Flowjo software (version v7.6.3 – v10.8), Tree Star Inc./ Becton, Dickinson & Company).

### Cell Culture

We generated primary cultures by seeding each cell type onto collagen I (PureCol, Advanced Biomatrix) coated 96-well cell culture dishes (Falcon 353075) containing M87 (M87+CT+X^50^) for Pops 1,2,3,4; M87 supplemented with an additional 10% fetal bovine serum (FBS) for Pops 5,6,8; and EGM2 (Lonza CC-3162) supplemented with an additional 10% FBS for Pops 7&9. We supplemented media with 1x Penicillin (100U/ml), 1x/Streptomycin (100μg/ml), and 0.5x Normocin (50μg/ml); EGM2 contains gentamicin and amphotericin and was therefore only supplemented with 0.5x normocin for the immediate culture post-FACS. A range of seeding densities were used in experiments, but 1500 cells per (96) well (∼4700 cells/cm^2^) was generally sufficient to produce colonies in most wells for each cell type. When the cells were at low cell densities in these initial cultures, the media was initially replenished after seven days post-seeding, then replenished every 2-3 days (Monday/Wednesday/Friday). We did not allow the epithelial cells (Pops1-4) to become confluent, but other cell types (Pops 6,7,8,9, & 12) all fared better when maintained at confluent densities and continued to divide in these conditions. Special attention was given when subculturing the cells, with the goal to minimize time exposed to trypsin. To subculture, we thoroughly rinsed each well 1x PBS, then applied Sigma Cell Dissociation Reagent (Sigma C5914 or Thermo 13150016—enough to cover the cells). Once the cells displayed early signs of detachment from the plate, which we monitored by phase microscopy, we added several drops (∼100ul) of 0.25% trypsin (or TrypLE). Typically, within 15-30 seconds, most cell types detach, at which point we gently and repeatedly pipetted the cells to detach them from the surface of the culture dish. We then inactivated the trypsin by adding an equal volume of 0.1% w/v soybean trypsin inhibitor. The cells were subsequently rinsed in excess PBS (10-12 ml), centrifuged, and either archived in cryopreservative medium (75% base medium, 15% FBS, 15% v/v DMSO) or re-seeded in their respective media onto PureCol coated dishes. All cell types were cultured under normoxic conditions at 37°C with 5%CO_2_, in a humidified cell culture incubator (Thermo/Forma).

Adipocyte cultures were created using a modified ceiling culture method^27^. Briefly, T-25 flasks were coated with bovine collagen I (PureCol®), turned upright, and filled with DMEM/F12 supplemented with 15% FBS and 1x Penicillin (100U/ml), 1x/Streptomycin (100μg/ml), and 0.5x Normocin (50μg/ml). 100-300μl media was removed and replaced with an equal volume of the lipid layer from the tissue/collagenase digests that had been prefiltered through a 40μm cell strainer. The flask was capped, inverted, inserted into a secondary container, and placed into a humidified incubator. Having the flask upside-down permits adipocytes to float to the surface and attach to the collagen-coated surface. After one week in culture, the medium is removed, the flask is returned to an upright position and replenished with 3ml fresh medium. The adipocyte cultures are subsequently maintained as a typical adherent culture—as described above for the other cell types. At first passage, the cells are harvested, stained, and FACs sorted for CD36 to remove potentially contaminating fibroblasts. Lipids were stained using Oil Red O (Sigma O0625) or BODIPY (4,4-difluoro-1,3,5,7,8-pentamethyl-4-bora-3a,4a-diaza-s-indacene (BODIPY® 493/503, Thermo D-3922).

### Adipogenesis/Osteogenesis lineage differentiation of Pop8 ADMSC/Fibroblasts

To test the lineage differentiation capabilities of Pop8 fibroblasts, we placed early passage Pop8 fibroblasts into modified adipogenic and osteogenic conditions^36–38, 51, 52^. Briefly, early passage (3-5p) Pop8 fibroblasts were seeded at a density of ∼53,000 cells/cm^2^ in PureCol coated 4-well tissue culture dishes (100,000 cells per well) in M87/10%FBS+P/S+N (pen-strep/normocin-described above). After cells were attached to the dish, the medium was replaced with adipogenic or osteogenic differentiation medium, which was made fresh weekly and replenished bi-weekly (every 3-4 days). Adipogenic Differentiation Medium consisted of DMEM/F12, 10%FBS, 1x Penicillin (100U/ml), 1x/Streptomycin (100μg/ml), and 0.5x Normocin (50μg/ml); Supplemented with: Dexamethasone (1μM final, Sigma D4902), IBMX (0.5mM final, Sigma I5879), Insulin (10μg/ml final, Sigma I6634), Indomethacin (0.2mM final, Sigma I7378), and Rosiglitizone (1μM, Cayman 71740) – added fresh. Adipocyte induction medium (AIM) is reportedly stable for three weeks without rosiglitazone and 1-2 days with rosiglitazone.

Differentiation was monitored by microscopy, and cells were maintained until they formed large lipid droplets, which usually took 10-21 days. Cells were stained by supplementing the complete growth medium with 2μg/ml Hoechst 33342 and 10μg/ml BODIPY, then incubated at 37°C for 1 hour. After the incubation, the cells were rinsed thrice with 1X PBS, and observed by fluorescent microscopy. Osteogenic Differentiation Medium consisted of Alpha-MEM supplemented with 10% FBS, 1x Penicillin (100U/ml), 1x/Streptomycin (100μg/ml), 100nM dexamethasone, 50μM L-Ascorbic acid 2-phosphate sesquimagnesium salt hydrate (Sigma A8960), 4mM beta-glycerophosphate (10mM), 0.5μM 1α,25-Dihydroxyvitamin D3 (Sigma D1530). After two weeks, 20μM Xylenol Orange (Fluka/Fisher 33825) was added to the medium and incubated for an additional 24 hours. The cells were then incubated an additional hour with Hoechst 33342, rinsed with PBS, and imaged by fluorescent microscopy.

### RNA Isolation and Sequencing

Total RNA was isolated from FACS-sorted cells using Qiagen RNA Mini (>100,000 cells, Qiagen 74104), RNA Micro (<100,000 cells, Qiagen 74004); and from the collagenase lipid layer using Qiagen RNA Lipid Tissue Mini Kit (Qiagen 74804) according to the to manufacturer’s instructions. Total RNA was treated with DNAse I for 30 minutes at 37°C (Ambion DNA-*free* DNA Removal kit, Thermo AM1906), then treated with DNAse Inactivation beads. RNA was then concentrated with RNA Clean & Concentrator™-5 columns (Zymo R1015), quantified by Nanodrop and Agilent Bioanalyzer assays, and stored at −80°C. Complete details of RNA sequencing and links to the dataset are provided in the adjacent article (Del Toro, et.al., in preparation). Briefly, mRNA was captured using poly-T coated dishes (Clontech) and SMARTer technology was used for library construction (SMARTer Stranded RNA-seq Kit).

Sequencing was performed on Illumina Trueseq v.2 technology, targeting approximately 30million paired-end reads (on average) for each cell population. We processed raw count data using R programming language (RStudio) and associated BioConductor packages. The count matrix was transformed for analysis, using two methods used in the DESeq2 package; i.e., (i) regularized log transformation (rlog), and (ii) variance-stabilizing transformation (vst)^53^. Quality control assessment of expressed transcripts across sorted cell samples was conducted and one sample was removed from statistical analyses, as it was an extreme outlier (Epithelial Population 4 Sample N239). This was determined by: (i) distance matrix, (ii) Principal Component Analysis, and (iii) Kolmogorov-Smirnov test that compares each sample to the reference probability distribution of the pooled data. This sample is however included in downstream visualizations, for example, the bar and donut graphs used throughout the manuscript, which display rlog transformed values on both log2 (bar graphs) and linear (donut graph) scales.

### Quantitative RT-PCR Array

A custom qRT-PCR array was generated by designing and validating PCR reactions for 148 genes related to breast cancer, cell adhesion and signaling, ECM and ligand pathways, and stem and developmentally related pathways, such as wnt/beta catenin, notch, TGFbeta, and hedgehog. Primer sets are provided in Figure 8—Table Supplement 1 and contain the official gene symbols, pathway/category information, ENSEMBL accession IDs, number of exons per gene, intron spanning information, and sequences for FOR and REV primers. Although not used, we have included also sequences of designed qRT-PCR Taqman probes. Total RNA was isolated from FACS-sorted cells (Qiagen RNA Mini Kit), were treated with with DNAse I (Ambion DNA-*free)*, and reverse transcribed using the SuperScript® VILO™ cDNA Synthesis Kit (Thermo 11754-050) in 20 ul reactions according to the manufacturer’s protocol. After cDNA synthesis, the cDNAs were diluted 8x, by adding 140 ul water to the 20 ul reactions. Transcripts were amplified, by qPCR, using the LightCycler® 480 SYBR Green I Master Mix in 10μl reactions, using 1μl diluted cDNA in each reaction. Primers, supplied as 300nM final concentration in the reactions were designed to be 80-200bp in length and overlapped exons, if possible. All PCR primers are provided in Figure 8—Table Supplement 1. PCR reaction parameters: 5min activation at 95°C for 5 min; 40 cycles each of melting at 95°C for 10sec., one-step annealing and extension at 60°C for 60seconds; and was followed by melting curve analysis. Relative levels of transcripts were calculated using the delta Ct method and normalized to those of the TBP reference transcript using the formula: %*TBP* = 2^−(*ct_Gene_*−*ct_TBP_*)^.

## Author Contributions

Conceptualization, WCH; Methodology, WCH, KT; Bioinformatics, KDT, RWS and WCH; FACS, WCH; Primary cell models and immunostaining, WCH, KT, KDT, YLM; Writing – Original draft, WCH; Writing – Review & Editing, WCH, KT, KDT, RWS, MB; Funding, RWS, MB and WCH.

## Acknowledgments

We thank Mina Bissell, Irene Kuhn, Alex Bazarov, Ritu Mukhopadhyay (Lawrence Berkeley Laboratory), Sandy Borowsky (U.C. Davis), Kornelia Polyak (Dana Farber Cancer Institute), Eric Prossnitz (University of New Mexico Health Sciences Center), Curtis Hines (Sandia National Laboratory, retired) for thoughtful scientific discussions and/or for critical review of the manuscript. We thank Stefan Klimaj, Selina Montoya, Roman Azharelleu, Kremena Karagyozova, Alyssa Cozzo, Gaelen Stanford-Moore, Maria Rojec, Anya Afasisheva, and Sun-Young Lee for their technical assistance. We thank and are very appreciative for the help of Ambrose Carr, Zhenmao Wan, Dana Pe’er (Columbia University) and the Columbia U. Genomic Sequencing Core for creating the mRNA sequencing libraries, Illumina RNA-sequencing, paired-end read alignment, data normalization, and count table generation used in the adjoining article and figures presented herein. We also thank Kerry Wiles and Erik Brooks of the Cooperative Human Tissue Network, and the anonymous women who graciously donated their remnant surgical tissue to science. We depended on the flow cytometry instruments provided at the University of New Mexico Flow Cytometry and High Throughput Screening Resource and thank Cecelia Jeanette Rietz and Wade Johnson for their assistance. A very special thank you to Michelle Scott, Lawrence Berkeley National Laboratory Advanced Microscopy and Flow Cytometry facility, for her expert technical guidance and support while developing the FACS panel and gating strategy.

---

**Figure 5—Table Supplement 1.**
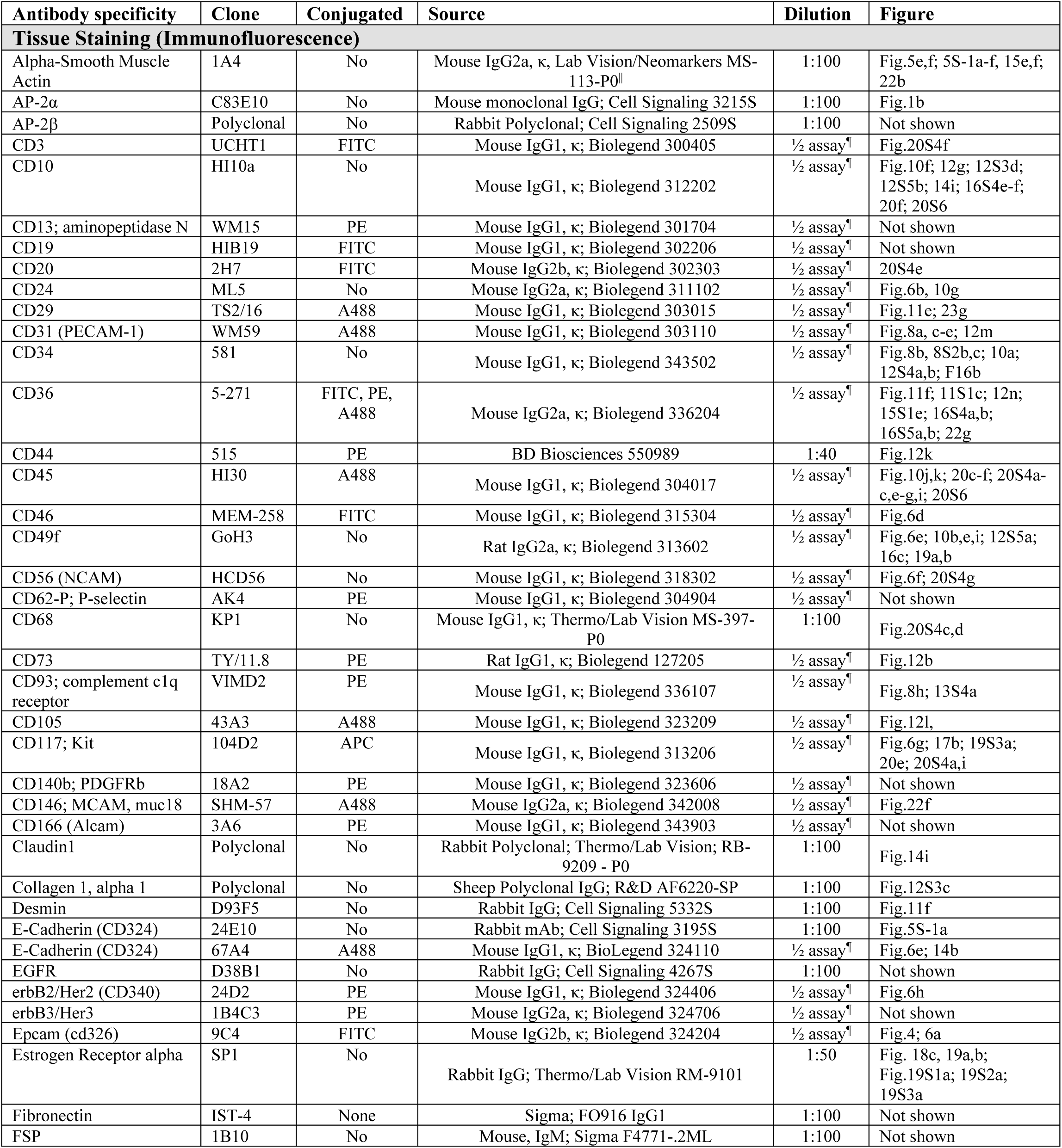
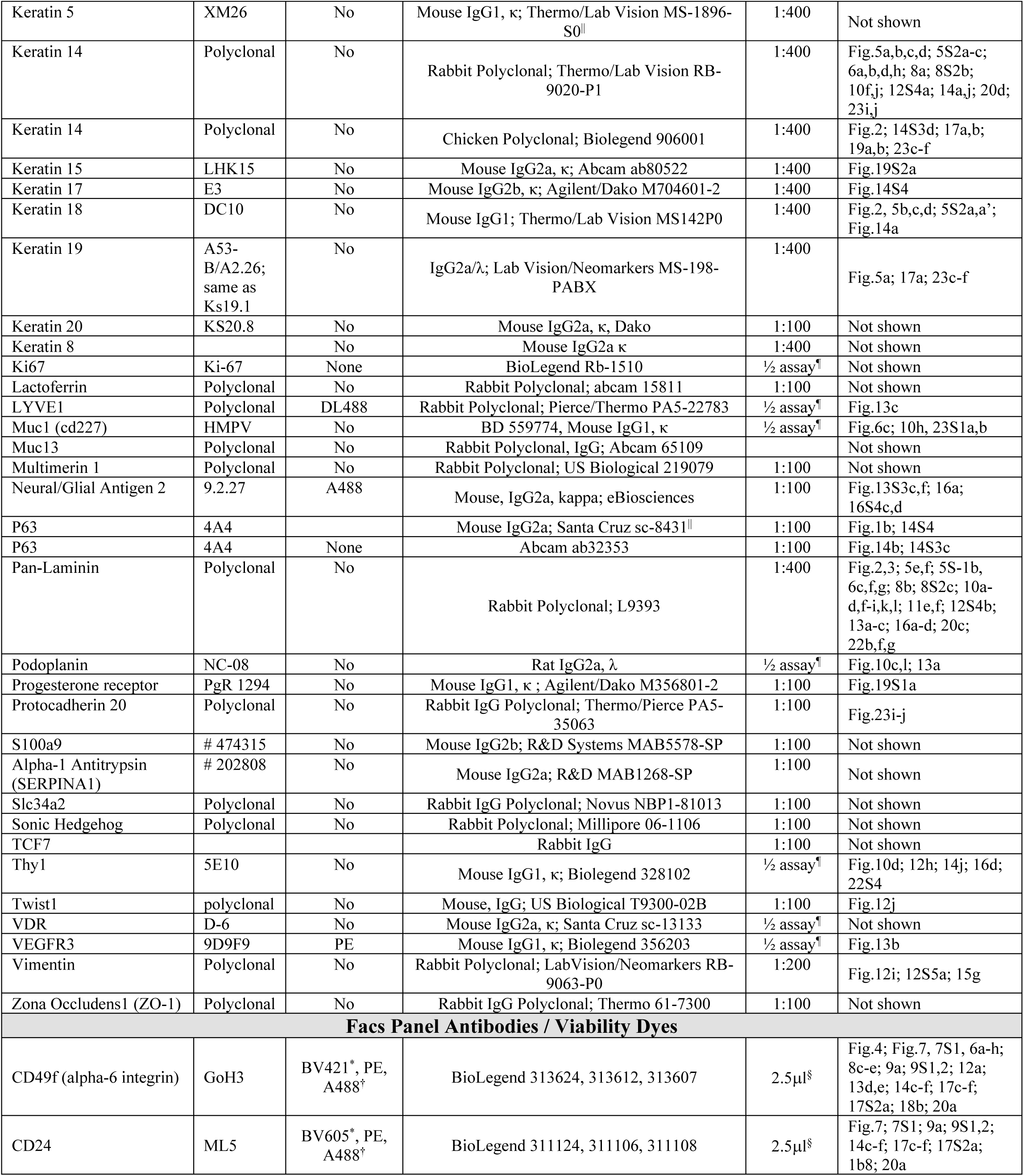
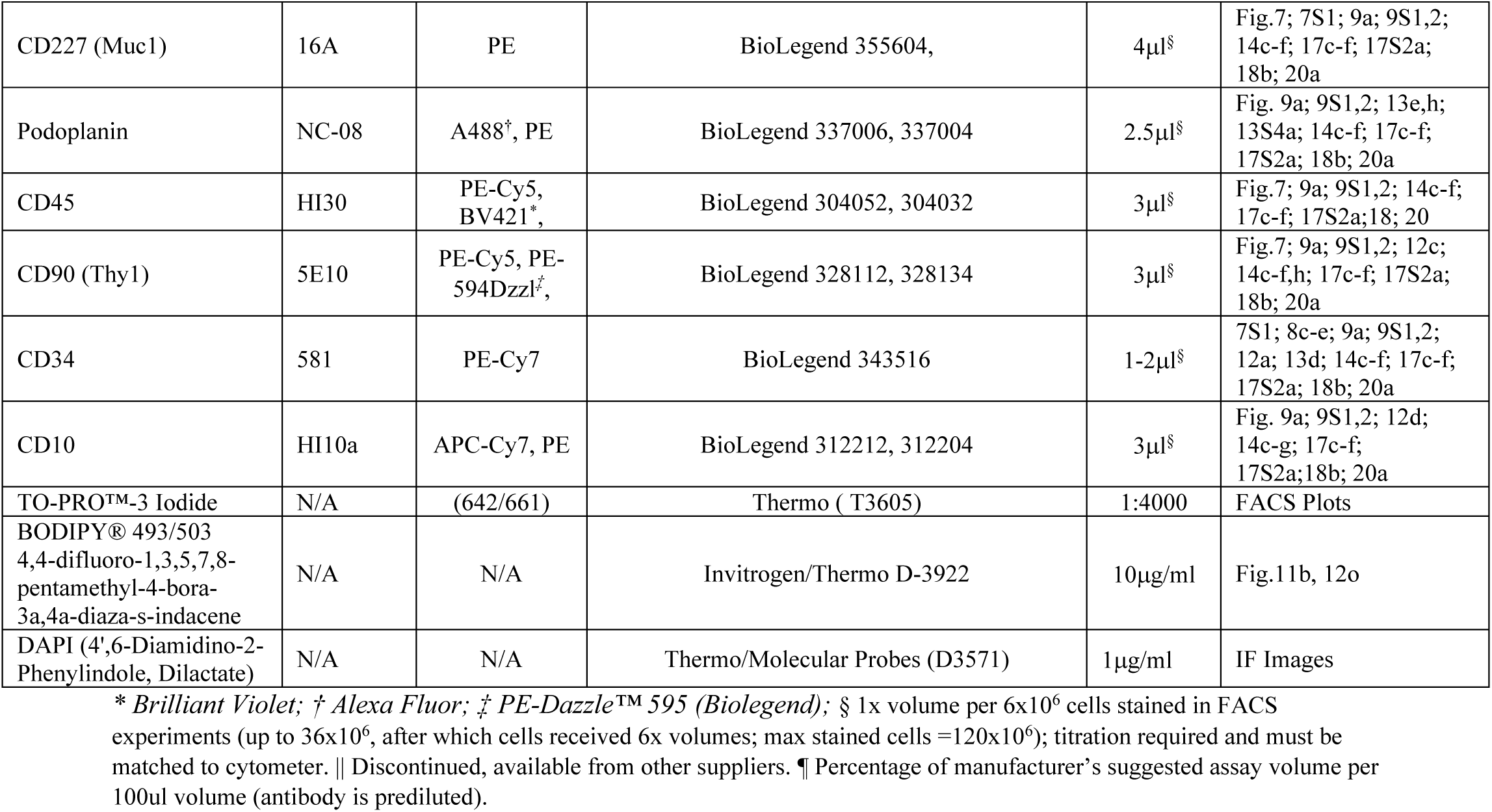
Antibodies and Reagents.

**Figure 8 Table Supplement 1.**
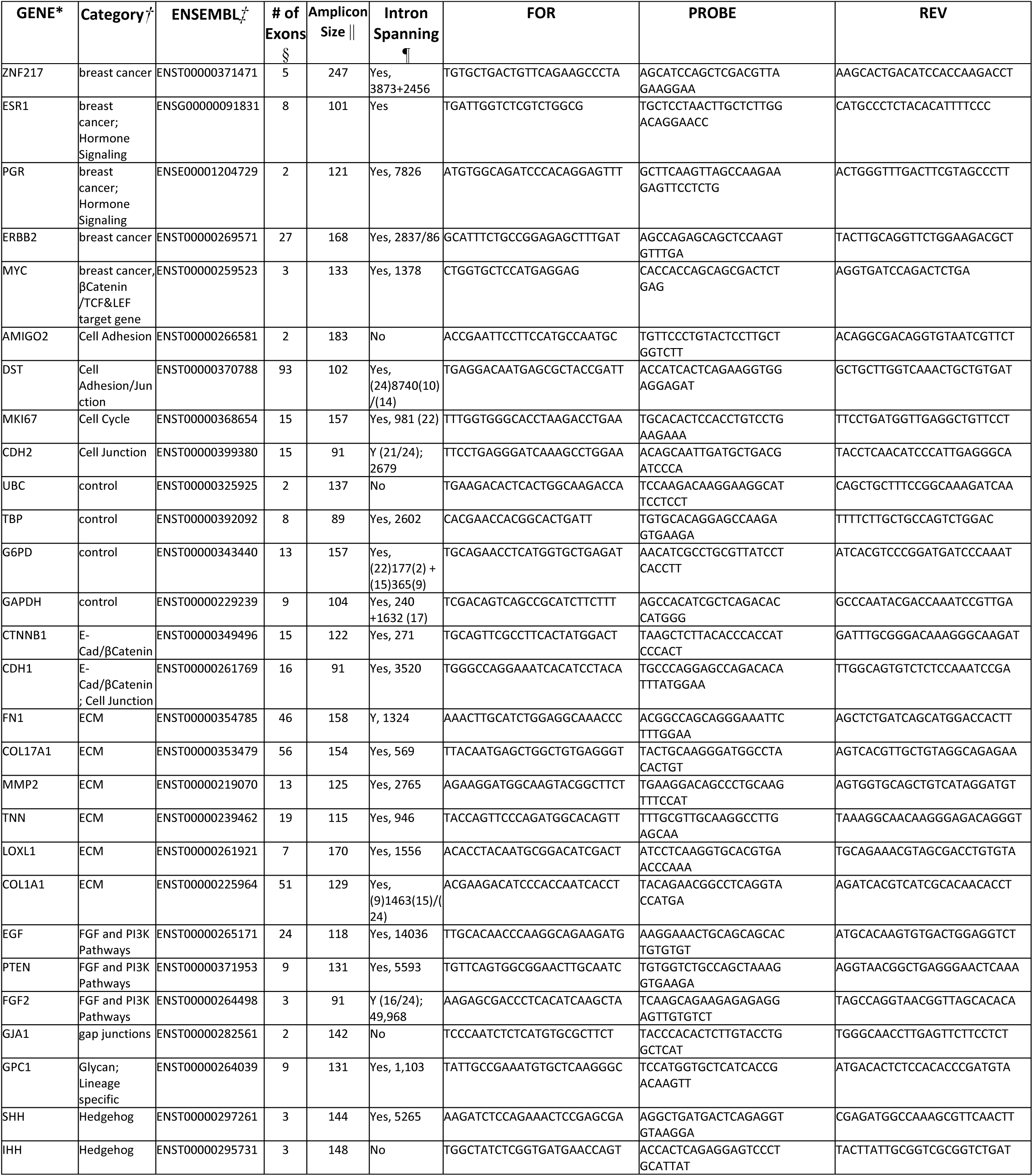
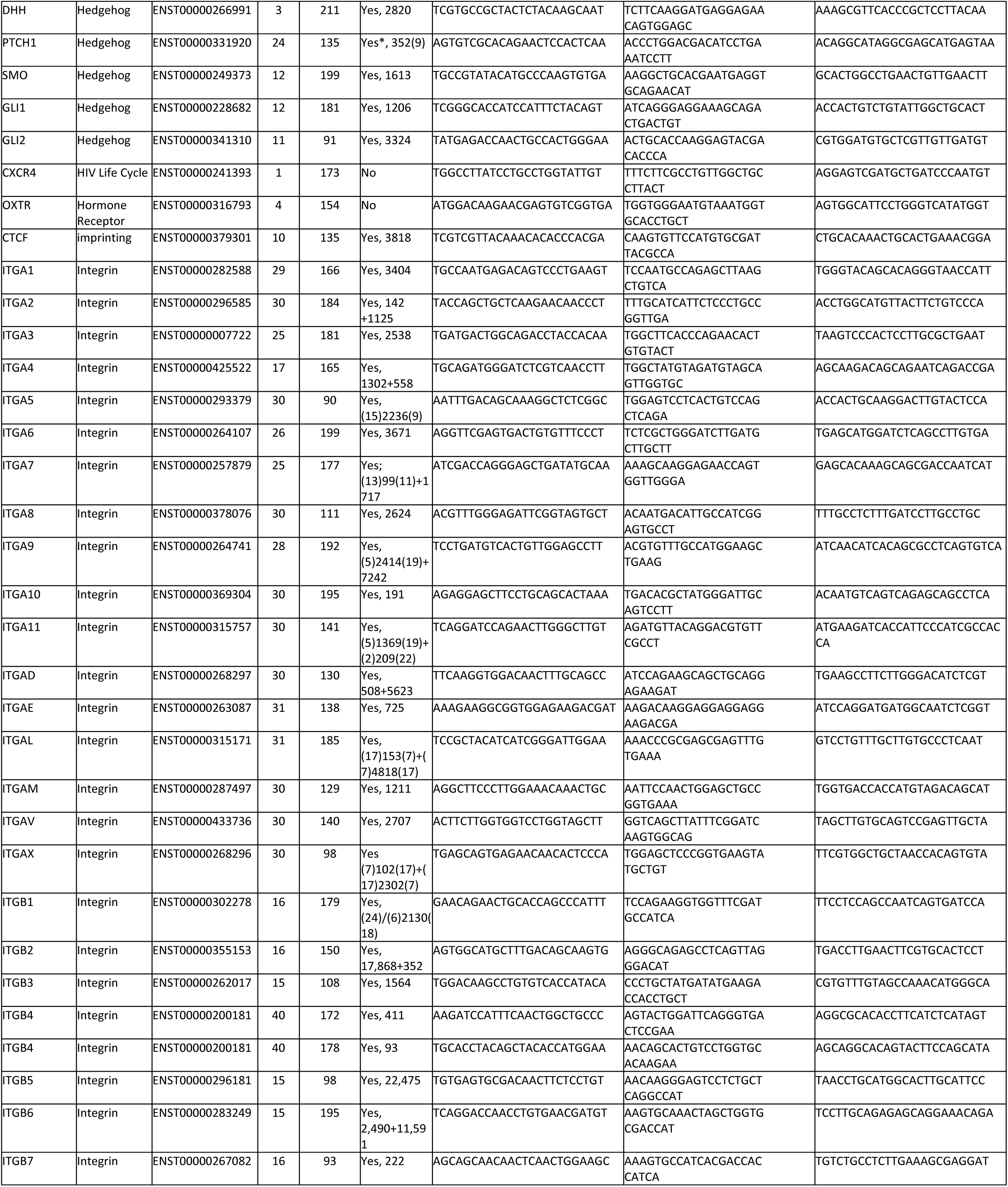
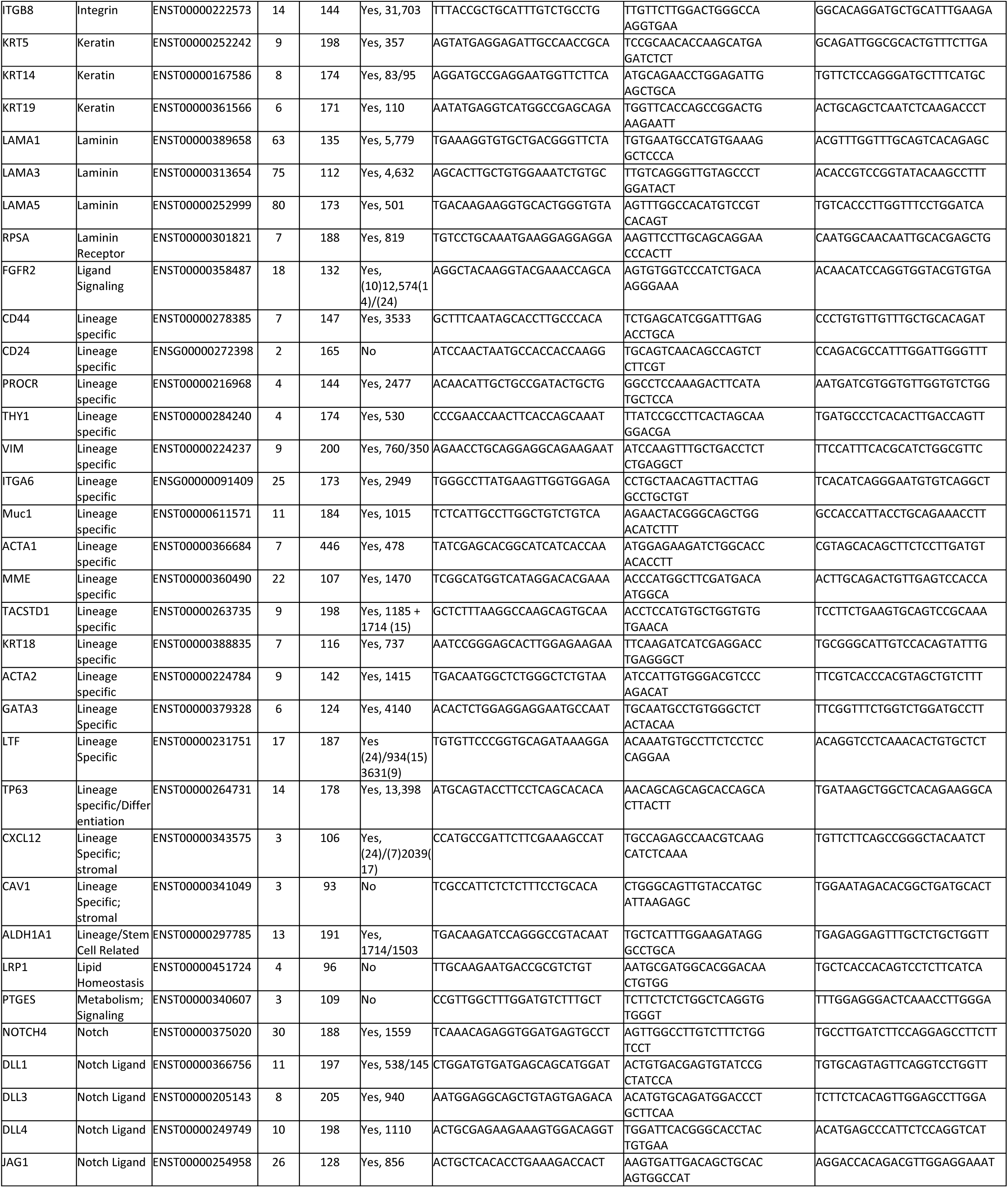
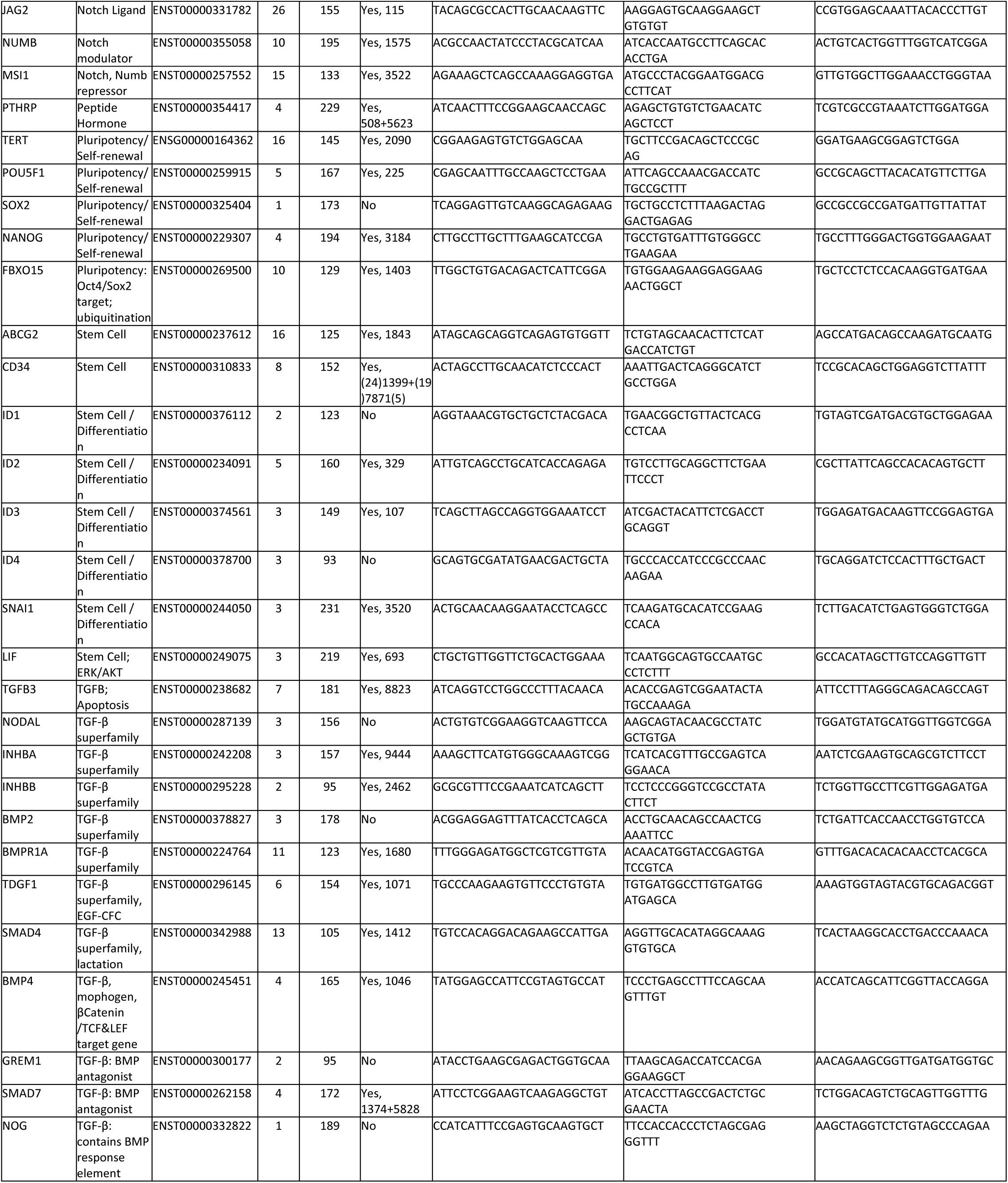
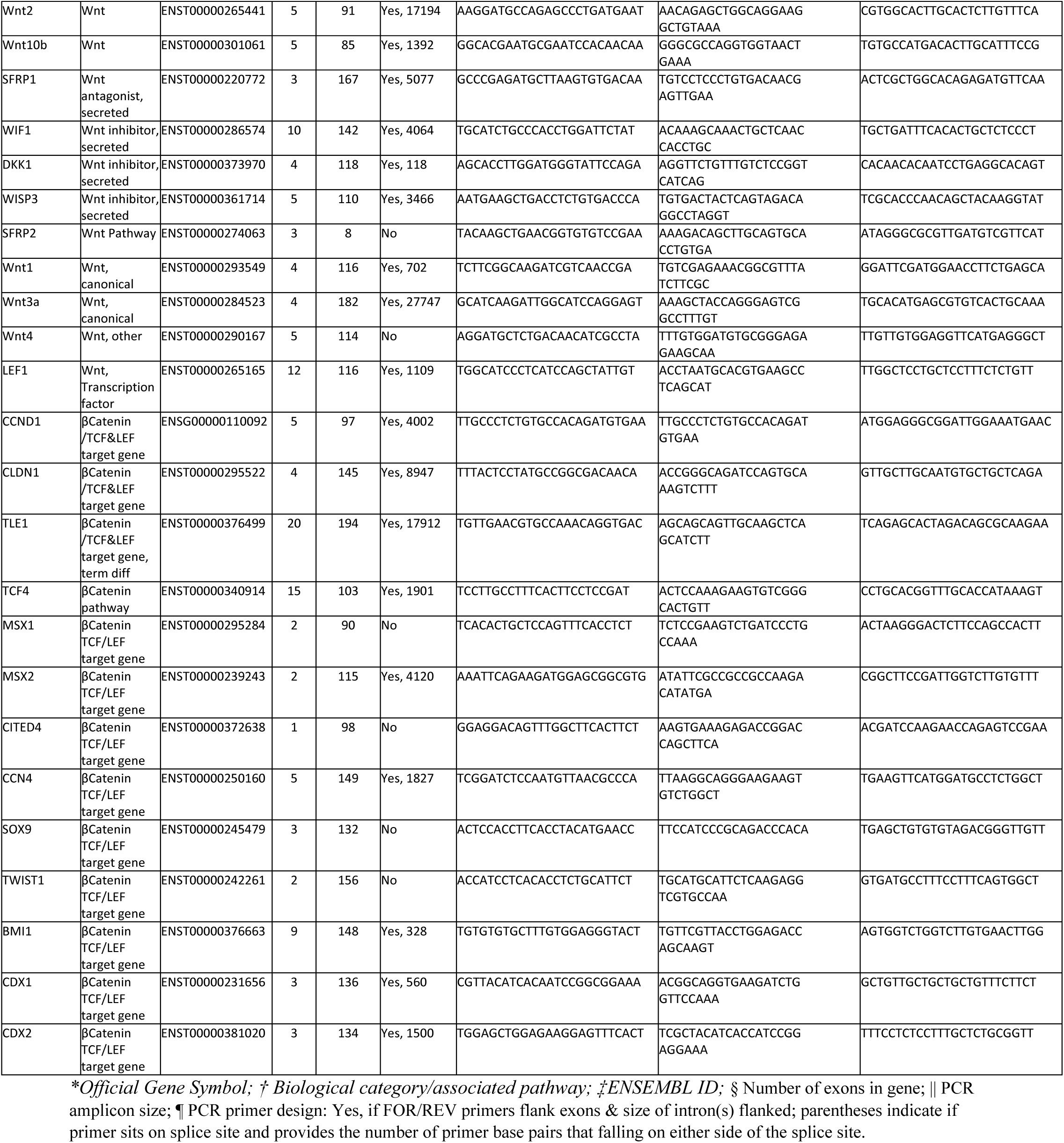
PCR Primers.

**Figure 9 Table Supplement 1.** Proportion of cell types in breast; percentages from cytometry gates. Excel File: Fig.9 Table 1_FACS_Percent_Pops.xlxs

**Figure 1 –figure supplement 1.**
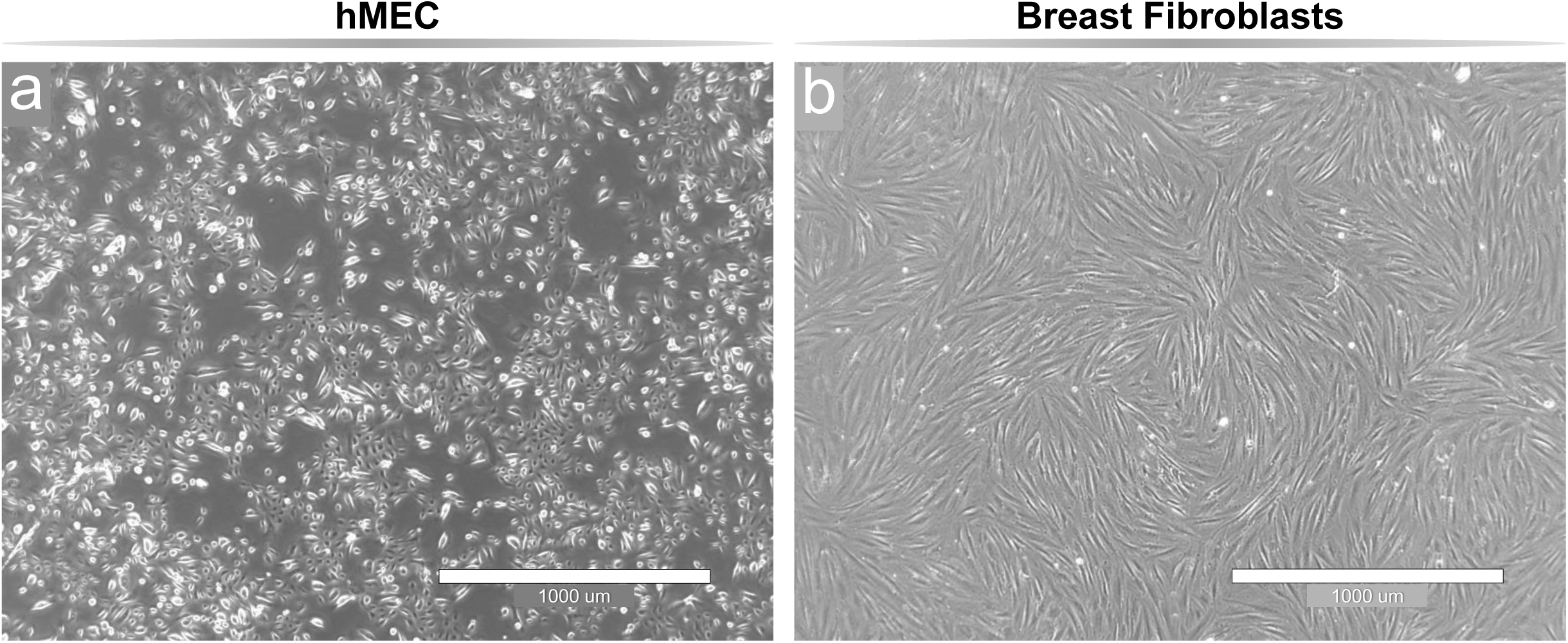
Morphology of common cell models. **(a)** Human mammary epithelial cell (HMEC) and **(b)** fibroblast cell lines were generated from primary cultures seeded with normal breast tissue organoids (collagenase-digested reduction-mammoplasty tissues). The cultures respectively exhibit the characteristic ‘cobblestone’ and ‘spindle-like’ morphologies commonly attributed to HMECs and fibroblasts. Imaged on an EVOS imaging station; 4x Ph Plan 0.13 NA objective. 1000μm scale

**Figure 5—figure supplement 1.**
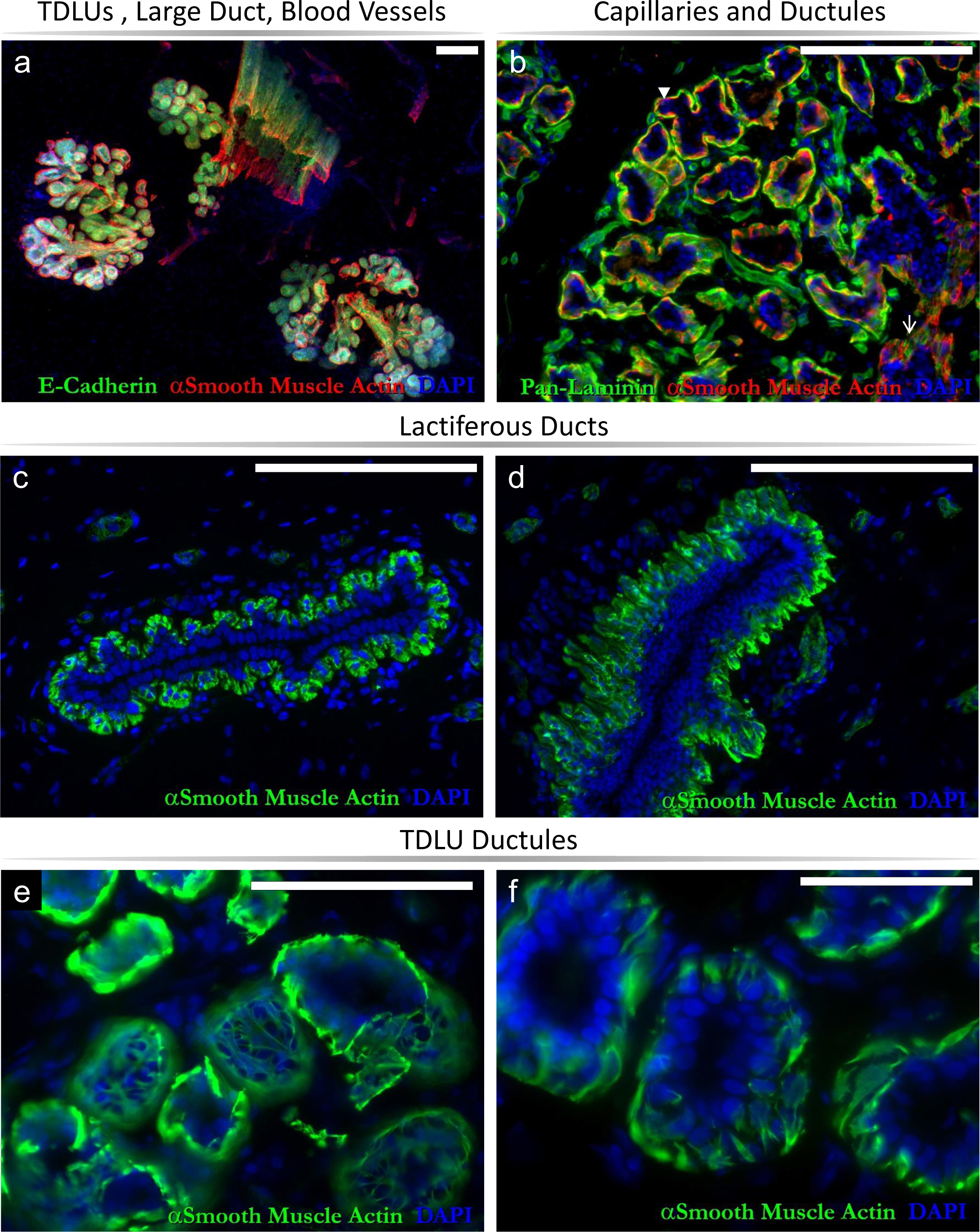
Main architectural elements in normal breast tissue revealed by immunostaining. Breast tissue staining exposes TDLUs, a large lactiferous duct, and capillaries in tissue sections stained for α-smooth muscle actin (ASMA) and **(a)** E-cadherin or **(b)** pan-laminin. ASMA is expressed in myoepithelial cells surrounding ductules (**b,** ▾), large ducts **(c,d)** and lobular tubules **(e,f)**. Scale =200μm (a-d); 100μm (e); 50μm (f).

**Figure 5—figure supplement 2.**
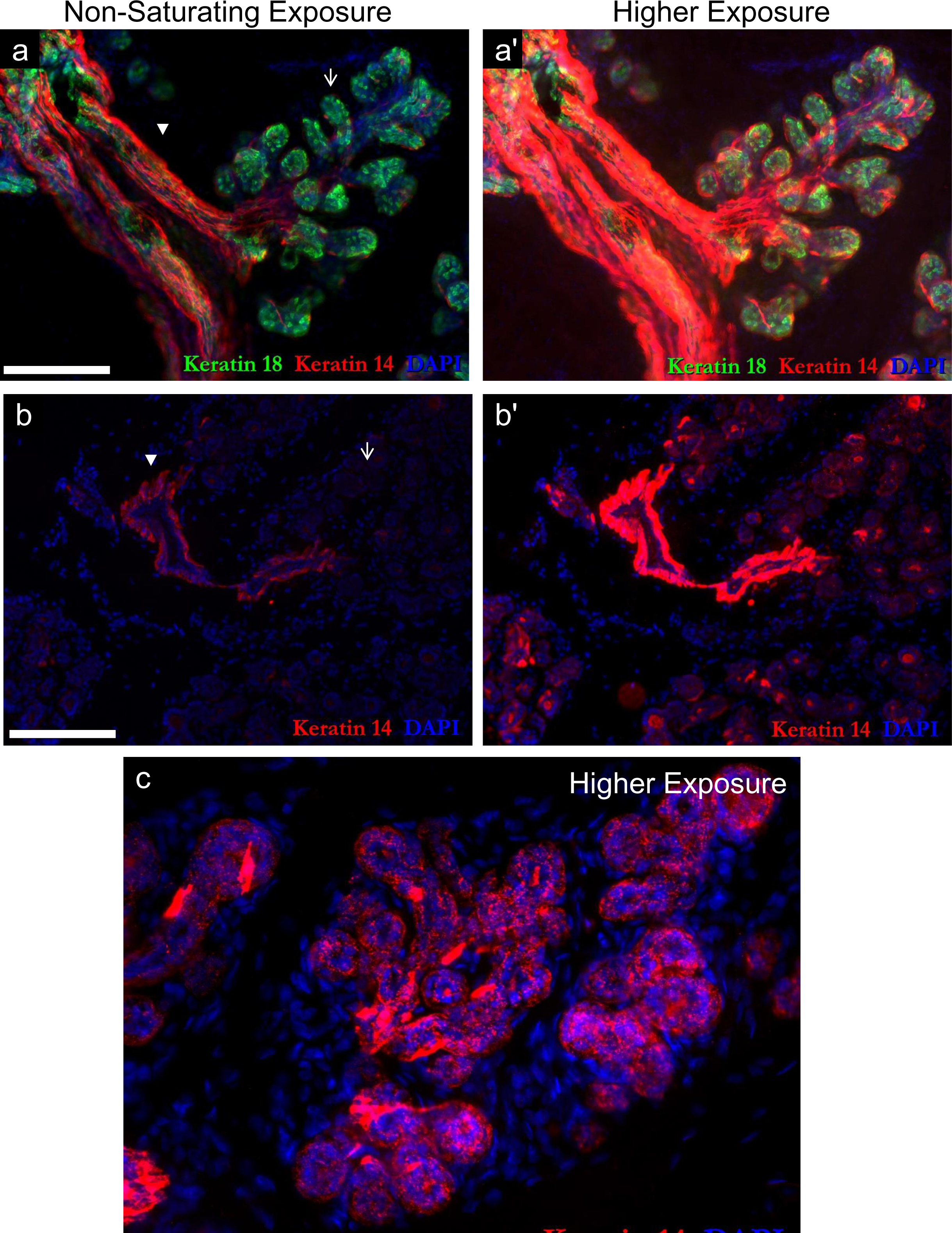
keratin 14 expression obscured by exposure settings. **(a,b)** Breast tissue immunostained with (a) keratin 18 and keratin 14 antibodies, imaged at a non-saturating exposure (as presented in Fig.5b). **(b)** A separate breast section, stained with K14 and also imaged at a non-saturating exposure. In both a&b, the ductal myoepithelial cells (▾) stain strongly for K14, whereas the staining of the ductule myoepithelial cells (↑) is indiscernible. **(a’ and b’)** Same fields as in a & b, imaged with a longer exposure time. At these exposures, k14 expression within ductules becomes apparent, visible as punctate staining within the myoepithelial cells **(c)**. Image of TDLU, stained with K14. The broad range of keratin expression in these different histologic areas (and technicalities associated with photographing it), likely contribute to the disparate claims of keratin 14 expression within ductal and lobular myoepithelial cells. Scale = 0.2mm.

**Figure 6—figure supplements 1-8.**
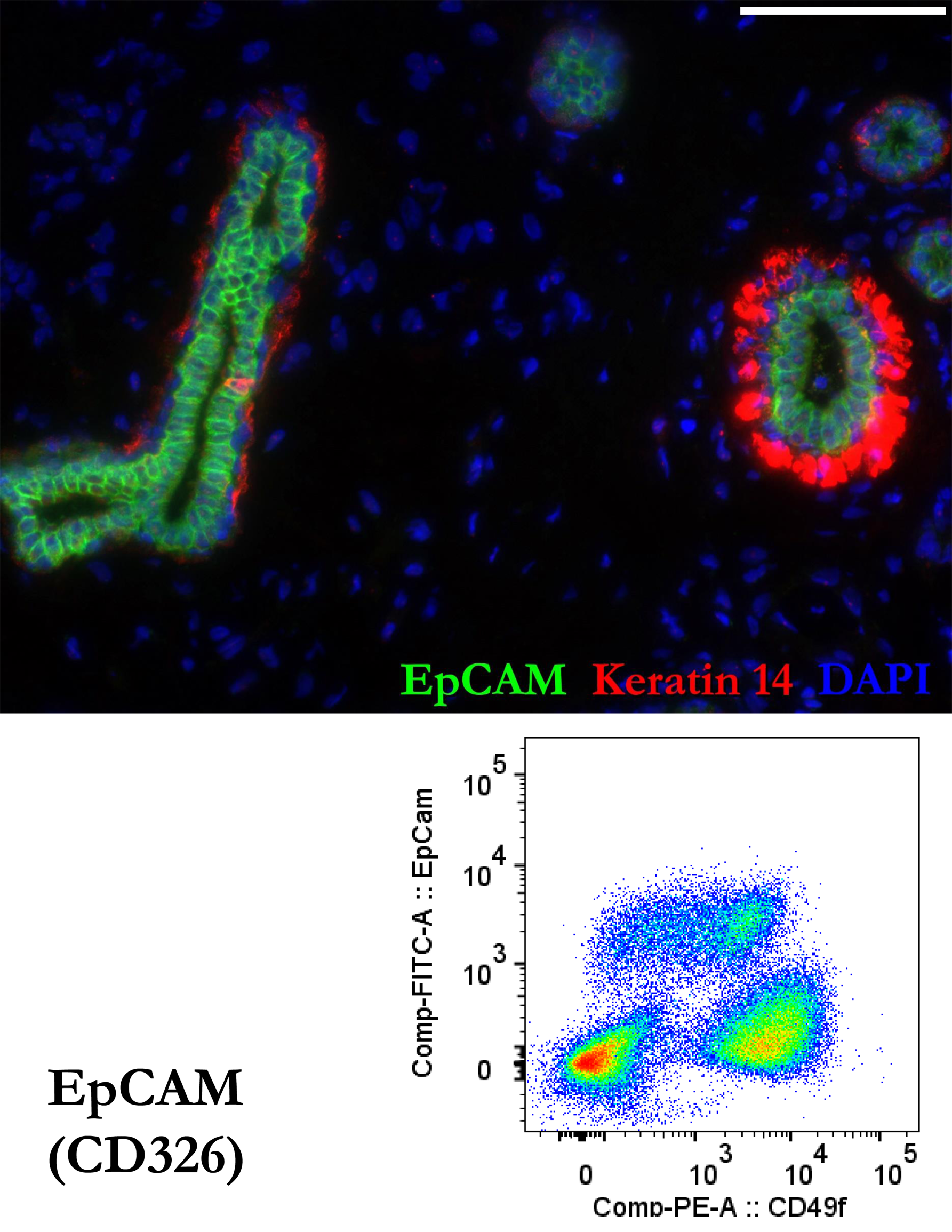
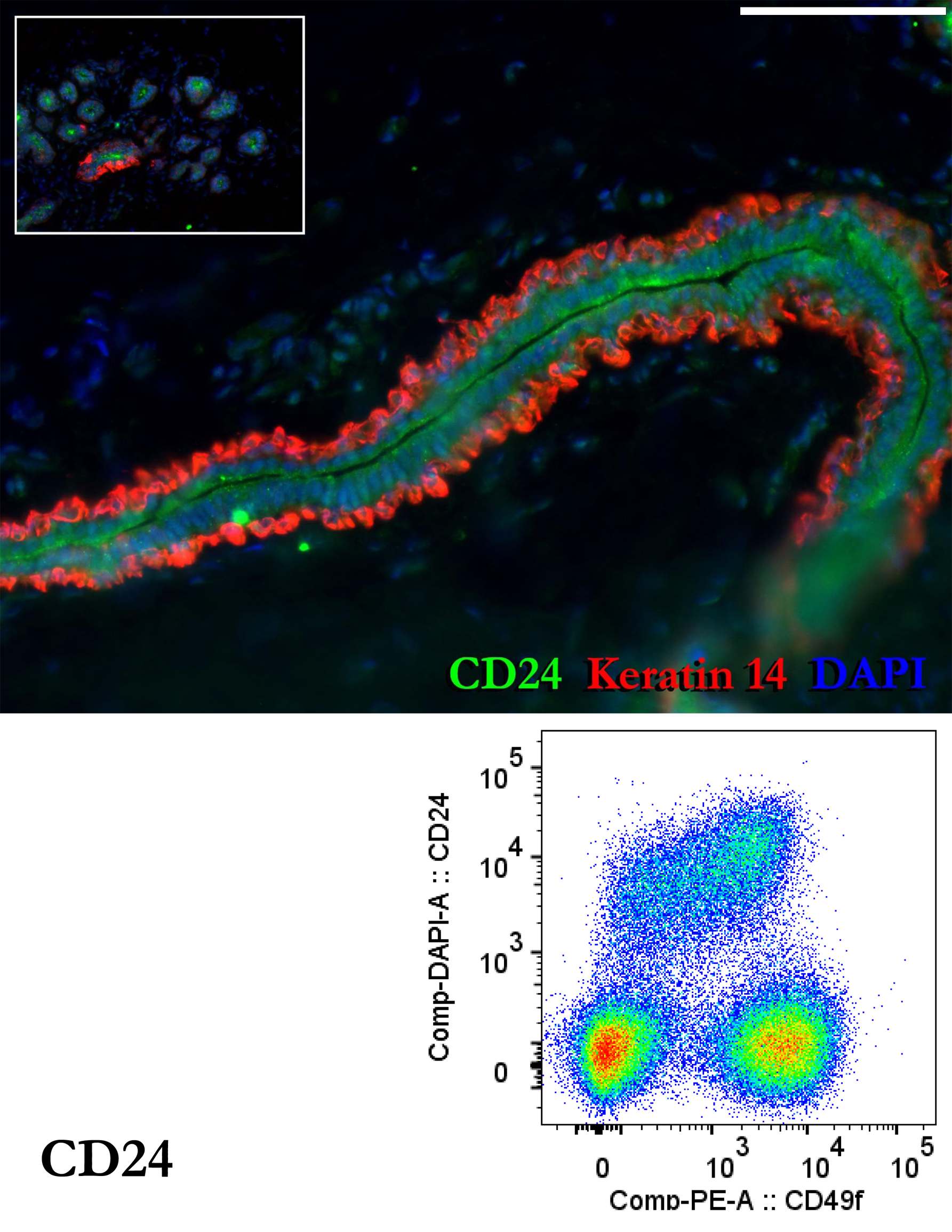
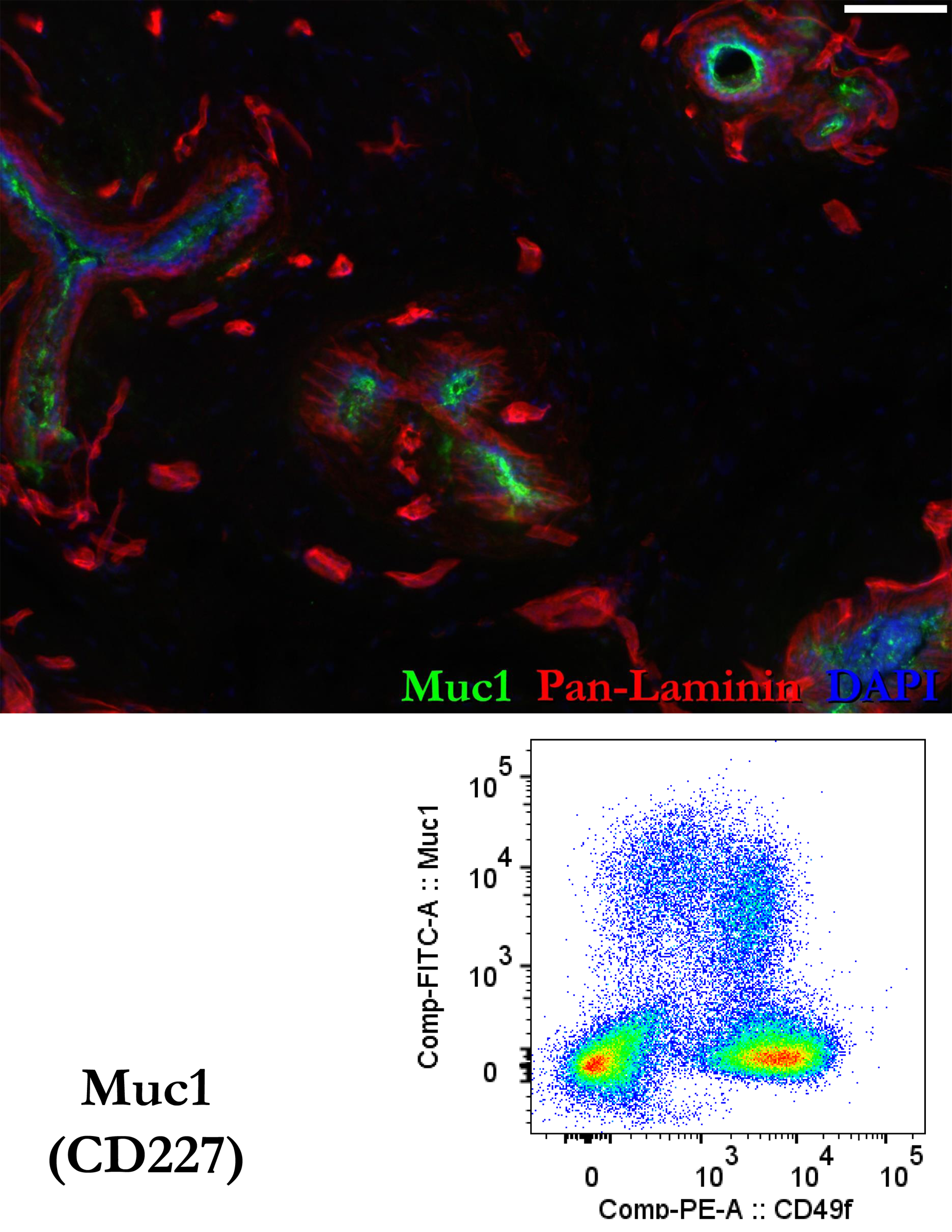
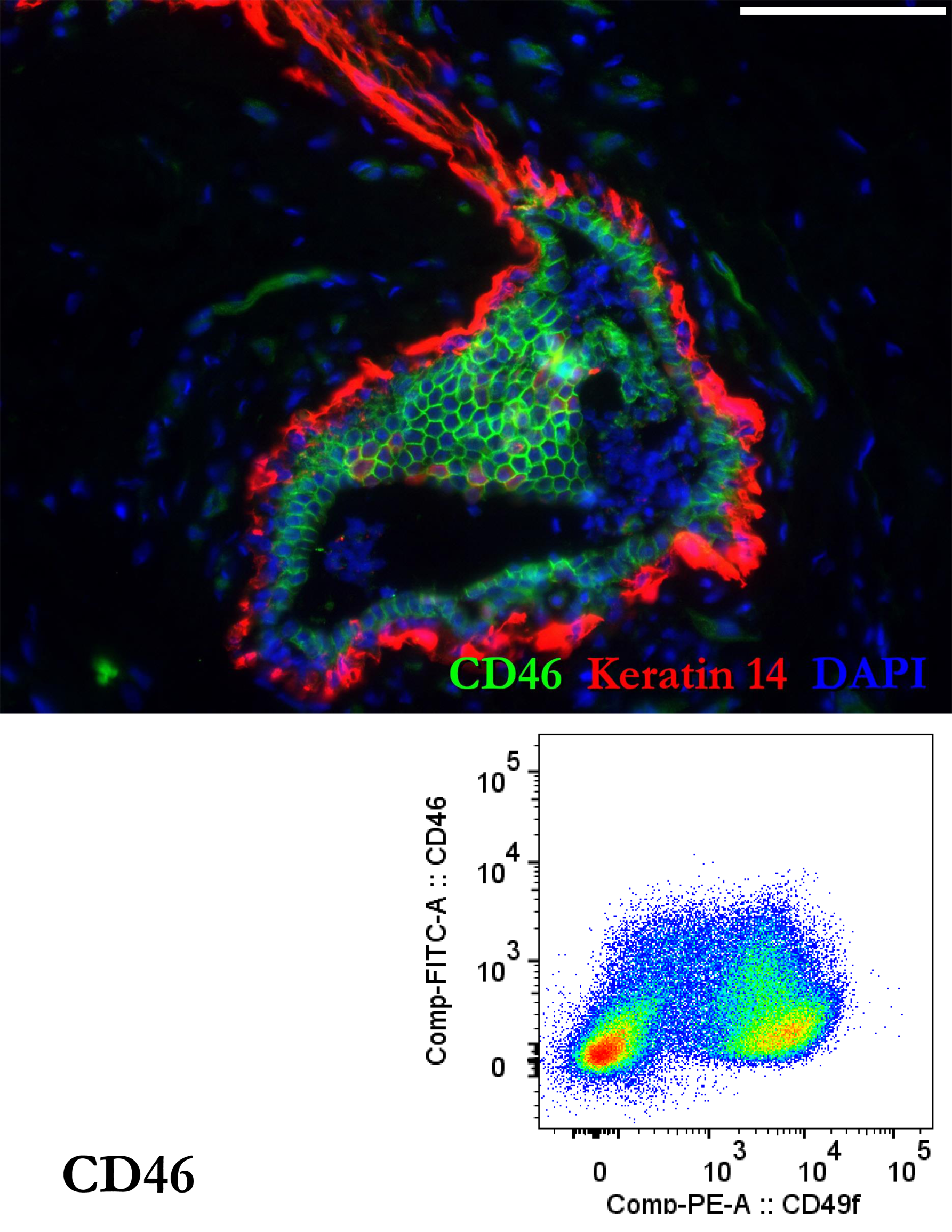
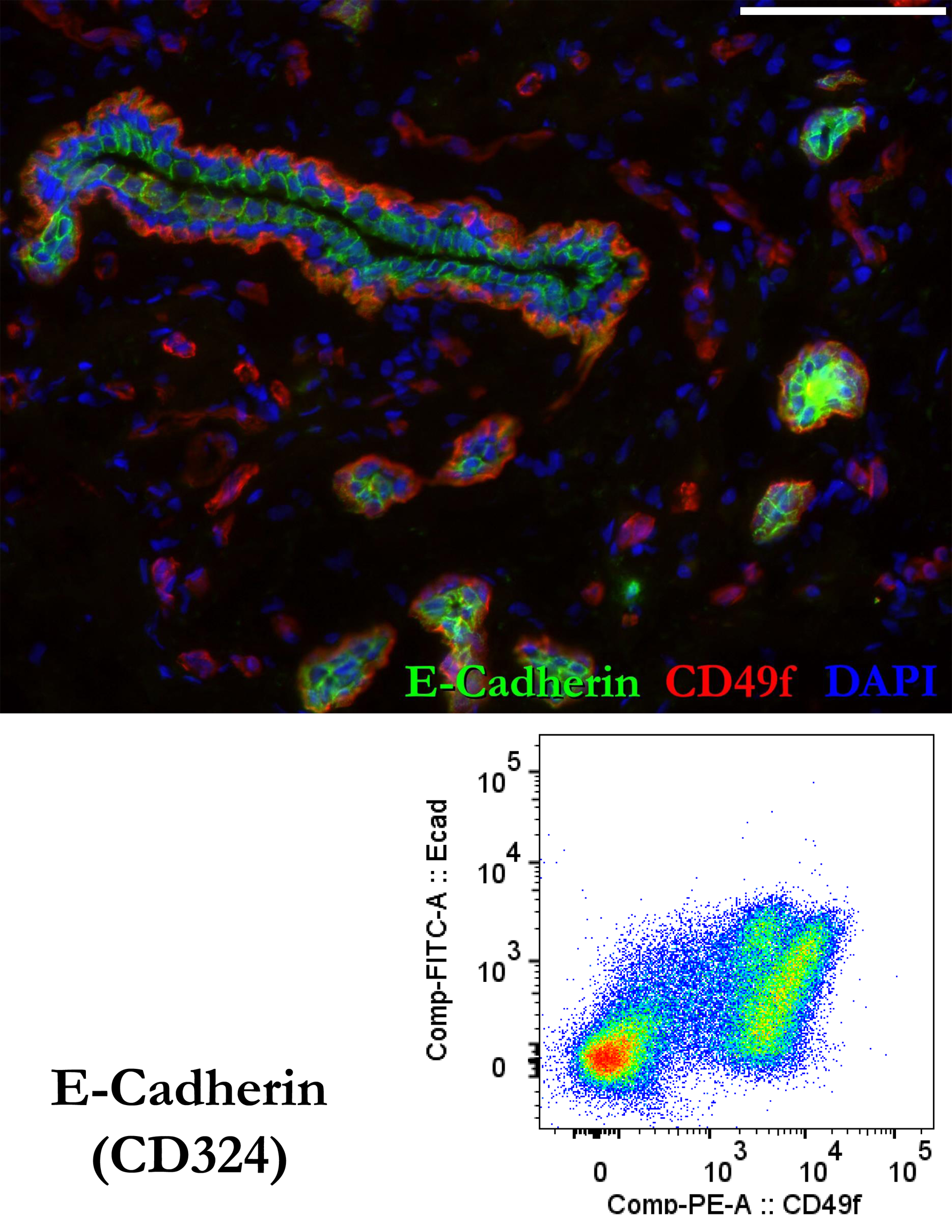
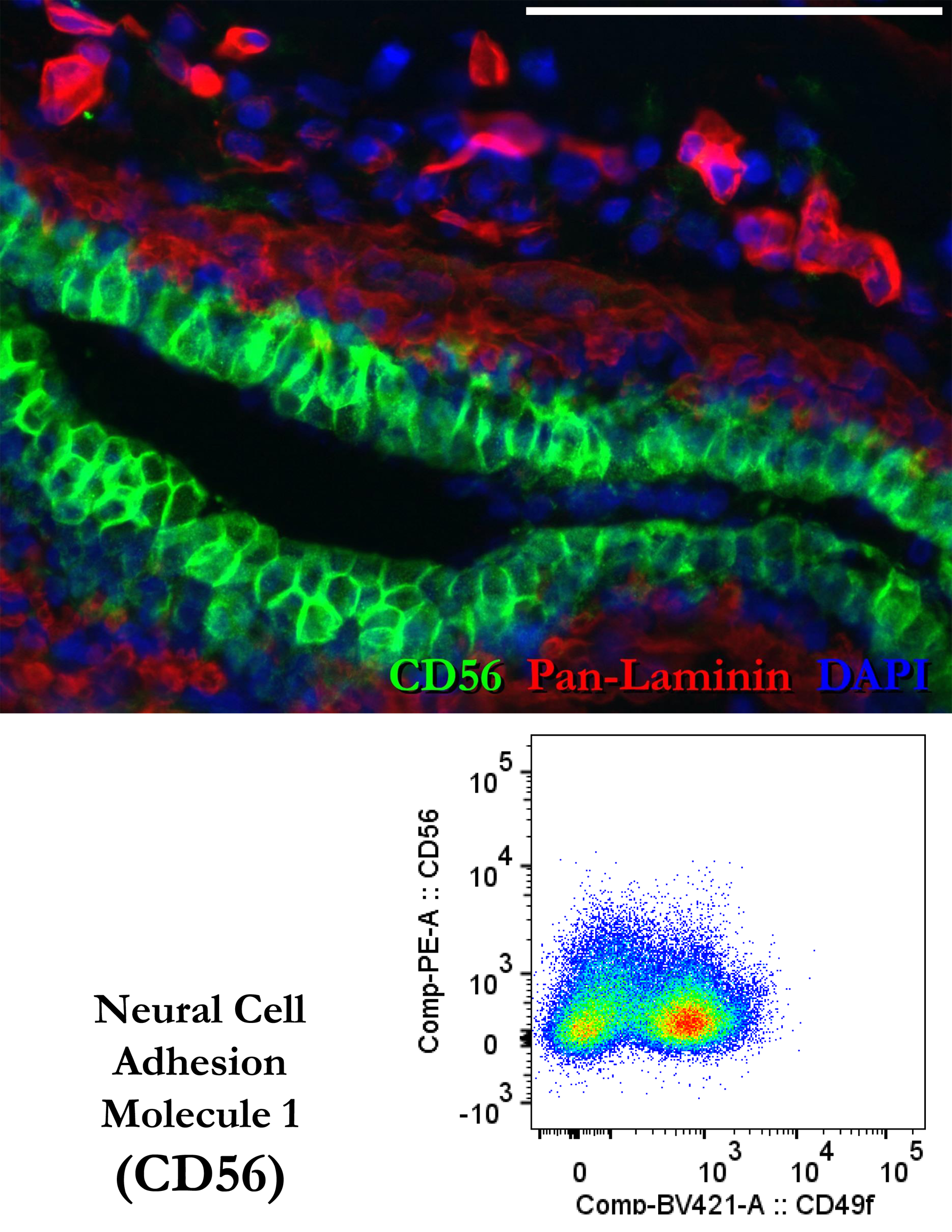
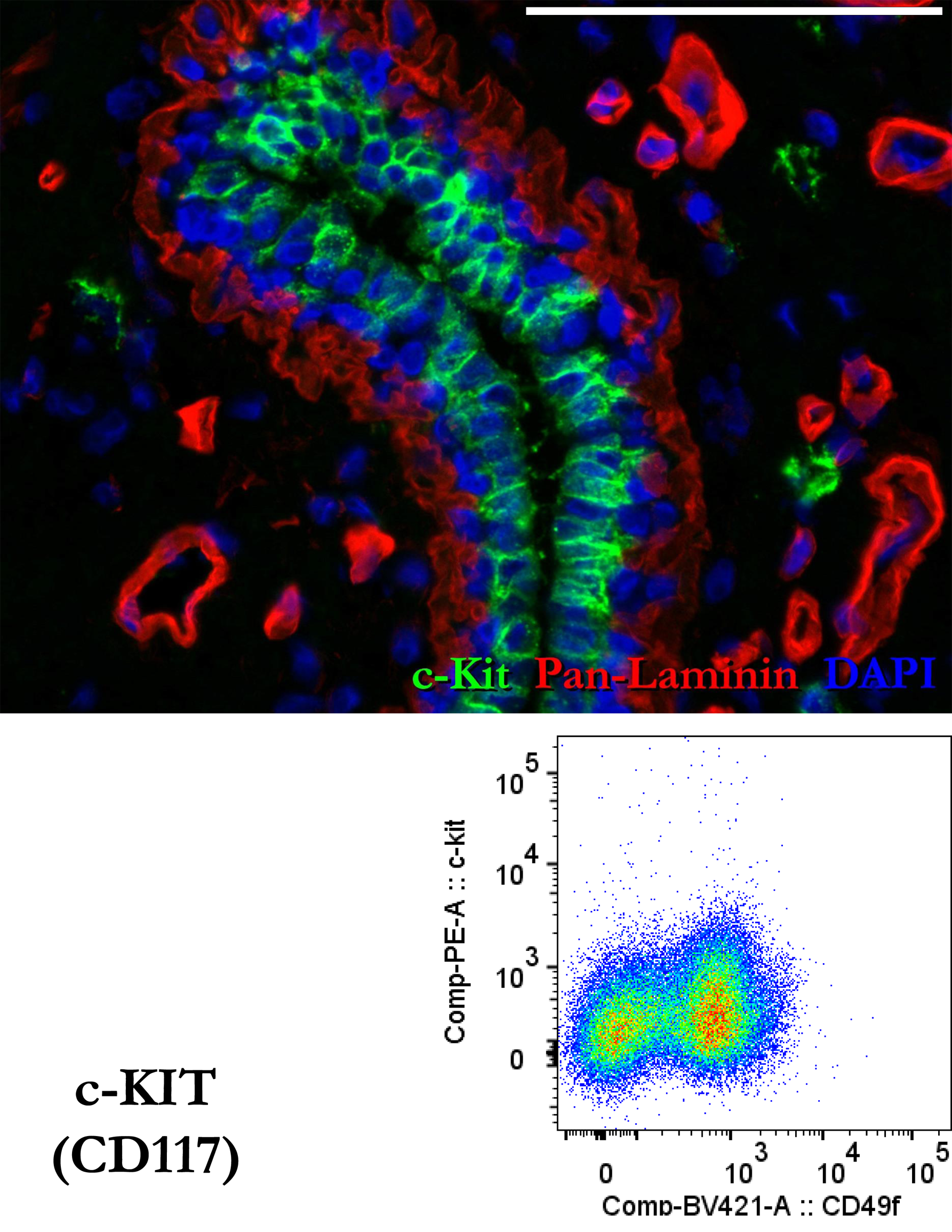
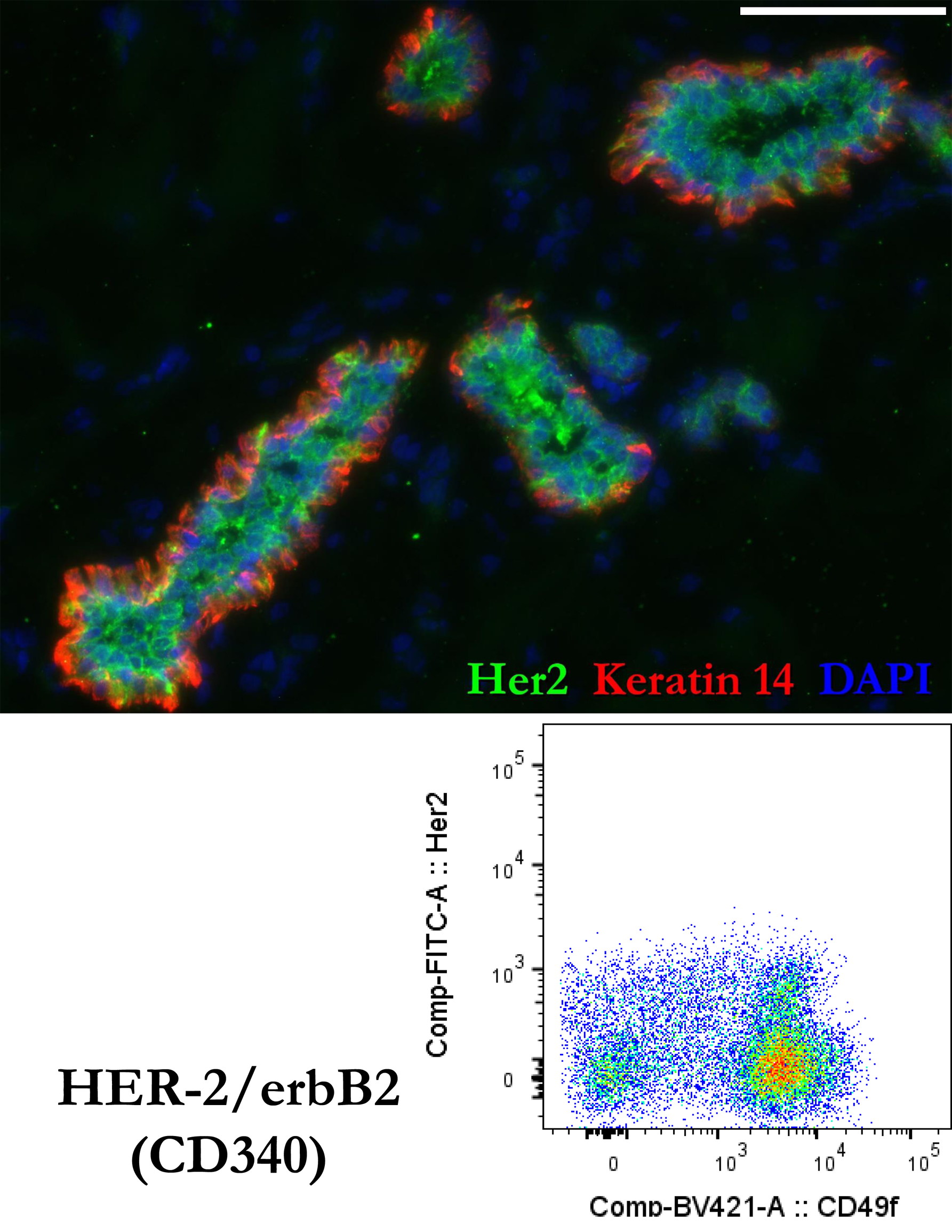
Full resolution images of Figs.6 a-h. scale = 0.1mm

**Figure 7 – figure supplement 1.**
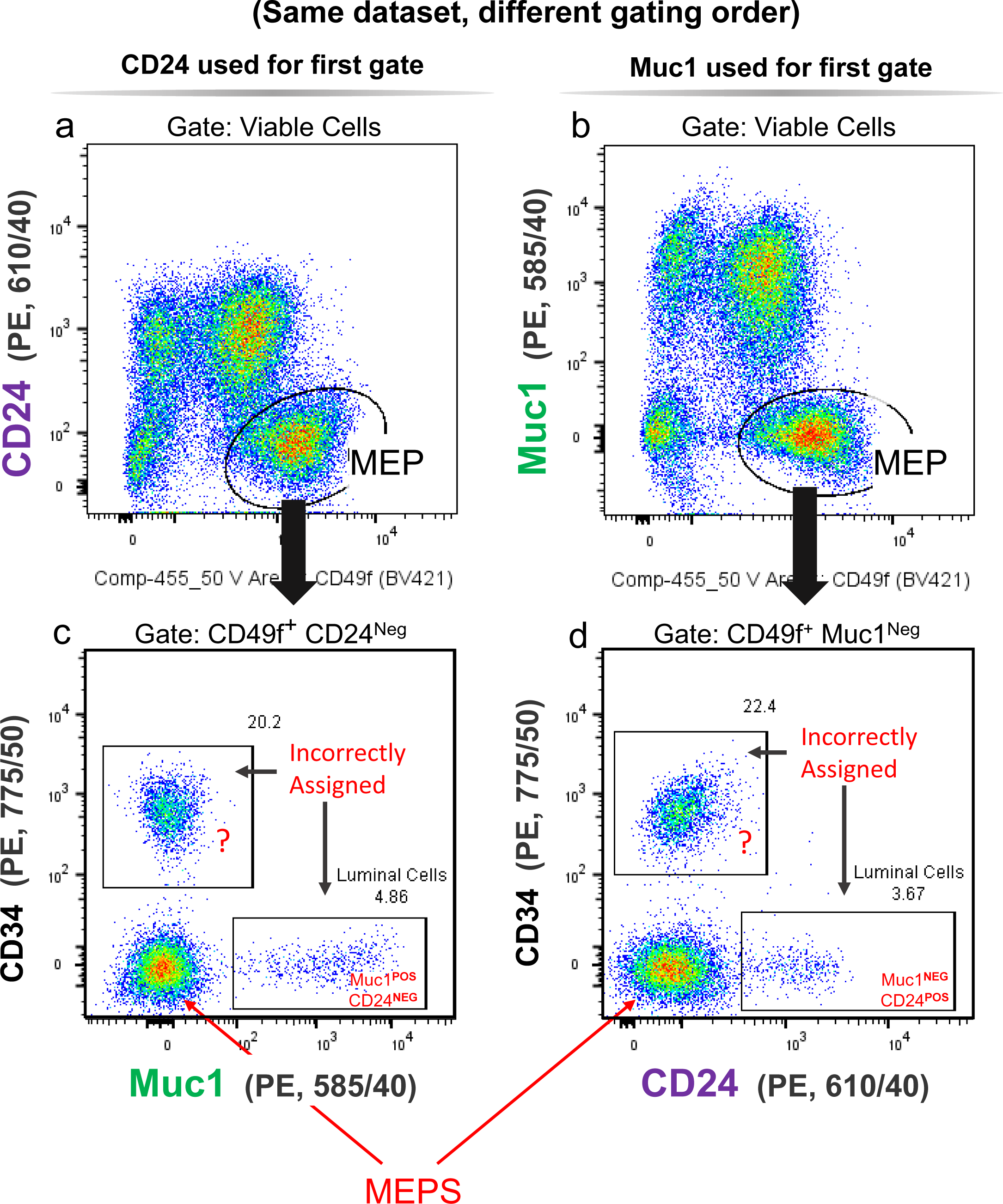
A common FACS strategy leads to incorrect cell assignments. **(a,b)** Representative FACS analysis of normal breast cells co-stained with CD49f, CD34, and both CD24 and Muc1 luminal markers. Myoepithelial cells (MEPs) can be identified by their high CD49f level and either their low (a) CD24 or (b) Muc1 levels. **(c,d)** FACS scatter plots of the gated ‘MEP’ fraction, analyzing CD34 and either (c) Muc1 or (d) CD24 (i.e., the luminal marker not used to define the MEPs in each above plot). This analysis reveals at least two cell fractions are incorrectly assigned as ‘MEPs.’ Roughly 3.5-5% were luminal epithelial cells, whereas 20-22% were an unidentified CD34^Pos^ cell fraction, marked with a ‘?’, which we later identified as endothelial cells.

**Figure 8—figure supplement 1.**
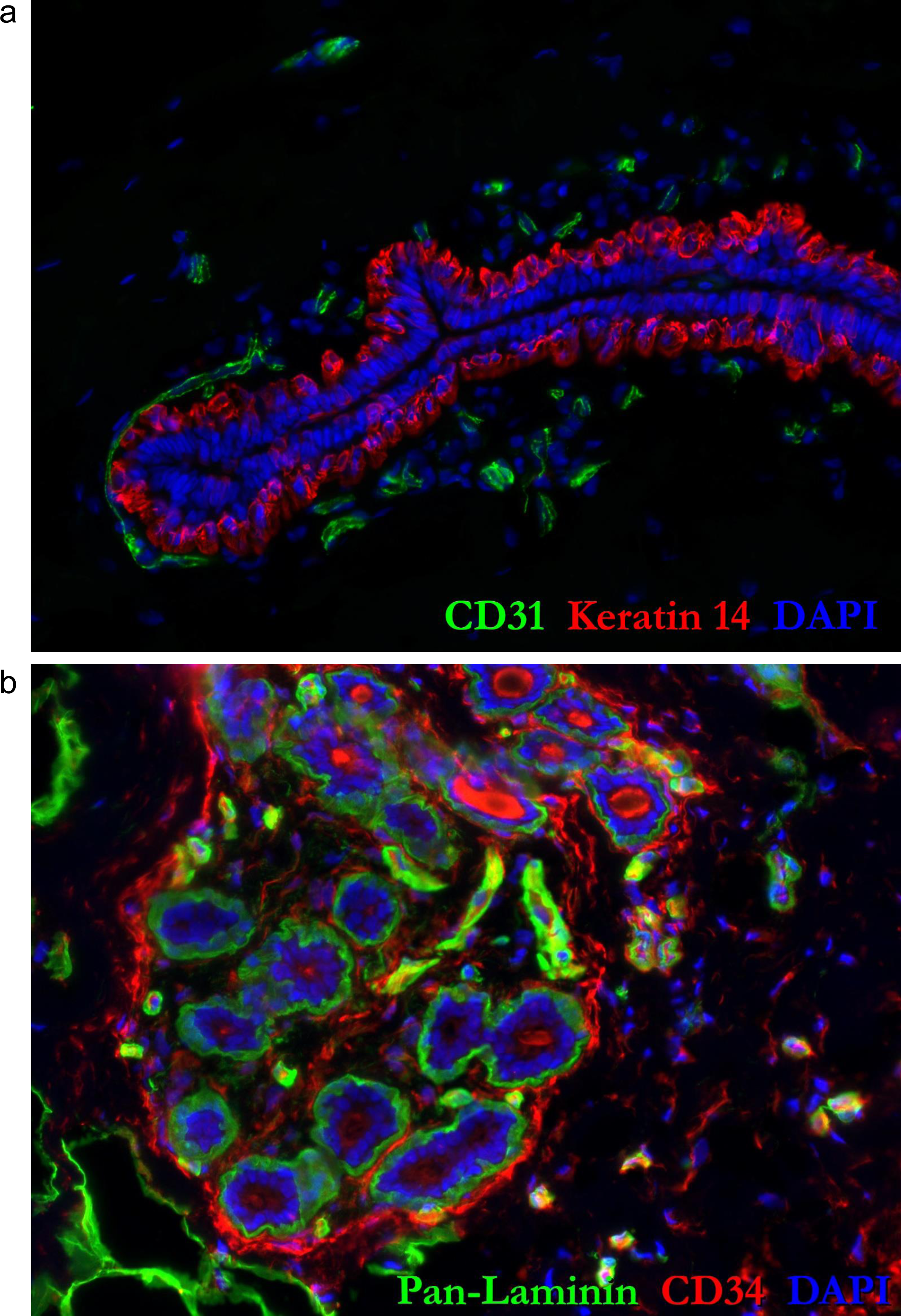
Full resolution images of Fig.8a-b

**Figure 8—figure supplement 2.**
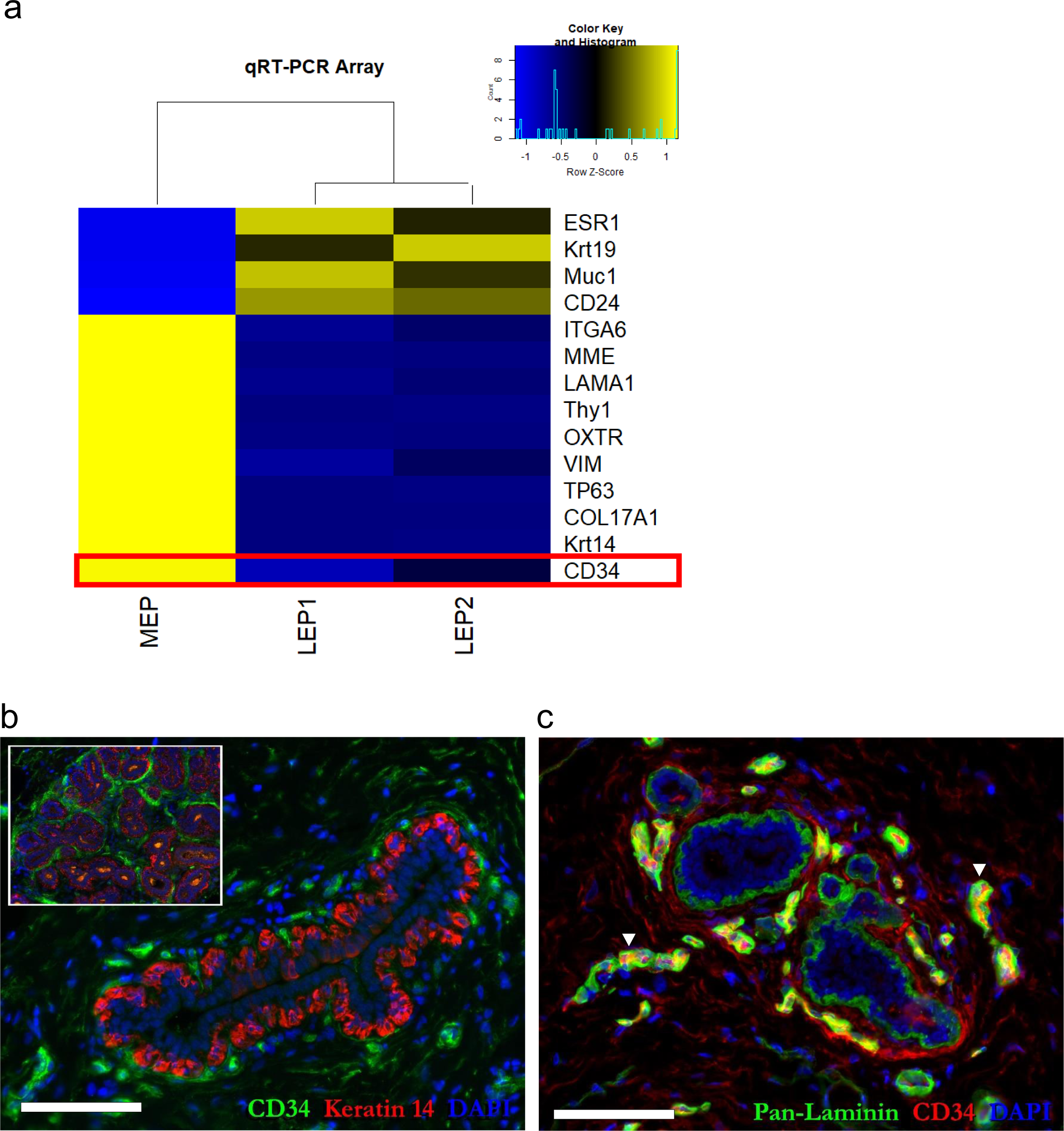
CD34 mRNA found in FACS-isolated myoepithelial fraction reveals endothelial contamination. **(a)** Heatmap of selected mRNA transcripts from a custom qRT-PCR array. mRNA levels were measured in FACS-sorted luminal and myoepithelial cell fractions (LEP1-CD24^Pos^CD49f^Neg^; LEP2-CD24^Pos^CD49f^Neg^; and MEPs-CD24^Neg^CD49^Pos^). The high CD34 level measured in FACS-sorted MEPs (red outline) was ultimately traced to endothelial cell contamination of the MEP gate. **(b)** Supporting this conclusion are tissues immunostained for CD34, demonstrating a lack of CD34 within keratin 14^Pos^ myoepithelial cells in a lactiferous duct or lobules (inset). **(c)** CD34 staining of capillary endothelial cells co-stained with pan-laminin (two of which are marked with ‘▾’).

**Figure 8—figure supplement 3.**
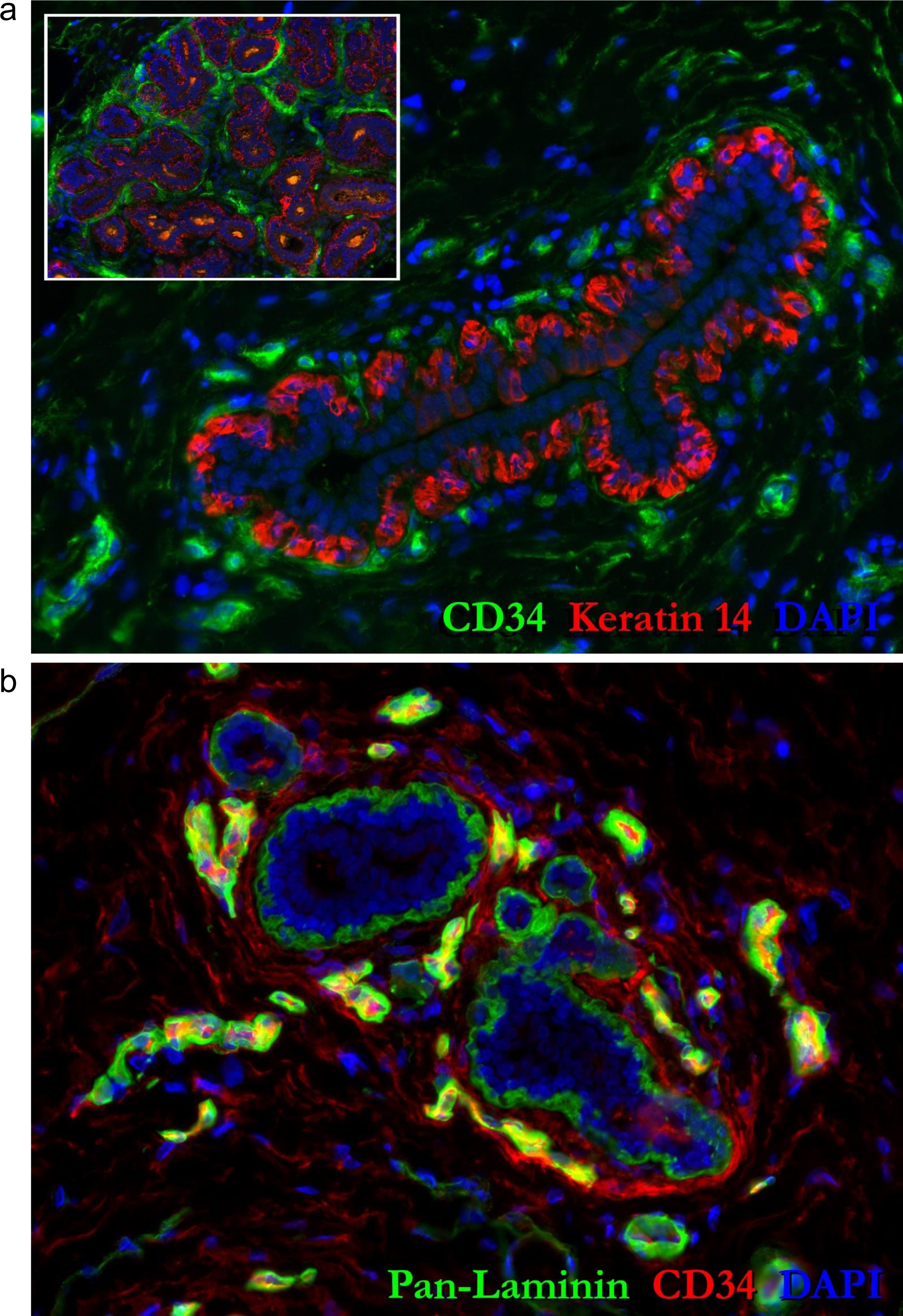
Full resolution images of Fig.8 supplement 2b,c

**Figure 9 - figure supplement 1.**
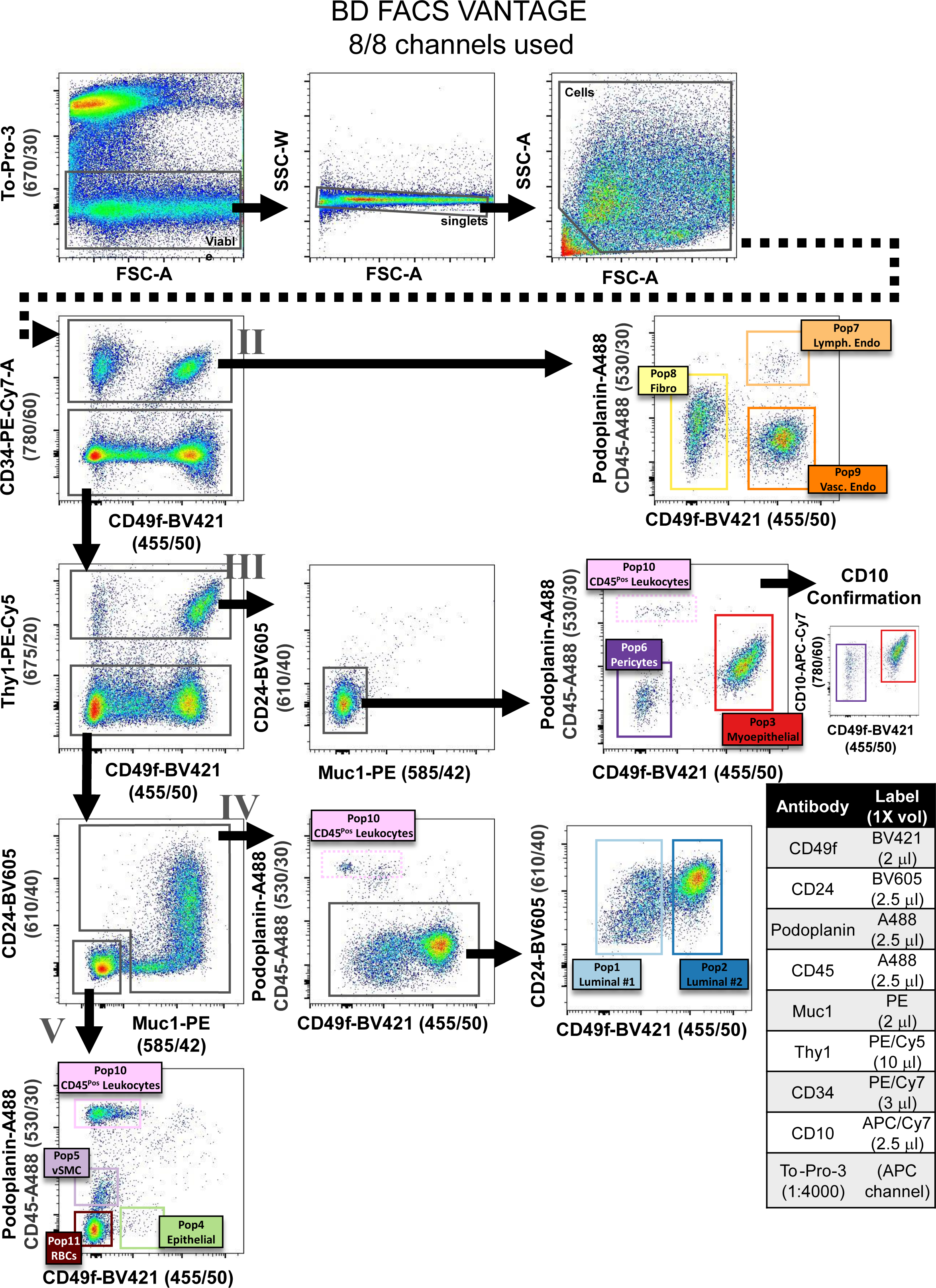
FACS strategy designed for an 8 channel FACS machine (BD FACS Vantage). This FACs gating scheme was used to isolate cell types for primary cultures and RNA-sequencing. The displayed data are of cells derived from normal breast tissue (reduction mammoplasty tissue from a 22-year-old female, sample #N239), and are representative of the 30+ samples analyzed. Because the number of antibodies required for cell sorting exceeded the number of available channels on this machine, both podoplanin and CD45 were used in the Alexa-488 channel (after validation experiments demonstrated a lack of marker overlap and feasibility of this strategy). The sizeable fluorescent spillover between PE-Cy5 and To-Pro-3 and heavy compensation typically needed to use these dyes together was avoided because PE-Cy5 (and all other markers) was analyzed only in To-Pro-3 negative cells (viable cells).

**Figure 9 - figure supplement 2.**
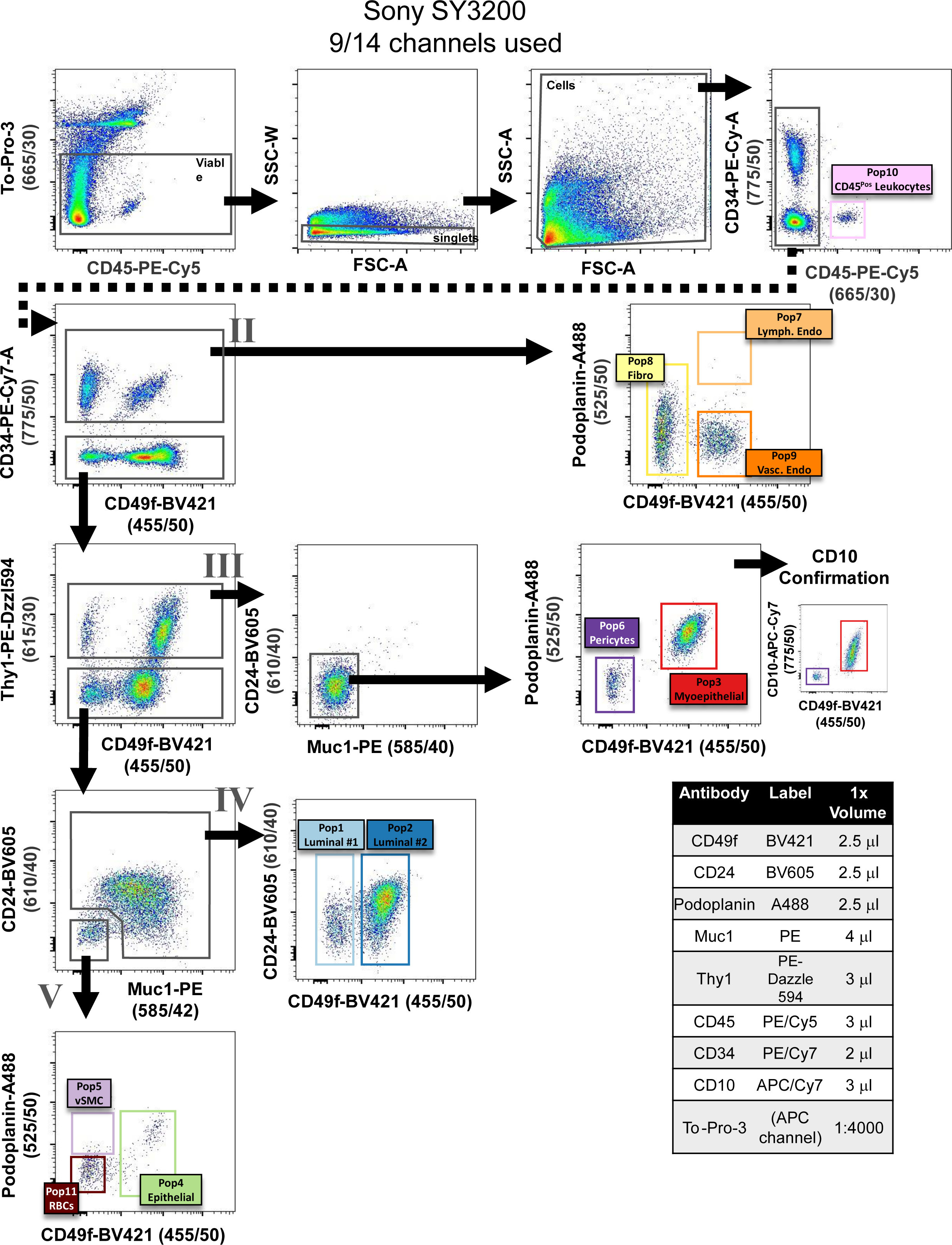
9-color FACS gating strategy optimized for a 14 channel FACS machine (Sony SY3200). This scheme was used to isolate cell types for primary cultures. Fluorochromes were exchanged and matched to this machine’s dichroic filter arrangement. CD45 was moved to an independent channel (PE-Cy5) and Thy1 was moved to PE-Dazzle-594. To-Pro-3 was again used as a viability dye (instead of DAPI) to permit use of BV421 (CD49f). The data displayed are cells from reduction mammoplasty breast tissue derived from a 37-year-old female (sample #N293).

**Figure 10—figure supplements 1-12.**
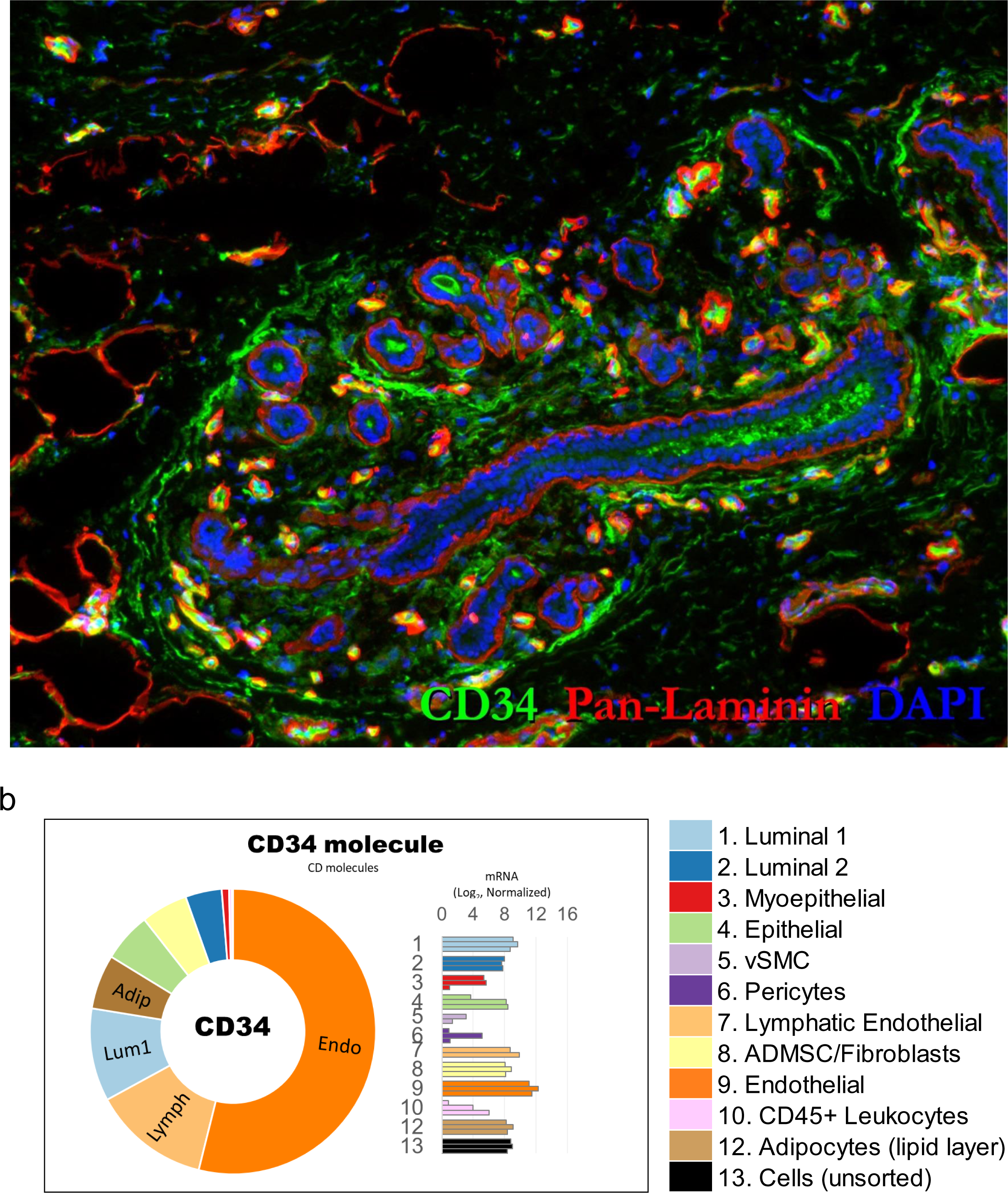
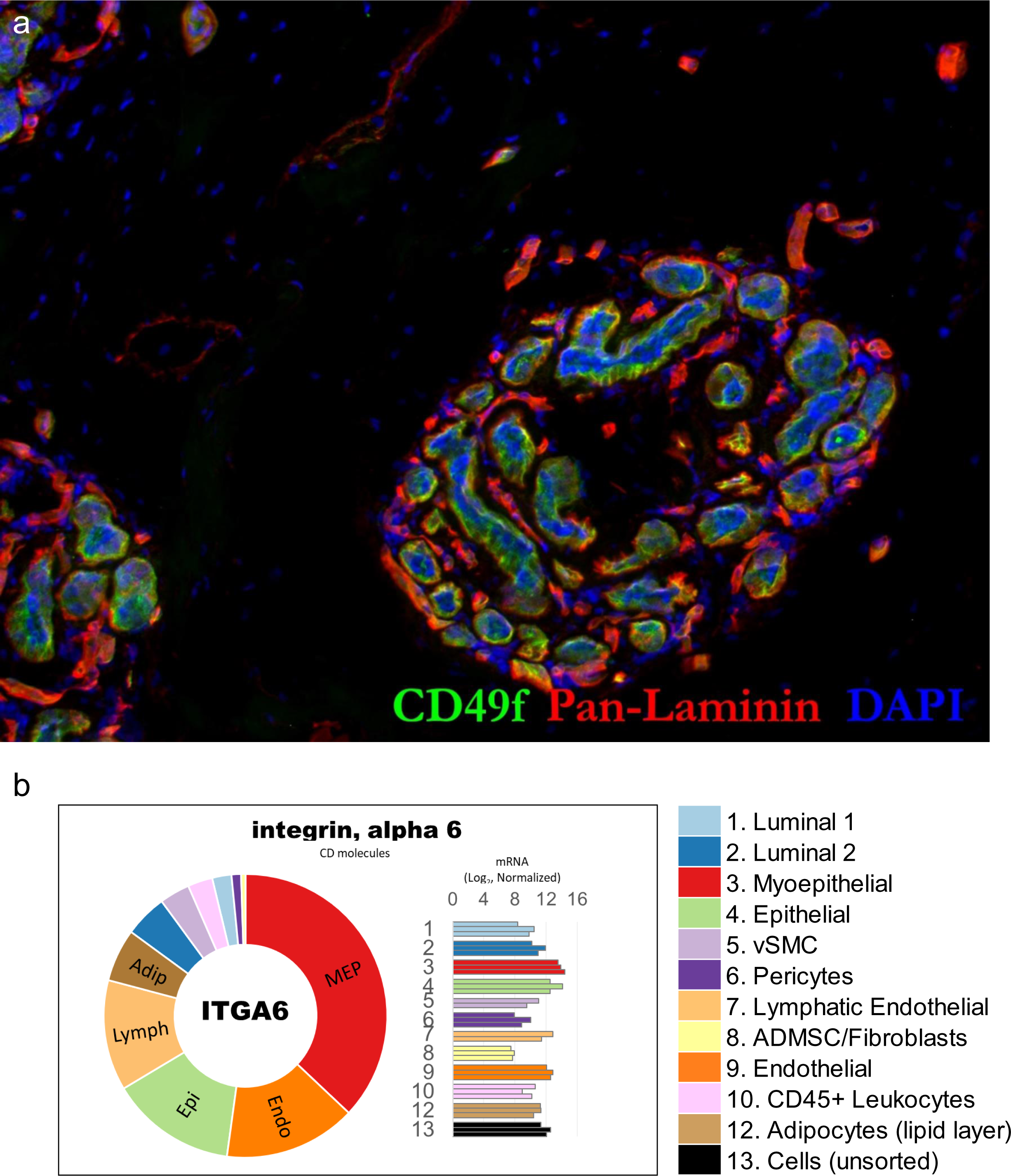
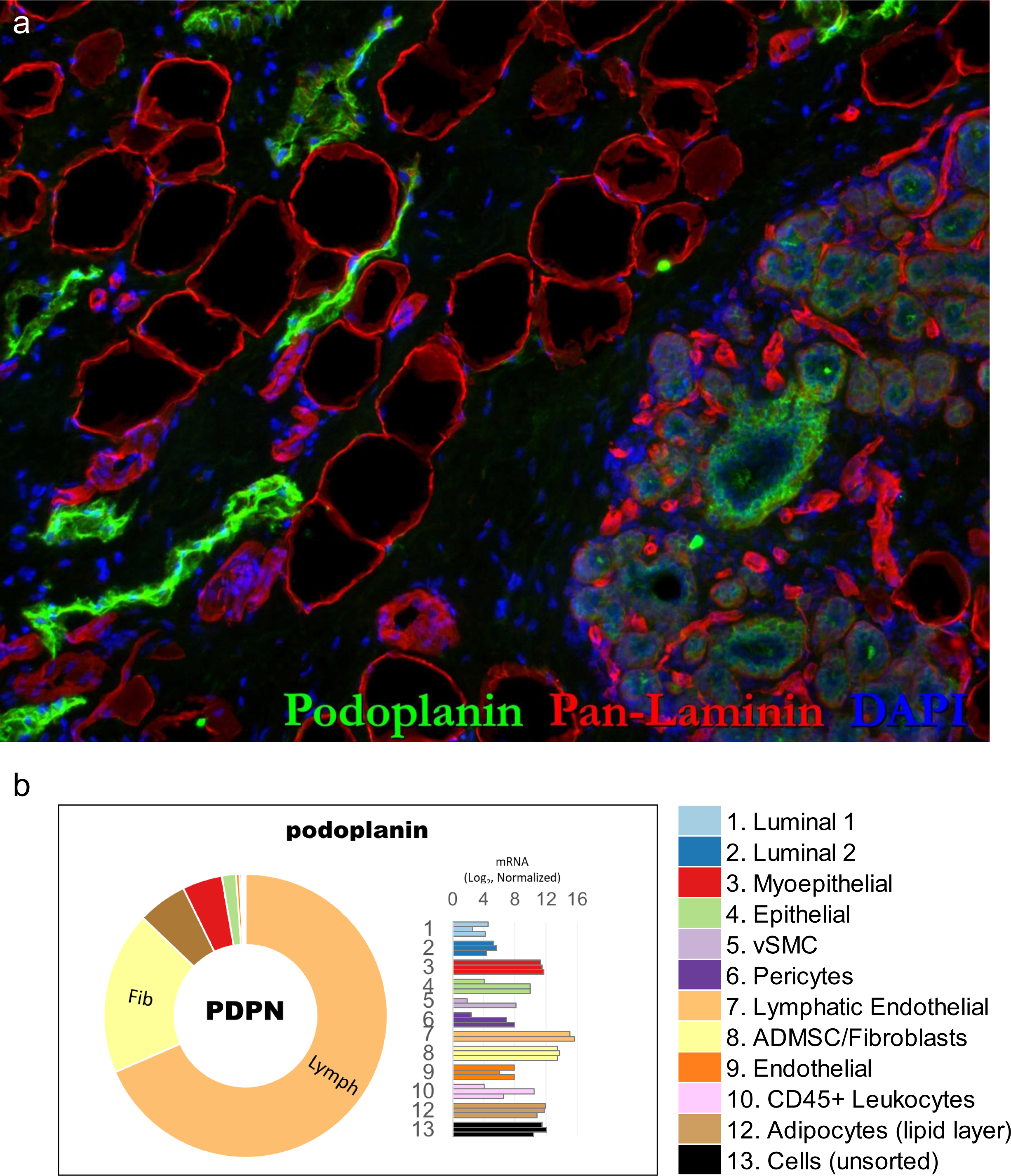
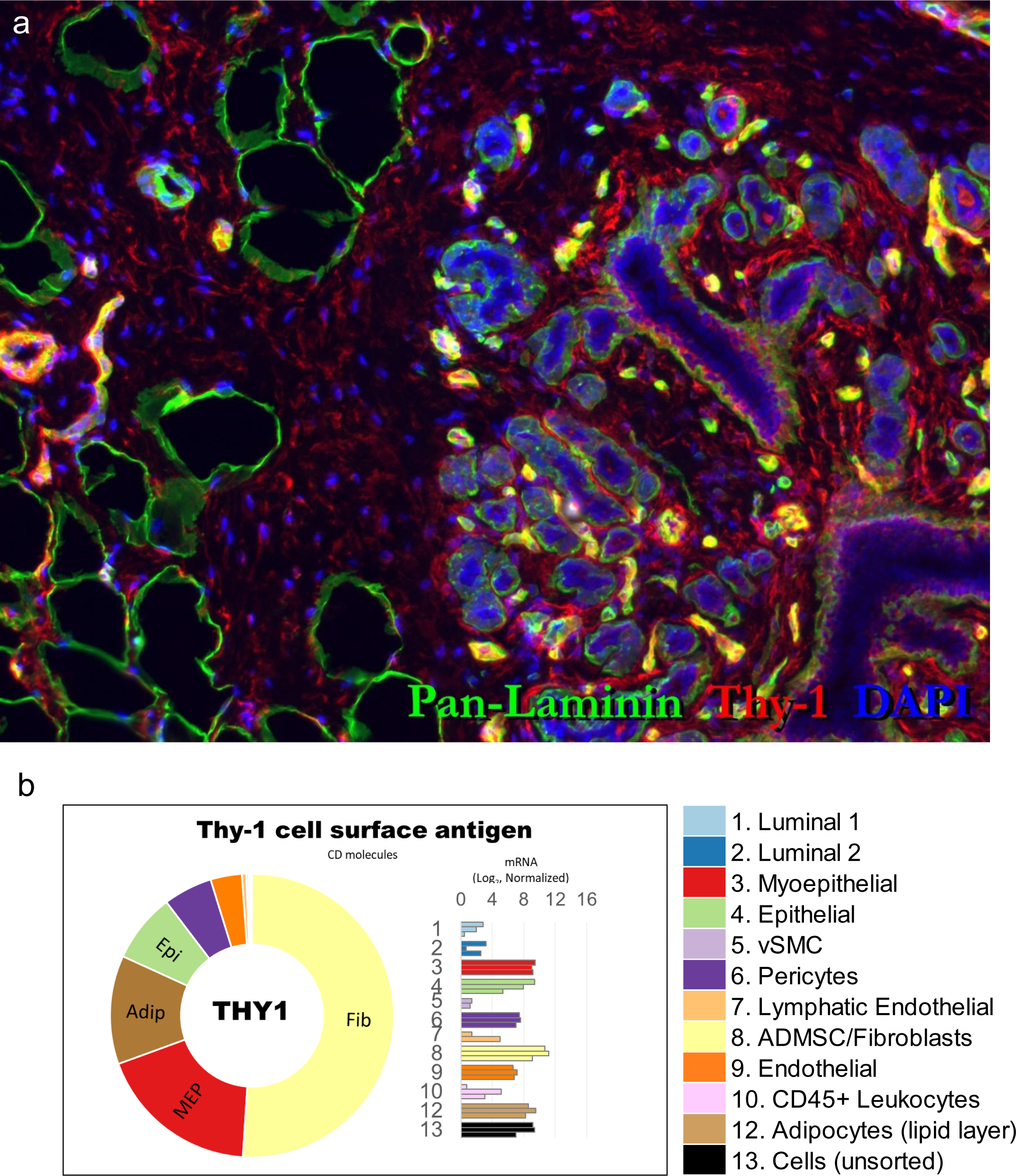
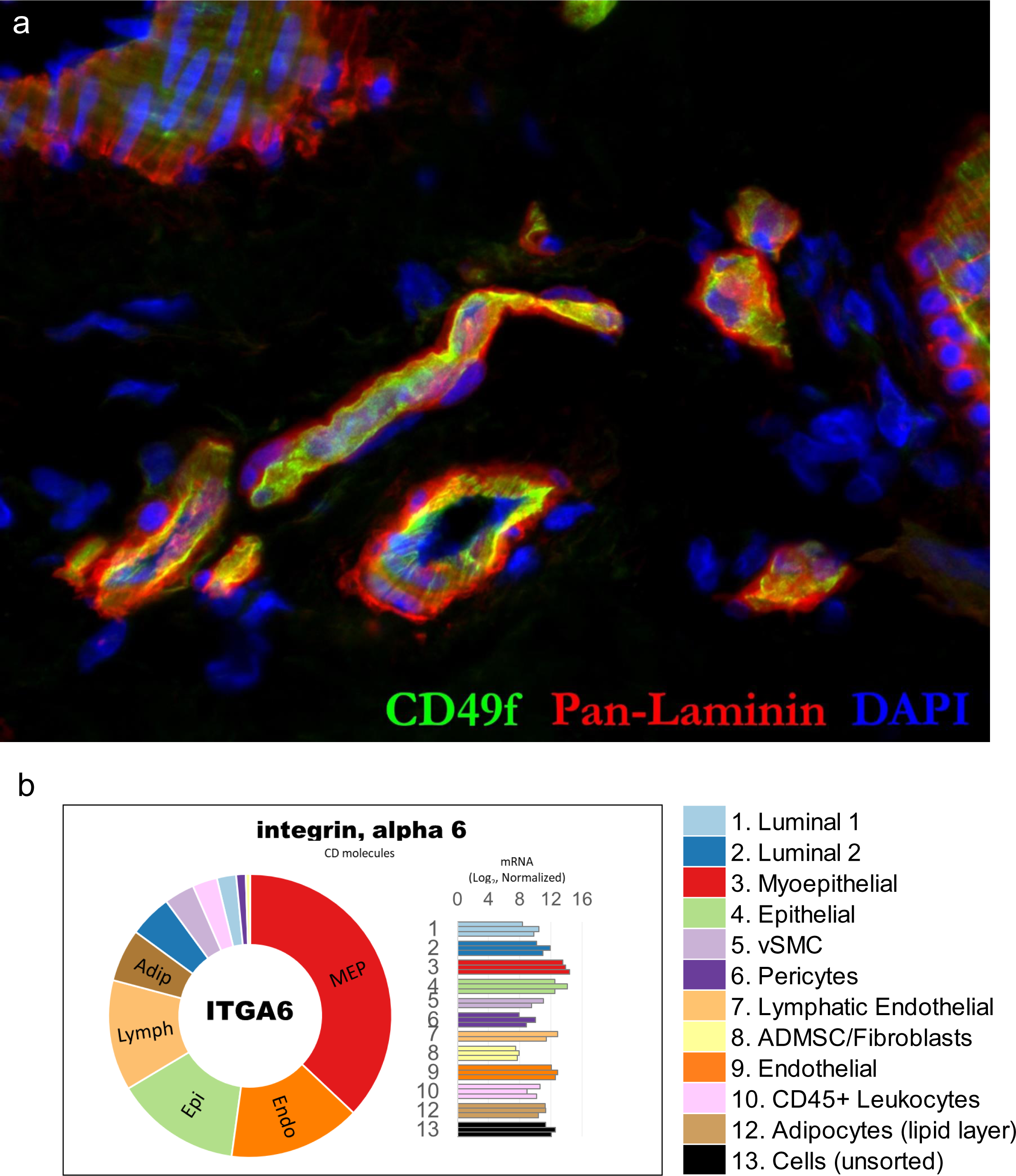
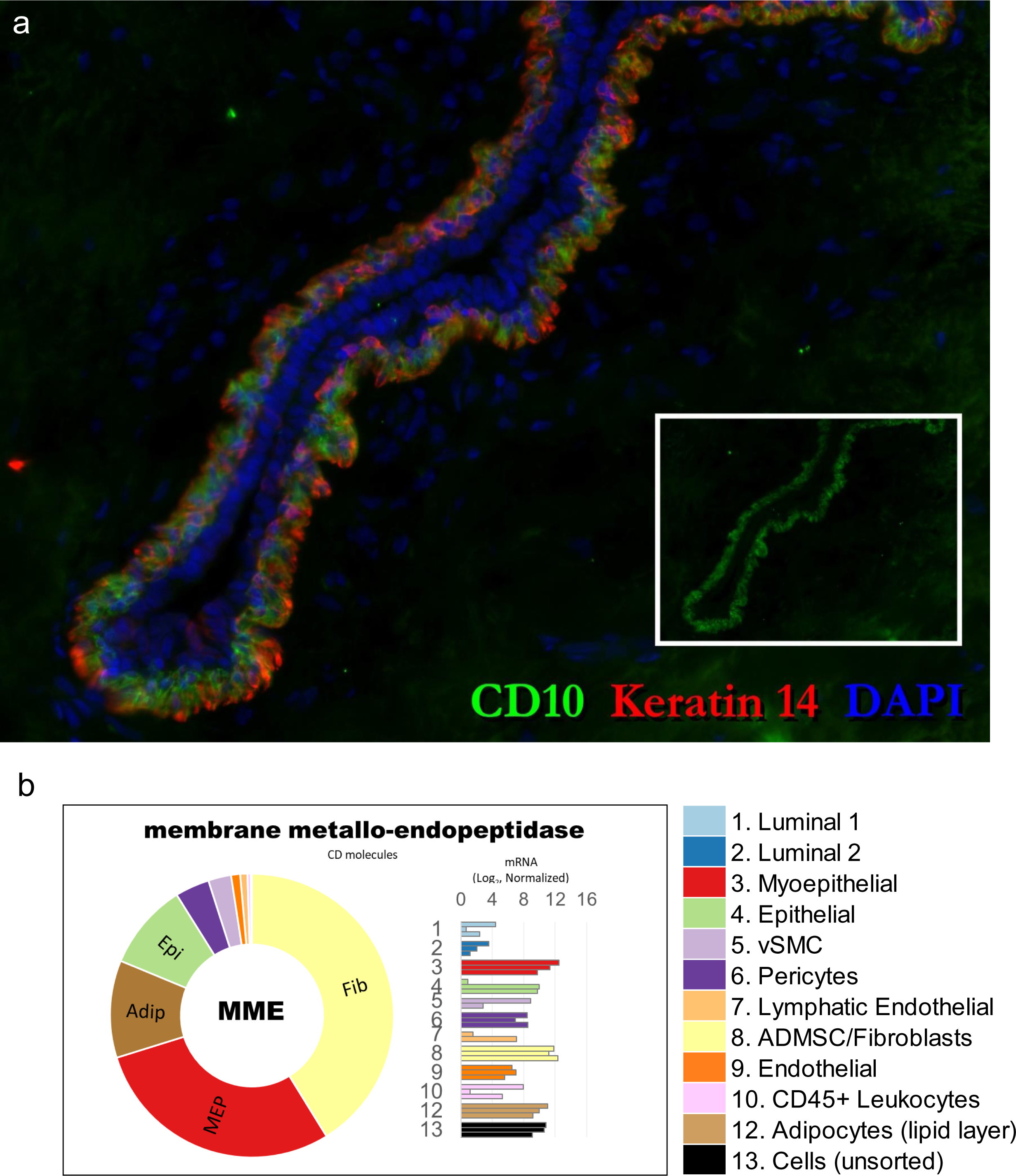
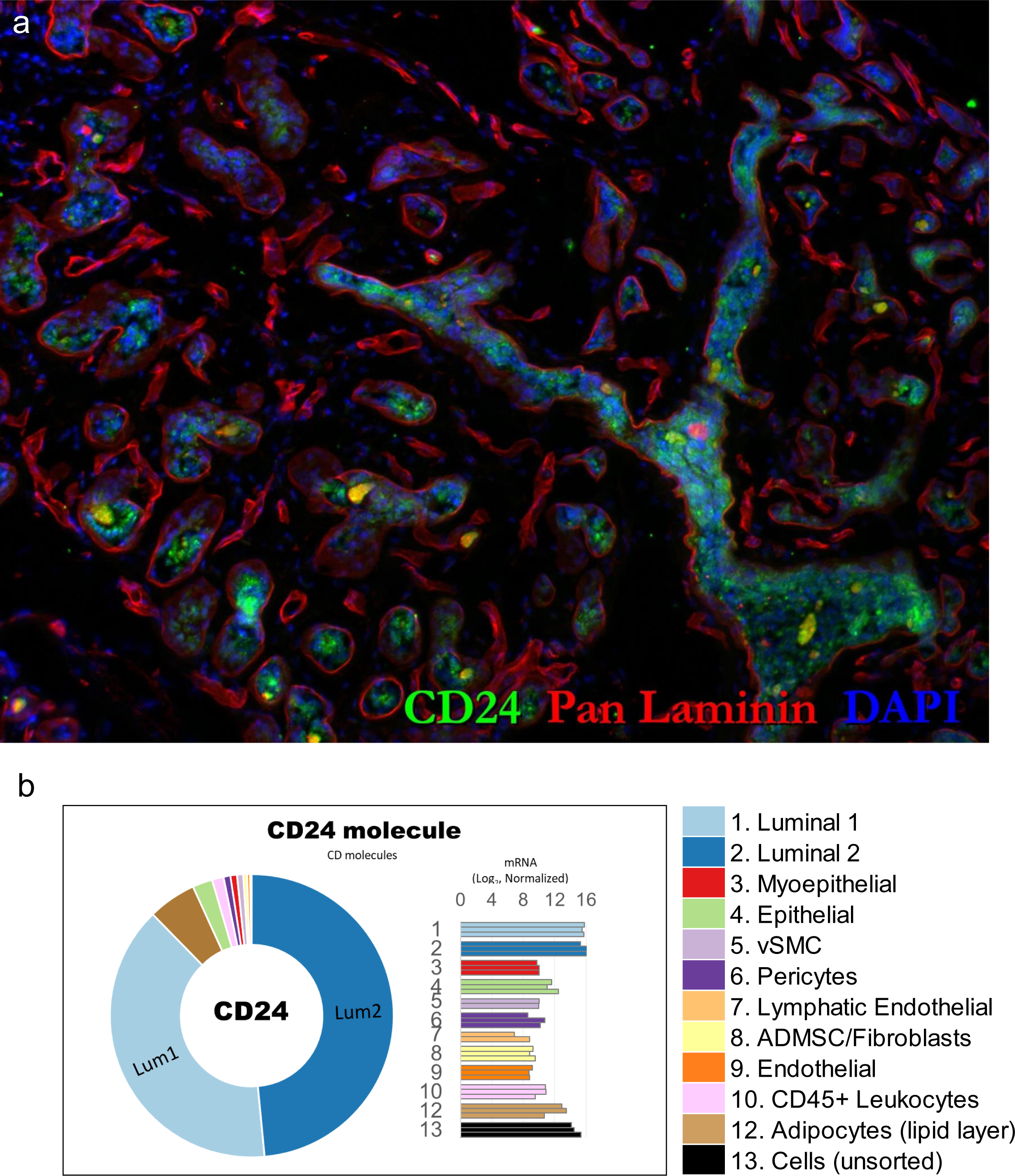
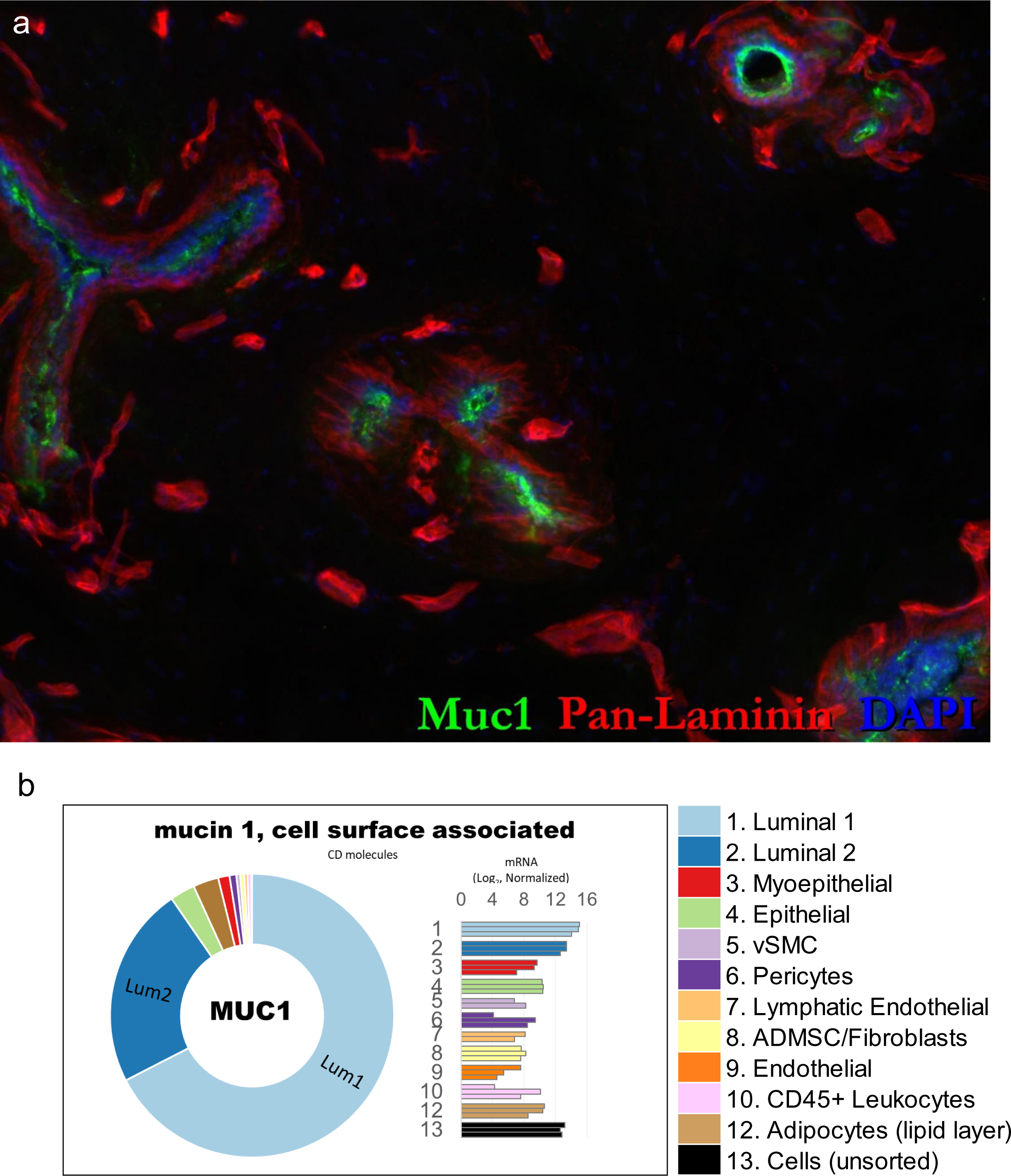
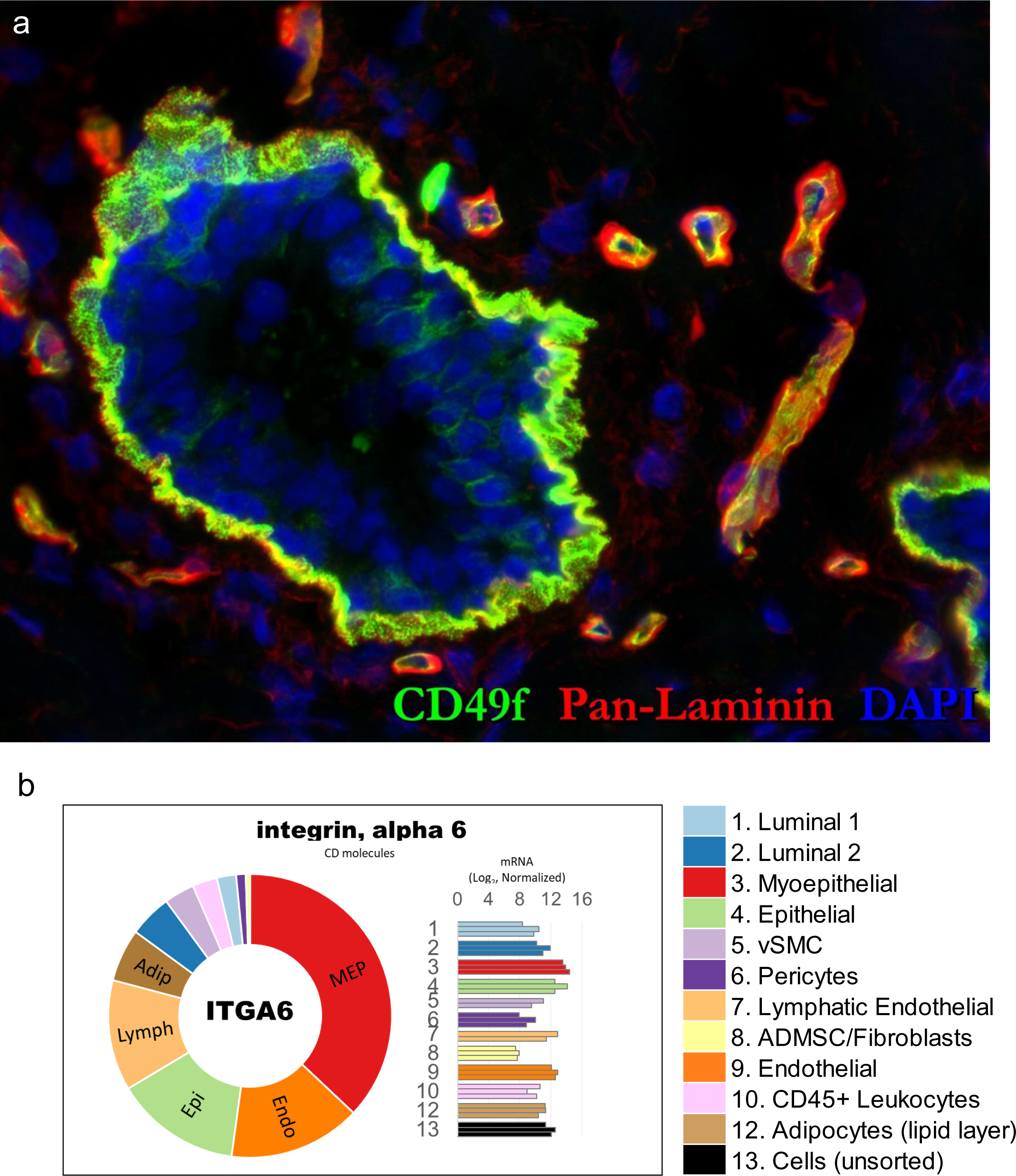
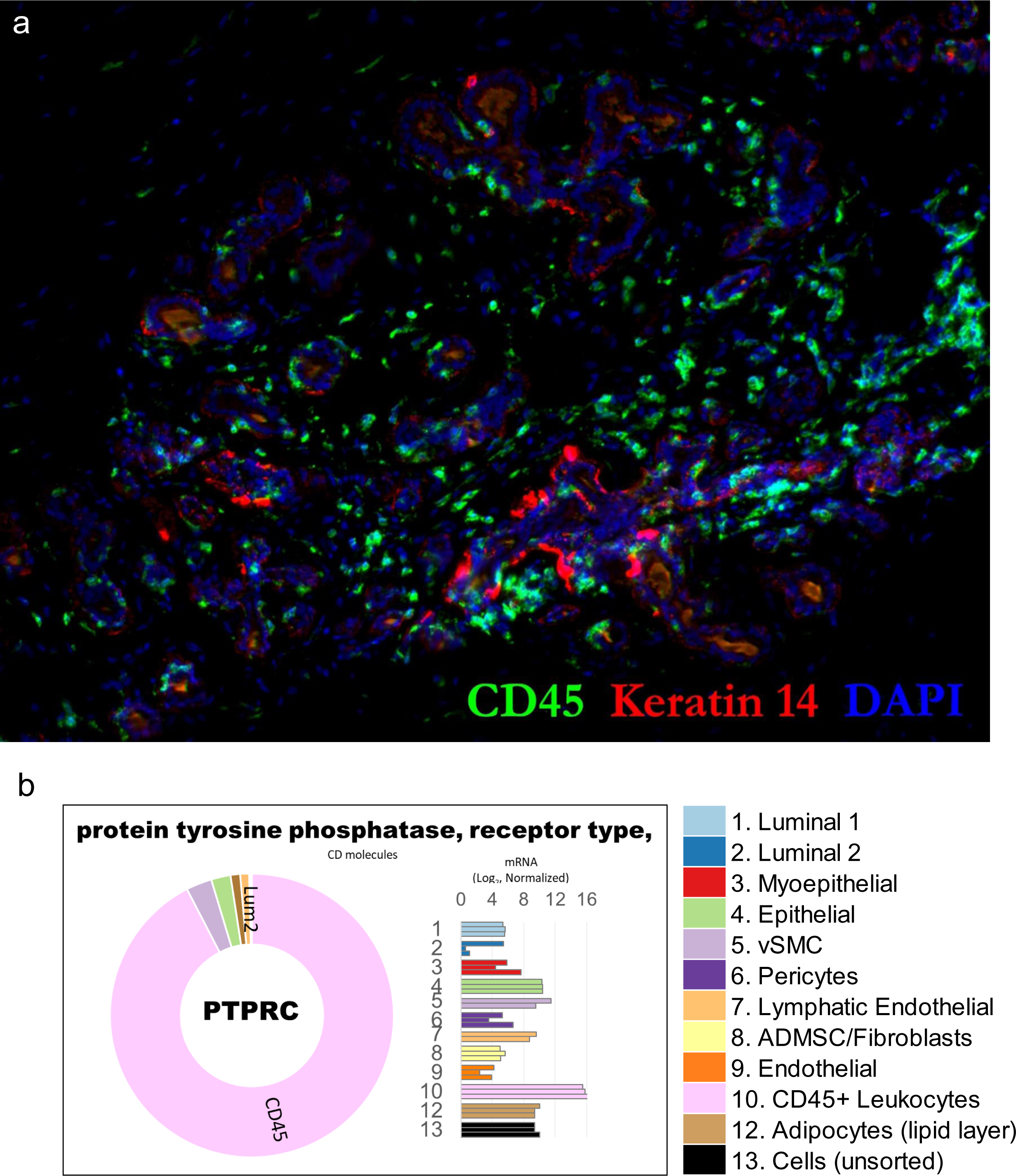
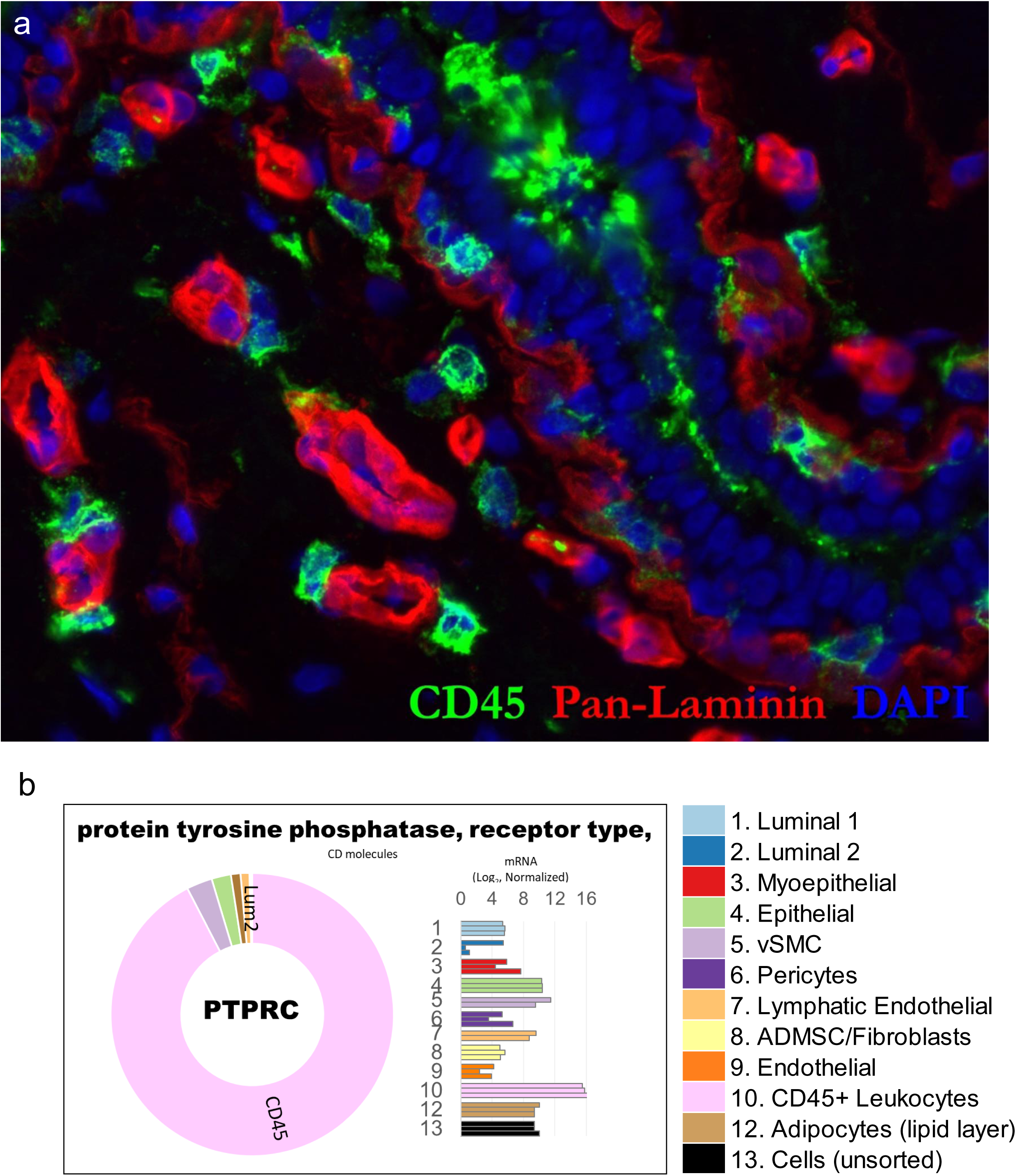
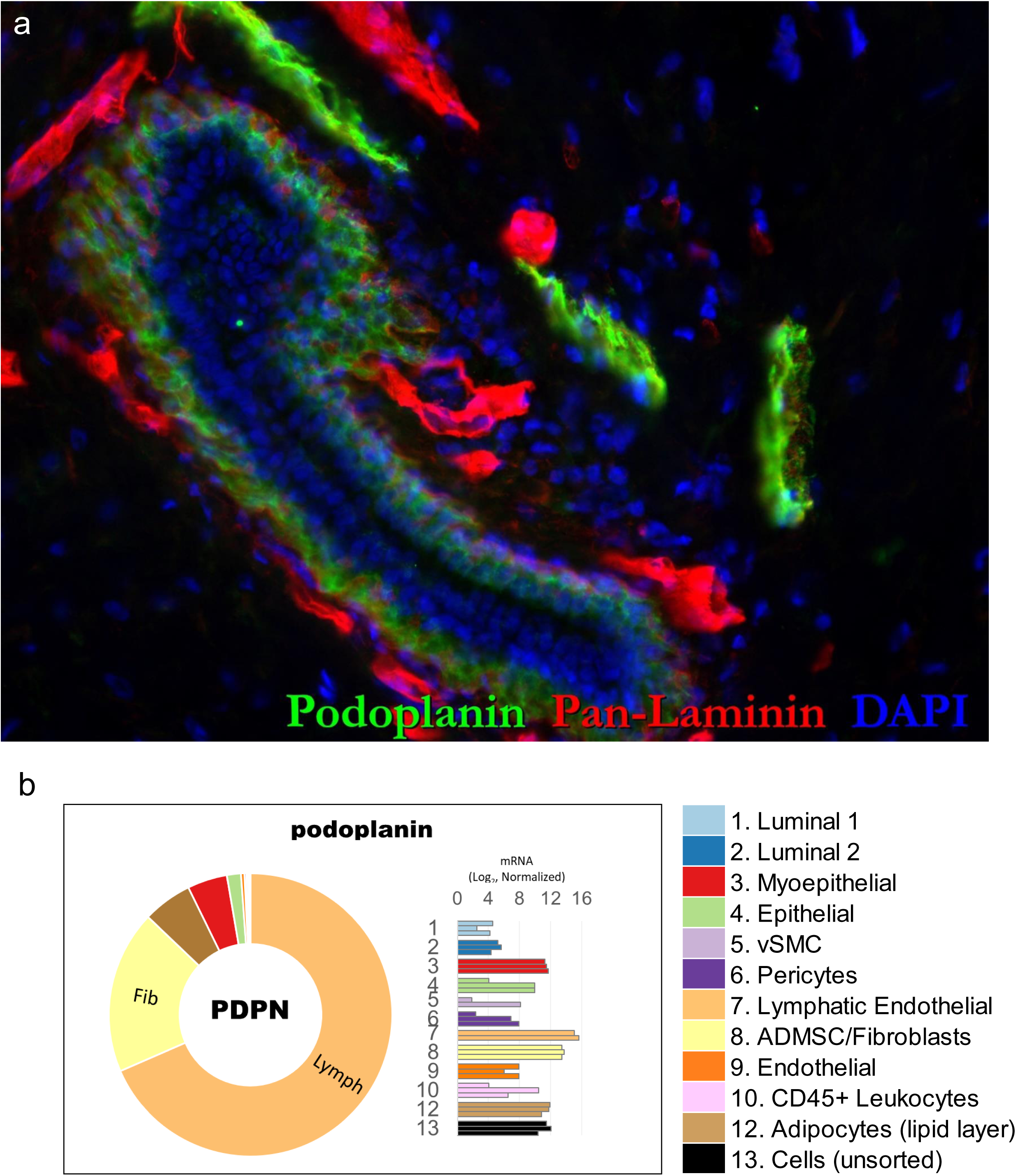
FACS markers: tissue staining and transcript levels. **(a)** Full resolution images of marker-stained normal breast tissues provided in Figs.10 a-l, along with **(b)** transcript levels measured in the FACS-purified cell types, as determined by RNA-sequencing. Normalized mRNA values (rlog, DEseq2) are provided on log2 scale (bar graph of each biological replicate) and linear scale (donut graph of median value), which are both color-coded by cell type.

**Figure 11 – figure supplement 1.**
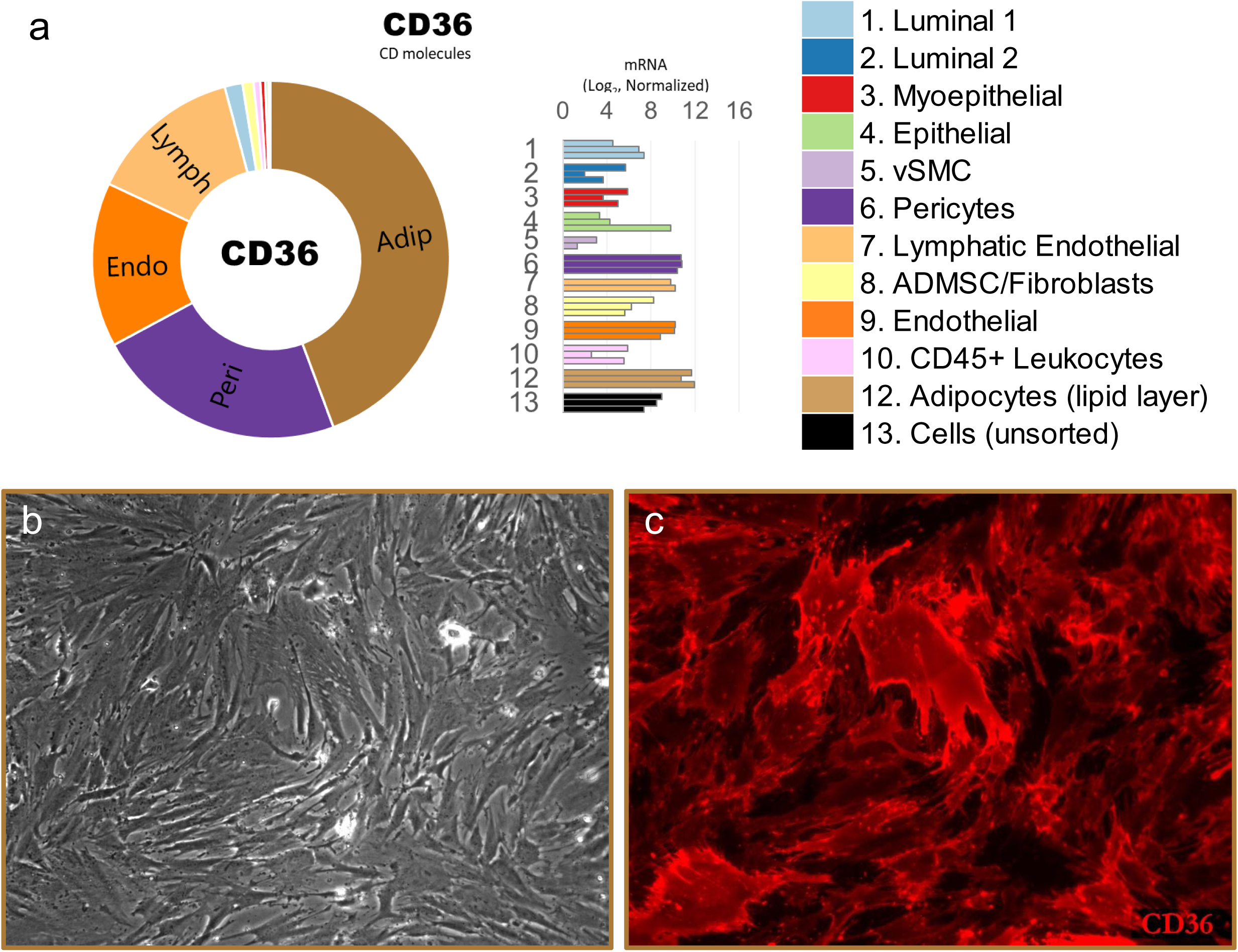
Adipocyte expression of CD36. **(a)** RNA-sequencing identified the cell surface protein CD36 as being differentially expressed by adipocytes (and pericytes). **(b)** Phase-contrast image of primary adipocyte culture. **(c)** CD36 immunostained culture of adipocytes.

**Figure 11—figure supplement 2.**
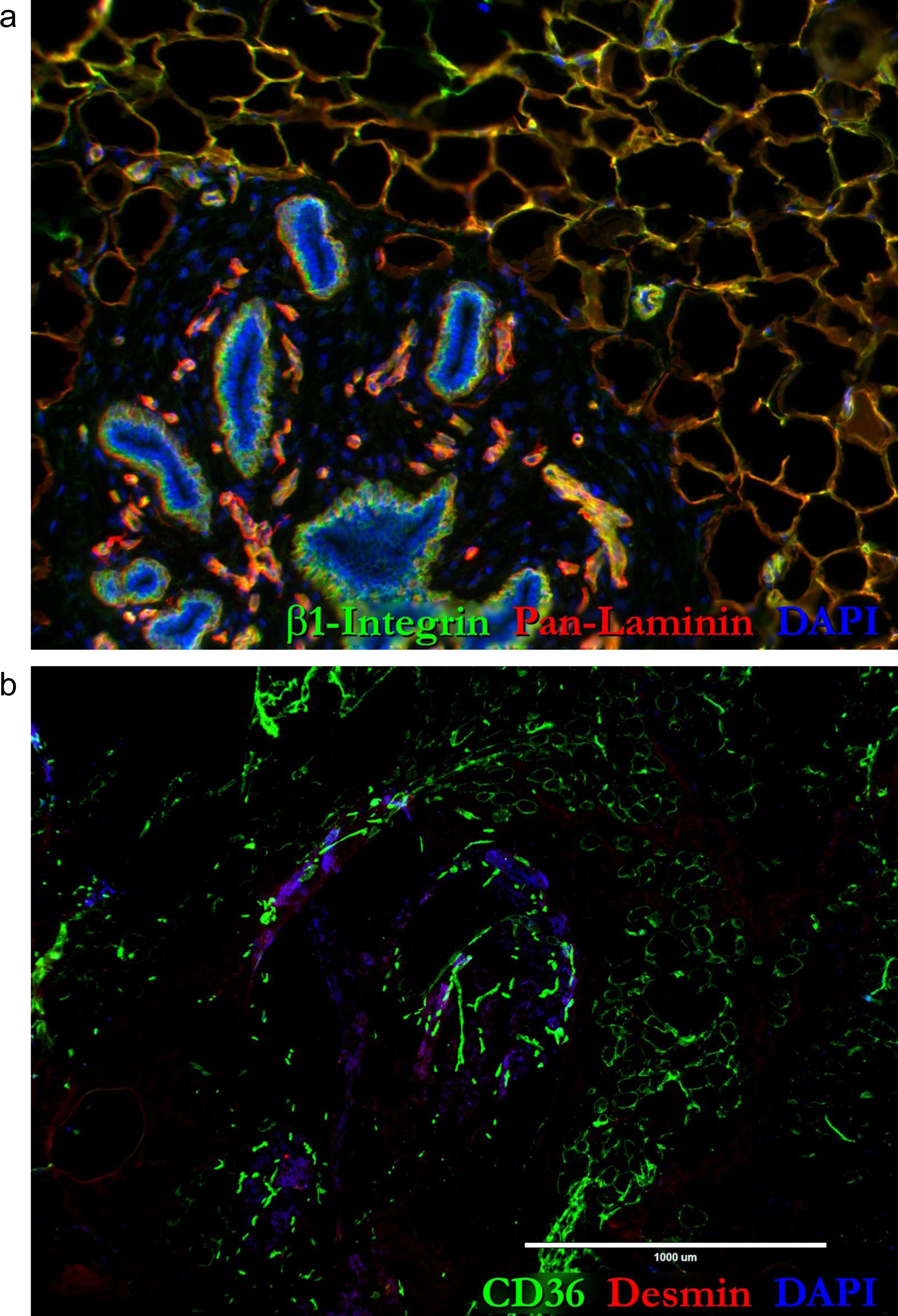
High-resolution images (a) High-resolution images of the immunostained breast tissues provided in Figs.11e,f.

**Figure 12 – figure supplement 1.**
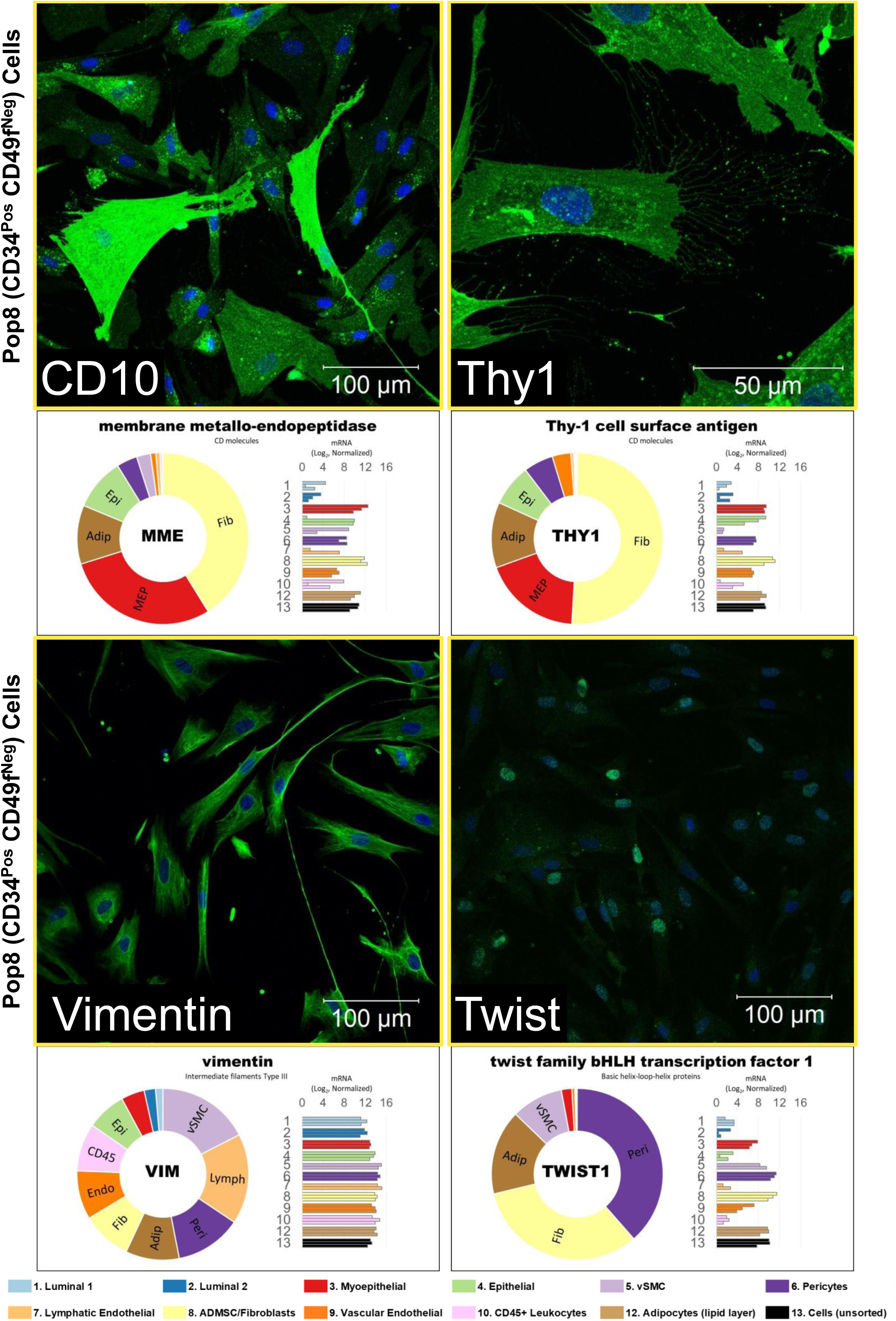
High resolution images of the immunostained Pop8 fibroblasts provided in Figs.12 g-j. Below each image are the RNA-seq determined transcript levels measured within uncultured FACS-purified cell types.

**Figure 12 – figure supplement 2.**
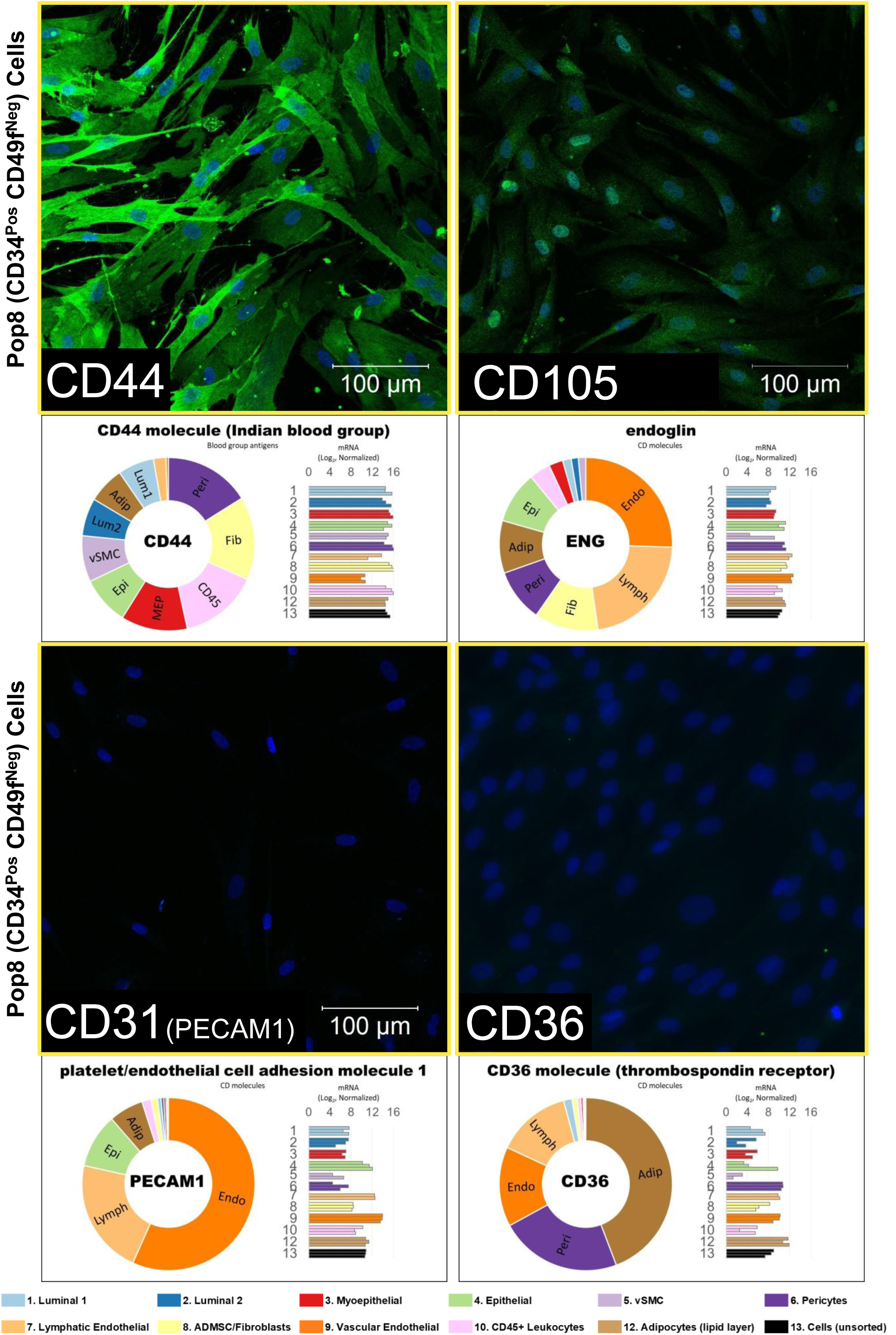
High resolution images of the immunostained Pop8 fibroblasts provided in Figs.12 k-n. Below each image are RNA-seq determined transcript levels measured within uncultured FACS-purified cell types.

**Figure 12 – figure supplement 3.**
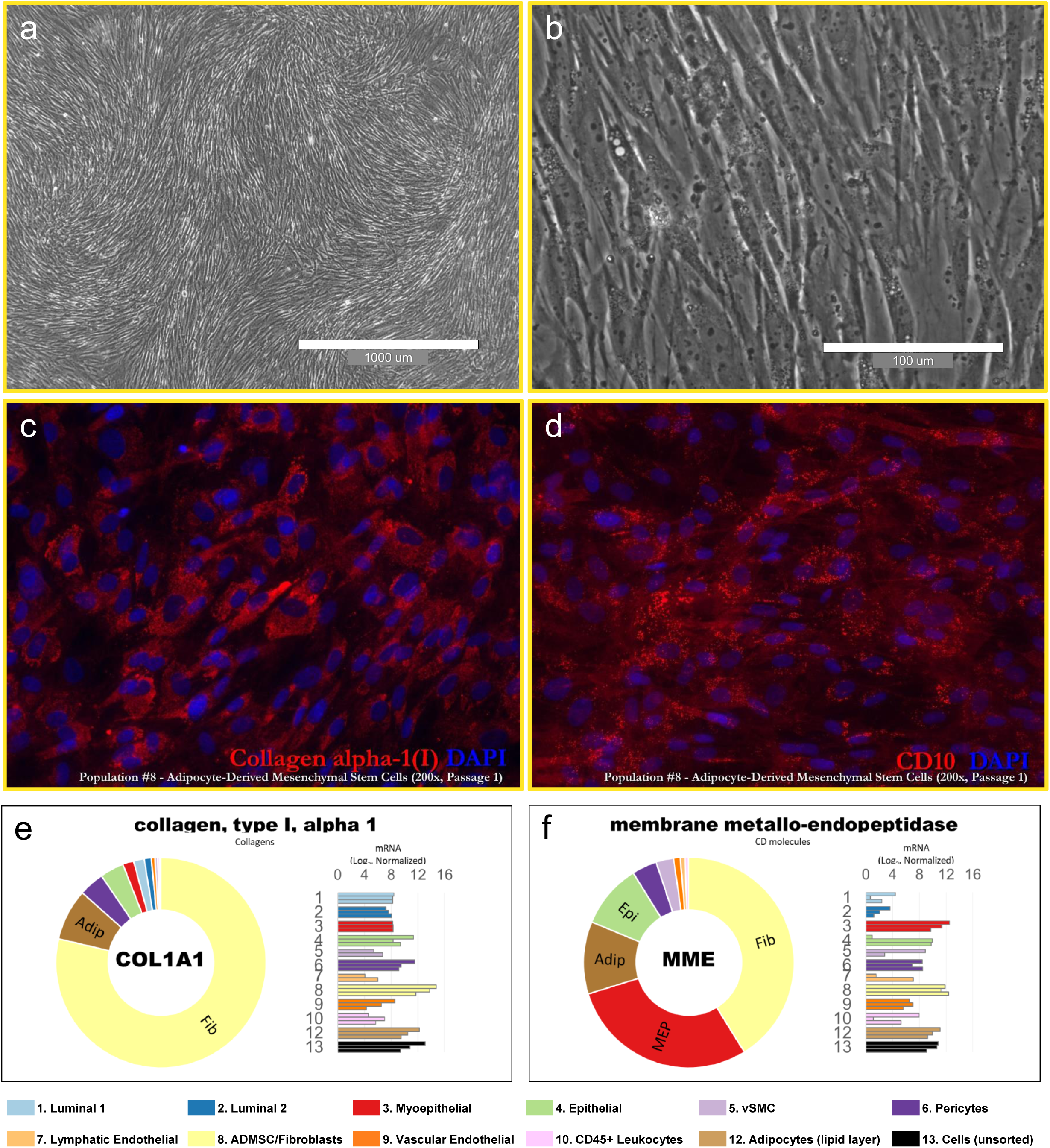
**(a,b)** Primary Pop8 (fibroblast) cultures imaged with (a) 4x and (b) 40x phase contrast objectives (scale = 1000μm and 100μm, respectively; imaged on EVOS imaging station; 4x Ph Plan 0.13 NA objective and 40x Plan LWD FL/PH 0.65NA objective). **(c,d)** Primary fibroblast cultures immunostained for (c) alpha I collagen and (d) CD10. **(e,f)** transcript levels of COL1A1 and CD10 (membrane metallo-endopeptidase, ‘MME’) measured in uncultured FACS-purified cell types, as determined by RNA-sequencing.

**Figure 12 – figure supplement 4.**
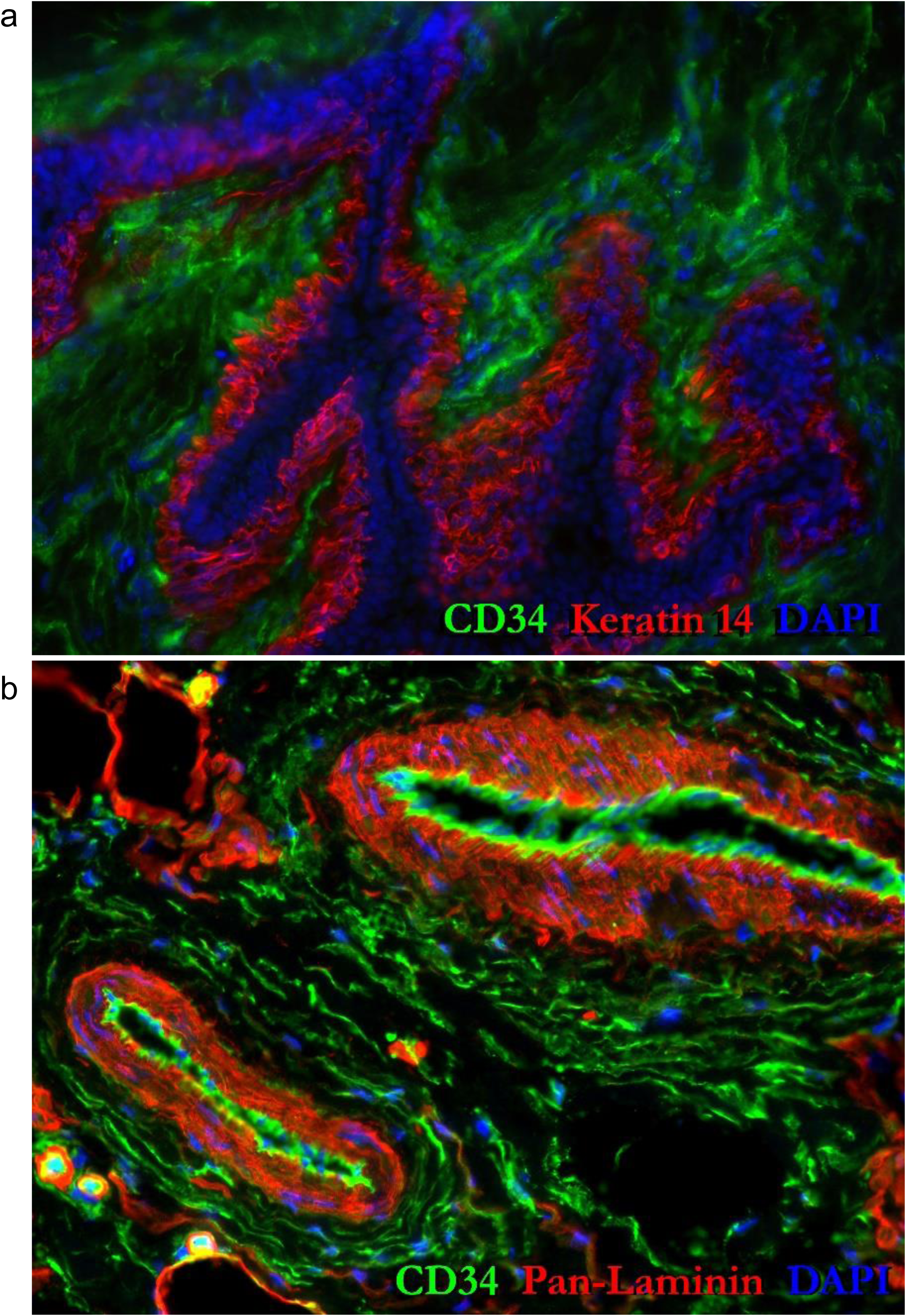
High resolution images of normal breast tissues immunostained for CD34, the main marker used to sort Pop8 fibroblasts (CD34^Pos^CD49f^Neg^). **(a)** CD34/keratin 14 and **(b)** CD34/pan-laminin.

**Figure 12 – figure supplement 5.**
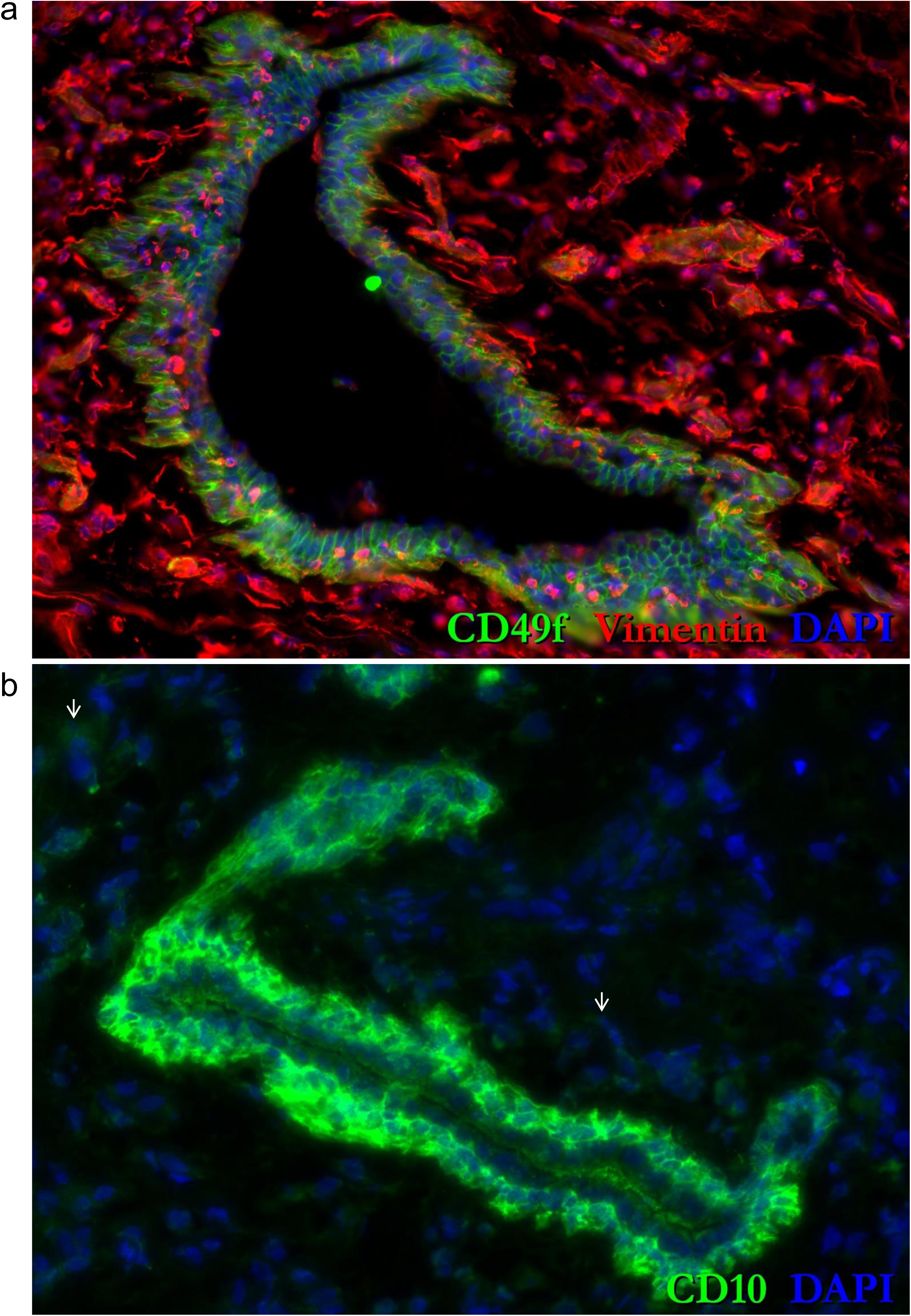
High resolution images of normal breast tissues immunostained for CD49f, a marker used to resolve the CD34^Pos^CD49f^Neg^ Pop8 (fibroblasts) in FACS experiments. **(a)** CD49f/Vimentin and **(b)** Tissue fibroblasts indeed expressed faint levels of CD10 (↑).

**Figure 13—figure supplement 1.**
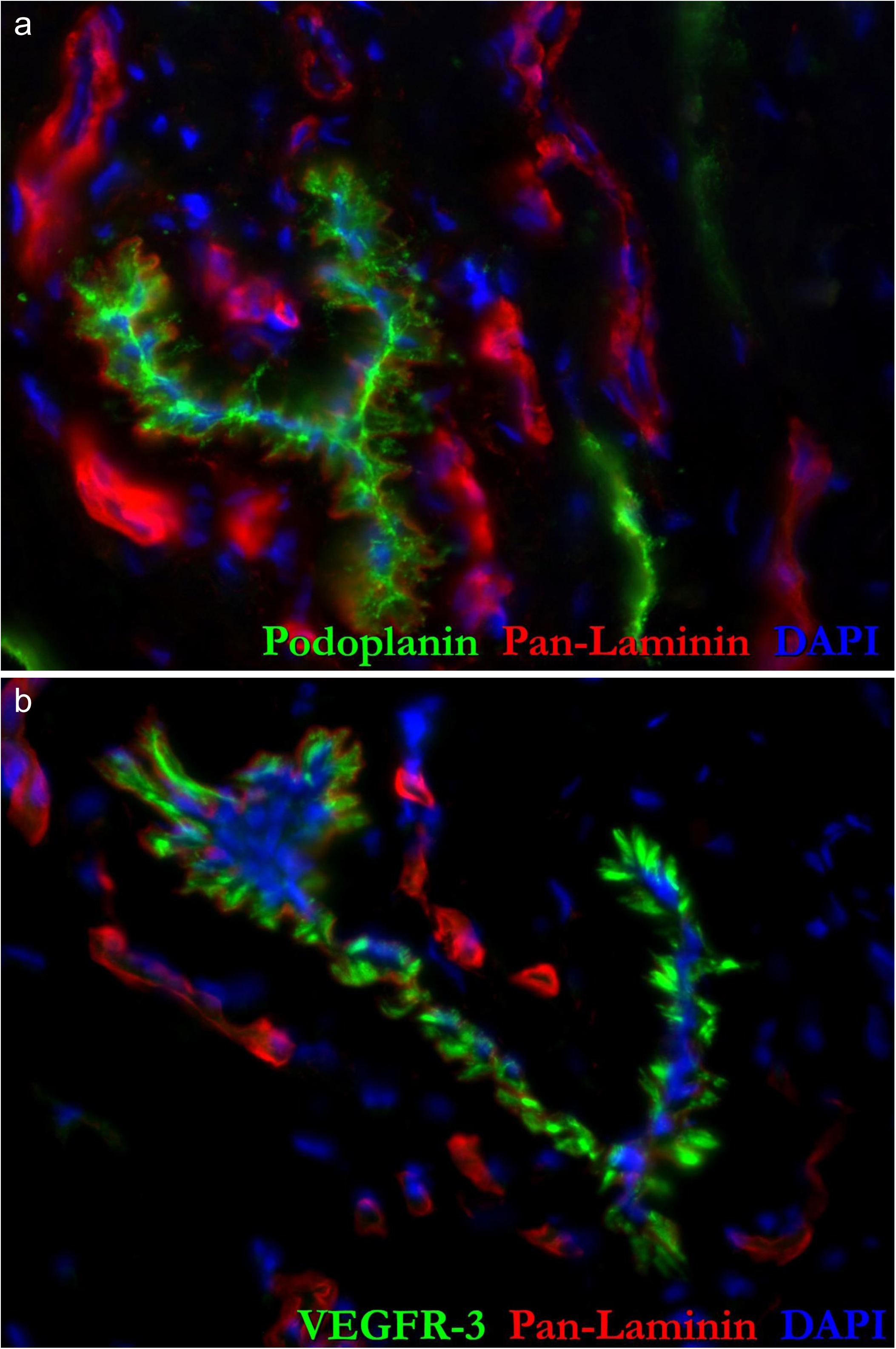
**(a,b)** High-resolution images of (a) podoplanin and (b) VEGFR-3 immunostained normal breast tissue sections provided in Figs.13a,b

**Figure 13—figure supplement 2.**
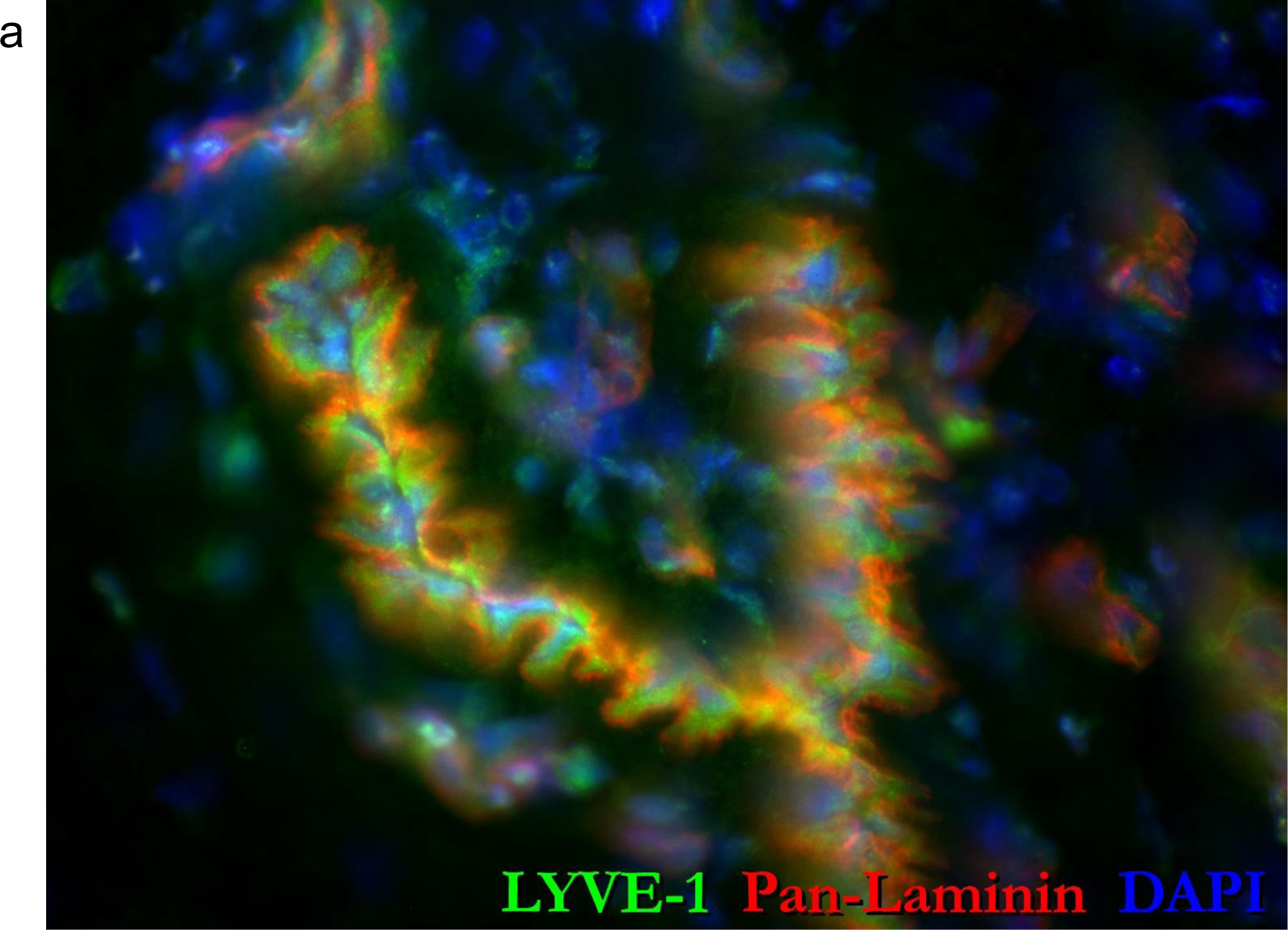
**(a)** High-resolution image of LYVE-1 immunostained breast tissue provided in Fig.13c.

**Figure 13—figure supplement 3.**
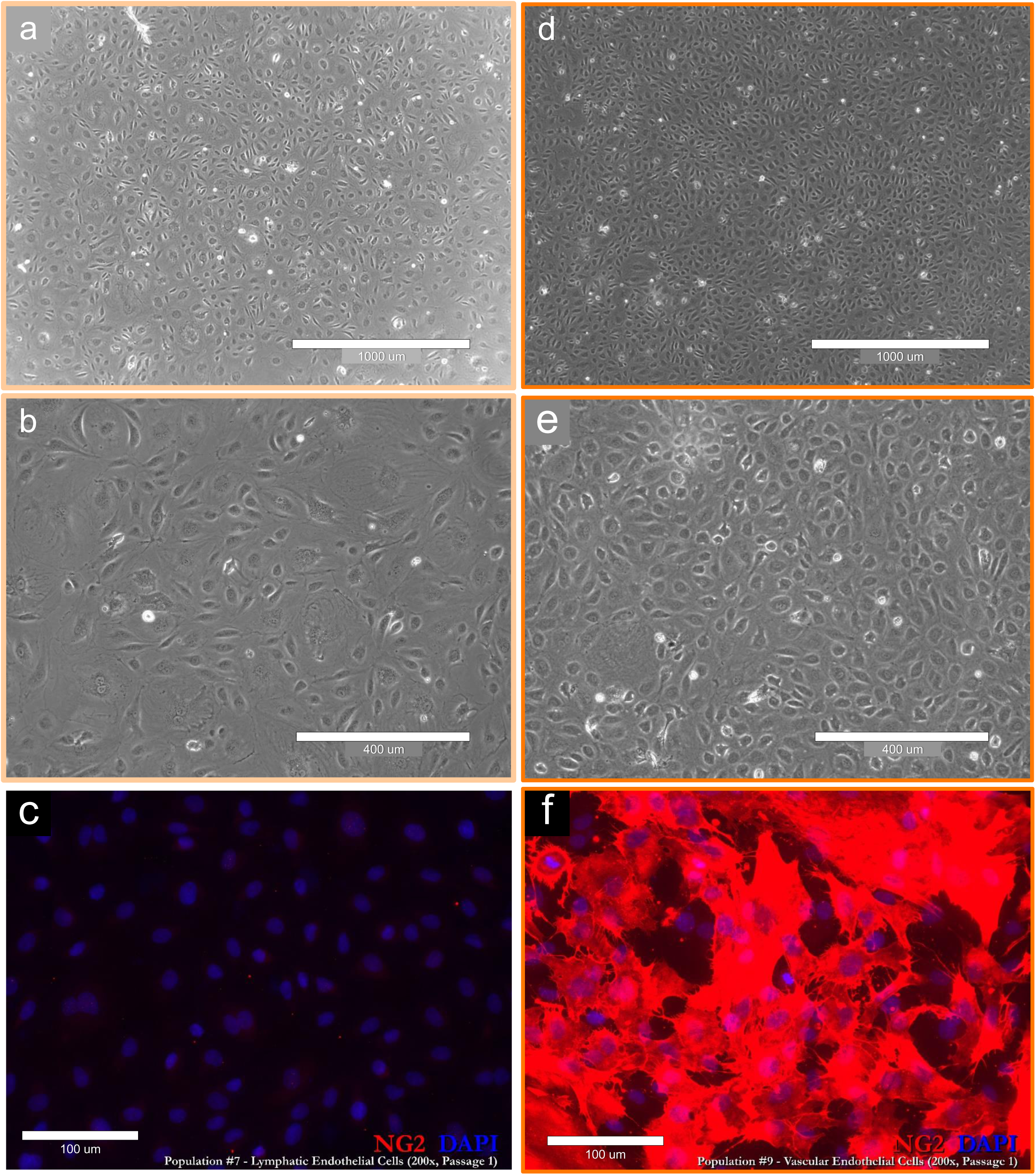
Primary culture of endothelial cell types. Phase-contrast images of Pop7 lymphatic (**a&b**, passage 5) and Pop9 vascular (**d&e**, passage 6) endothelial cell cultures derived from reduction mammoplasty tissues, imaged at low and high resolution (scale=1000μm (a,d); and 400μm (b,e). **(c,f)** Differential NG2 immunostaining of (c) Pop7 lymphatic (uniformly NG2 negative) and (f) Pop9 vascular (uniformly and strongly NG2 positive) endothelial cultures demonstrate the distinct nature of these two cell fractions. Both are passage 1 cells; scale = 100μm. Breast tissues were derived from 26-year-old (#N204; a,b,d,e) and 22-year-old (#N239; c,f) females. Cultured in EGM-2 medium supplemented with an additional 10% FBS (Lonza) at 37°C with 5% CO2.

**Figure 13—figure supplement 4.**
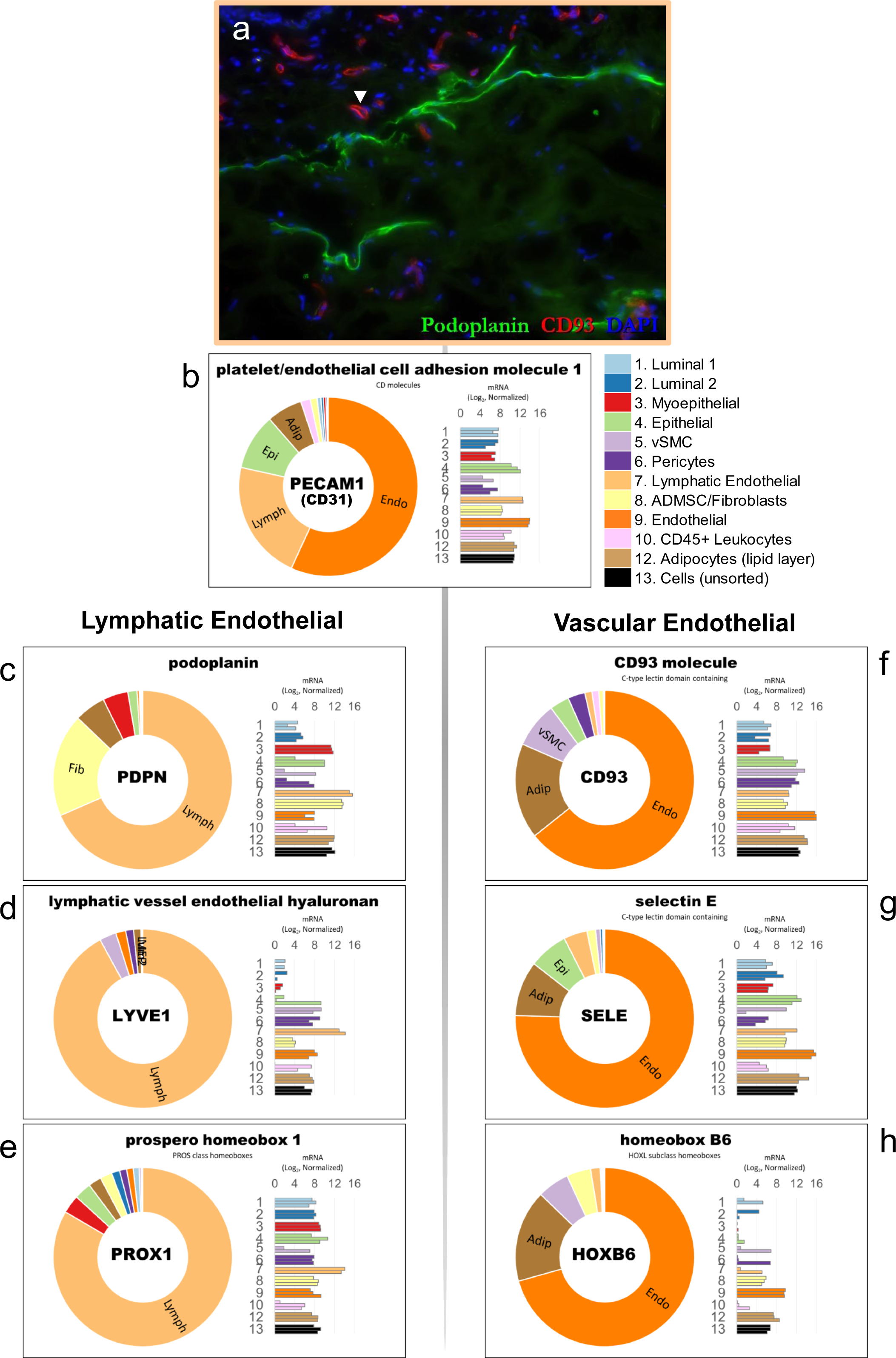
Validation of endothelial cell fractions by RNA-sequencing. **(a)** Normal breast tissue immunostained with podoplanin and CD93 antibodies. The vascular endothelial-specific expression of CD93 (▾) was identified by RNA-sequencing uncultured Pop7 (lymphatic) and Pop9 (vascular) cell fractions. **(b)** PECAM1, the gene encoding the pan-endothelial CD31 antigen, was expressed by both endothelial cell fractions. **(c-e)** Genes encoding the established lymphatic markers: podoplanin (PDPN), lymphatic vessel endothelial hyaluronan receptor 1 (LYVE1), and prospero Homeobox 1 (PROX1), were all differentially expressed by Pop7 cells. **(f-h)** Among the genes differentially expressed Pop9 cells were: CD93, endothelial selectin (SELE), and homeobox protein hox-b6 (HOXB6). For each gene, normalized mRNA values (rlog, DEseq2) are provided on both a log2 scale (bar graph of biological replicates) as well as on a linear scale (donut graph of median values). Each chart is color-coded by cell type; Endothelial cells are orange: lymphatic (light orange) and vascular (dark orange).

**Figure 14—figure supplement 1.**
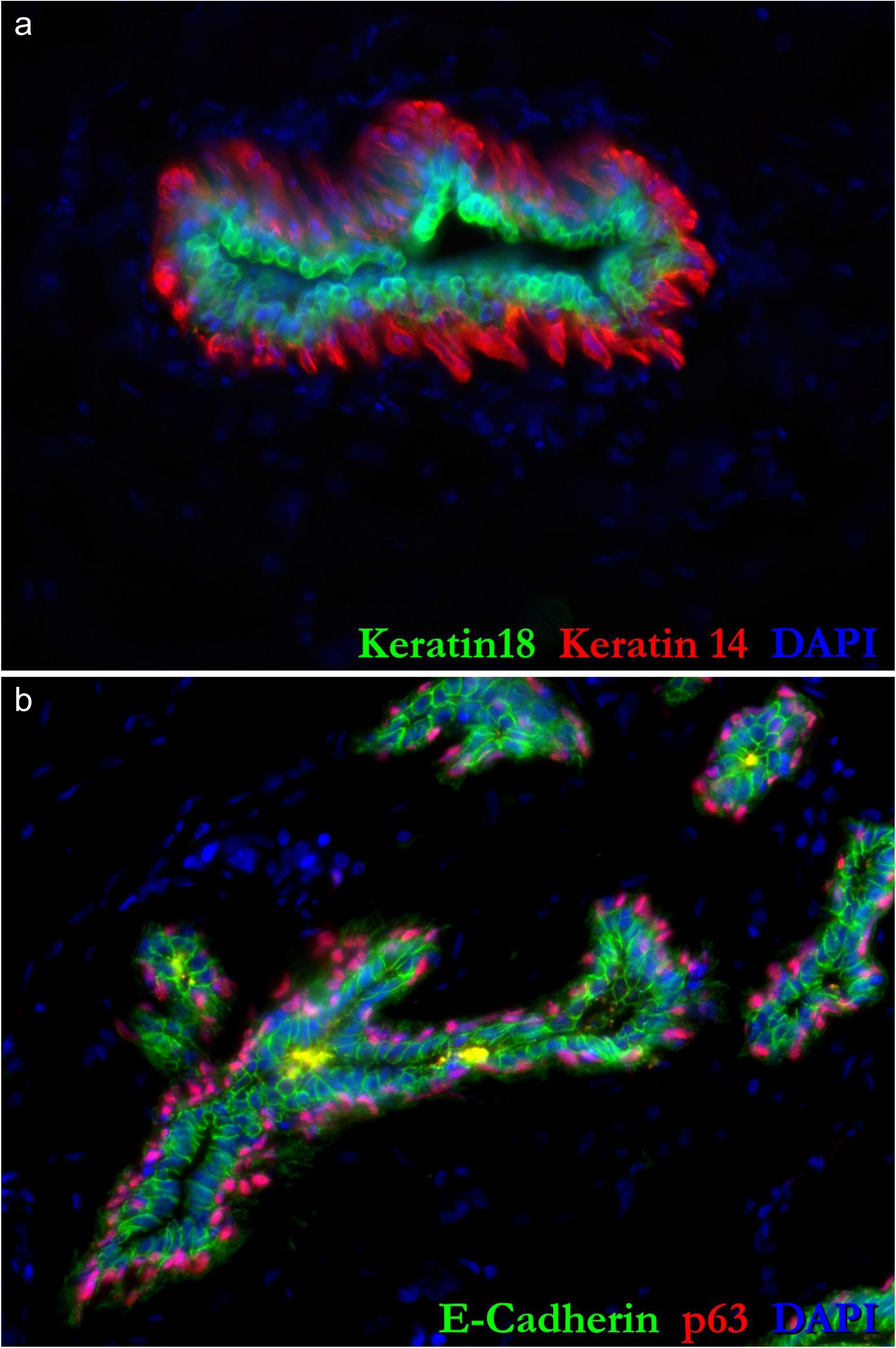
**(a,b)** High resolution images of immunostained breast tissue provided in Fig.14 a,b.

**Figure 14—figure supplement 2.**
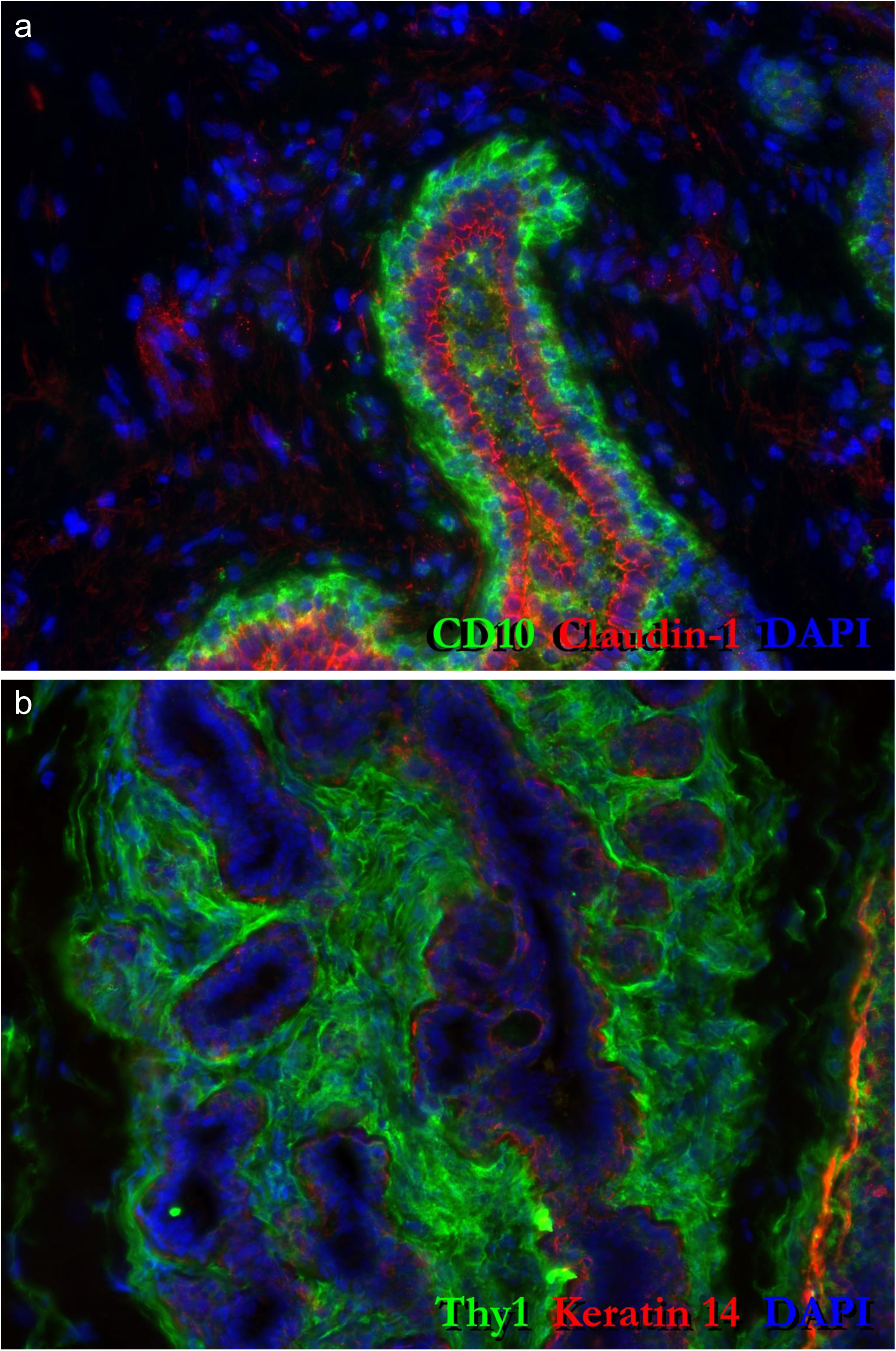
**(a,b)** High resolution images of immunostained breast tissue provided in Fig.14 i, j.

**Figure 14—figure supplement 3.**
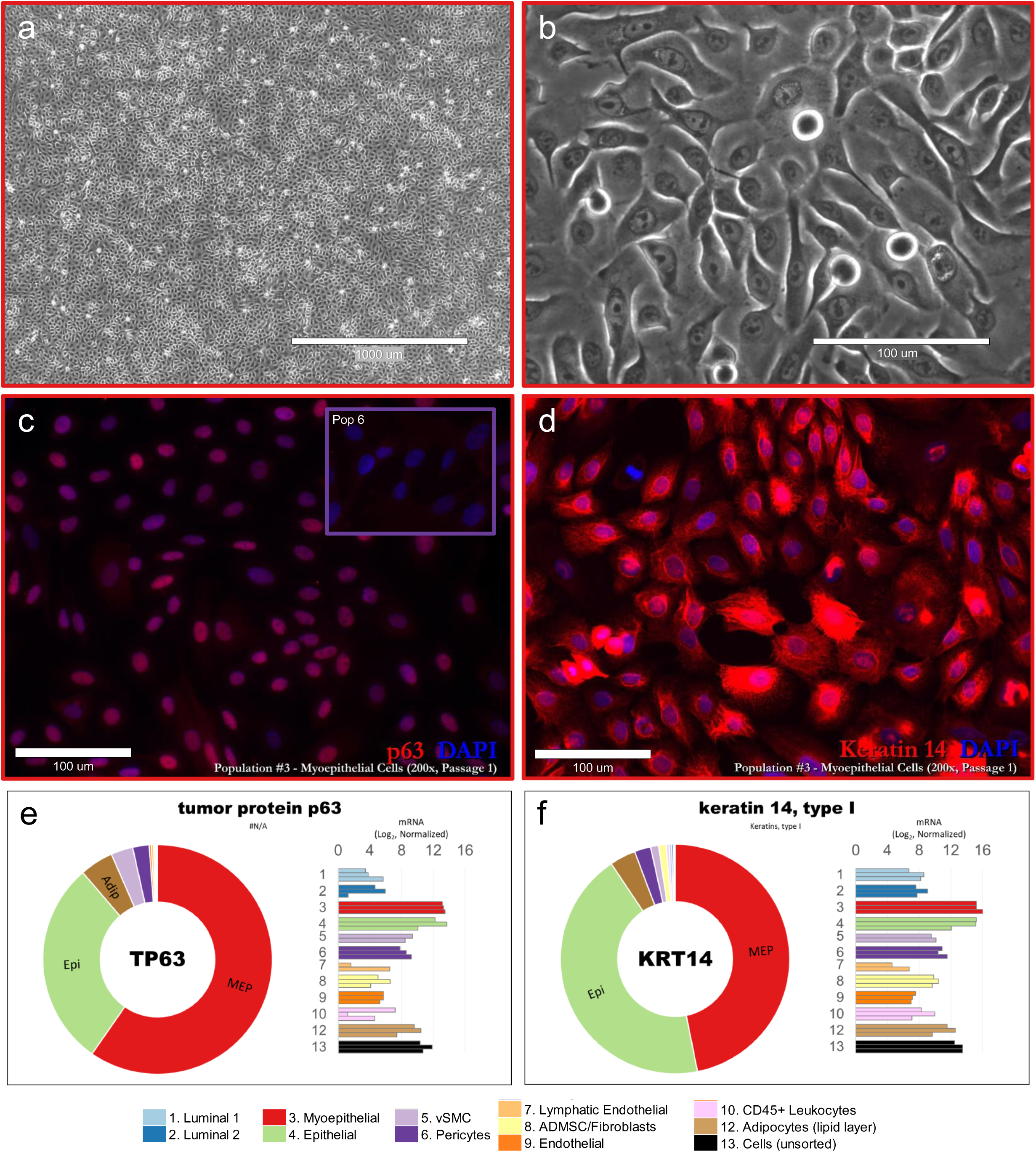
Primary Pop3 (myoepithelial) cultures imaged with **(a)** 4x and **(b)** 40x phase contrast objectives (scale = 1000μm and 100μm, respectively; imaged on EVOS imaging station; 4x Ph Plan 0.13 NA objective and 40x Plan LWD FL/PH 0. **(c,d)** Pop3 cultures immunostained for (c) p63 and (d) keratin 14; inset in c displays Thy1^Pos^ Pop6 cells immunostained for p63. The Pop3 cells were sorted from reduction mammoplasty tissues surgically excised from a 20-year-old female; sample #N238 (a,b) and a 23-year-old female; sample #N239 (c,d). **(e,f)** transcript levels of TP63 (p63) and KRT14 (keratin 14) measured in uncultured FACS-purified cell types, as determined by RNA-sequencing. Normalized mRNA values (rlog, DEseq2) are provided on log2 scale (bar graph of each biological replicate) as well as a linear scale (donut graph of median value), each color-coded by cell type (myoepithelial cells are red).

**Figure 14—figure supplement 4.**
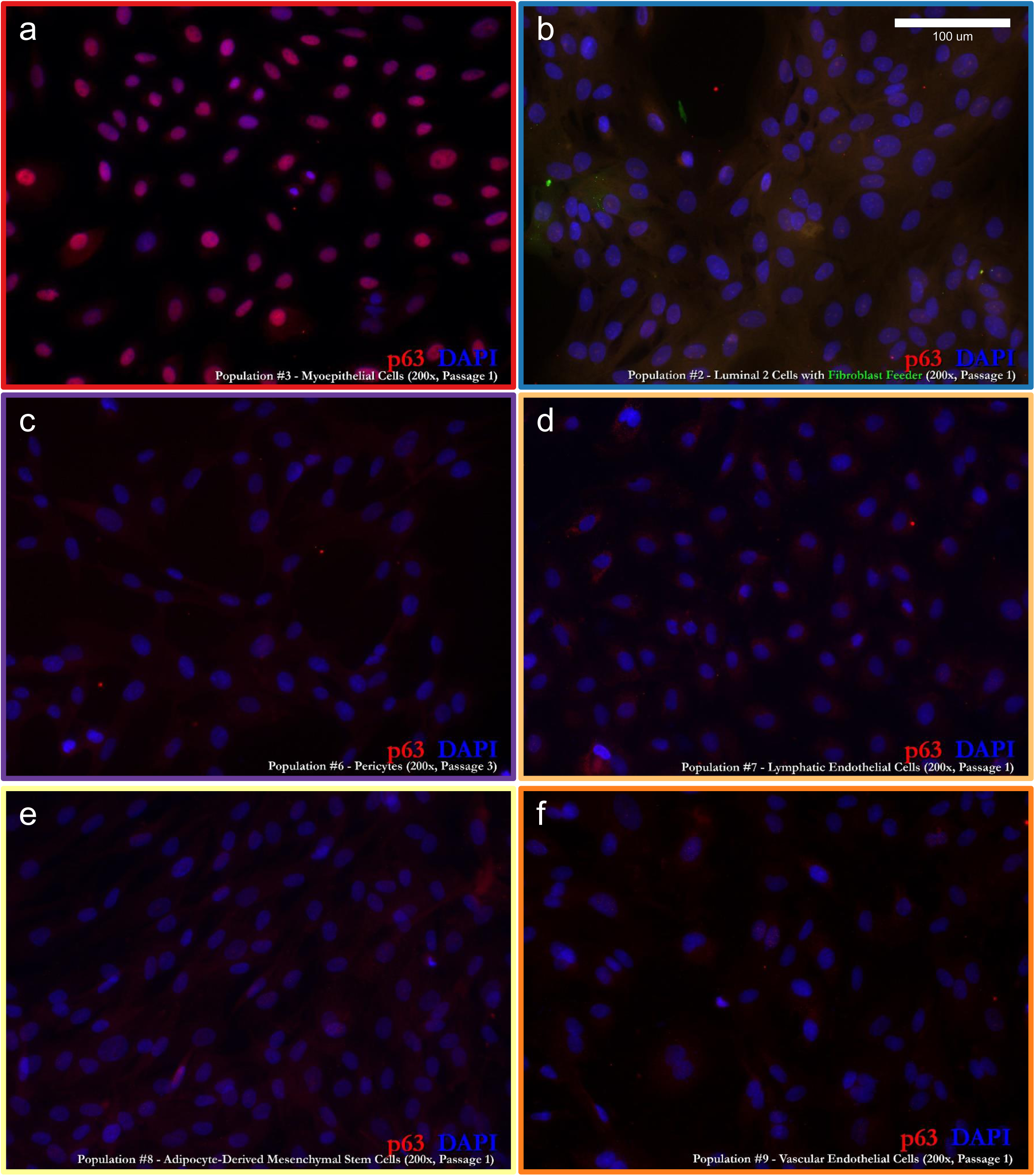
Specificity and purity of primary myoepithelial cell models. p63 immunostaining of primary cultures of (a) Pop3 myoepithelial cells, (b) luminal epithelial cells, (c) pericytes, (d), lymphatic endothelial cells, (e) fibroblasts, and (f) vascular endothelial cells. Only the Pop3 culture is stained by p63, confirming its myoepithelial identity. The absence of staining in other cultures demonstrates the rigor of each cell type’s purification strategy (as myoepithelial cells are robust and will easily overtake other cultures). Scale=100μm.

**Figure 15—figure supplement 1.**
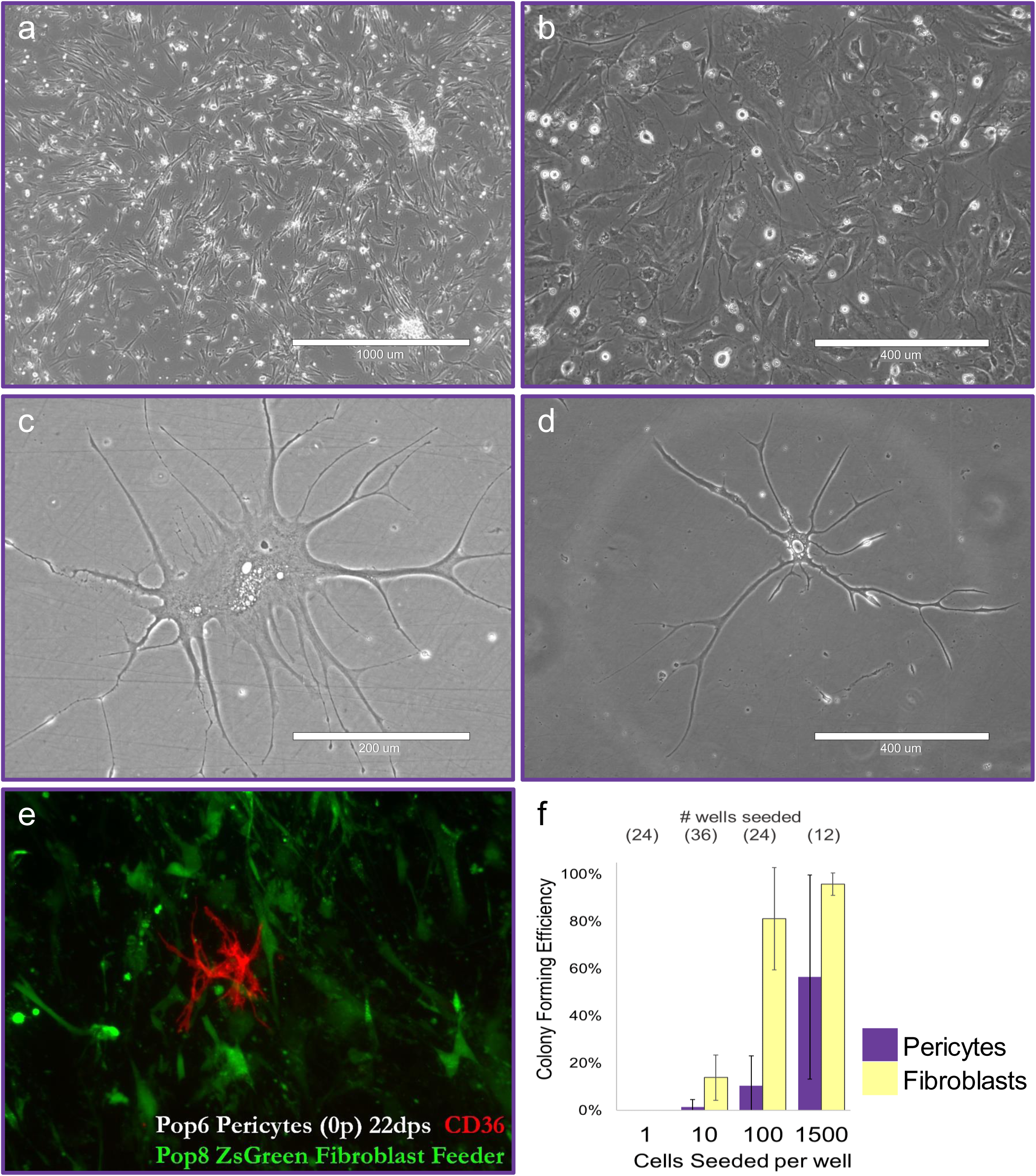
Pericyte characteristics. **(a-d)** Phase contrast images of primary Pop6 perictyes. Pericytes resembled fibroblasts, but often exhibited (a,b) a unique bi-, tri-, or multi-polar phenotype; and, at low confluence, (c,d) a prominent stellate phenotype. **(e)** CD36 immunostained co-culture of primary (passage 0) Pop6 pericytes seeded at low density into a feeder layer of ZsGreen-labeled pop8 fibroblasts, imaged 22-days post seeding (dps). (f) Colony forming efficiency of Pop6 pericytes (purple) and Pop8 fibroblasts (yellow), seeded onto a collagen I substratum at limiting densities of 1500, 100, 10, and 1 cell per well (distributed by FACS robotics). Cells were seeded into two 96-multi-well culture dishes (0.32cm^2^/well), fed weekly, and analyzed after ∼3 weeks in culture (23 days); For each cell type, n= 24 (1 cell/well), 36 (10 cells/well), 24 (100 cells/well), and 12 (1500 cells/well).

**Figure 16—figure supplement 1.**
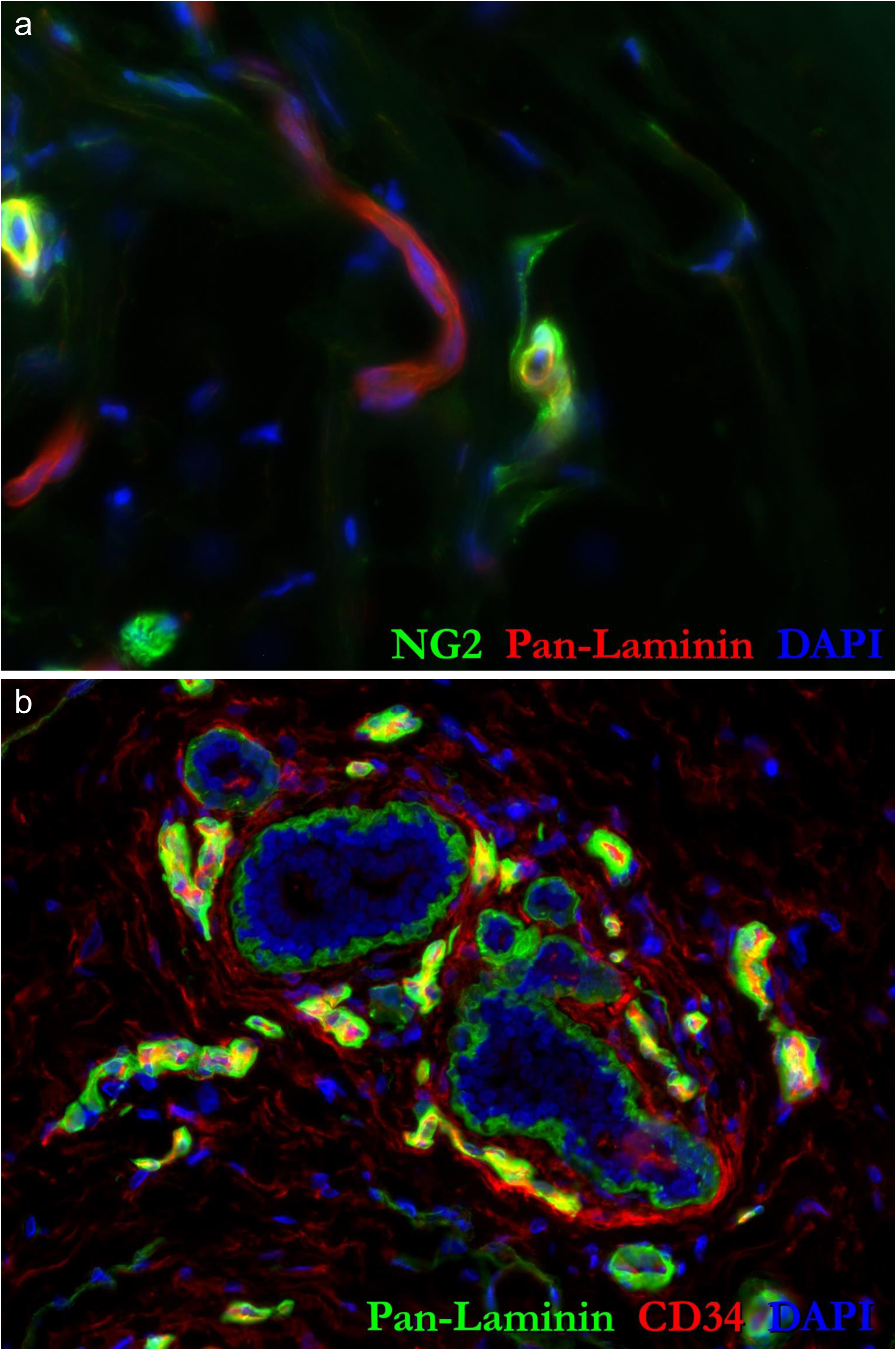
High-resolution images of pericytes. **(a)** High resolution images of the immunostained breast tissues provided in Figs.16a&b.

**Figure 16—figure supplement 2.**
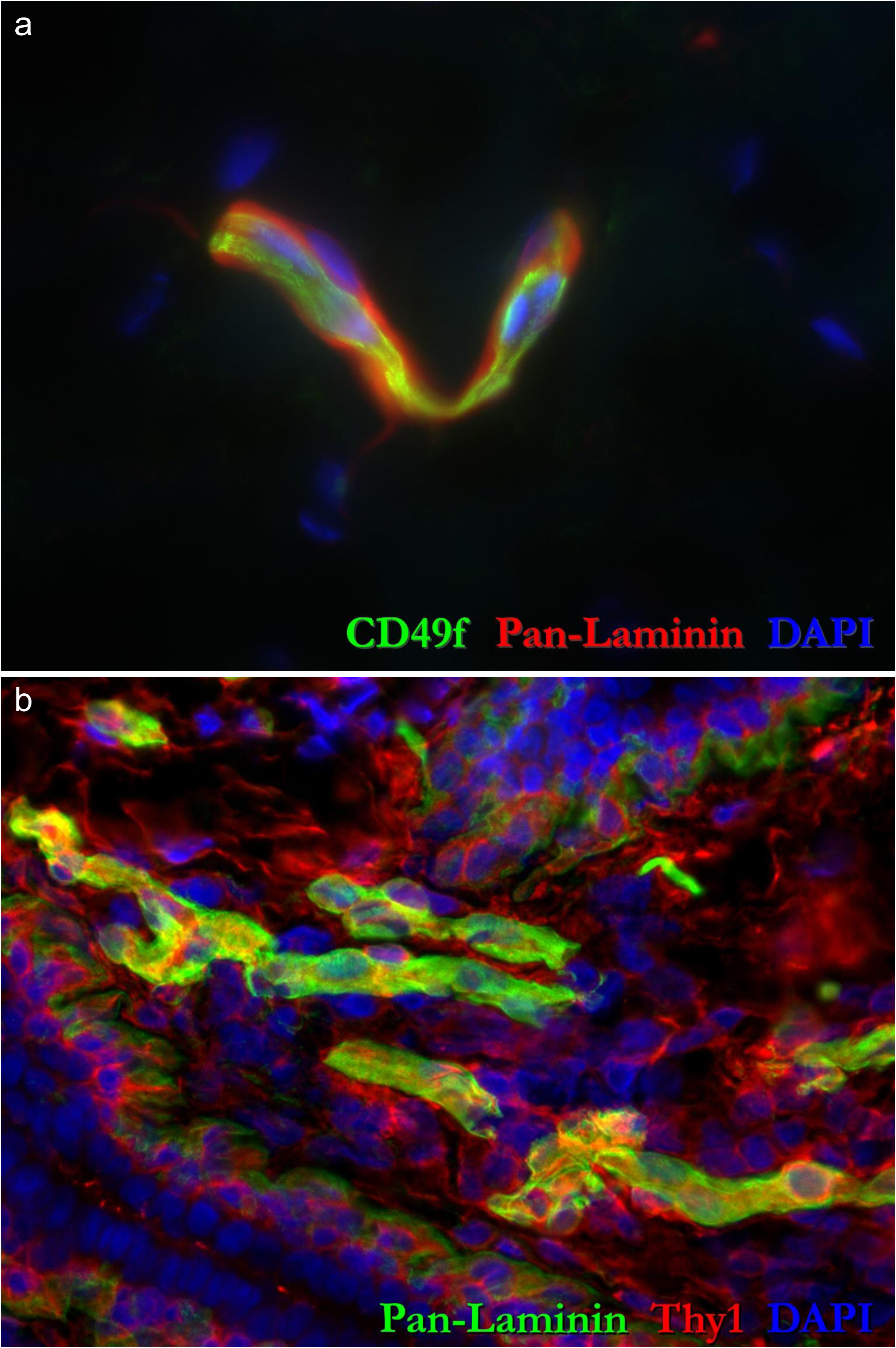
High-resolution images of pericytes. **(a)** High resolution images of the immunostained breast tissues provided in Figs.16c&d.

**Figure 16—figure supplement 3.**
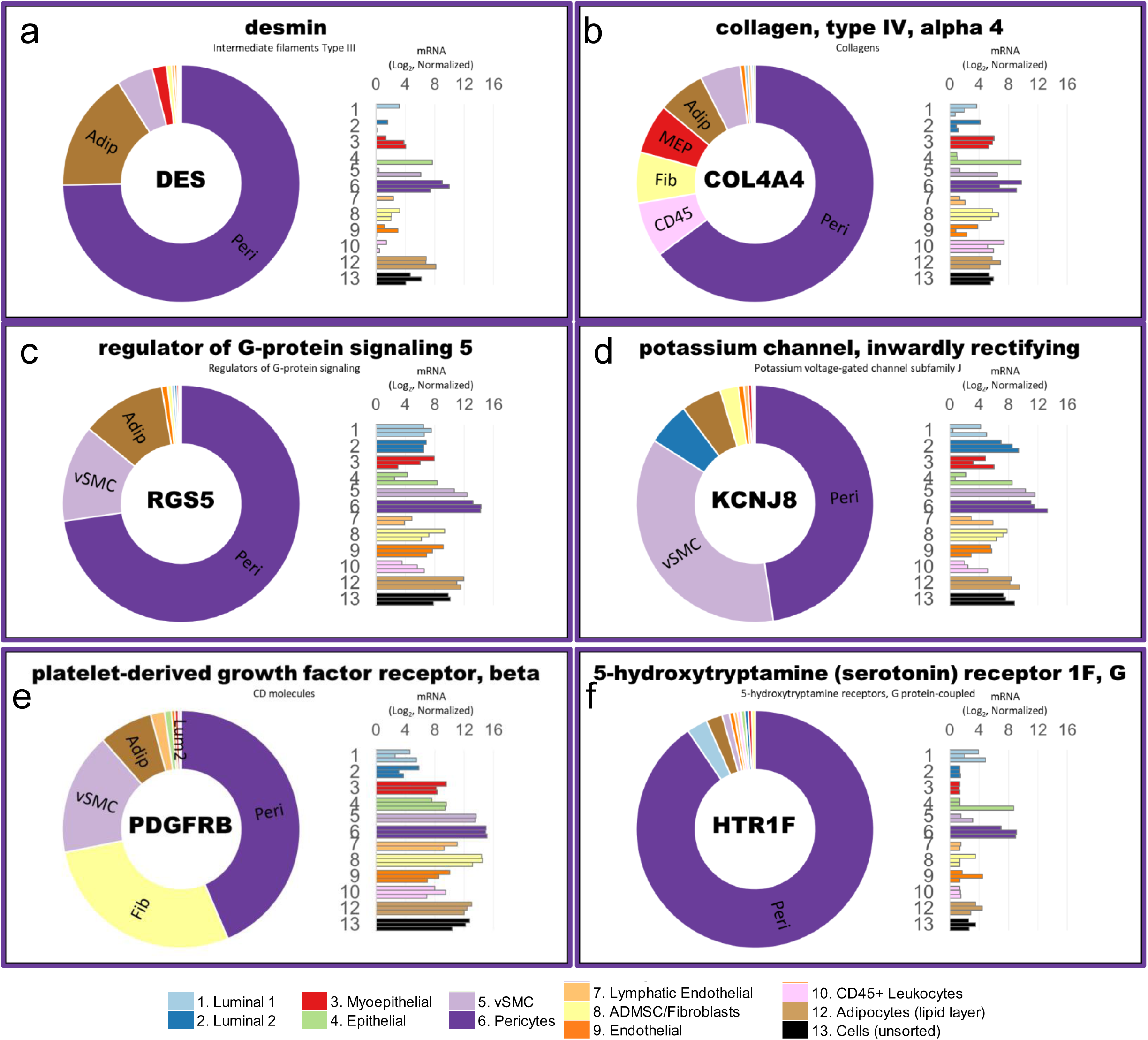
Pericyte validation via RNA-sequencing. **(a-f)** Transcript levels of pericyte-associated genes (a) desmin, (b) basement membrane collagen (Col IV alpha-4), (c) RGS5, (d) KCNJ8, and (e) PDGFRβ (aka CD140b); and (f) serotonin receptor HTR1F, a gene we discovered to be uniquely expressed by pericytes. Transcript levels were measured in freshly sorted (uncultured) cell types by RNA-sequencing. Normalized mRNA values (rlog, DEseq2) are provided on log2 scale (bar graph of each biological replicate), as well as on a linear scale (donut graph of median value) that are color-coded by cell type (pericytes are purple).

**Figure 16—figure supplement 4.**
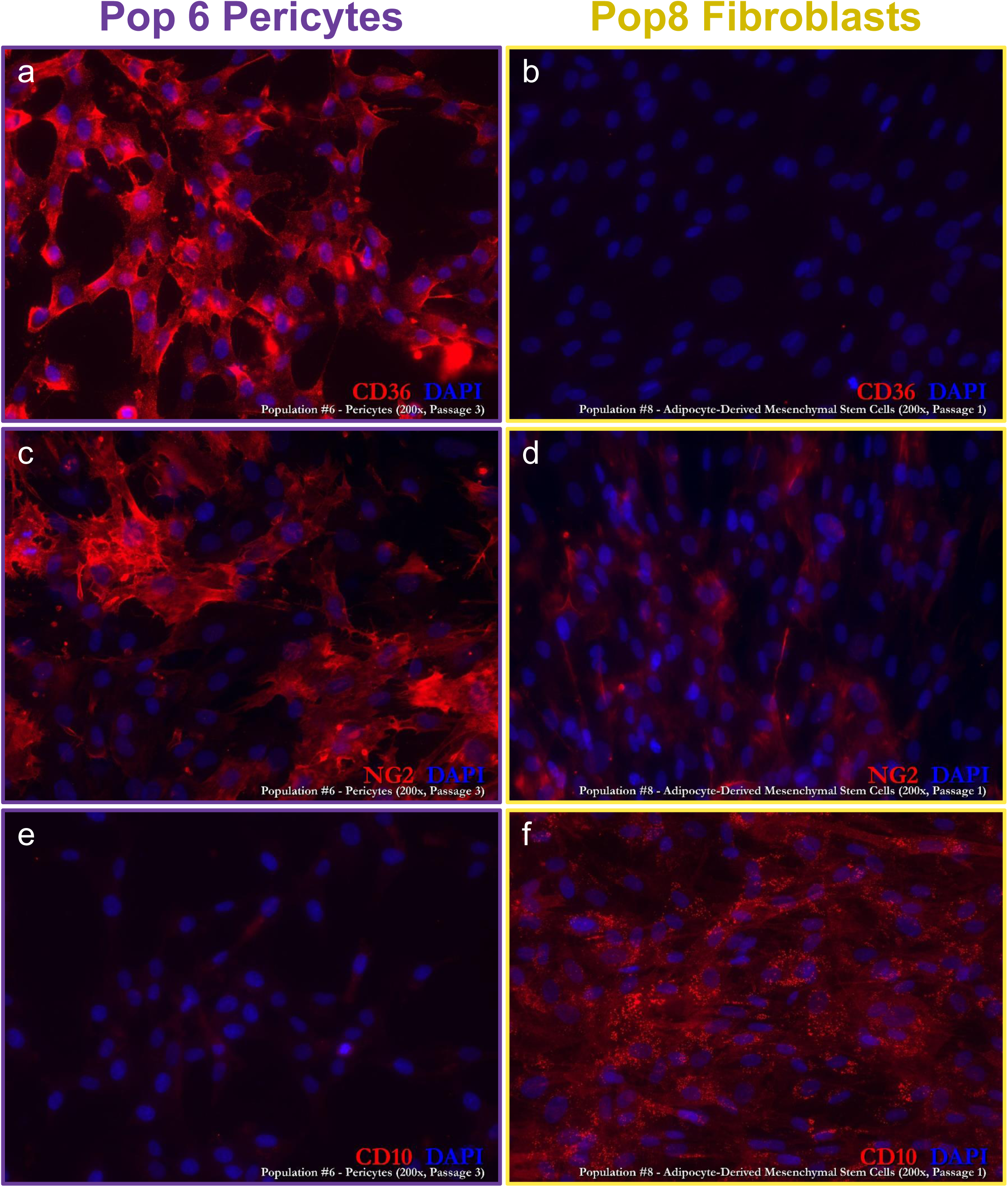
Validation of primary pericyte cultures. **(a-f)** Isogenic cultures of Pop6 pericytes (a,c,e) and Pop8 fibroblasts (b,d,f), immunostained with (a,b) CD36, (c,d) NG2, and (e,f) CD10. The observed differential staining helped establish the distinct nature of Pop6 and Pop8 cells. These cultures were both purified from sample#239, a 23 y/o female (notably, #N239 was one of four individuals whose cell populations were RNA-sequenced).

**Figure 16—figure supplement 5.**
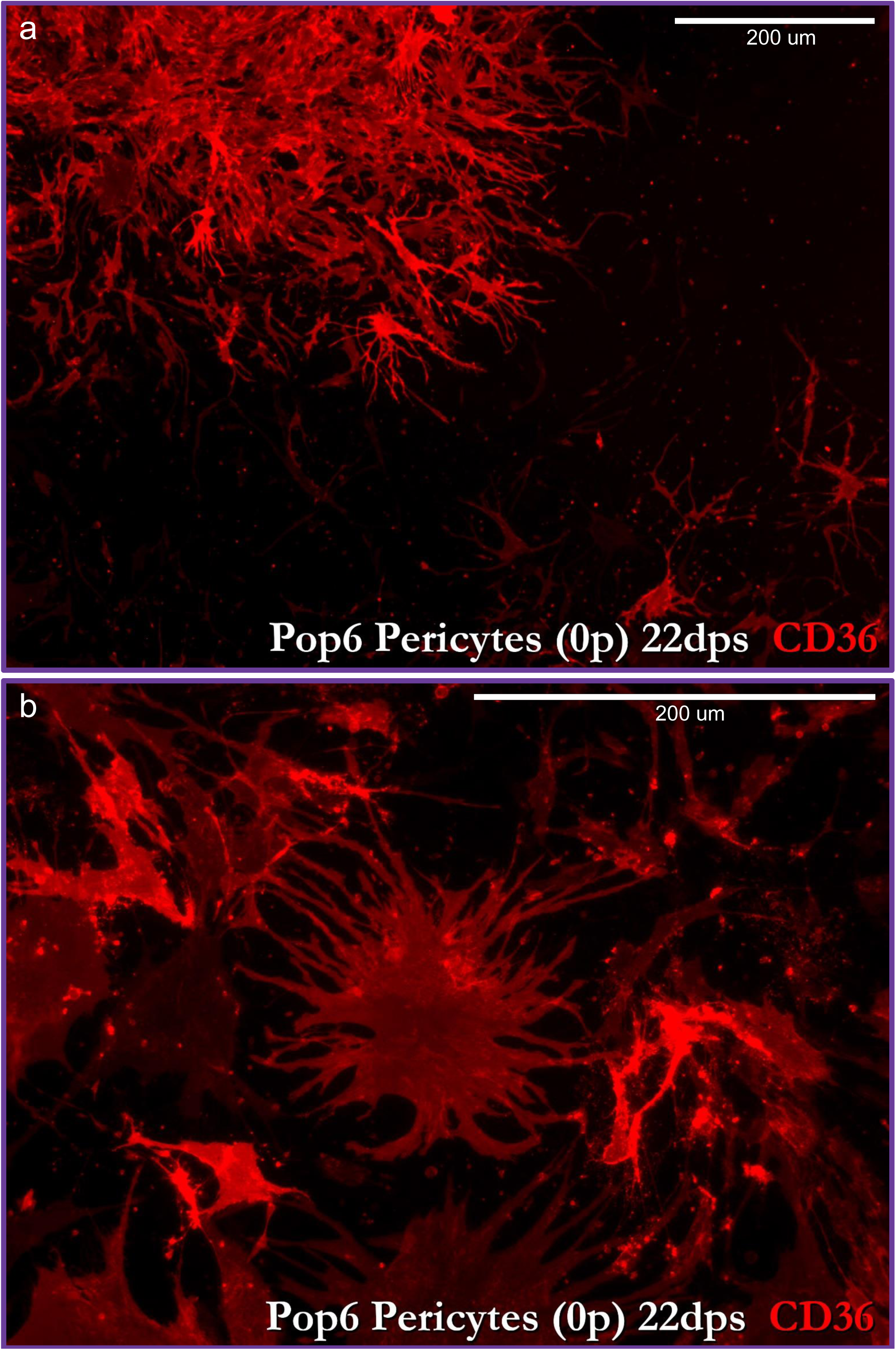
A unique morphology exhibited by pericytes. **(a,b)** High-resolution images of an early (22-days post seeding) primary culture of Pop6 (pericytes), stained with CD36. The multi-polar phenotype exhibited by pericytes with multiple cellular projections emanating from the cell body (akin to astrocytes) is unique among the breast cell populations.

**Figure 17—figure supplement 1.**
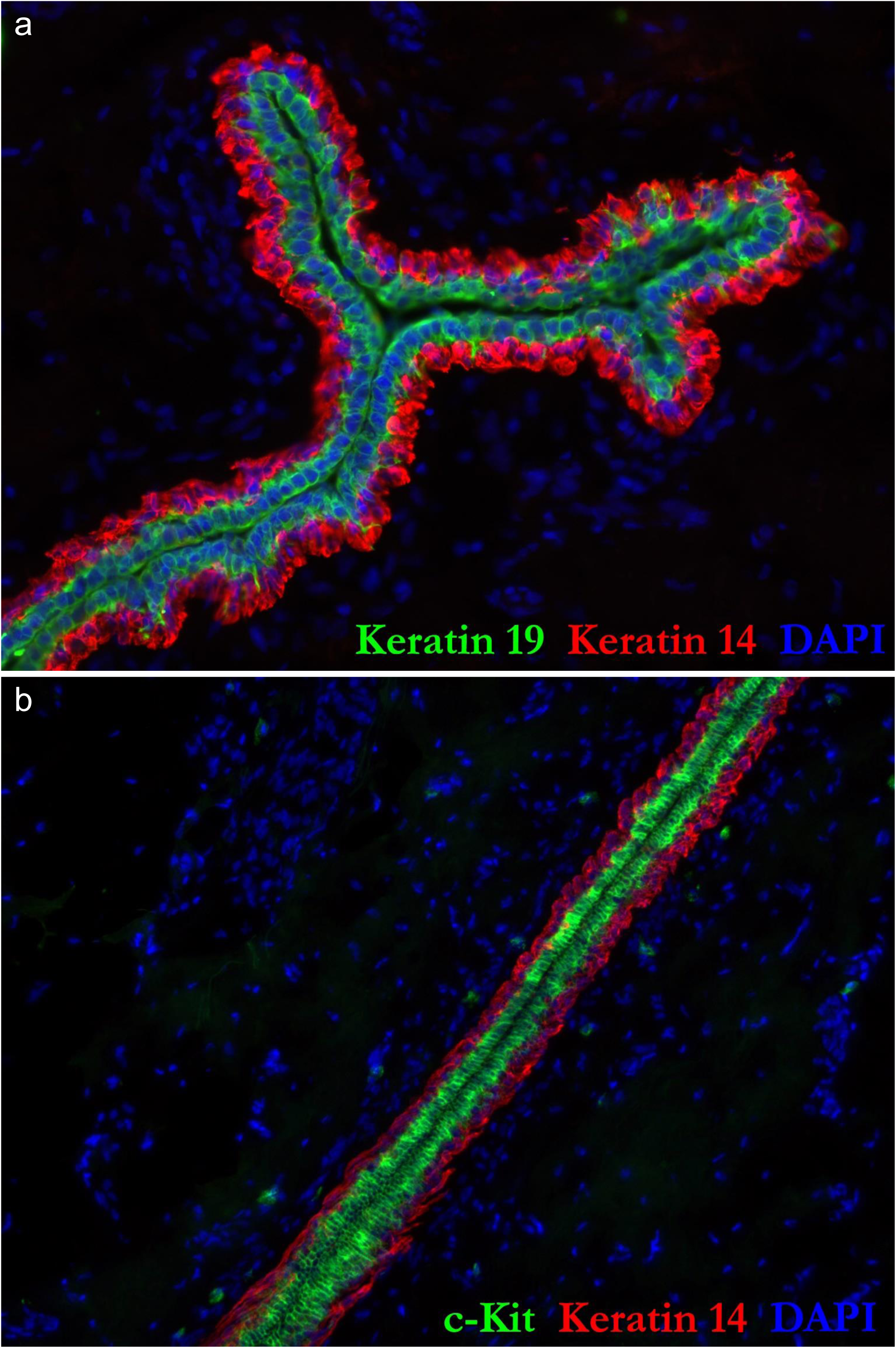
High-resolution images of luminal epithlelial cells. **(a,b)** High resolution images of the immunostained breast tissues provided in Figs.17a&b.

**Figure 17—figure supplement 2.**
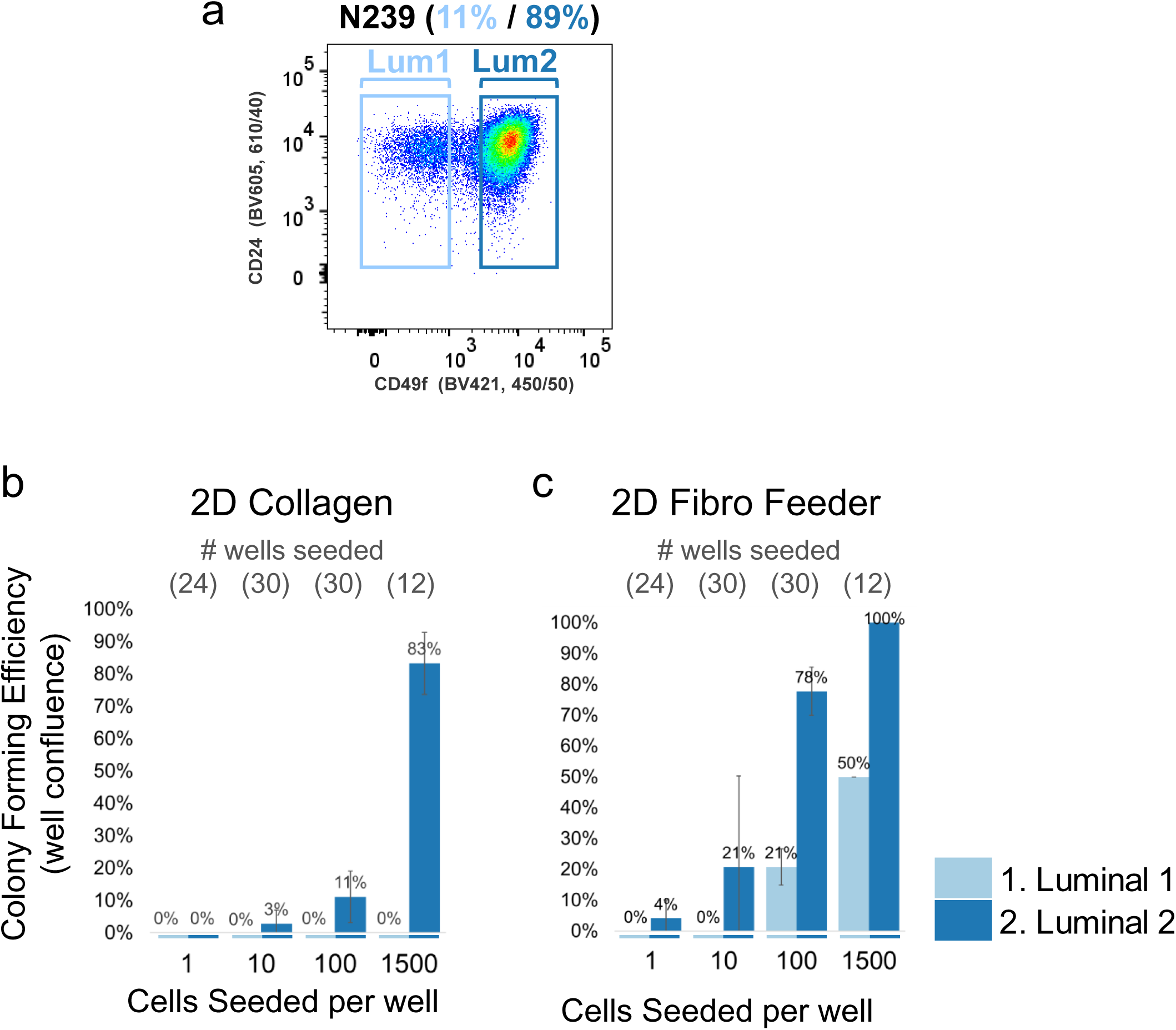
Luminal cell types possess different colony forming efficiencies that are both enhanced by stromal fibroblasts. **(a)** FACS plot of Lum1 and Lum2 cells isolated from the tissue of a 23 y/o female (sample# N239). Cells were backgated for viable single cells and CD34^Neg^ Thy1^Neg^ CD45^Neg^ CD24^Pos^ ∨ Muc1^Pos^. Of this entire luminal epithelial cell fraction, 11% were categorized as Pop1 (CD49f^Neg/Low^) and 89% were identified as Pop2 (CD49f^Pos^) luminal epithelial cells. **(b,c)** Immediately after the second round of sorting, Lum1 and Lum2 cells were sorted a final time, into (b) collagen I (Purecol®) coated 96-well dishes, where every other well contained (c) fluorescently labeled fibroblasts (N141 Pop8 fibroblasts-ZsGreen^Pos^; 1500/well), ‘2D Fibro Feeder’. The luminal cells were challenged by seeded them at limiting densities of 1500 cells/well (12 wells), 100 cells/well (30 wells), 10 cells/well (30 wells), and 1 cell/well (24 wells), comprising a total of 192 wells (one 96-well dish for each luminal cell type). After 35 days in culture, the wells were scored for cell growth (i.e, presence of large colonies that had reached 80-100% well confluency). Lum1 and Lum2 cells displayed differential colony forming efficiencies that were enhanced by stromal fibroblasts; and in the case of Pop1, were necessary for them to grow at these seeding densities.

**Figure 18—figure supplement 1.**
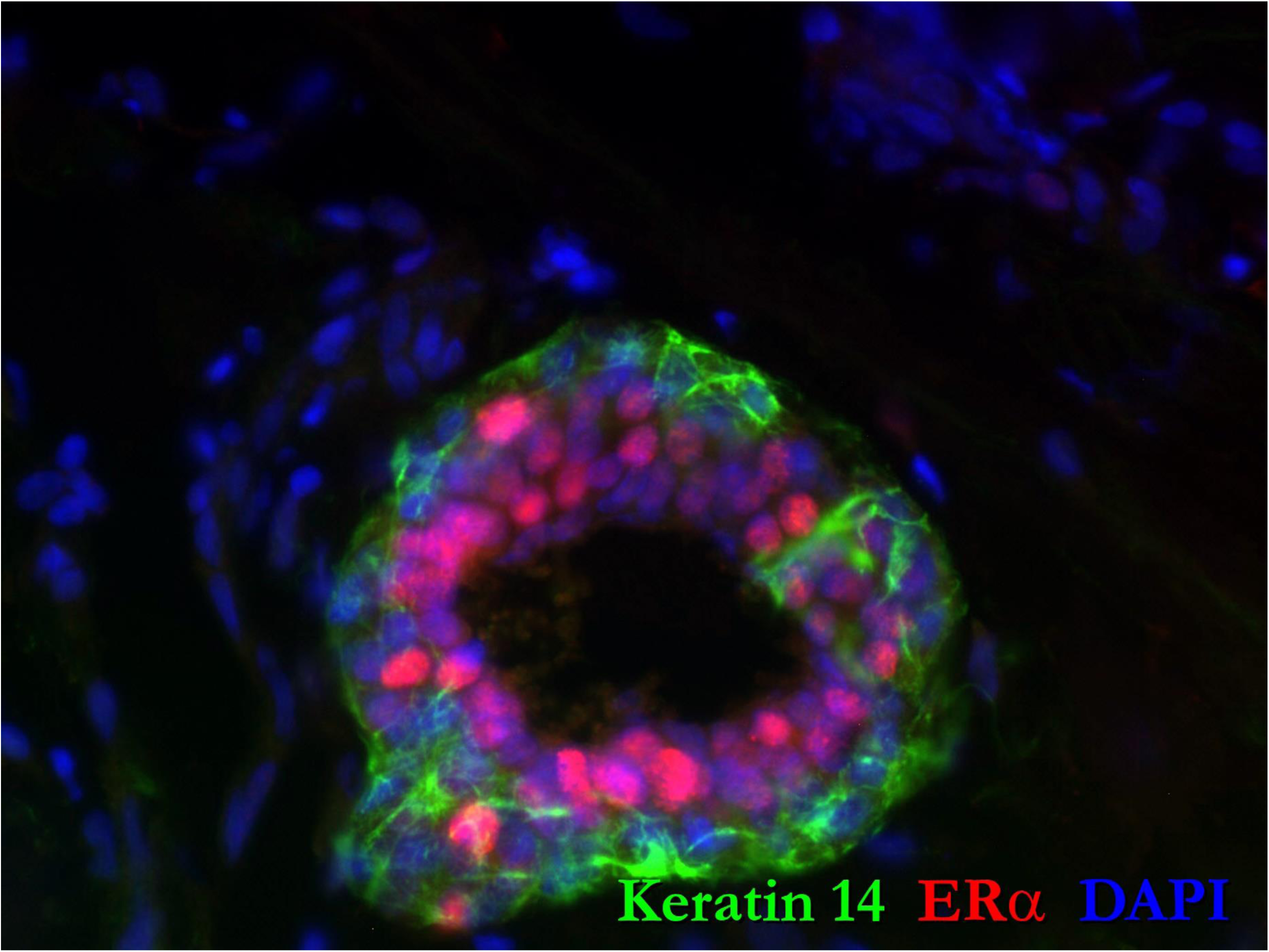
High resolution image of the immunostained breast tissue provided in Fig 18c. Sample N277, reduction mammoplasty tissue from a 22 y/o female, stained for keratin 14 (green) and estrogen receptor alpha (red). The luminal cell compartment in this tissue is dominated by ER^Pos^ luminal cells. Immunostaining results matched the high proportion of Pop1 luminal cells observed by FACS.

**Figure 19 – video supplement 1.**
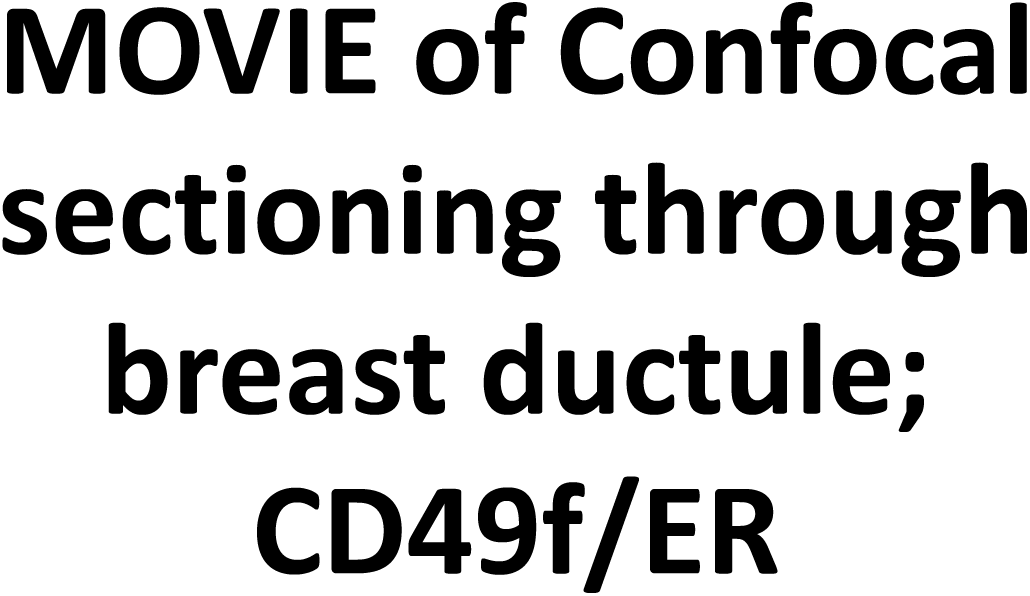
AVI movie of Z-stacked confocal images illustrated in Fig.19. The movie shows a breast ductule, stained with CD49f (green), ERα (red), keratin 14 (blue), and DAPI (white). Mutually exclusive expression of ERα and CD49f was found among the luminal cells.

**Figure 19 – figure supplement 1.**
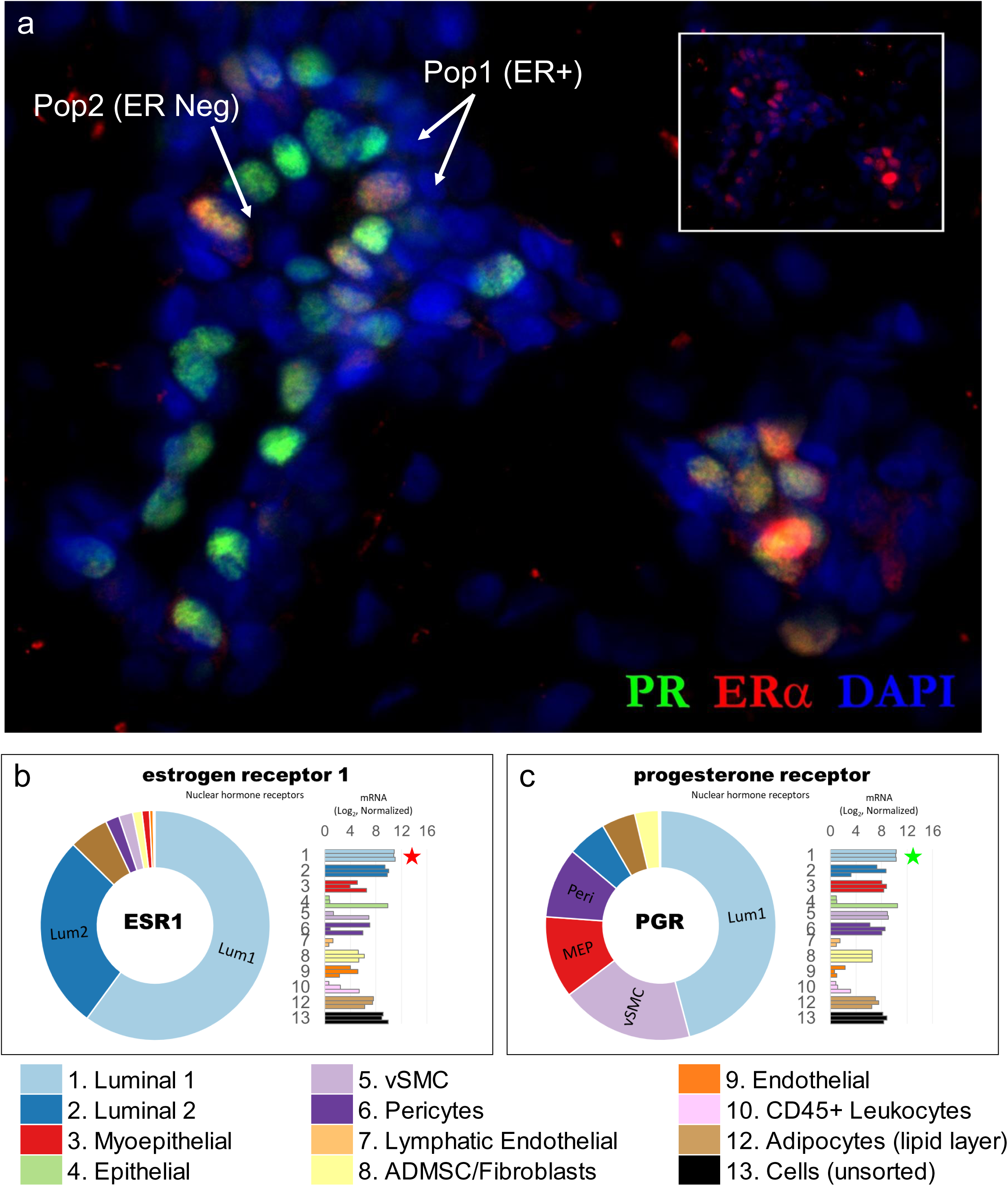
co-incident expression of estrogen and progesterone receptors within Pop1 luminal epithelial cells. **(a)** Immunostaining of normal breast tissue reveals coincident expression of progesterone receptor (PR, green) and estrogen receptor alpha (ERα, red), which are both localized to a subfraction of luminal epithelial cells. Digital removal of green ERα signal exposes the dimmer PR staining within each ERα^Pos^ cell (inset). **(b&c)** Tissue immunostaining is consistent with the higher mRNA levels of the genes encoding (b) ERα (ESR1) and (c) PR (PGR) within Pop1 luminal epithelial cells (light blue bars, marked with red and green stars, respectively). Normalized mRNA values (rlog, DEseq2) are provided on log2 scale (bar graph of each biological replicate) and a linear scale (donut graph of median value), which are both color-coded by cell type.

**Figure 19 – figure supplement 2.**
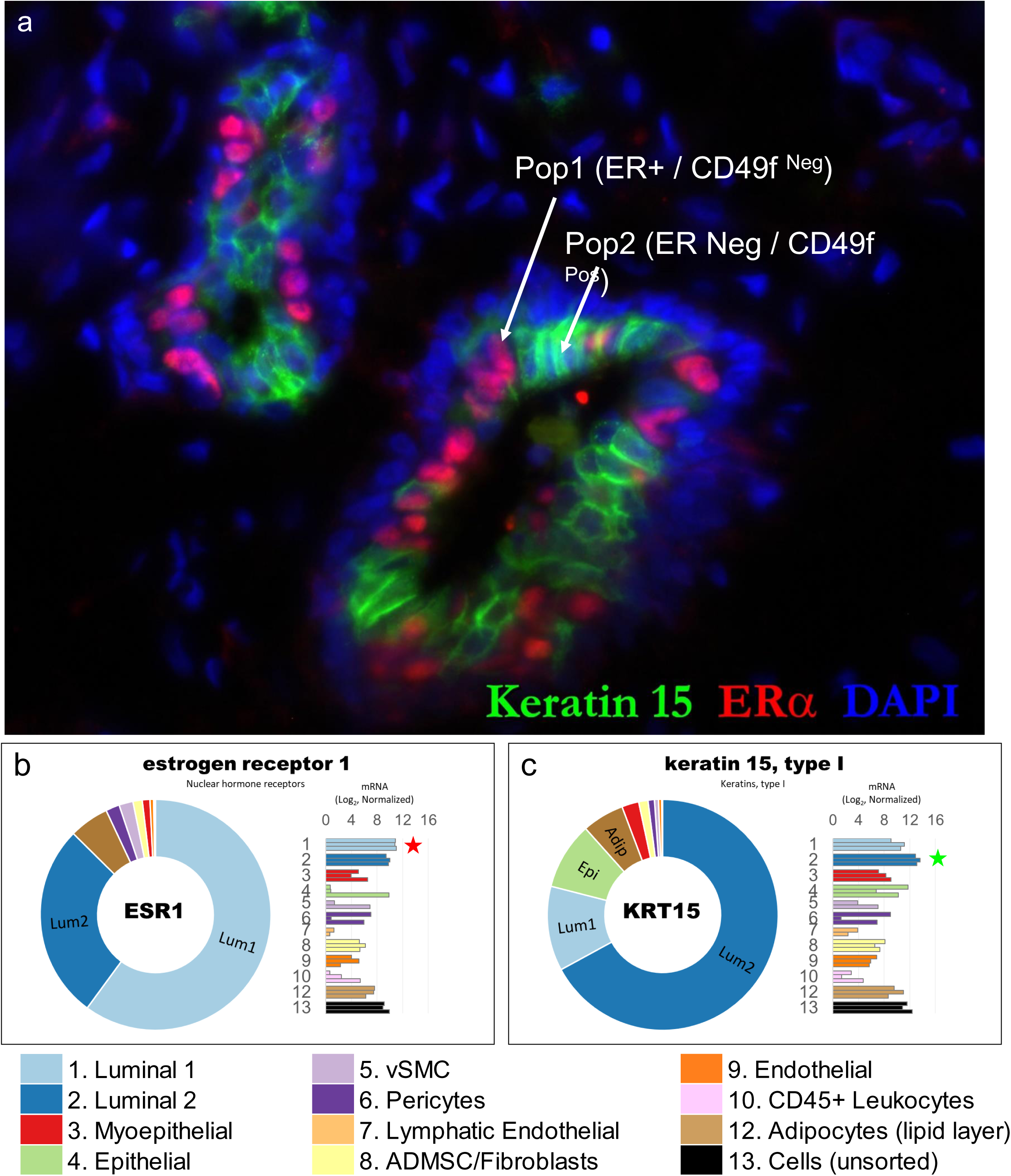
Mutually exclusive expression of estrogen receptor and keratin 15 within luminal epithelial cells. **(a)** Immunostaining of normal breast tissue with ERα and keratin 15 (K15). **(b&c)** RNA sequencing indicates (b) ERα is differentially expressed by Pop1 luminal cells, whereas (c) K15 was found differentially expressed within Pop2 luminal cells. The mutually exclusive expression of these two proteins between the two luminal cell types was confirmed by immunostaining.

**Figure 19 – figure supplement 3.**
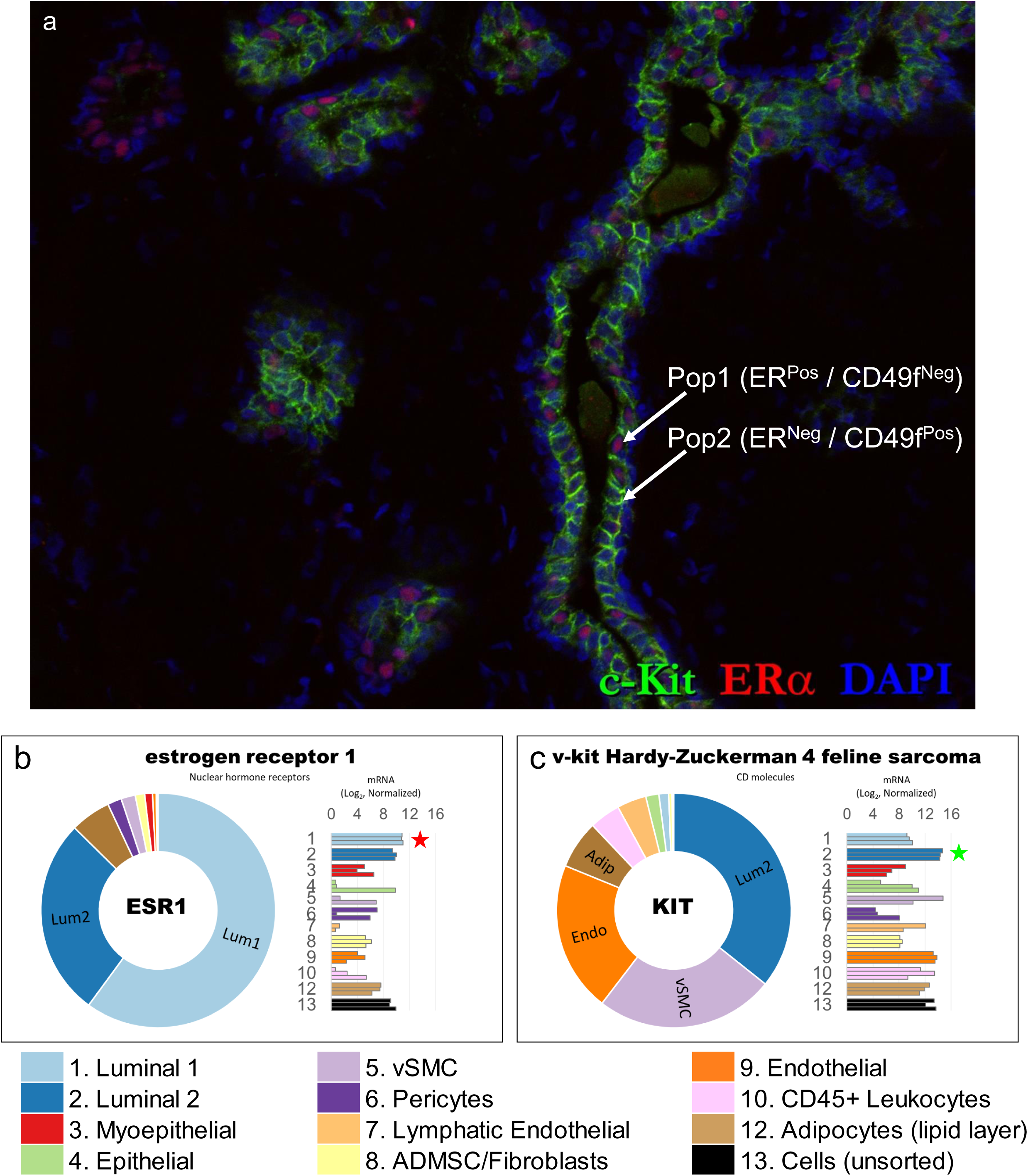
Mutually exclusive expression of estrogen receptor and c-Kit within luminal epithelial cells. **(a)** Immunostaining of normal breast tissue with ERα and c-Kit **(b&c)** RNA sequencing indicated (b) ERα (ESR1) was differentially expressed by Pop1 luminal cells, whereas (c) c-kit (KIT) was found differentially expressed within Pop2 luminal cells. The mutually exclusive expression of these two proteins between luminal cell types was confirmed by immunostaining.

**Figure 20— video supplement 1.**
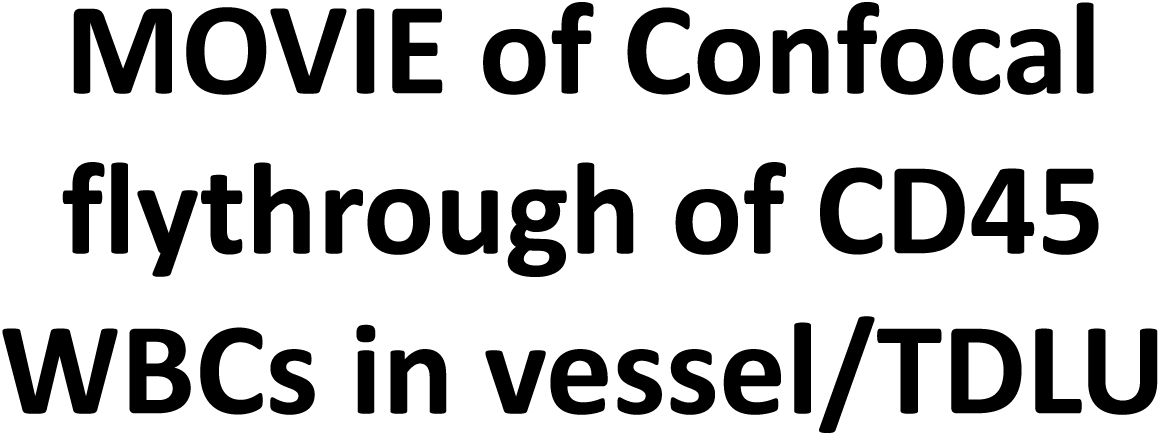
Intravascular and extravascular leukocytes within the breast. AVI movie zooming and rotating through the 3D projection of confocal images illustrated in Fig.20c. CD45 and pan-laminin immunostaining of normal breast tissue reveals a breast lobule firmly enveloped by blood vessels. CD45-stained intravascular leukocytes are present within the large vessel in the foreground, and extravascular leukocytes are present amid the epithelial cells within the breast lobule (on right). These data are from a normal reduction mammoplasty breast tissue specimen resected from a 33-year-old female (sample #N209).

**Figure 20 – figure supplement 1.**
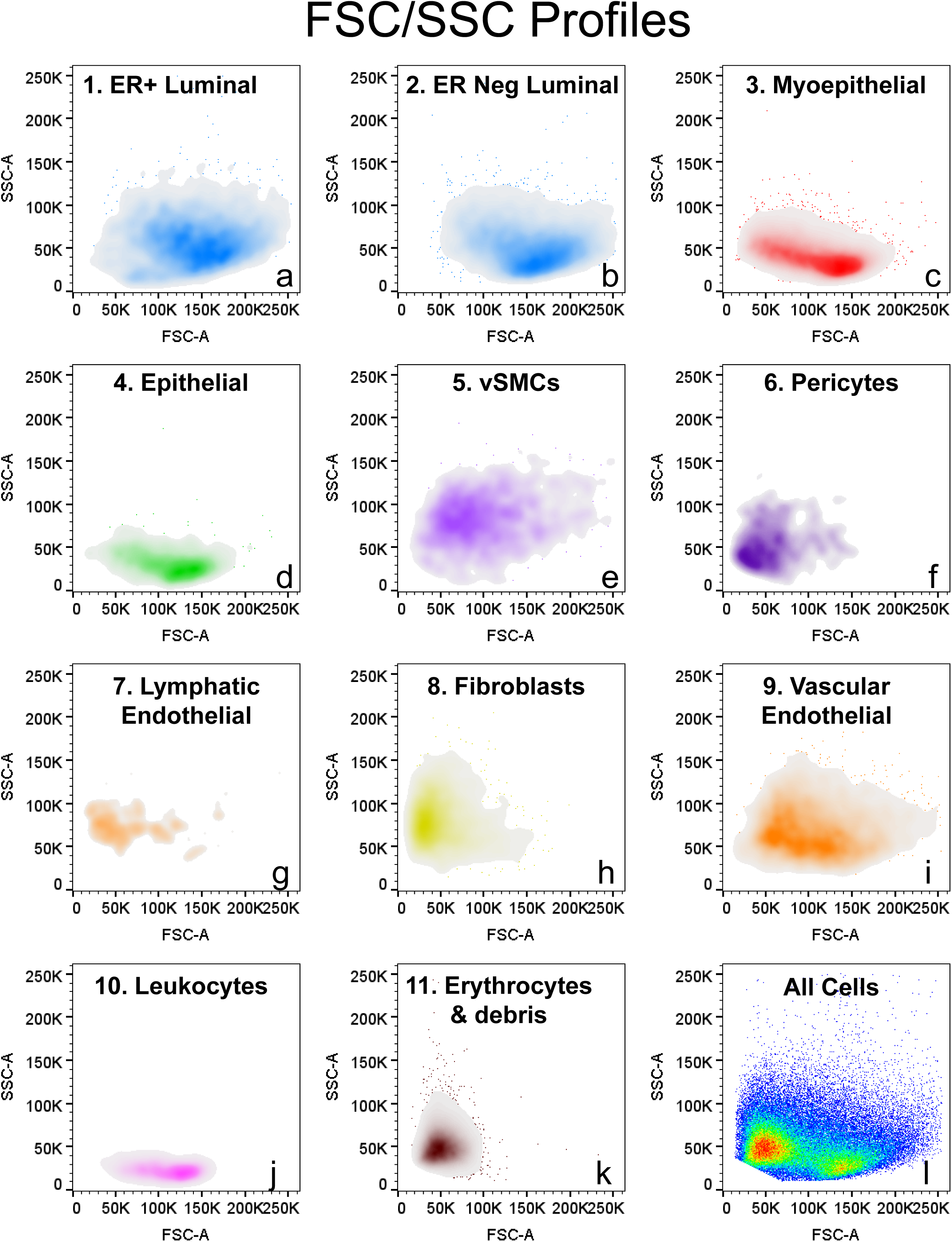
Physical characteristics the breast cell populations. Plots of forward and side scatter reveal the physical characteristics of each of each gated cell population/cell type (forward scatter (FSC) is correlated with cell size; Side scatter (SSC) is associated with cell shape and internal vesicles/granularity).

**Figure 20—figure supplement 2.**
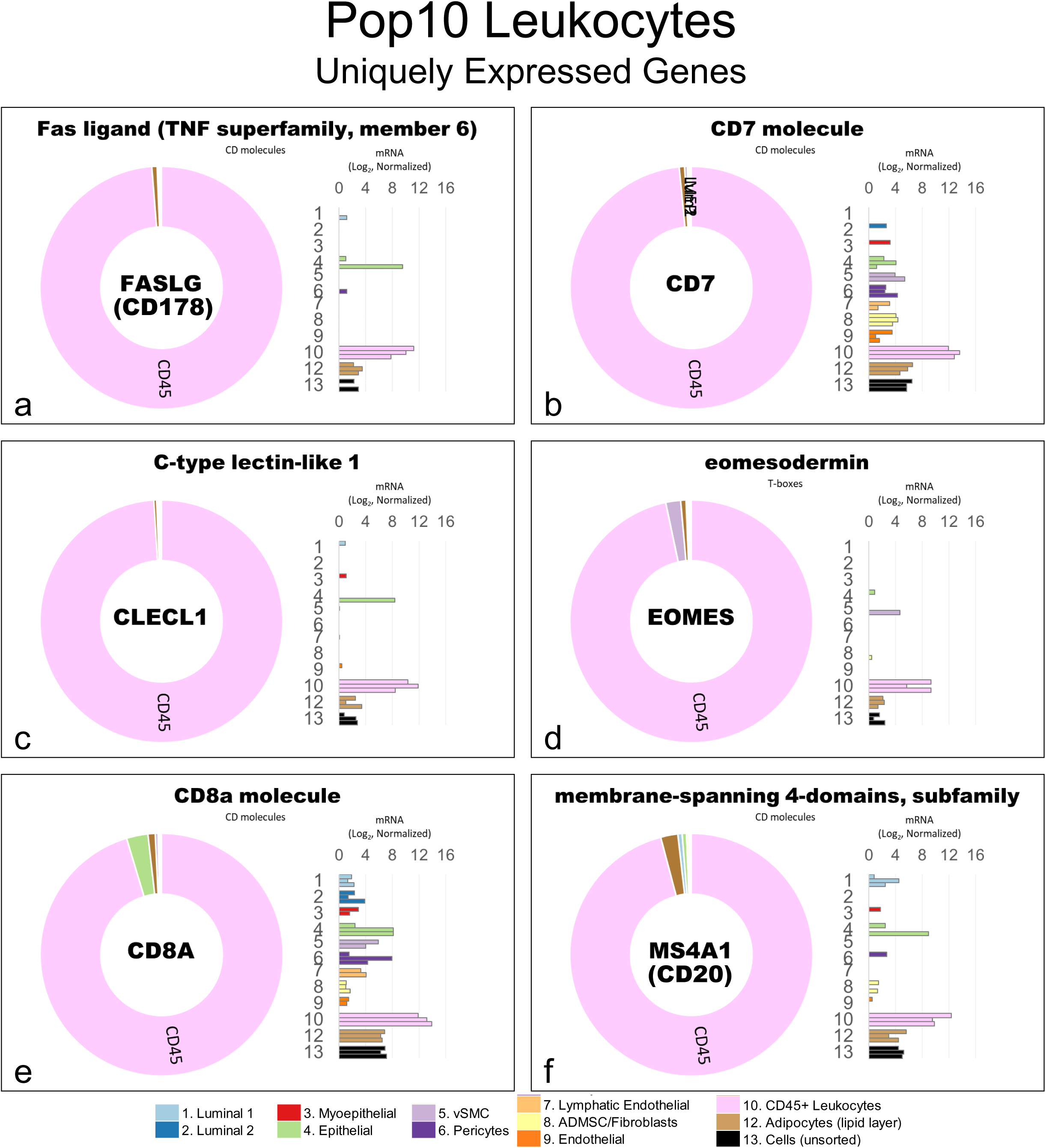
Leukocyte validation via RNA-sequencing. **(a-f)** Transcript levels of genes uniquely-expressed by Pop10 leukocyes included: (a) Fas-ligand (CD178), (b) CD7, (c) C-type lectin-like 1 (CLECL1), (d) eosmesodermin (EOMES), (e) CD8a, and (f) B-lymphocyte antigen, membrane-spanning 4A1 (MS4A1). Transcript levels were measured in freshly sorted (uncultured) cell types by RNA-sequencing. Normalized mRNA values (rlog, DEseq2) are provided on log2 scale (bar graph of each biological replicate), as well as on a linear scale (donut graph of median value) that are color-coded by cell type (leukocytes are pink).

**Figure 20—figure supplement 3.**
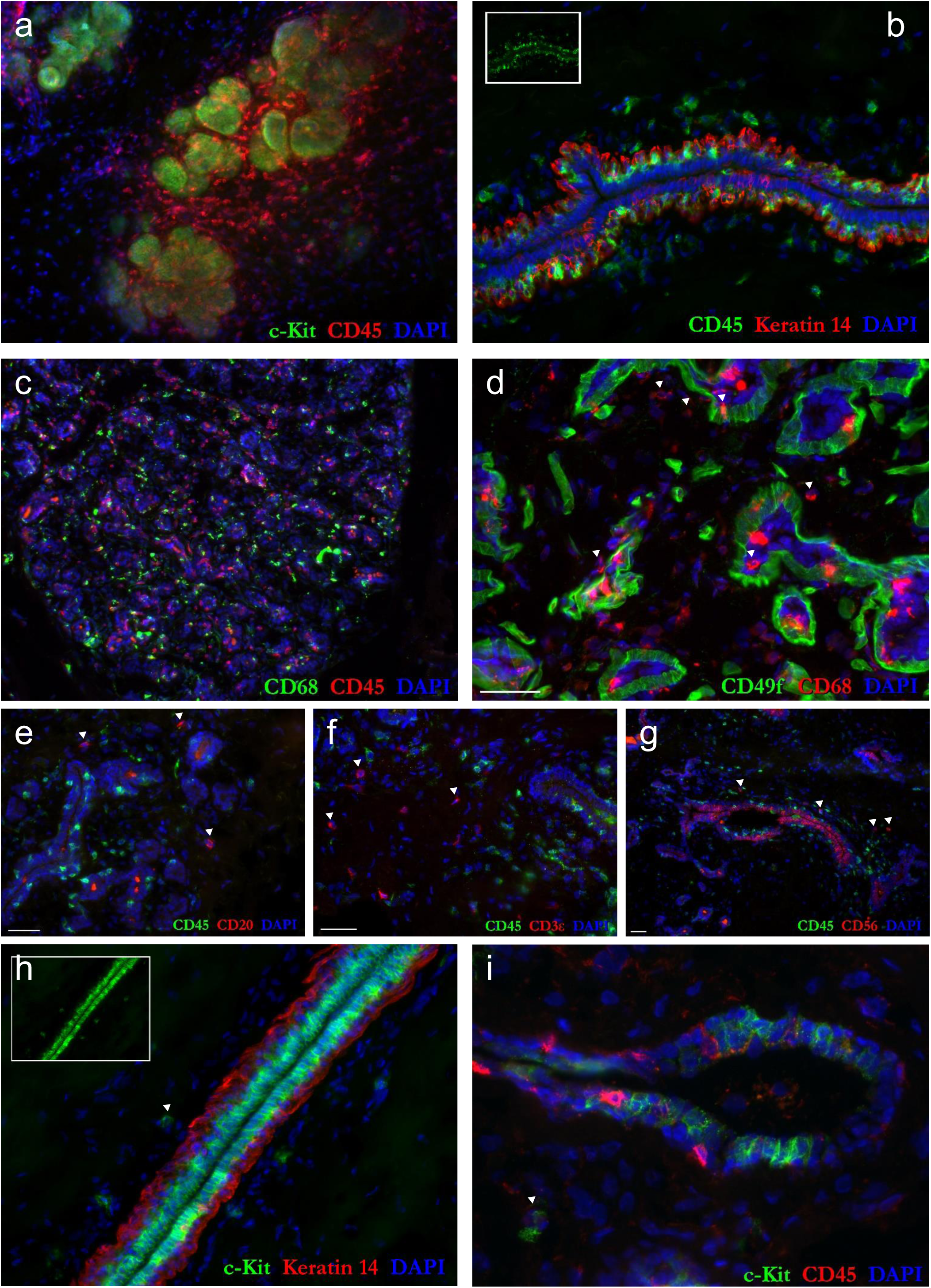
Histological location of leukocyte subtypes. **(a-i)** Normal breast tissue stained with the pan-leukocyte marker CD45 and (a,h,i) c-kit (which labels mast and luminal epithelial cells), (b) Keratin 14 (labels myoepithelial cells and reveals epithelium), (c) CD68 (labels macrophages), (e) CD20 (B-cells), (f) CD3ε (T-cells), and (g) CD56 (which natural killer (NK) cells—and surprisingly also luminal epithelial cells). **(a)** a strikingly abundant number of leukocytes were often found in the breast, almost always concentrated near (a) TDLUs and (b) ductal epithelium. The vast majority of leukocytes were found in the stroma proper, but many leukocytes had transmigrated past the basement membrane, where they existed alongside myoepithelial and luminal epithelial cells. **(c,d)** Most breast leukocytes expressed the macrophage marker CD68 (marked by ‘▾’s in d). Other leukocyte types observed were **(e)** CD20^Pos^ B-cells, **(f)** CD3ε^Pos^ T-cells, **(g)** CD56^Pos^ NK cells, and **(h,i)** mast cells, each marked by ‘▾’ in the corresponding images.

**Figure 20—figure supplements 4-5.**
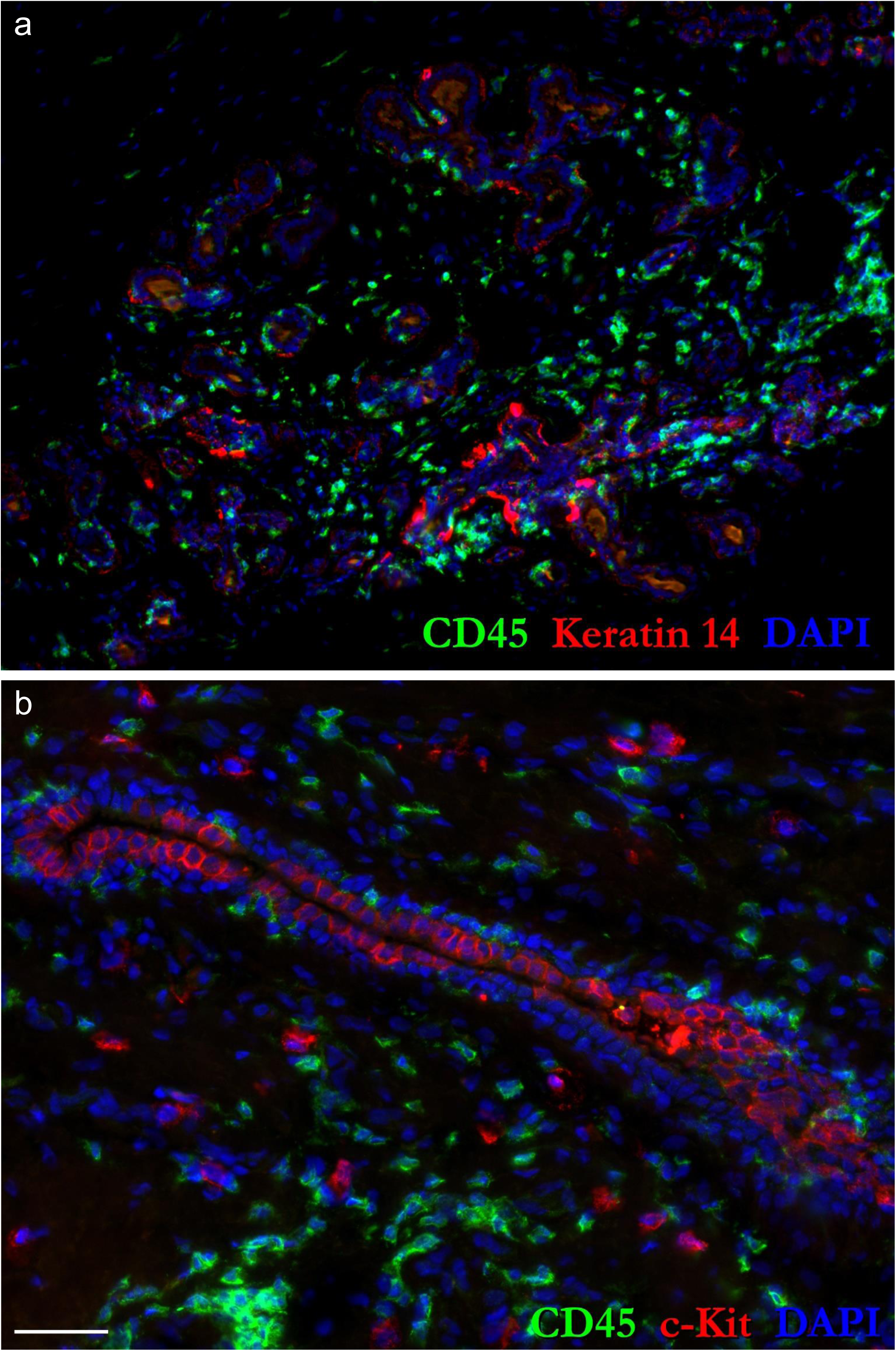
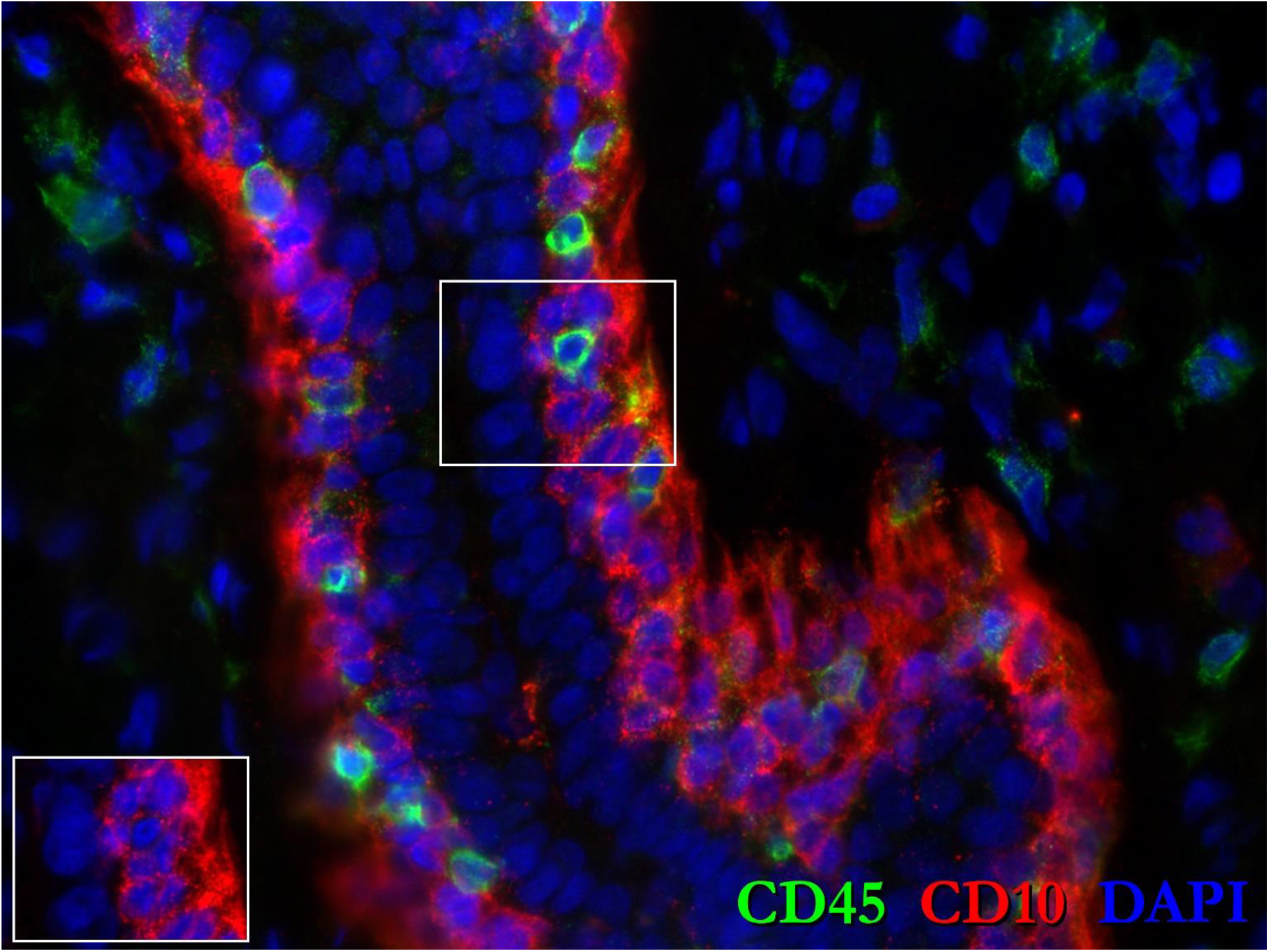
High resolution images of Figs. 20d-f

**Figure 20—figure supplements 6-10.**
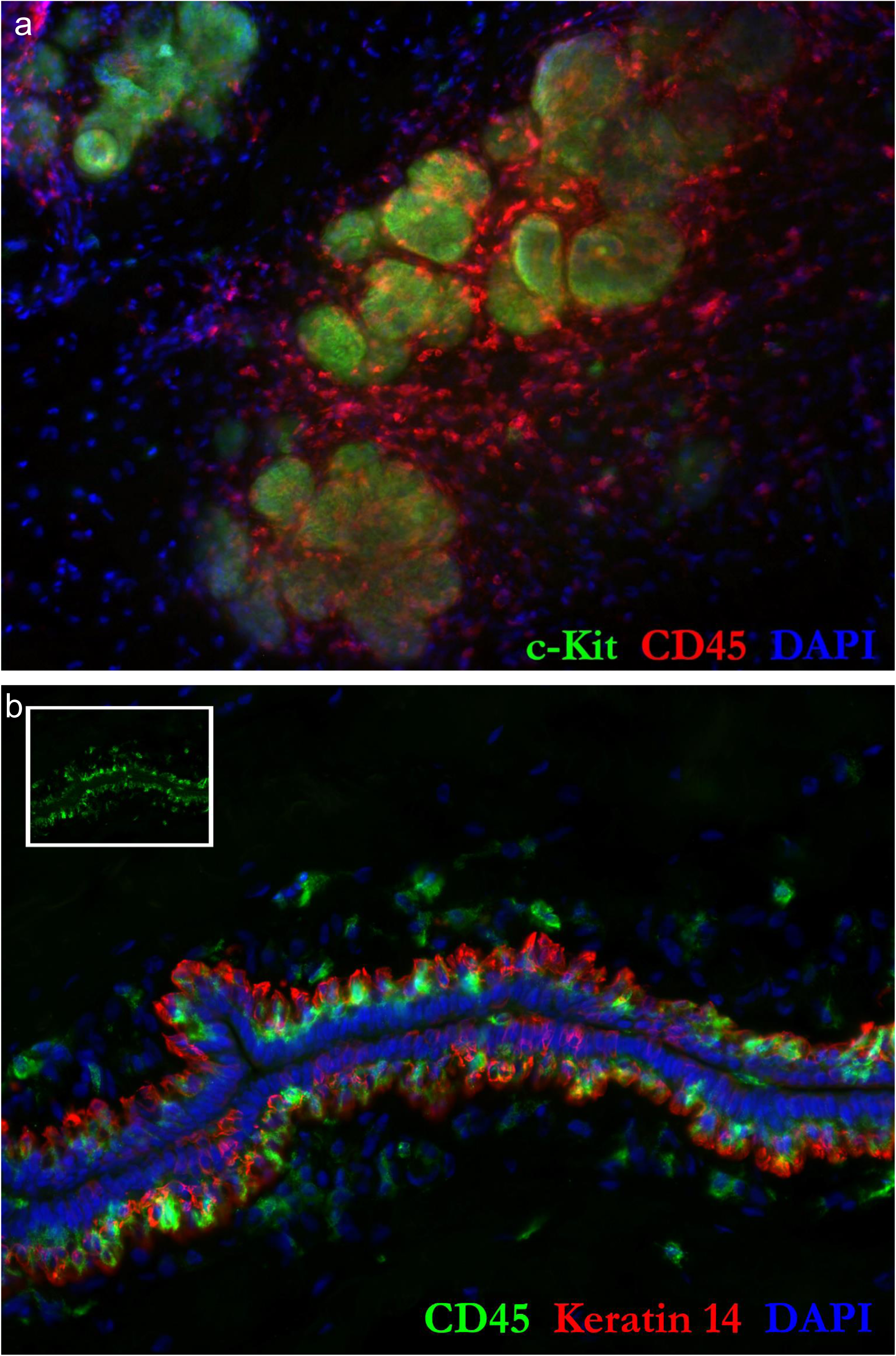
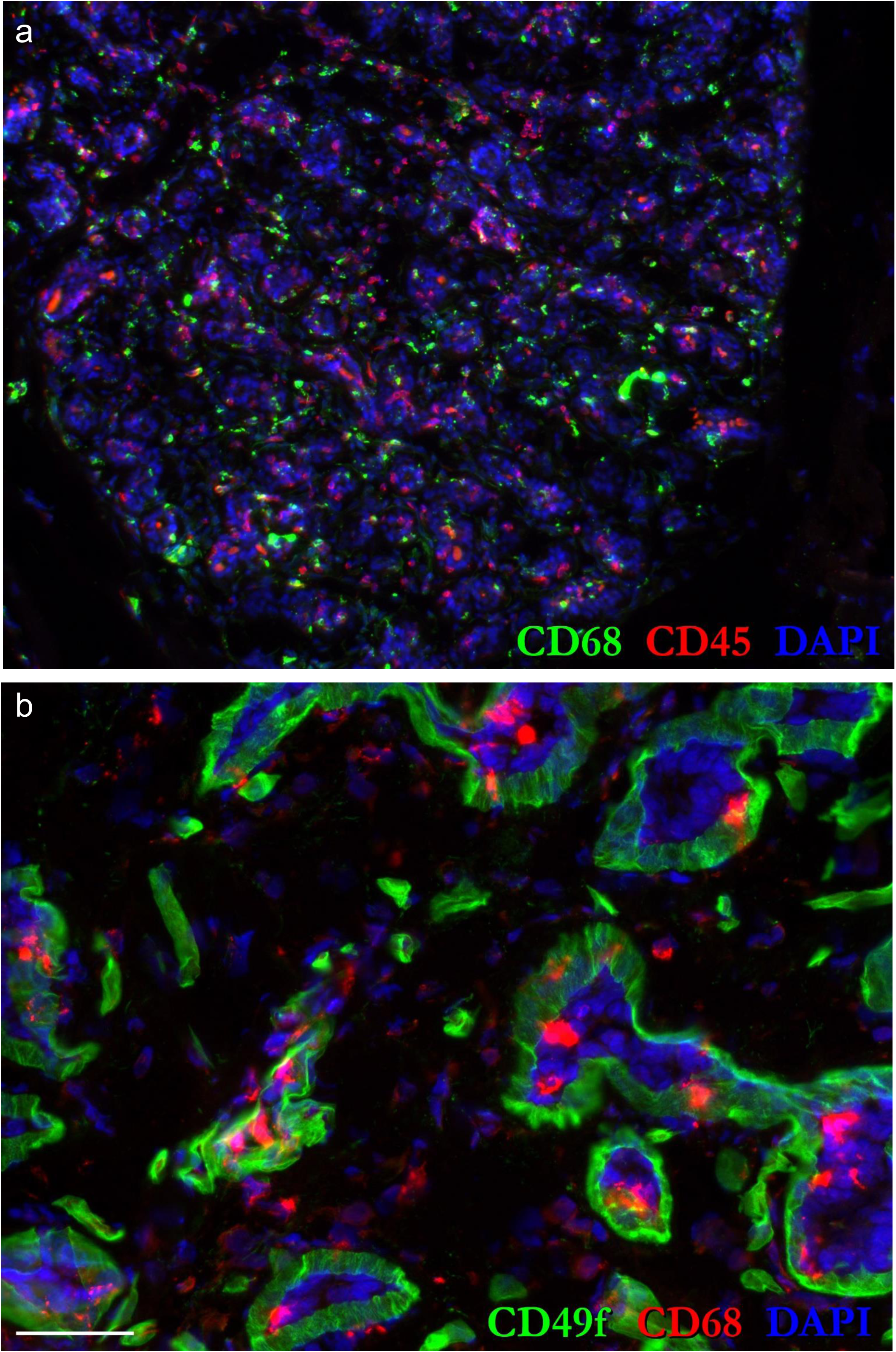
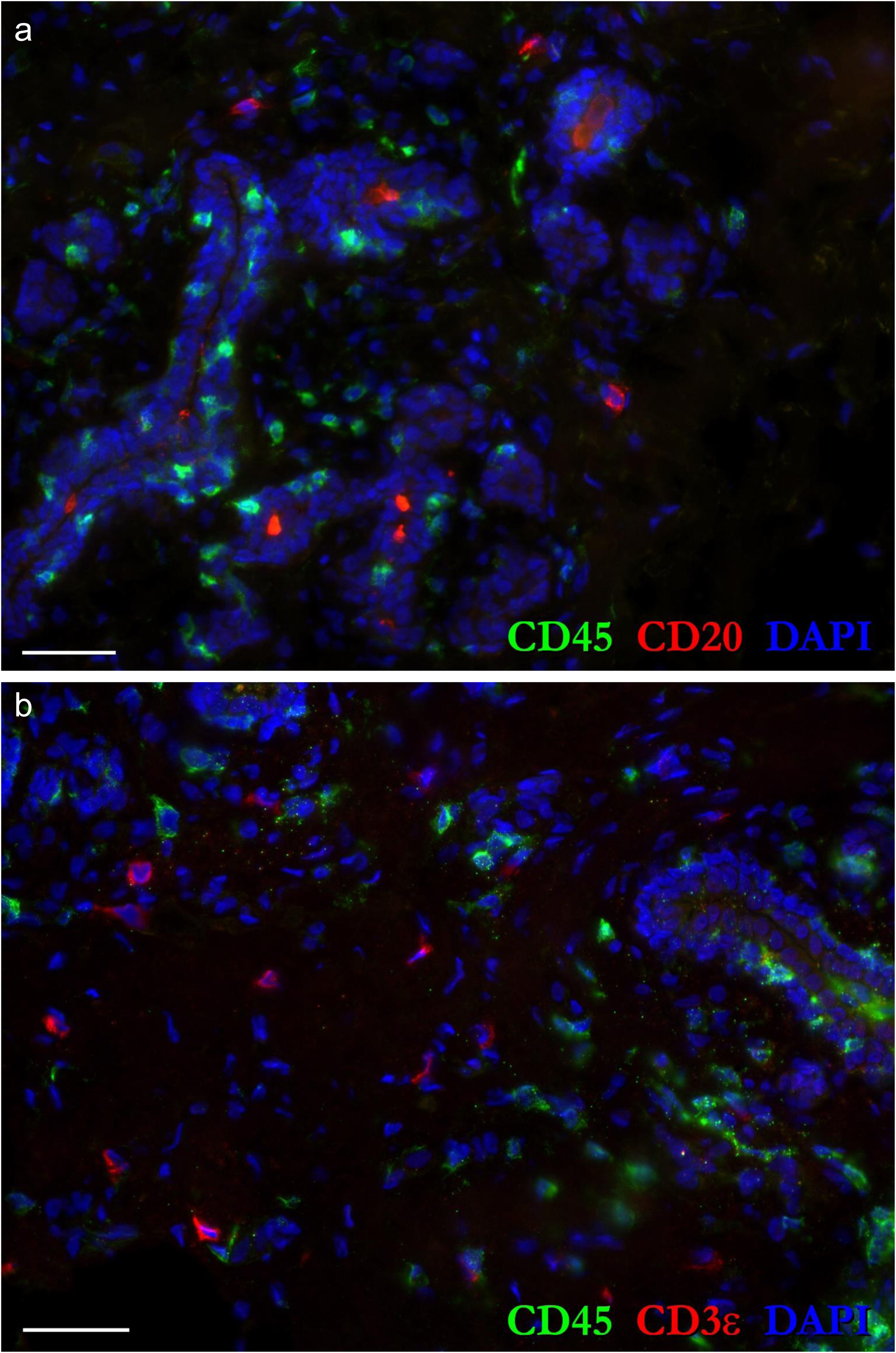
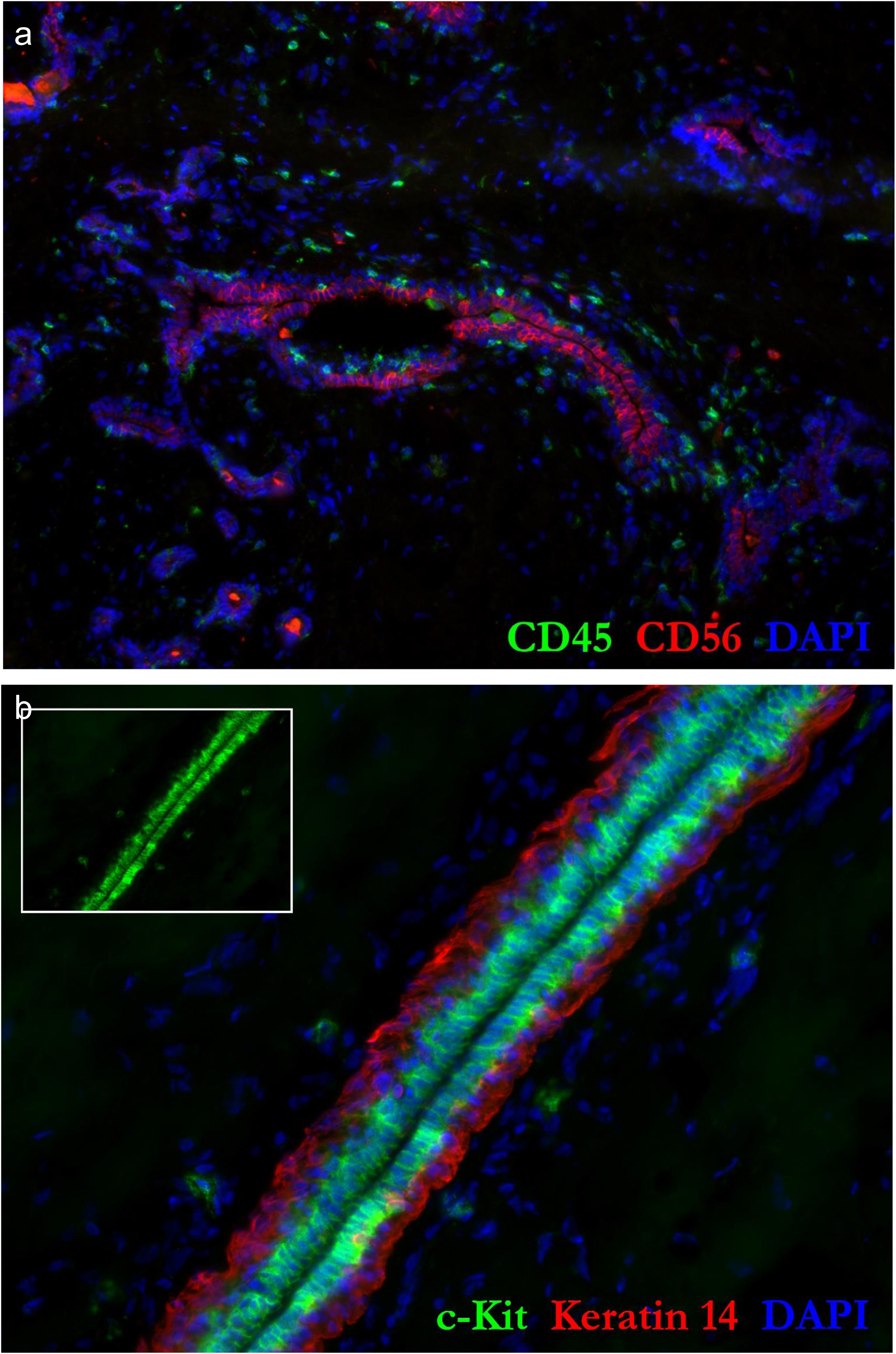
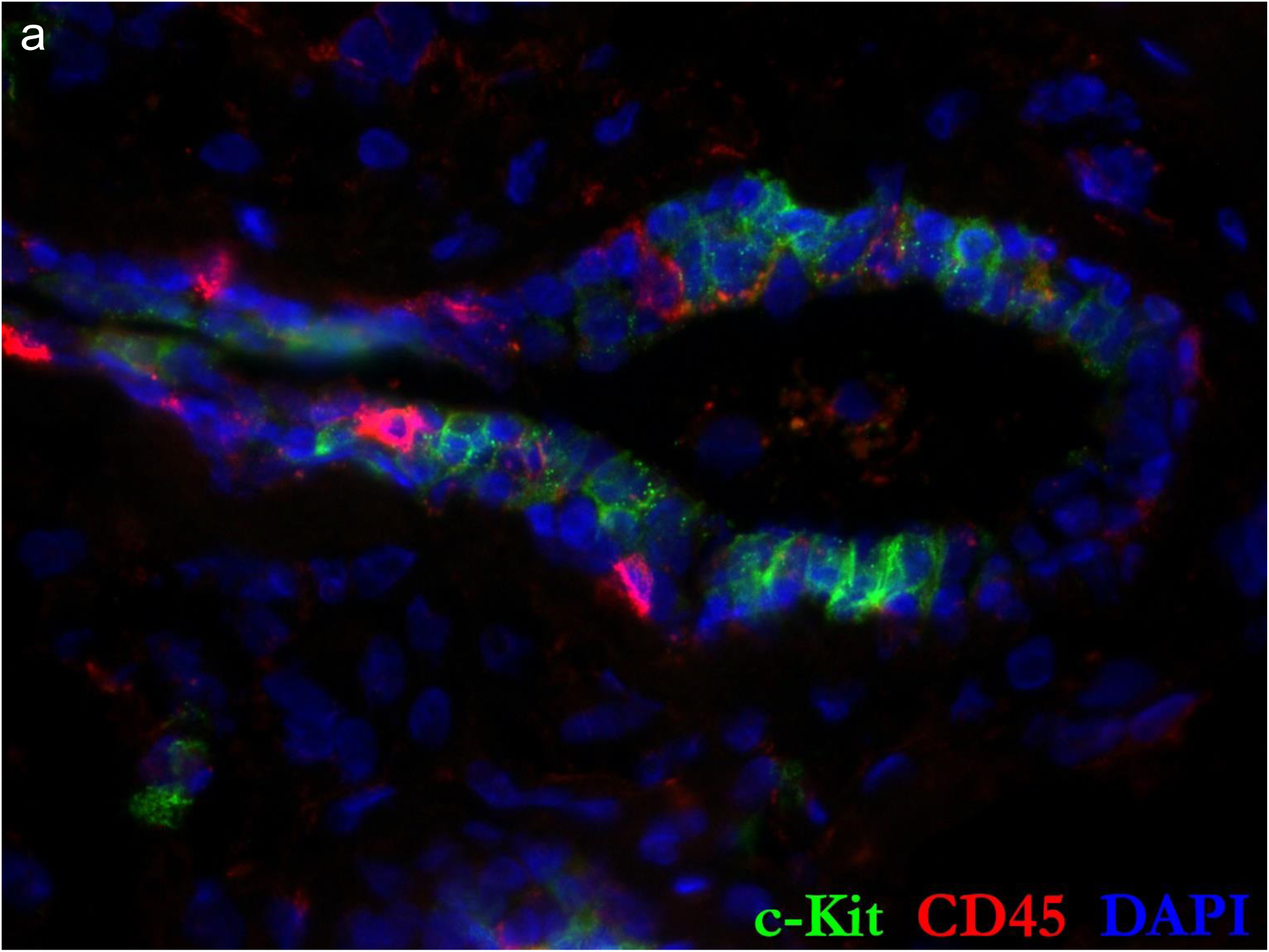
High resolution images of Figure 20– figure supplement 3a-i

**Figure 22 – figure supplement 1.**
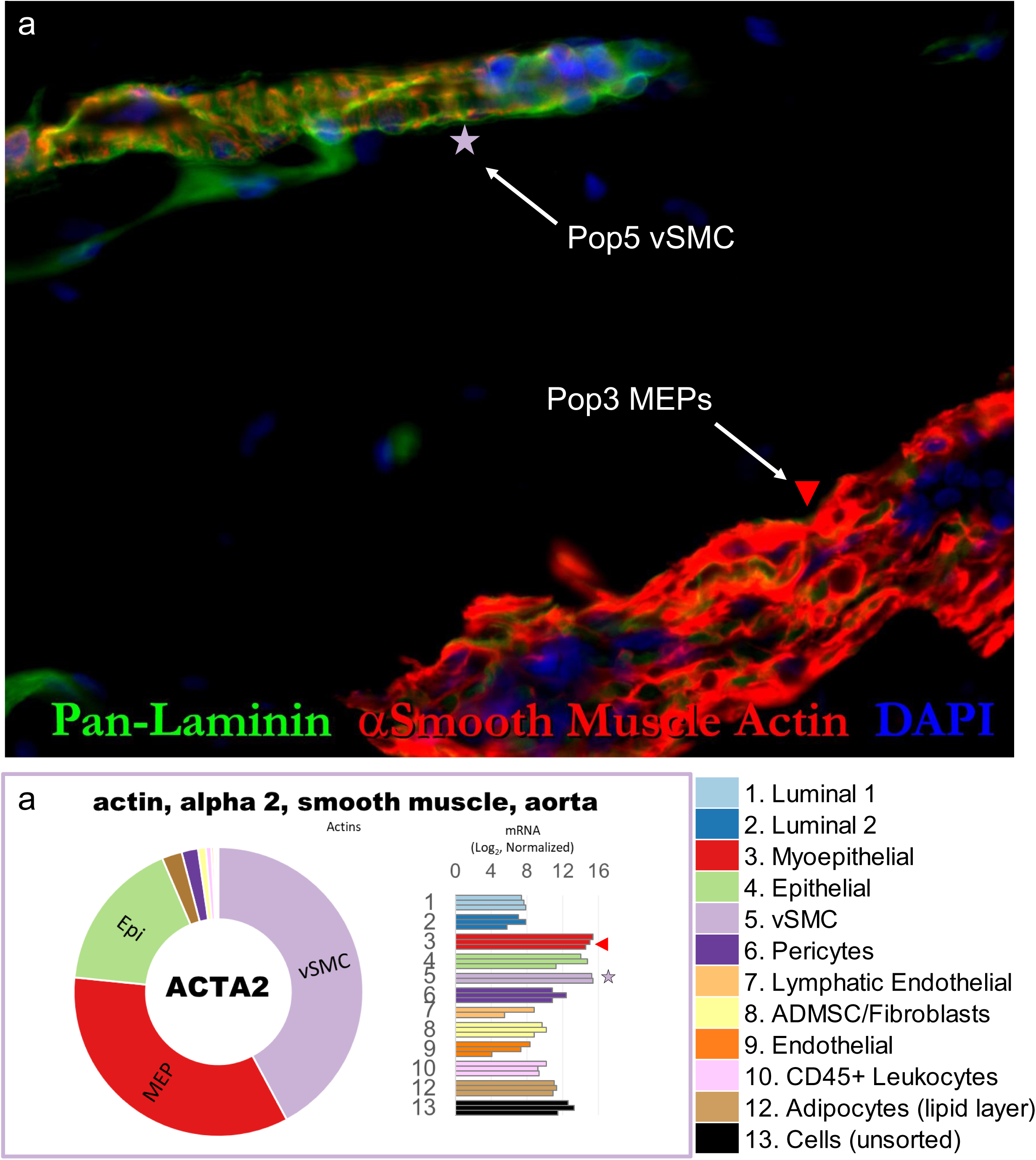
αSmooth muscle actin expression in human breast cells. High resolution image of the immunostained breast tissue provided in Fig. 22b. Below the image are the RNA-seq determined transcript levels of smooth muscle actin (ACTA2) measured within the uncultured FACS-purified cell types.

**Figure 22 – figure supplement 2.**
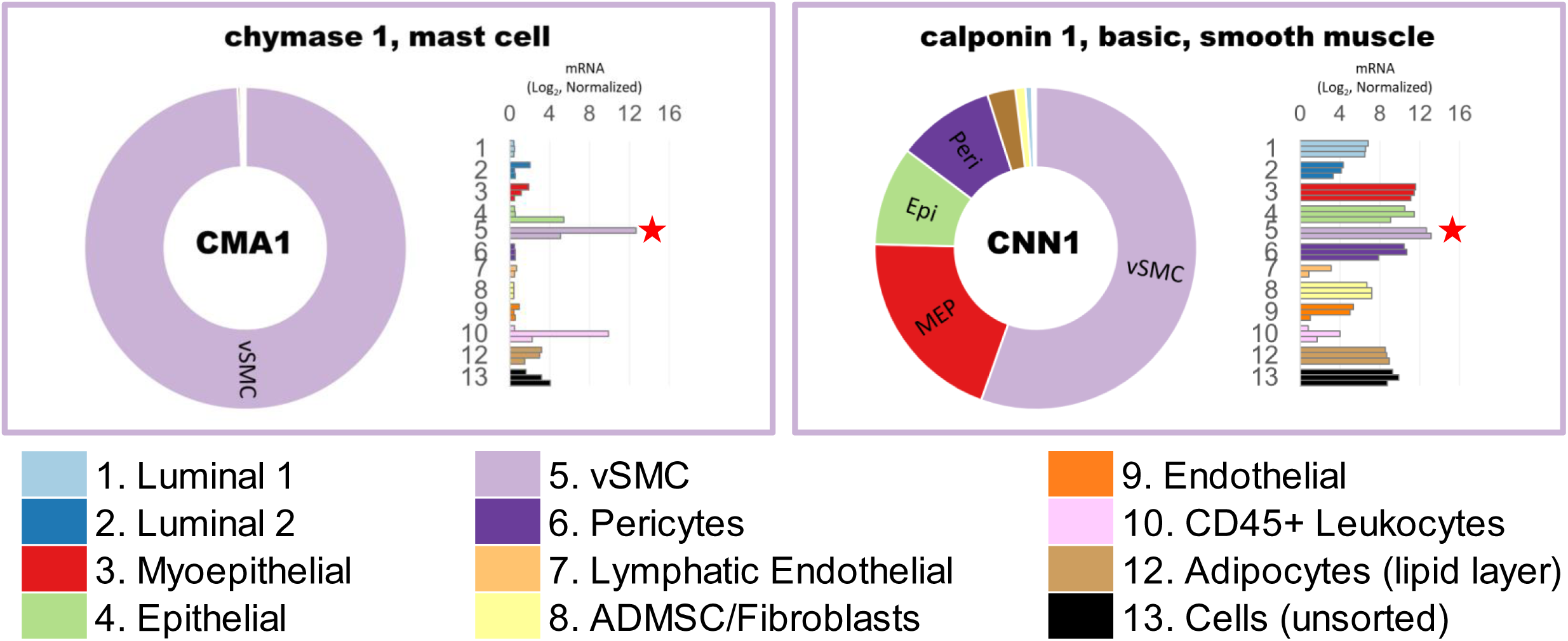
Transcript levels of vSMC-associated genes. **(a)** chymase 1 (CMA1) and **(b)** calponin1 (CNN1). Transcripts were measured in freshly sorted (uncultured) cell types by RNA-sequencing. Normalized mRNA values (rlog, DEseq2) are provided on log2 scale (bar graph of each biological replicate), as well as on a linear scale (donut graph of median value), each color-coded by cell type (vSMCs are light purple).

**Figure 22 – figure supplement 3.**
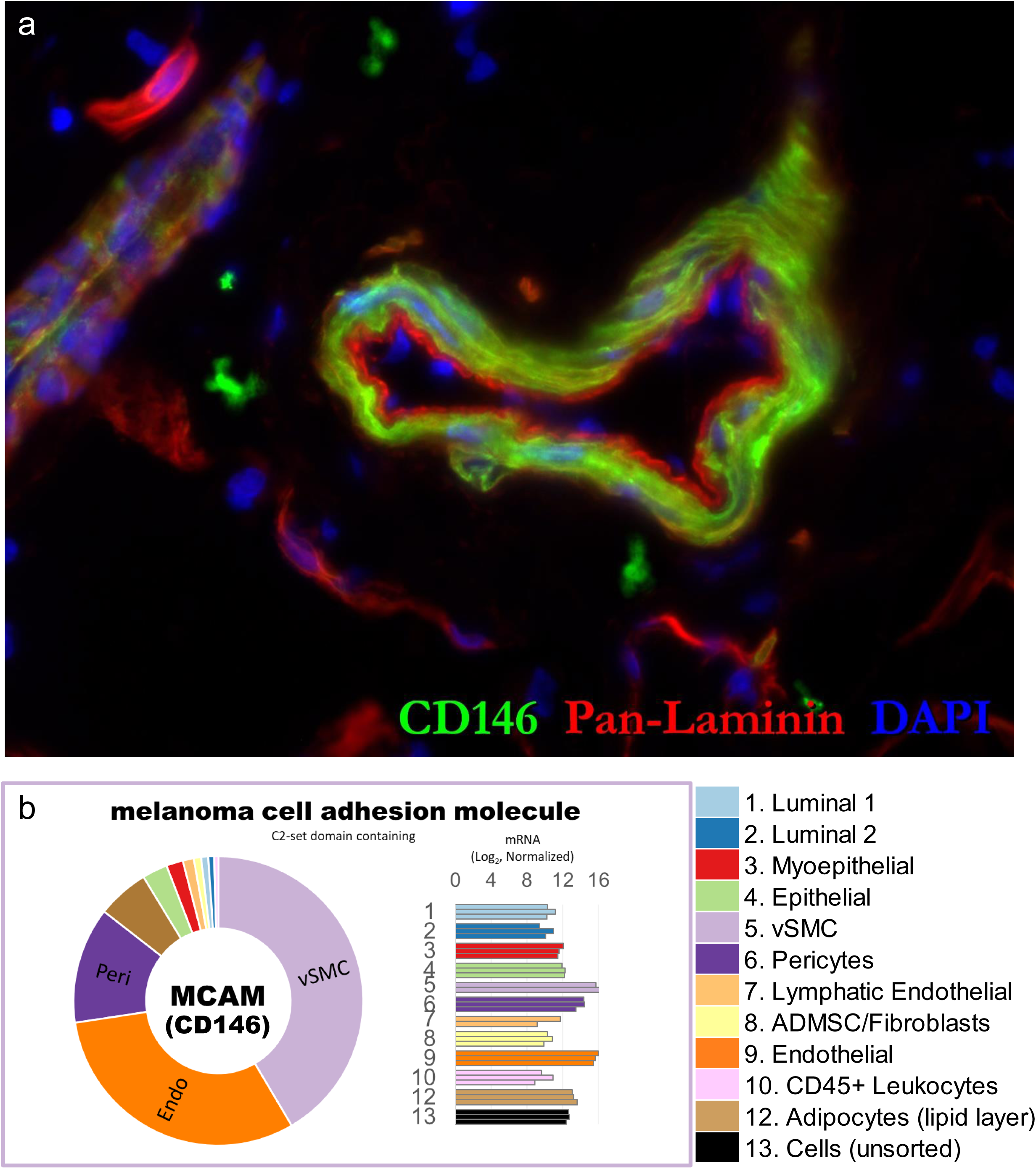
Melanoma cell adhesion molecule expression in human breast cells. High resolution image of the immunostained breast tissue provided in Fig. 22d. Below the image are the RNA-seq determined transcript levels of melanoma adhesion molecule (MCAM—also referred to as CD146), measured within each uncultured FACS-purified cell type.

**Figure 22 – figure supplement 4.**
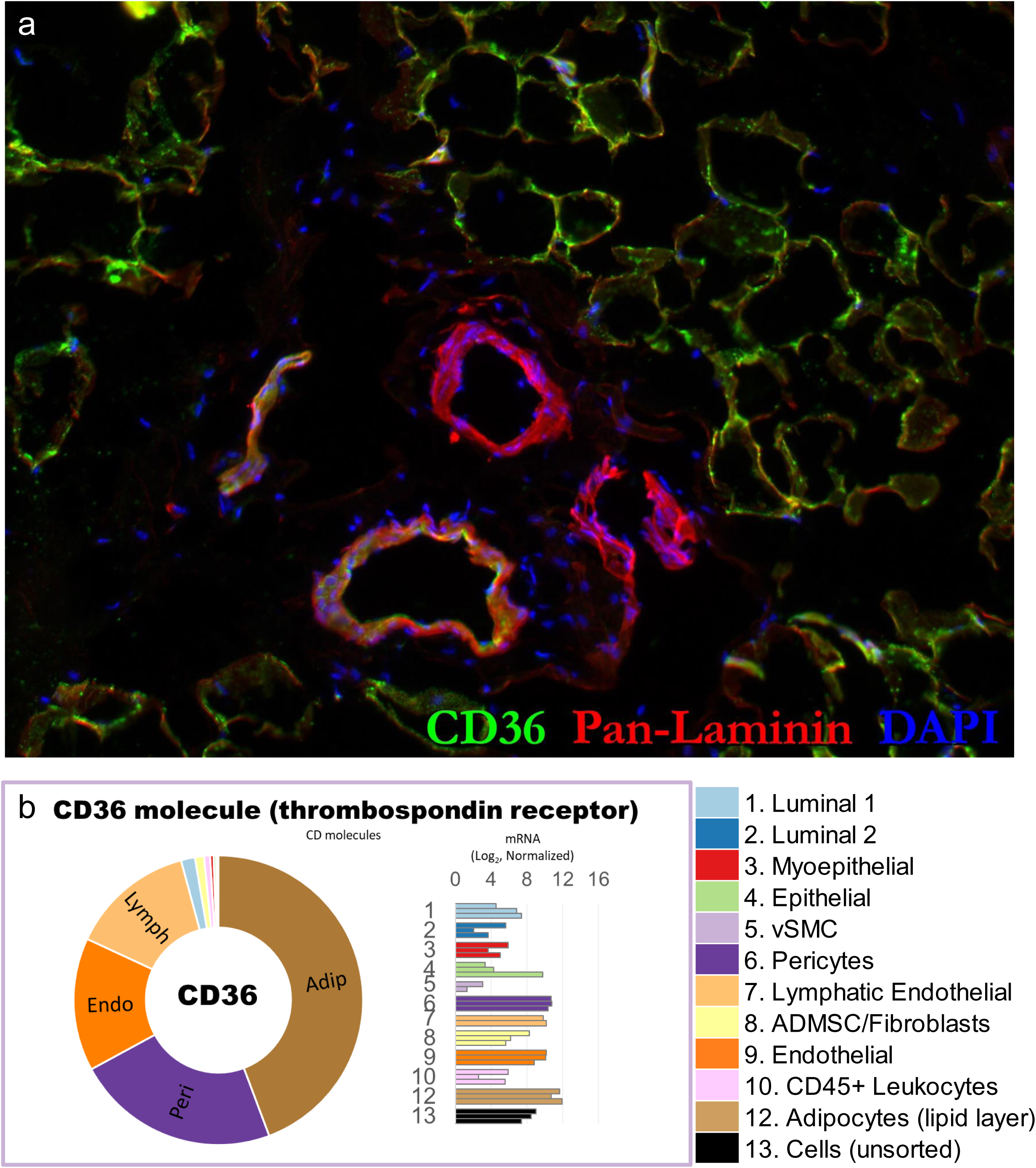
CD36 expression in human breast cells. High resolution image of the immunostained breast tissue provided in Fig. 22e. Below the image are the RNA-seq determined transcript levels of CD36, measured within each uncultured FACS-purified cell type.

**Figure 22 – figure supplement 5.**
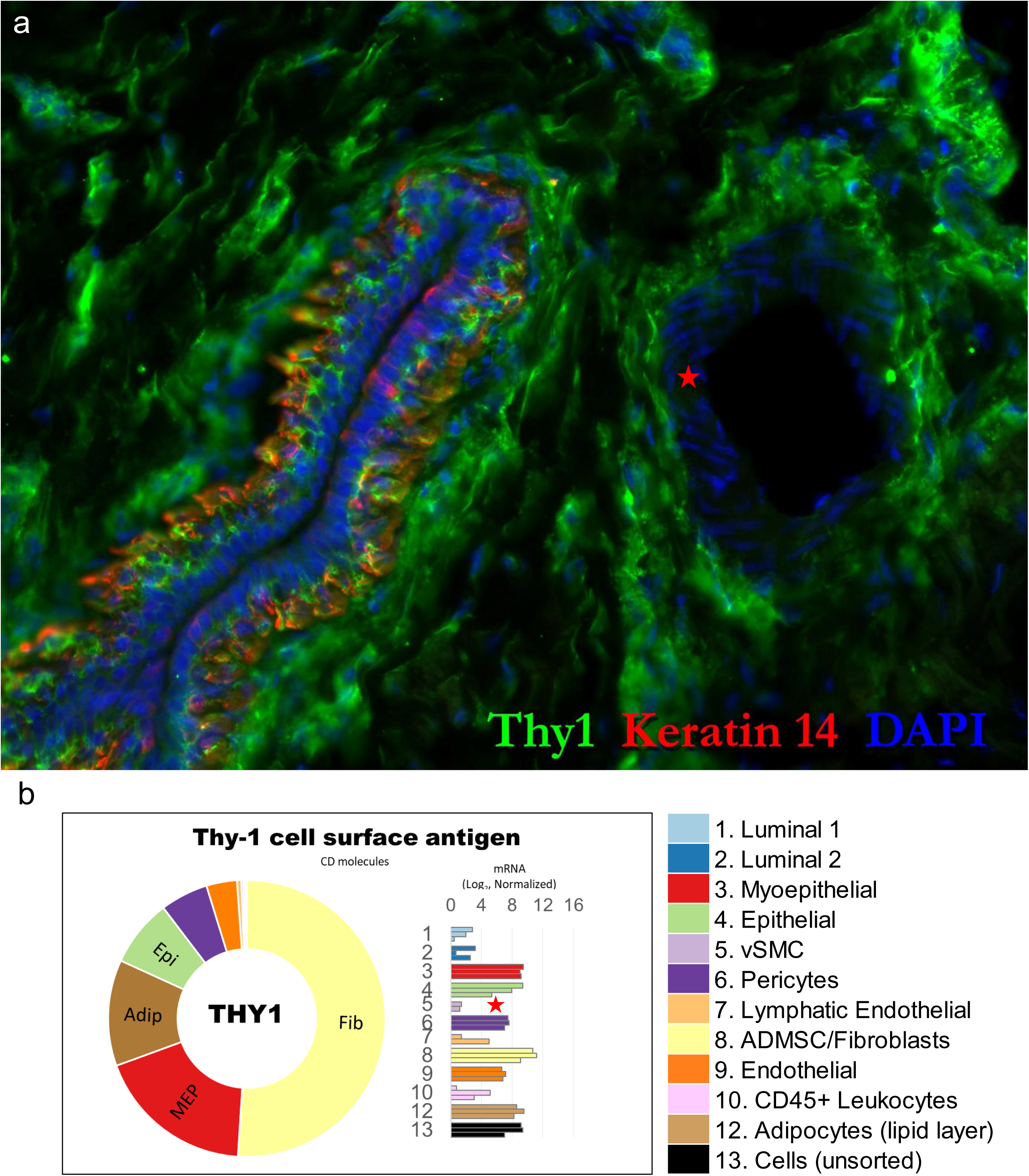
Thy1 is differentially expressed between pericytes and vascular smooth muscle cells (vSMCs). **(a)** A key marker in the FACS cell isolation strategy is Thy1, as it is used to identify and isolate Pop6 pericytes and myoepithelial cells. As is predicted by this gating scheme, we indeed found Thy1 was differentially expressed between Pop6 pericytes (isolated from the Thy1^Pos^ fraction, shown in dark purple) and Pop5 smooth muscle cells (isolated from the Thy1^Neg^ fraction, shown in light purple). **(b)** High resolution image of normal breast tissue immunostained for Thy1 and Keratin 14. The lack of Thy1 staining within vSMCs (identified by their hallmark cross-hatched nuclei, ‘red star’) is consistent with the lack of Thy1 staining by Pop5 (in FACS) and lack of Thy1 gene expression in FACS-purified cells shown in a.

**Figure 23 – figure supplement 1.**
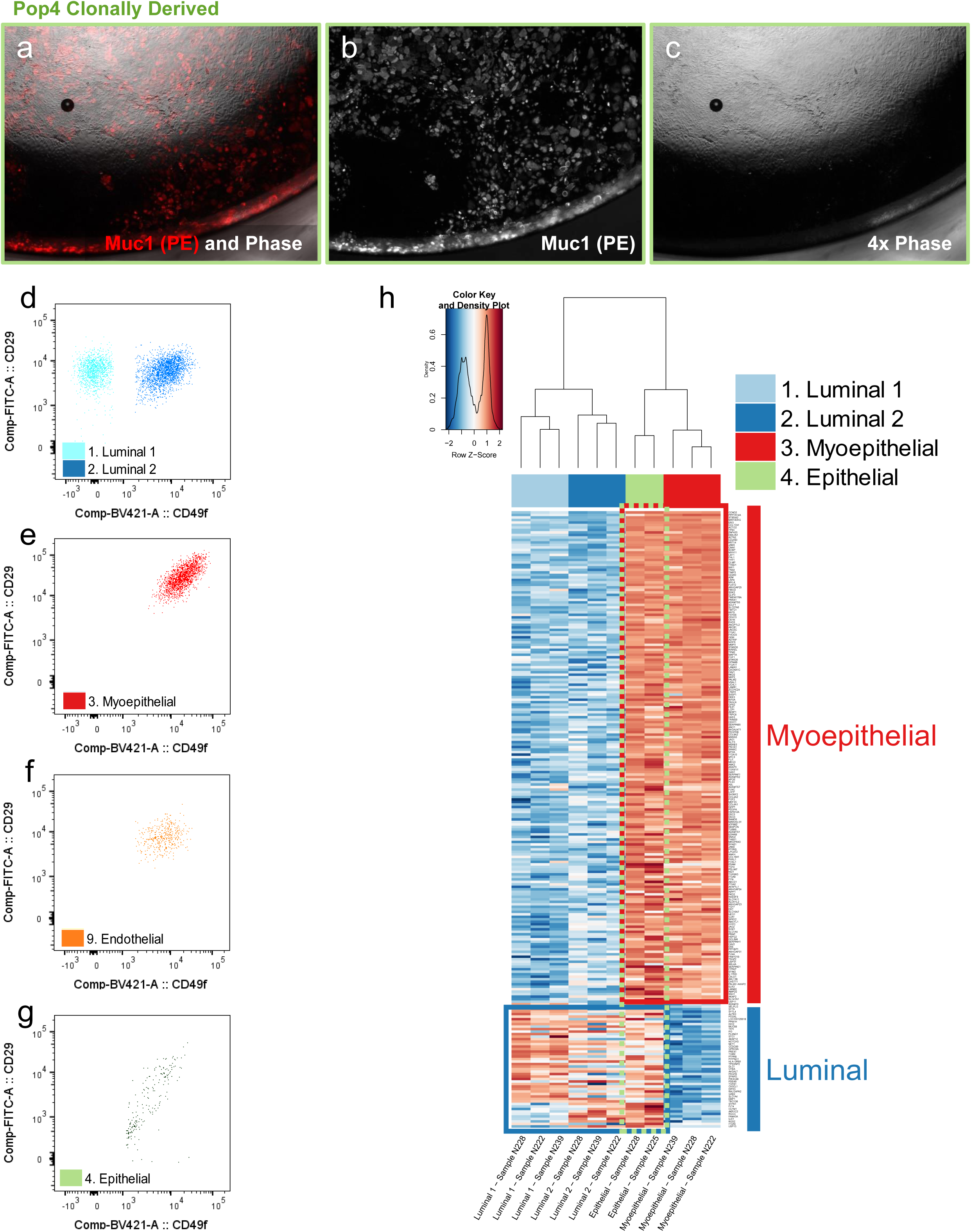
The atypical phenotype of Pop4 epithelial cells. **(a-c)** The clonal heterogeneous outgrowth of a single Pop4 cell, seeded by FACS into a 96-well culture dish; the **(a)** Phase/IF merged imaged produced by overlapping images of **(b)** Muc1-PE fluorescent immunostaining and **(c)** phase-contrast microscopy. **(d-g)** Individual FACS scatterplots of those used in the overlay shown in Fig. 23g, showing **(d)** Pop1 and Pop2 luminal cells, **(e)** Pop3 myoepithelial cells, **(f)** Pop9 endothelial cells, and **(g)** Pop4 epithelial cells, which display a unique CD29 staining profile. **(h)** Hierarchical clustering of RNA-seq measured transcripts from uncultured luminal epithelial cell types. Gene clusters of Pop1&2 (light and dark blue), Myoepithelial cells (red), and Pop4 epithelial cells (green) demonstrate the similar—but distinct—gene expression between Pop4 (green dashed outline) and myoepithelial cells (red outline), as there is a set of genes expressed by Pop4 that diverges and more closely resembles that of luminal epithelial cells (blue outline).

